# Integrative structural biology of a giant marine virus reveals architectural and functional complexity beyond icosahedral symmetry

**DOI:** 10.64898/2026.11.16.766480

**Authors:** Sara Otaegi-Ugartemendia, Roberto Marabini, Pablo Herrera-Nieto, Gabriela N. Condezo, Lars Mühlberg, Haina Wang, Joost Snijder, Rosa Viner, Lucía Ortiz Loo, Fan Liu, Ruth-Anne Sandaa, Carmen San Martín

## Abstract

Knowledge of giant virus diversity has expanded rapidly; however, isolated representatives remain scarce, and few have been examined at high resolution. Here, we characterized haptophyte-infecting *Prymnesium kappa* virus RF02 (PkV-RF02) using integrative structural biology. The ∼3 Å resolution structure obtained by block-based cryo-electron microscopy reconstruction shows a *pseudo*T=219 icosahedral shell built from over 20 different proteins and approximately 10,000 polypeptides, a multi-layered stabilization network reinforced by 20,000 disulphide bridges, two penton variants evenly distributed in the capsid, and a protein channel crossing the internal membrane. Cross-linking mass spectrometry defines the intricate capsid-membrane-core interaction network, which includes a second major capsid protein, a third penton variant, and multiple biochemical functions packaged within the virion. Phospho- and glycoproteomics reveal widespread post-translational modifications, with approximately 50% of proteins phosphorylated and uncommon methyl-N-acetylated viosamine glycans. This study provides a comprehensive structural and interaction map that highlights organizational and functional innovations underlying giant virus assembly.

## Main

Eukaryotic viruses with capsids larger than 150 nm are denoted as ‘giant viruses’^1^ and included in the nucleocytoplasmic large DNA virus group (*Nucleocytoviricota* phylum). Giant viruses are relevant not only in human or animal health (poxviruses, African swine fever virus (ASFV), iridoviruses), but also in the whole ecological balance on Earth. Many of them infect unicellular algae and protozoa, significantly affecting nutrient and energy cycles^2^. Detection of giant viruses in environmental surveys has significantly accelerated, with reported genomes now reaching the thousands^2,3^. Yet only a few have been actually isolated, and fewer studied at high resolution.

Giant viruses belong to the *Varidnaviria* realm, whose main hallmark is the major capsid protein (MCP) fold as a double jelly roll (DJR) perpendicular to the capsid surface^4^. For icosahedral giant viruses, reported triangulation numbers range between T = 147 and T = 499^5,6^, although *Acanthamoeba polyphaga* mimivirus (APMV) may reach the T = 972-1,200 range^7^. The large virions present challenges for high resolution cryo-electron microscopy (cryo-EM) data processing, such as the defocus variation within the same particle caused by exceeding the microscope depth of field, intrinsic capsid flexibility, and other deviations from icosahedral symmetry^1^. Despite the development of block-based reconstruction methods that help to overcome these limitations^8^, only one giant virus has been solved at resolution better than 4 Å (the limit at which polypeptide chains can be traced with confidence in cryo-EM maps)^9,10^, and few more in the 4-5 Å resolution range^11–14^. Most of these studies provide incomplete structures, due to limited resolution for vertex or minor capsid proteins (mCPs), or components deviating from the icosahedral symmetry. Block-based reconstruction methods also help unmask asymmetric capsid components as they do not enforce overall icosahedral symmetry. In this way, six different MCP variants, a singular vertex, and mCPs that break the icosahedral symmetry, were localized in the T = 169 *Paramecium bursaria* chlorella virus 1 (PBCV-1) capsid^9,10^.

Beyond these departures from symmetry, many components of giant virus particles, such as the inner lipid membrane, the genome and its associated proteins in the viral core, do not follow the symmetry of the capsid and may even be organized in a different way in each particle, precluding their observation by single particle averaging. Therefore, cryo-EM analyses cannot describe core component interactions or asymmetric capsid components happening in low copy numbers. Cross-linking mass spectrometry (XL-MS), however, can detect protein-protein interactions (PPIs) happening between both ordered and disordered virion components. XL-MS has been used to extract molecular proximity information in vaccinia^15^, decipher spatial distribution in human cytomegalovirus and Melbournevirus particles^16,17^, and facilitate assignment of protein identity to mCP cryo-EM densities^11,17^.

Here, we present the structural characterization of *Prymnesium kappa* virus RF02 (PkV-RF02, *Tethysvirus bergenense* under the current ICTV nomenclature). PkV-RF02 is a giant icosahedral virus infecting haptophyte algae, with a 583 kbp linear double-stranded dsDNA genome from which 635 open reading frames (ORF) were predicted^18,19^. Since haptophytes contribute 30-50% of primary productions in the ocean^20^, their viruses have great ecological importance. PkV-RF02, taxonomically closer to the amoeba-infecting APMV than to the well-characterized algavirus PBCV-1, is included in the *Imitervirales* order^21^. We have previously shown that at least 136 different proteins are present in purified PkV-RF02 virions, of which 66% have unknown functions, and only the MCP has been unequivocally identified as capsid component^19^. We now use an integrative approach combining block-based cryo-EM reconstruction, XL-MS, phosphoproteomics, and glycoproteomics, to show that PkV-RF02 virions are highly organized biological machines carrying novel architectural features, functional enzymes, and abundant post-translational modifications (PTMs), likely crucial for infection and environmental fitness.

## Results

### Overall PkV-RF02 capsid structure

Cryo-EM showed that PkV-RF02 has an icosahedral capsid of diameter between 185 nm (facet-to-facet) and 210 nm (vertex-to-vertex), enclosing an internal membrane (**Extended Data Fig. 1a**). In some particles, the viral genome appears to adopt a toroidal arrangement (**Extended Data Fig. 1b**). A 4.7 Å resolution icosahedrally averaged map showed that the PkV-RF02 capsid has a *pseudo*T = 219 triangulation number (**Extended Data Fig. 1c-d, Supplementary Table 1**), previously observed only for *Phaeocystis pouchetii* virus at 30 Å resolution^22^. Capsomers are arranged in 12 pentasymmetrons and 20 trisymmetrons^23^ (**Extended Data Fig. 1c**). The icosahedral asymmetric unit (AU) contains 109 MCP monomers (36 trimers, plus a single monomer for the trimer centred at the icosahedral 3-fold axis), and a monomer of the vertex protein (penton) (**Extended Data Fig. 1e**). The internal membrane is fastened to the 8.5 nm thick capsid shell by contacts near the 3-fold icosahedral axis (**Extended Data Fig. 1f**). In the averaged map, the genome produces a pattern of concentric shells separated by approximately 2.8 nm (**Extended Data Fig. 1f**), slightly more than the 2.5 nm described for toroidal naked DNA^24^.

Refining and reconstructing the AU in ten overlapping blocks^8^ yielded ∼3 Å resolution maps (**Extended Data Fig. 2, Supplementary Fig. 1, Supplementary Table 2**) which allowed us to solve the structure of the pseudo-hexameric and pentameric capsomers, as well as mCPs on the inner and outer surfaces of the capsid (**Extended Data Table 1, Supplementary Tables 3-4, Supplementary File 1, Supplementary Movie 1**). Using mass-spectrometry, we previously identified two MCP candidates present in the virion^19^, hereafter referred to as MCP variant 1 (MCPv1, ORF313) and MCP variant 2 (MCPv2, ORF450), with a protein sequence identity of 23.34% (**Supplementary Fig. 2**). MCPv1 was the most abundant protein in the virion, while MCPv2 was approximately 100 times less abundant^19^. MCPv1, most similar to *Emiliana huxleyi* virus 201 (EhV-201) MCP (PDB: 8rbt)^25^ (**Supplementary File 2a-b**), was traced in all the 6-fold coordinated capsomer positions (**Extended Fig. 1e, Supplementary Table 4**) and shows the DJR fold conserved in the *Varidnaviria* realm (**Fig. 1a**). The MCPv1 N-terminal region and the loops located above the DJR extend to clasp neighbouring monomers to form the pseudo-hexagonal trimer (**Fig. 1b**), stabilizing it by intra- and inter-chain π-π contacts, non-covalent interactions between aromatic residues^26^ (**Fig. 1c, Supplementary Table 5**). Extra density protruding from serine and threonine residues (Ser357, Ser366, Thr380, Ser382) indicates possible O-linked glycosylations on the capsid surface (**Fig. 1d, Extended Data Table 2**). No density corresponding to MCPv2 was detected in any position of the AU and thus, the protein could not be modelled (**Supplementary Fig. 2**).

**Figure 1.**
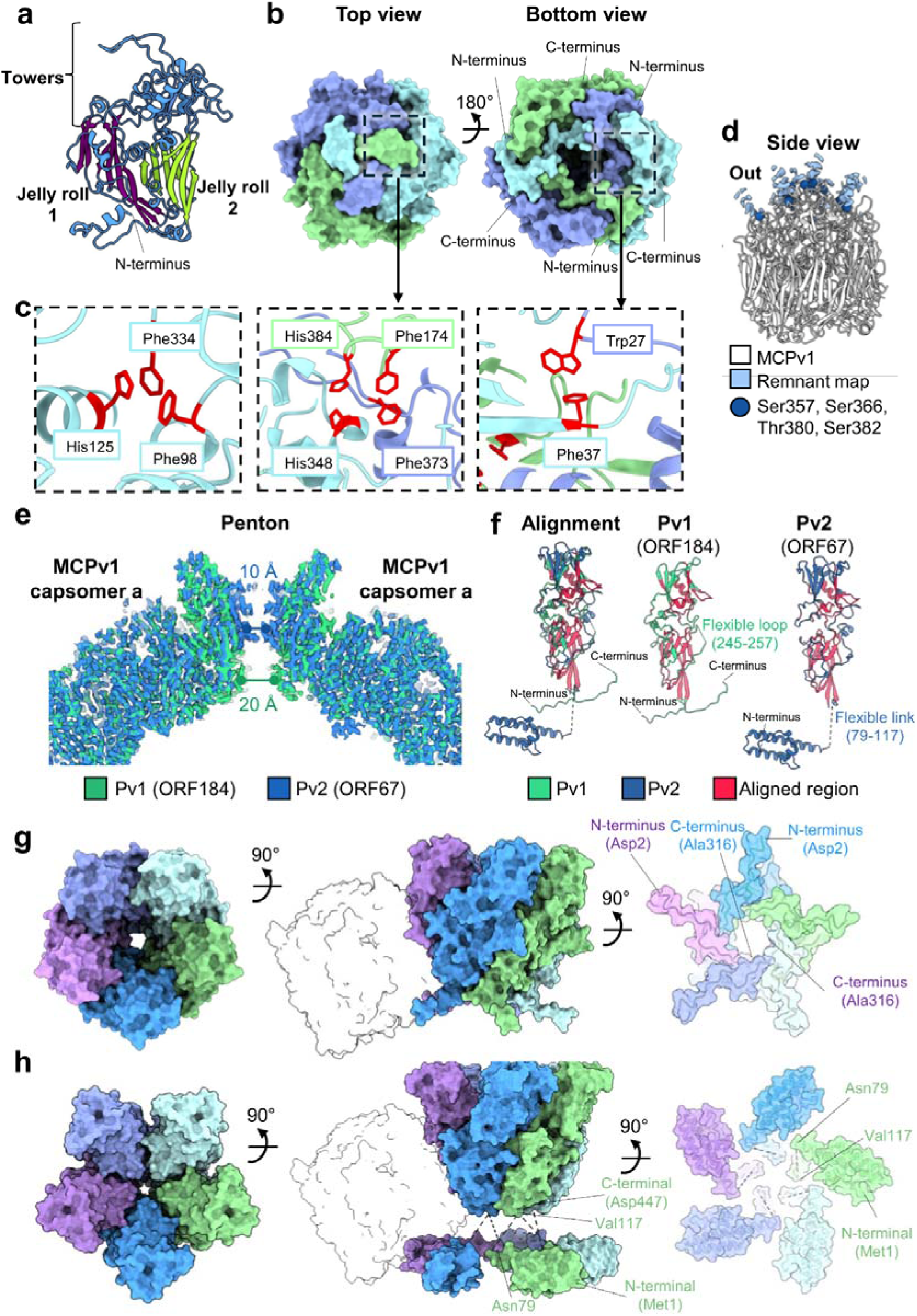
Structure of main capsid components. (**a**) Molecular model of the MCPv1 monomer, with β-strands in maroon for jelly roll 1 and in green for jelly roll 2. (**b**) MCP trimer capsomer, from the top as viewed from outside the capsid (left) and bottom (right). (**c**) Intra- and inter-chain π-π interactions in MCPv1. (**d**) MCP remnant densities extending from serine and threonine residues. (**e**) Overlap of the two solved maps for the 5-fold symmetry block. The view is a section across the capsid. Pore diameters measured at their narrowest point are indicated for each penton variant. (**f**) Structural alignment of molecular models for penton variants: Pv1 (ORF184) and Pv2 (ORF67). (**g**) Pentameric structure of Pv1, from the top, side and bottom. The N-terminal region clasps the adjacent MCPv1 capsomer (shown in silhouette). (**h**) Pentameric structure of Pv2, from the top, side and bottom. The N-terminal region forms an α-helical domain that clasps neighbouring MCP trimers (silhouette).

### Two penton variants evenly distributed in the capsid

3D classification of the block centred at the 5-fold axis yielded two high-resolution maps that were different exclusively in the penton protein (**Fig. 1e, Extended Data Fig. 2**). Approximately 40% of the block sub-particles contributed to each of the maps, while the rest yielded low resolution maps (**Supplementary Table 2**). Two different penton proteins were traced: penton variant 1 (Pv1, ORF184) and penton variant 2 (Pv2, ORF67) (**Fig. 1f**). Both show a well-structured single jelly roll (SJR) at the base and a seven-strand β-sandwich at the top (**Fig. 1f**). When assembled as pentamers, both proteins feature a central pore, 20 Å in diameter at its narrower point for Pv1, and 10 Å for Pv2 (**Fig. 1e, g-h**). Pv1 and Pv2 share 30.7% protein sequence identity. Pv1 has a shorter N-terminal region, which acts as a hook to the peripentonal capsomers (**Fig. 1f-g**). The larger N-terminal domain in Pv2 forms a coiled-coil structure connected via a flexible link (residues 79-117) that could not be traced, also clasping the peripentonal capsomers (**Fig. 1f, h**). The C-terminal domain of Pv1 stabilises the pentamer by interacting with adjacent monomers, similarly to the MCPv1 N-terminal arm (**Fig. 1g**), while the C-terminal domain of Pv2 remains confined to the main body of the pentamer (**Fig. 1h**). Two vertex types frequently occur in icosahedral capsids, particularly in tailed phages but also in varidnaviruses (PRD1, PBCV-1, APMV)^27^, always arranged as 11 identical vertices plus a singular one. Our observation of two equally populated penton classes represents a marked departure from this conserved vertex specialization, pointing to a previously unrecognized mode of vertex composition in icosahedral virus capsids. We assessed the possibility that each particle may preferentially incorporate one penton variant, or that a variant at one vertex influences the composition of neighbouring vertices, by comparing our experimental penton distribution against simulated assemblies (**Supplementary Note 1**). These analyses support a near stochastic mode of penton incorporation, with little evidence for preferential placement or local coordination.

### Multilayered capsid reinforcement by an exceptionally extensive disulphide network and minor coat proteins

Apart from the non-covalent interactions described above, MCPv1 trimers are further stabilized by covalent disulphide bonds between Cys174 in one monomer and Cys413 in the next (**Fig. 2a**). In the capsid, although inter-capsomer interactions are not energetically favoured (ΔG∼0, **Supplementary Table 6**), all possible MCP-MCP interfaces are stabilized by inter-capsomeric disulphide bonds (**Fig. 2b**). Together, MCPv1 intra- and inter-capsomer disulphide bridges create an extensive cross-link network of ∼20,000 covalent bonds across the capsid surface (**Supplementary Fig. 3a**). Further, at the vertices, Pv2 Cys293 forms a disulphide bond with Cys316 of the peripentonal MCPv1 capsomer, whereas the corresponding Pv1 residue (Cys145) does not, but proximity indicates that a disulphide bond could form with a minor conformational adjustment (**Fig. 2c, Extended Data Table 2, Supplementary Table 7**).

**Figure 2.**
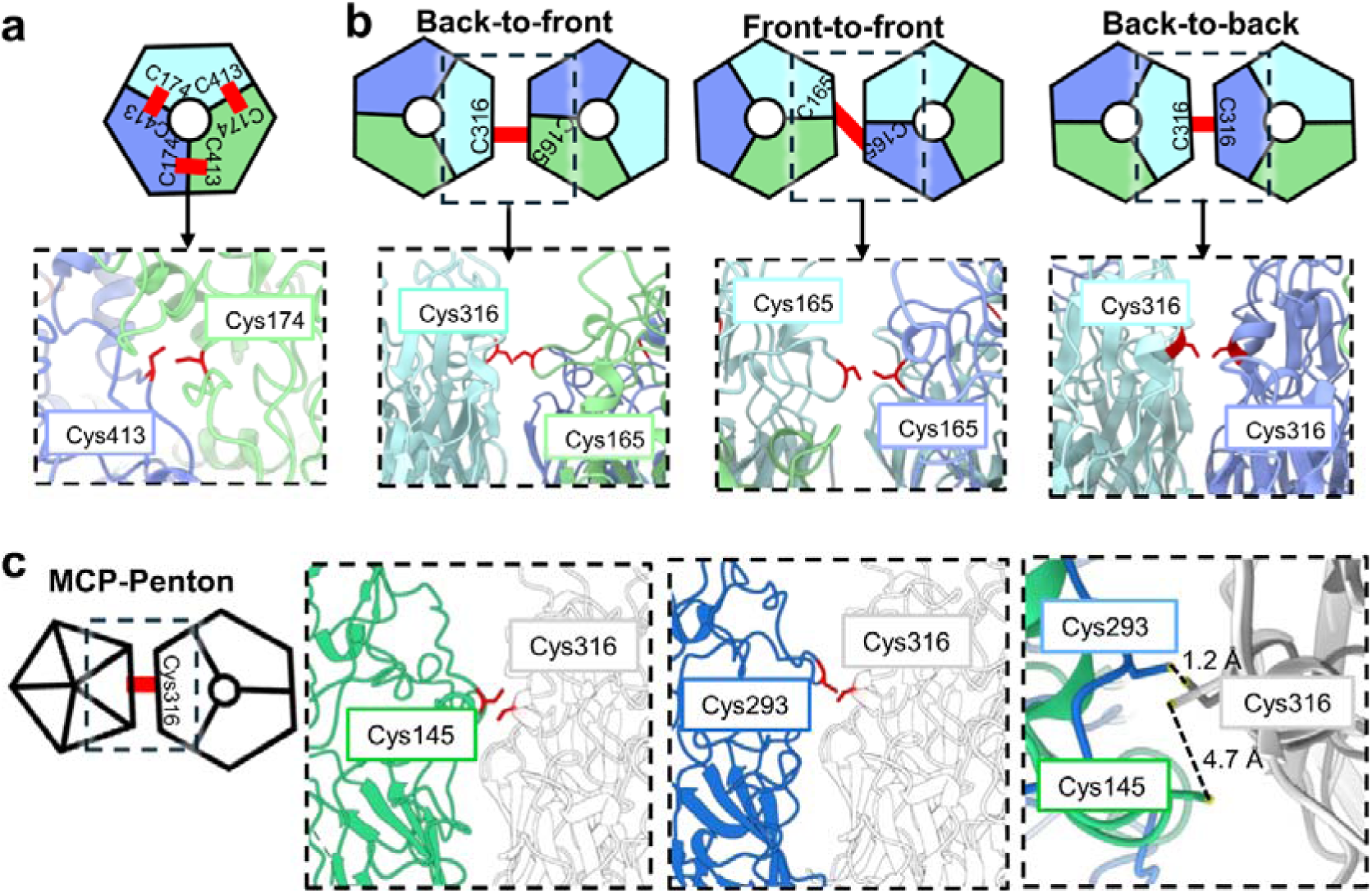
Disulphide bond stabilization of major capsid components. (**a**) Intra-capsomeric disulphide bond between MCPv1 Cys174 and Cys413. (**b**) Inter-capsomeric disulphide bonds in the three types of MCP capsomer interfaces. Residues involved in the described interactions are coloured in red in (a-b). (**c**) Pv1/Pv2 interaction with MCPv1 (grey) by cysteine residues. The two panels on the left display contacts of cysteines coloured in red. The panel on the right illustrates the interaction distances between these cysteines.

Besides trimeric and pentameric capsomers, we traced eighteen mCPs, six of them on the outer capsid surface (external mCPs) and twelve on the inner one (internal mCPs) (**Extended Data Fig. 3-4, Extended Data Table 1, Supplementary Table 4, Supplementary Movie 1**). Nine mCPs contribute to the pentasymmetron, four on the outer surface and five on the inner one (**Fig. 3a-c**). On the outer side, ORF76 connects the penton protein with the first two rings of MCP capsomers; ORF281 links the peripentonal ring with the next two, reaching the pentasymmetron edge; while ORF289 and ORF535 bridge the pentasymmetron with the trisymmetron (**Fig. 3a, Extended Data Fig. 3-4**). These proteins occupy the valleys between the MCP trimers, in some cases (ORF76, ORF281, and ORF535) obstructing inter-capsomer disulphide bonds. To compensate, they contain cysteine residues near MCPv1 cysteines, forming MCP-mCP disulphide bonds (**Fig. 3a**). These three proteins also present O-linked glycosylations (**Fig. 3a, Extended Data Table 2**). Remarkably, ORF289 molecules are deeply embedded within the capsid valleys, locked in by inter-capsomer disulphide bonds formed above them (**Fig. 3a**). On the inner side, ORF357 is located between capsomers a, b and c; and ORF404 links ORF214 (the zip protein according to the designation in PBCV-1^9^) to the second ring of capsomers in the pentasymmetron, as does the C-terminal region of the tape measure protein (TmP, ORF401). Finally, ORF481 protrudes towards the capsid lumen at the interface between the second and third ring of capsomers (**Fig. 3b**). This protein displays a globular fold unusual among the less structured cementing proteins (**Extended Data Fig. 4**). Structural homology searches revealed similarity to *Yersinia* (PDB: 8ovt) and human (PDB: 7ui6) glycosyltransferases (**Supplementary File 2e-f**). Internal pentasymmetron mCPs do not interact with the membrane (**Extended Data Fig. 1f**), unlike those in PBCV-1, which contact the lipid bilayer near the 5-fold axes^10^.

**Figure 3.**
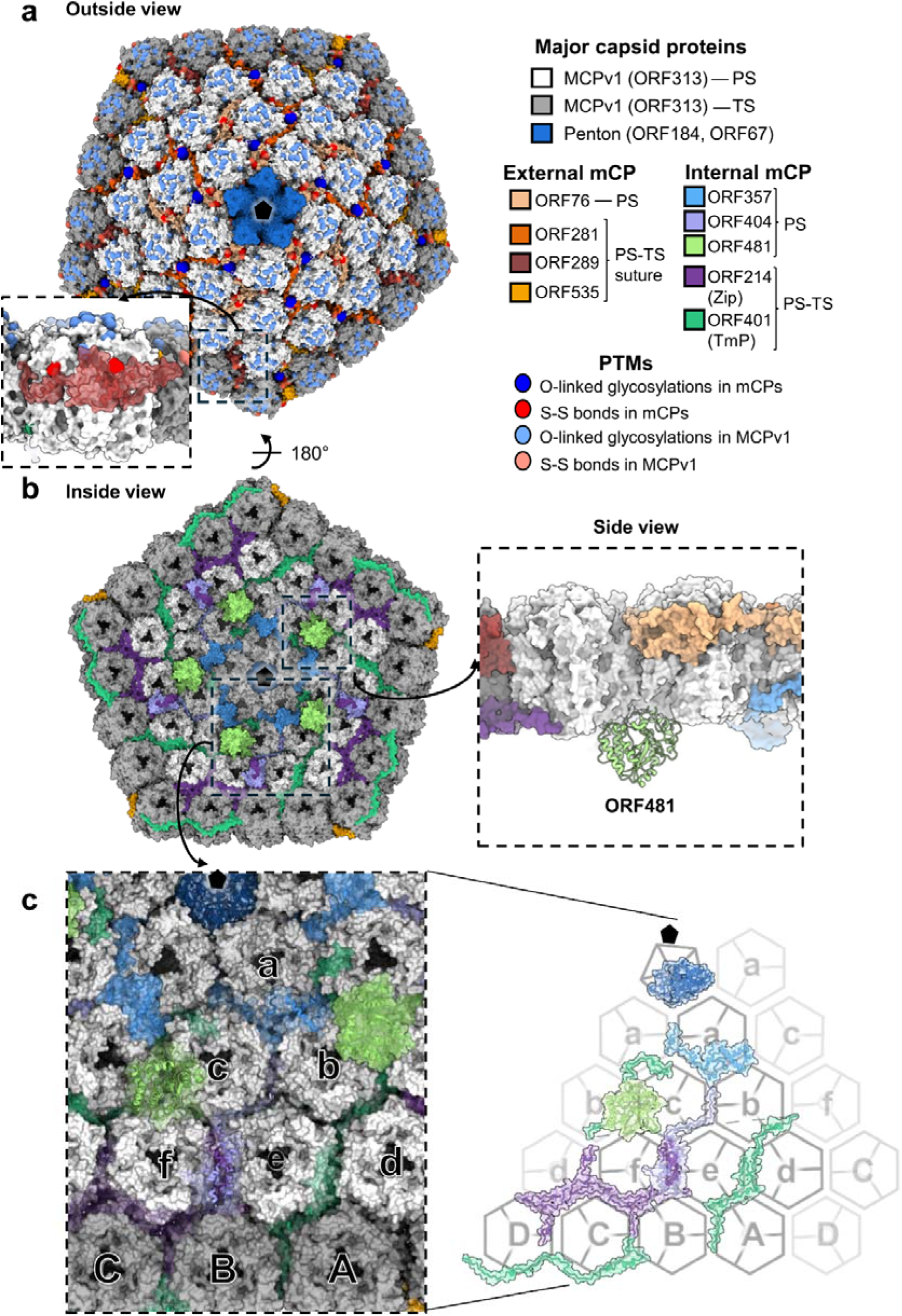
Pentasymmetron organization of PkV-RF02 virions. (**a**) Outside view of the pentasymmetron. Disulphide bridges and O-linked glycosylations on the MCP are shown in light red and blue, while those in the mCPs are depicted in dark red and blue. TS, trisymmetron. PS, pentasymmetron. The region enclosed by the dashed square is zoomed in to show ORF289 locked in place by disulphide bonds between MCPv1 capsomers (red). (**b**) Inside view of the pentasymmetron. The panel at the right shows a side view to appreciate the position of the putative glycosyl transferase ORF481, which is protruding towards the lumen of the virus particle. (**c**) Closer look at the internal mCP organization in the pentasymmetron. For clearer visualization, on the right the MCP trimers have been computationally removed to show the mCP network.

We traced thirteen mCPs in the trisymmetron, four on the outer surface and nine on the inner one (**Fig. 4a-b, Extended Data Fig. 3-4**). On the external surface, trisymmetron mCPs fill the valleys between MCP trimers in a specific pattern: ORF535 and ORF289 fill valleys at the trisymmetron-pentasymmetron boundary as indicated above, ORF287 is located on the valleys at trisymmetron edges, and ORF140 occupies the rest. Similary to ORF289 in the pentasymmetron, both ORF140 and ORF287 are locked in by inter-capsomer disulphide bonds formed above them (**Fig. 4c**). The organization of internal trisymmetron mCPs can be described as two groups: proteins aligned parallel to the trisymmetron suture, and proteins extending from the trisymmetron periphery towards the 3-fold axis. The zip protein reinforces the edge of the trisymmetron, while two copies of the elongated TmP run in antiparallel fashion from vertex to vertex at each trisymmetron boundary, one capsomer away from the suture defined by the zip protein (**Fig. 4b, Extended Data Fig. 3-4**). ORF132, ORF169, ORF356 and ORF399 extend inwards from the TmP towards the 3-fold axis.

**Figure 4.**
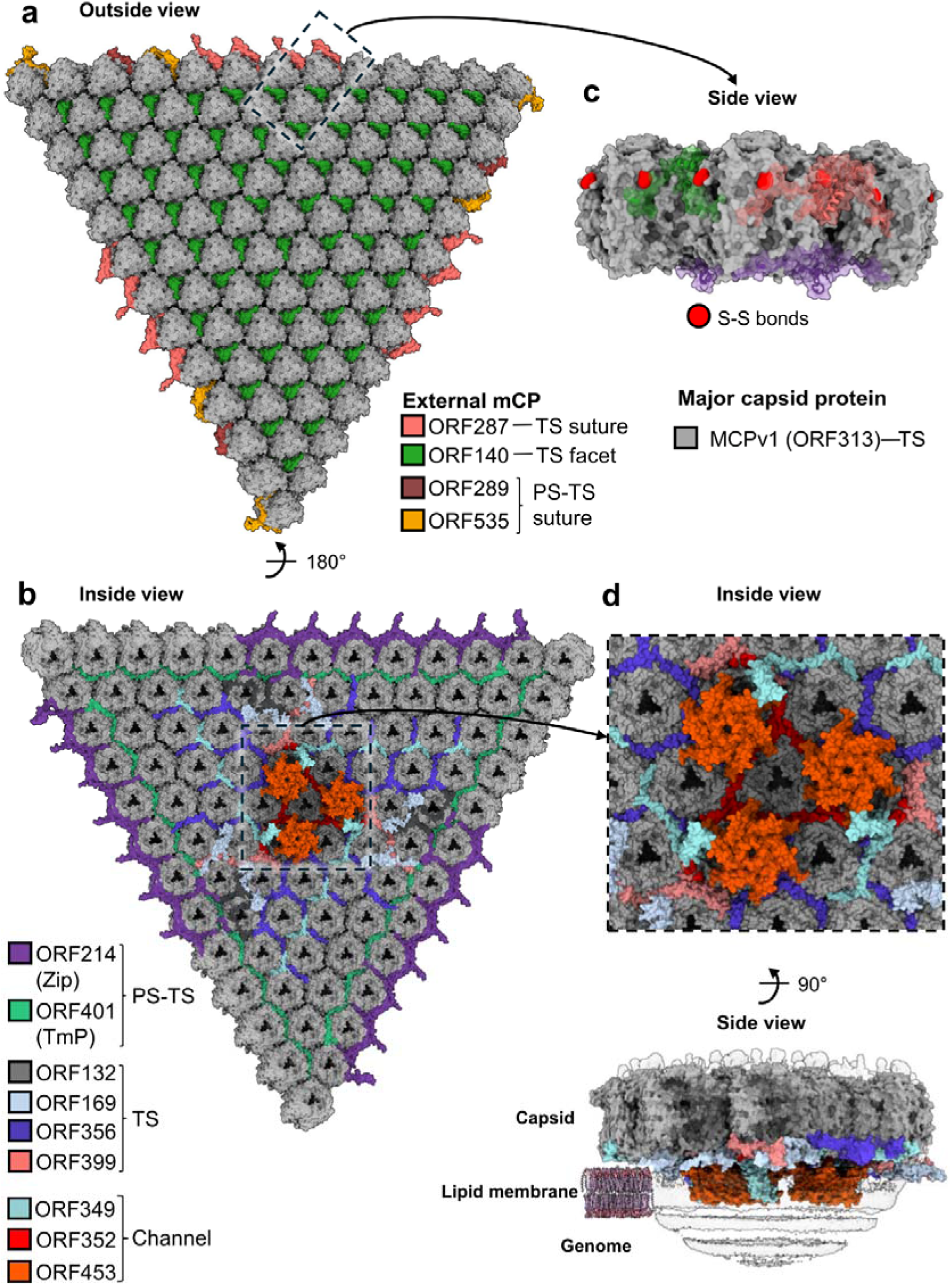
Trisymmetron organization of PkV-RF02 virions. (**a**) Outside view of the trisymmetron (TS). A side view of the region enclosed by the dashed square is depicted in (c). (**b**) Inside view of the trisymmetron. The region enclosed by the dashed square is zoomed in (d). (**c**) Side view of the capsid section showing ORF140 and ORF287 embedded in the valleys between the MCPs and locked in place by disulphide bonds between MCPv1 capsomers (red). Proteins coloured as in (a). (**d**) Inside and side views of the channel structure. In the bottom pannel, the cryo-EM map is depicted as a transparent surface.

Altogether, the PkV-RF02 capsid features a multilayered reinforcement architecture comprised by external mCPs, internal mCPs, and an extensive network of disulphide bridges between MCP capsomers, penton proteins, and mCPs (**Supplementary Fig. 3a**). This reinforcement is reflected in the exceptional stability of purified PKV-RF02 virions, which remained largely intact after harsh treatments that disrupt related viruses^28–30^ and were only partially disrupted by 25 mM TCEP (**Supplementary Fig. 3b**), well above concentrations typically used for protein denaturation.

### A membrane channel within the viral particle

The set of proteins traced under the trisymmetron is completed by ORF352, ORF349 and ORF453, which form the distinctive feature clasping the capsid to the internal membrane near the 3-fold axis (**Fig. 4b, d, Extended Data Fig. 1f, Supplementary Movie 1**). An ORF453 hexamer is inserted in the lipid bilayer, with Phe73 and Phe82 contributing to a largely hydrophobic central pore, 2.3 Å at its narrowest (**Supplementary Fig. 4a-c**). ORF349 is also inserted in the membrane and bridges the capsid shell to the channel hexamer (**Supplementary Fig. 4a-b**). ORF352, lacking transmembrane α-helices, is specifically positioned in the capsid near the channel. Foldseek identified potassium channels as the top eight matches for ORF453 (**Supplementary File 2g**), although the high E-values, as well as low DALI Z-scores (**Supplementary File 2h**), indicate only weak structural similarity. Although predicted potassium channels are common in giant viruses^31^, their organization within virions had not been directly observed. In PkV-RF02, the channel complex is the sole ordered feature associated with membrane-capsid contacts.

### Completing the spatial map of PkV-RF02 virions by XL-MS

Only 21 of the 136 proteins identified in the virion by mass spectrometry^19^ could be traced in the cryo-EM maps, likely because the remaining ones lack icosahedral symmetry, occur at low copy number, vary in organization among particles, or are flexible, causing their signal to be averaged out. To overcome this limitation, we used XL-MS to resolve the spatial distribution of the PkV-RF02 proteome in intact virions. We first evaluated particle stoichiometry by shotgun proteomics, using the protein copy numbers provided by the cryo-EM identified capsid proteins and their corresponding proteomics intensity as an internal standard to determine copy numbers for all remaining proteins (**Supplementary Fig. 5a, Supplementary File 3**). These estimates indicate that each viral particle contains near 14,300 protein copies, of which ∼63% correspond to traced capsid components (**Supplementary Fig. 5b**). Out of 172 different viral proteins detected in the shotgun proteomics experiment, 152 appeared in at least one copy (**Supplementary File 3**). XL-MS of virus particles detected 5,390 unique residue-to-residue connections between viral proteins (at 1% false discovery rate (FDR), **Supplementary File 4**). These connections amount to 561 PPIs between 112 different viral proteins, indicating that a substantial part of the virion proteome was covered.

Using our previous functional annotations and transmembrane predictions^19^, and unsupervised community clustering of the PPI network^32^, we defined four clusters consistent with the spatial distribution of virion components derived from cryo-EM (**Fig. 5**). Cluster 1 contains capsid proteins; cluster 3 is formed by proteins at the capsid-membrane interface; and genome-interacting proteins appear exclusively in cluster 4, the viral core. Proteins in cluster 2 were annotated as unknown. Only a very small proportion of cross-links occurred between clusters (2.4% between cluster 3 and 4), indicating that virion structural integrity was preserved during cross-linking. In contrast, intra-cluster cross-links were much more common: 55% in cluster 3, and 41.3% in cluster 4, consistent with the physical separation of these environments by the lipid bilayer (**Supplementary Fig. 5c**).

**Figure 5.**
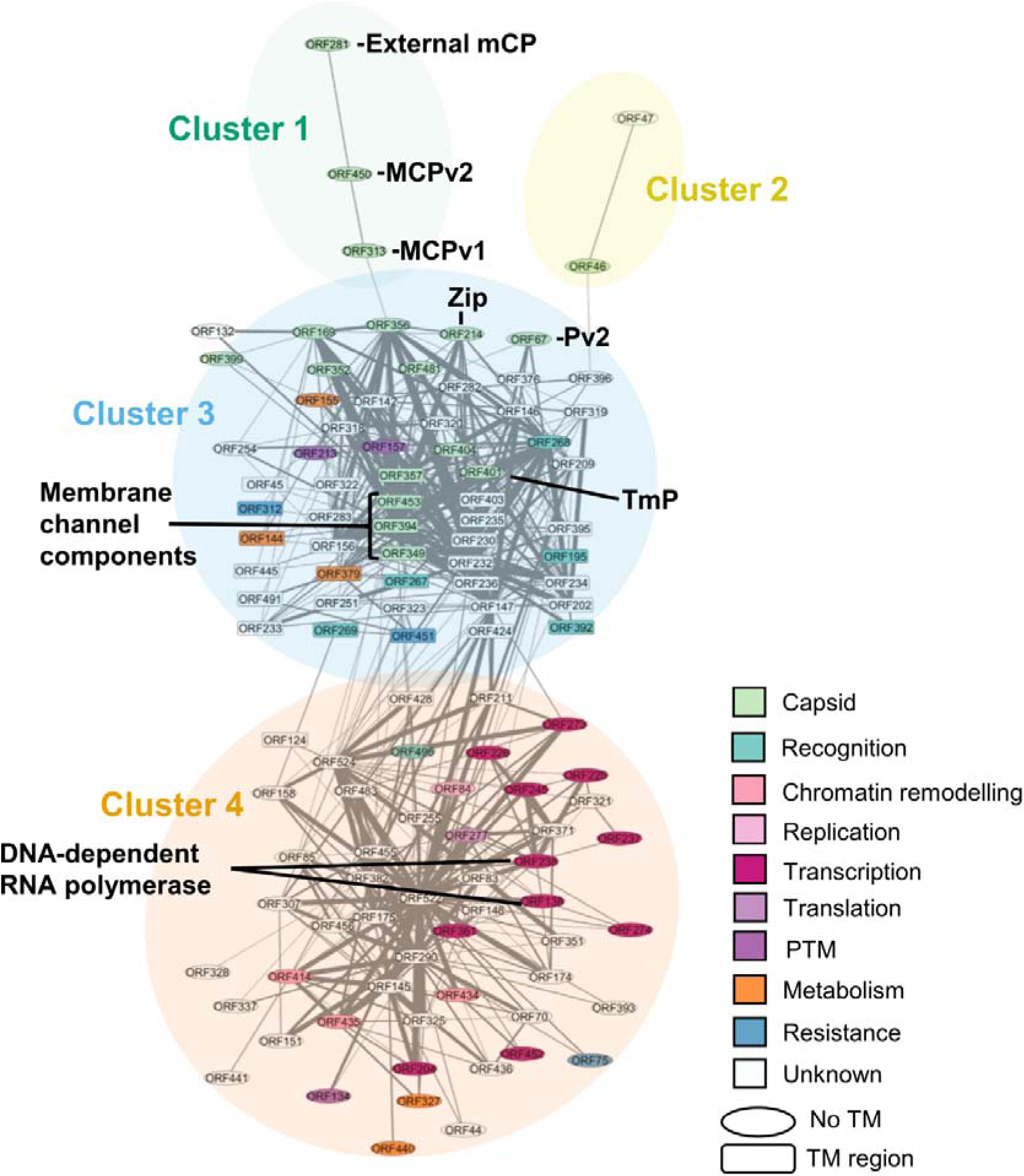
Protein-protein interactions (PPI) detected by XL-MS. General PPI network clustered in four different groups that correspond to their spatial distribution. Names of proteins with transmembrane motifs predicted in^19^ are enclosed in rectangular boxes. Box colours correspond to functional groups as assigned in^19^.

### Identification of additional capsid components: MCPv2, a third penton variant, and a fourth channel component

In cluster 1, we localized MCPv2, cross-linked to MCPv1 and to external pentasymmetron mCP ORF281 (**Supplementary Fig. 6a**). Cross-links involving ORF281 C-terminal residues (Lys199, Lys223) position MCPv2 at the pentasymmetron-trisymmetron interface (**Fig. 6a**). Additionally, a homomeric-interlink involving residue 428 of two different MCPv2 molecules complies with DSSO cross-link distance constraints (≤ 35 Å between Cα^33^) only in front-to-front capsomer interfaces (**Supplementary File 5a**). All assigned cross-links satisfied the expected distance restraints, except for one that exceeded the threshold slightly but remained within 40 Å. Together, they place MCPv2 at specific positions in the B and f capsomers (**Fig. 6a**). Nevertheless, these results do not exclude additional MCPv2 locations in the capsid. Cryo-EM refinement focused on capsomer B did not yield high resolution maps compatible with MCPv2 sequence, likely reflecting its low abundance (44 molecules per virion, 0.73 per AU) (**Extended Data Table 1**, **Supplementary File 3**). AlphaFold yields only low confidence predictions for MCPv1/MCPv2 heterotrimers (not shown), suggesting that incorporation of MCP variants differs from that of PBCV-1, where both homo- and heterotrimers were observed^10^.

**Figure 6.**
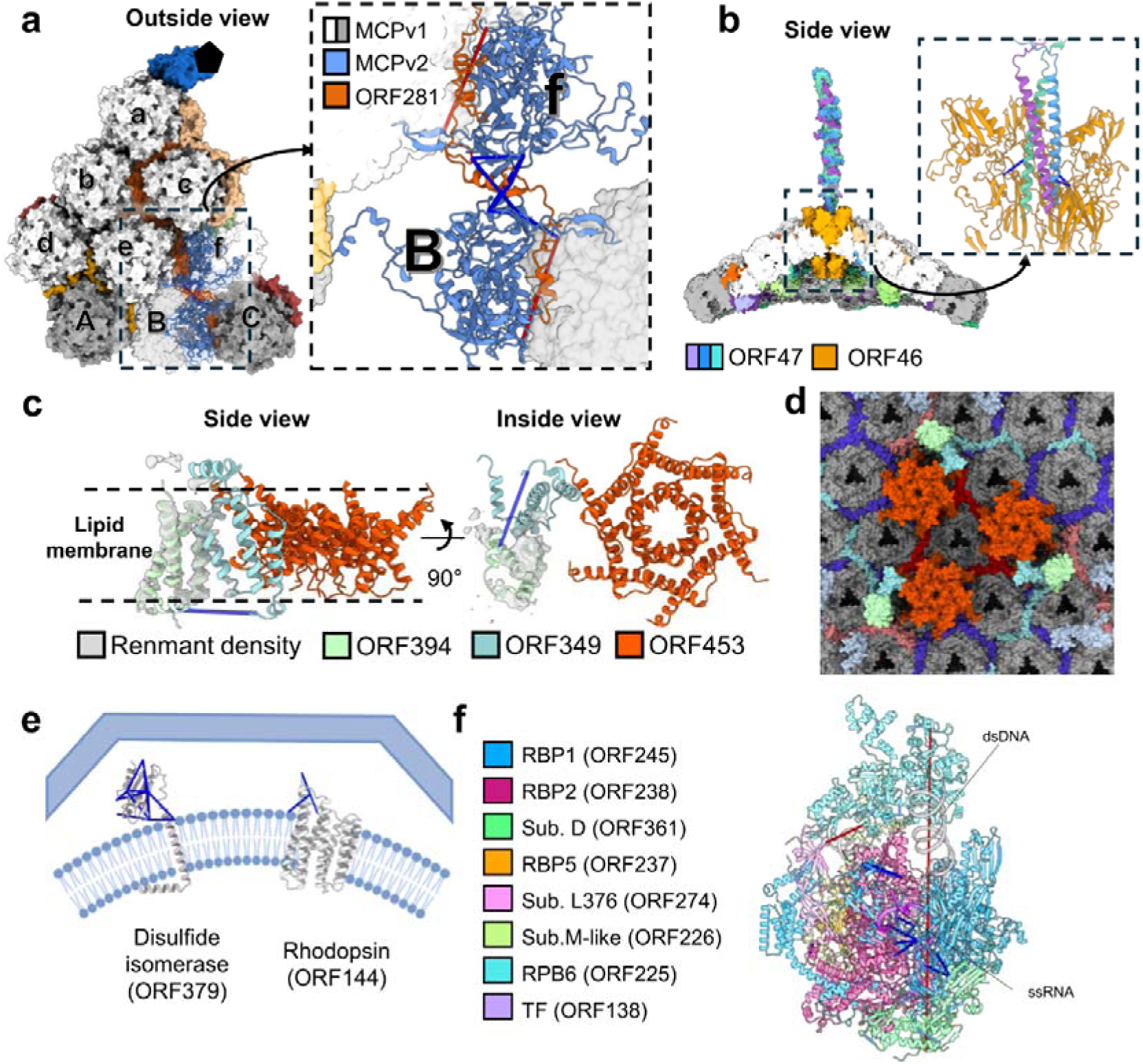
Localization of proteins in the viral particle based on XL-MS. (**a**) AlphaFold prediction for an MCPv2 (ORF450) monomer fitted in capsomer B chain 3 and capsomer f chain 1 positions according to XL-MS data. The panel at the right shows a closer look to appreciate satisfied crosslinks in blue (≤35 Å Cα-Cα), and non-satisfied in red. (**b**) Pentameric ORF46 and trimeric ORF47 AlphaFold predictions manually fitted in the vertex region according to XL-MS data. The panel at the right shows a closer look to appreciate satisfied crosslinks in blue. (**c**) Channel structure with AlphaFold prediction of ORF394 (residues 1-89) fitted in the remnant density, together with the traced models of ORF349 and ORF453. Only one cross-link is displayed, as it falls within the traced region of ORF349 and satisfies the DSSO distance constraint. (**d**) Inside view of the trisymmetron in the channel region, with ORF394 included. (**e**) Predicted models of transmembrane proteins disulphide isomerase (ORF379) and rhodopsin (ORF144) present in cluster 3. In the AlphaFold prediction of ORF379, nine out of eleven inter-protein cross-links passed the pLDDT 50 criterion, and all of them satisfied distance constraints. In the AlphaFold prediction of ORF144, one out of two intra-protein cross-links satisfied the pLDDT >50 threshold and DSSO distance constraints when mapped on the predicted ORF144 structure. (**f**) AlphaFold prediction of DNA-dependent RNA polymerase II complex present in cluster 4. Cross-linked residues are coloured according to DSSO distance constraint: satisfied interlinks in dark blue, satisfied intralinks in light blue, non-satisfied in red. Most inter-protein cross-links passed the pLDDT cutoff of >50 (62 out of 145), and 93% of them satisfied distance constraints. Distance measurements of identified cross-linked sites on PkV-RF02 proteins are summarized in Supplementary File 5.

In cluster 2, ORF46 and ORF47 (**Supplementary Fig. 6b**), previously annotated as unknown^19^, cross-link with cluster 3, pointing to spatial relation to the capsid. AlphaFold predicted with high confidence both a SJR fold reminiscent of penton proteins, and pentameric multimerization for ORF46, which shares 28.7% protein sequence identity with Pv1, and 26.3% with Pv2 (**Supplementary Fig. 6c**). Structural homology search indicated similarities to PBCV-1 singular vertex P1v1 (PDB: 8h2i)^10^ and EhV-201 penton protein (PDB: 8rbt)^25^ (**Supplementary File 2i-j**). These results, together with its calculated copy number (10 copies per particle, **Extended Data Table 1, Supplementary File 3**), strongly suggest that ORF46 is a third penton variant (Pv3). ORF47 is a remarkably long protein for which AlphaFold predicts a disordered N-terminal segment (residues 1-400) which could form a triple β-helix architecture reminiscent of bacteriophage tailspikes^34^ and PBCV-1 fibre-like proteins^10^, followed by a low confidence α-helical region (residues 400-700), and a C-terminal half with five well-defined domains (residues 700-1,568) (**Supplementary Fig. 6d-e**). These show structural homology to fibronectin domain-containing proteins (PDB: 2jll), interleukin (PDB: 7zan), and a cellular adhesion domain containing protein (PDB: 4ofy) (**Supplementary File 2k-n**). Copy number estimation for ORF47 (3.7 copies per particle, **Extended Data Table 1, Supplementary File 3**) supports the presence of one trimer per virion. Given the structural resemblance to fibre proteins involved in virus entry, and the presence of adhesion-related features, we propose that ORF47 may form a spike in a Pv3 singular vertex involved in PkV-RF02 host recognition or attachment. This hypothesis is further supported by the observation that all inter- and intra-protein cross-links satisfied DSSO distance constraints in an AlphaFold prediction of pentameric ORF46 in complex with a trimer of the ORF47 N-terminal segment (**Fig. 6b, Supplementary File 5b**).

In cluster 3, we found multiple cross-links between transmembrane proteins ORF349 (traced in the channel complex) and ORF394 (**Extended Data Fig. 5a**). AlphaFold predicted with high confidence three transmembrane α-helices (residues 1-89) for ORF394 (**Extended Data Fig. 5b**), in agreement with the shape of cryo-EM weak density forming a three-helix bundle whose identity could not be assigned. Together with the distance constraints for the N-terminal cross-link between the two proteins, these observations identify ORF394 as a component of the channel complex (**Fig. 6c-d, Supplementary File 5c**).

### Complex functions packaged in the viral particle

Clusters 3 and 4 exhibited significantly more complex PPI networks than clusters 1 and 2. Here, we highlight selected functions found in these networks. In cluster 3, ORF379 is structurally homologous to endoplasmic reticulum protein disulphide isomerases (ERdj5, PDB: 5ayk) and malaria thioredoxin (PDB: 4o32), enzymes that catalyse reversible oxidation of cysteine thiol groups^35^ (**Fig. 6e, Extended Data Fig. 5c**, **Supplementary File 2o-p**). ORF379 interacts with thirteen other proteins, including eight capsid proteins and channel components (**Extended Data Fig. 5d**), and cross-links place its globular domain facing the capsid interior. It harbours a catalytic Cys76-X-X- Cys79 motif consistent with disulphide isomerase activity^36^ (**Extended Data Fig. 5e**). Copy-number estimates indicate ∼35 copies per virion (**Supplementary Table 8, Supplementary File 3**).

Also in cluster 3 we find ORF144, with structural homology to phycodnavirus rhodopsins OLPVR1 (PDB: 9in9) and OLPVRII (PDB: 6sqg) (**Fig. 6e, Extended Data Fig. 5f, Supplementary File 2q-r**). ORF144 interacts with five proteins, four of which are located under the trisymmetron (**Extended Data Fig. 5g**). Giant virus rhodopsins bind retinal through a conserved lysine, and act as proton pumps^37,38^. Structural alignment of the ORF144 prediction with OLPVR1^39^ shows that Lys204 occupies the equivalent position within the transmembrane retinal-binding pocket, supporting its potential retinal-binding function (**Extended Data Fig. 5h**). Its very low abundance (∼0.15 molecules per virion) suggests that ORF144 is passively incorporated through host membrane carryover during assembly rather than functioning as a structural virion component (**Supplementary Table 8**, **Supplementary File 3**).

Cluster 4 forms the virion core and is dominated by proteins involved in genome organization, replication, and transcription (**Fig. 5**). We previously showed that PkV-RF02 packages a near-complete transcription machinery, including all eight virus-encoded DNA-dependent RNA polymerase subunits^19^. XL-MS revealed that seven RNA polymerase II subunits and the transcription factor ORF138 form a complex within the virion (**Fig. 6f, Extended Data Fig. 6**), indicating that the polymerase is packaged in a pre-assembled state to support immediate transcription upon infection. The major subunits RBP1 (ORF245; 133 copies) and RBP2 (ORF238; 161 copies), as well as ORF138 (110 copies), occur at similar abundances, whereas the remaining subunits are present at only 10-35 copies per particle, indicating that only a subset of packaged polymerases may be fully functional immediately after entry (**Supplementary Table 8, Supplementary File 3**).

An intriguing core component is the dinoflagellate/viral nucleoprotein (DVNP, ORF414)^19^, thought to contribute to genome organization through dsDNA-condensing activity. Although its high pI (10.7) and net charge (+28) could aid genome charge neutralization (**Supplementary Fig. 7a**), complete neutralization of the 583-kbp genome would require ∼41,700 DVNP molecules, far exceeding the ∼43 copies estimated per virion (**Supplementary Table 8, Supplementary File 3**). XL-MS data indicate that DVNP associates with other highly basic, largely disordered proteins (**Supplementary Fig. 7b-c**), hinting at a collective action to condense and organize the viral genome.

### Post-translational modifications in the PkV-RF02 virion

Phosphoproteomics identified 83 phosphorylated proteins in purified PkV-RF02 particles (∼50% of 161 viral proteins detected, **Supplementary File 6**). Categorization by function and PPI cluster revealed no obvious spatial or functional distribution (**Supplementary Fig. 8a-b**), indicating that phosphorylation is widespread across both structural and non-structural virion proteins. We detected 172 phosphorylation sites across 69 viral proteins (**Supplementary File 6**), with most proteins carrying a single site (**Supplementary Fig. 8c**). Exceptions include MCPv1, with six phosphorylation sites at its base (**Extended Data Table 2**), and proteins ORF455 and ORF147, each containing ∼10 phosphorylation sites clustered in disordered regions (**Supplementary Fig. 8d**).

Cryo-EM revealed densities consistent with O-linked glycans, most prominently at MCPv1 residues S357/S366/T380/S382 (**Extended Data Table 2**). In glycoproteomic analysis of purified PkV-RF02 virions to identify site-specific O-glycosylation (**Supplementary note 2**), raw MS/MS spectra revealed a striking and recurring pattern of oxonium ions, compatible with the presence of HexNAc (204 Th), HexHexNAc (366

Th), and the rare methyl-N-acetylated viosamine (MeVioNAc, 202 Th), previously described in bacteria and in APMV fibril glycans^40,41^ (**Supplementary Fig. 9a**). Glycans associated to the combined T357/T366 and T380/S382 sites reached masses up to ∼2700 Da and are likely linked to the peptide through a HexNAc residue (**Supplementary note 2**). Targeted search considering all glycans up to 2,700 Da containing at least 2 HexNAc, 1 Hex, and 1 MeVioNAc residue, yielded 28 high-confidence glycopeptide identifications, indicating. O-linked glycans composed of 4-7 monosaccharides and containing HexNAc-HexNAc, Hex-HexNAc, and MeVioNAc-MeVioNAc linkages, although the precise placement of MeVioNAc within the glycans remains unresolved (**Supplementary Fig. 9b**).

## Discussion

Although PkV-RF02 is a relatively small giant virus, its capsid and genome sizes are comparable to those of mycoplasmas^42^. Integrative combination of block-based cryo-EM reconstruction and XL-MS enabled high-resolution tracing of capsid proteins, assignment of weak densities, generation of a spatial PPI map, and, notably, identification of virion components that occur at low copy number or deviate from icosahedral symmetry (**Extended Data Fig. 3; Figs. 1, 5, 6a-c**). Whereas the former capabilities have been demonstrated previously^16,17,43^, our study expands this approach by providing insight into the latter. These analyses reveal a highly complex virion architecture featuring a multilayered capsid reinforcement system, three different vertex proteins, a membrane-spanning channel, extensive PTMs, a genome likely organized by DNA-condensing proteins, and a diverse repertoire of packaged enzymes (**Extended Data Fig. 7**).

The extensive network of intra- and intermolecular disulphide bonds in PkV-RF02 covalently cross-links the virion, complementing the stabilization provided by cementing mCPs. Together, these covalent and non-covalent interactions constitute one of the most elaborate capsid reinforcement systems described to date. Disulphide-mediated stabilization by ∼10^3^ bonds per particle has been reported in the baculovirus helical nucleocapsid^44^, and in siphophage Jabs^45^. The cumulative avidity of more than 20,000 inter-protein disulphide bonds, exceeding previous examples by roughly one order of magnitude, likely underlies the remarkable resistance of PkV-RF02 virions to harsh physicochemical treatments. For viruses infecting non-blooming haptophytes such as *Prymnesium kappa*, which typically occur at low abundances (up to 10^5^ cells/ml)^46^, enhanced particle stability may improve persistence in the ocean while awaiting host encounter. However, successful infection also requires capsid disassembly. Several features point to structurally weaker regions that may facilitate this process. None of the penton variants forms inter-subunit disulphide bonds, and their ability to form disulphide links with MCPv1 differs. AlphaFold models place no cysteines near MCPv1 Cys316 in Pv3, whereas the corresponding cysteines in Pv1 are more distant than in Pv2. In addition, MCPv2 lacks intramolecular disulphide bonds and is not predicted to form intermolecular disulphide links. These observations suggest that penton and MCP variants may contribute to regulating capsid opening. Notably, mCPs ORF140, ORF287 and ORF289 are locked in place by overlying disulphide bonds between MCPv1 capsomers (**Fig. 3a, 4c)**, pointing to an assembly pathway in which external mCPs are incorporated before inter-capsomer disulphide bonds are established. Both assembly and disassembly of the extensive disulphide network are likely tightly regulated. PkV-RF02 assembles in cytoplasmic viral factories^19^, where the reducing environment necessitates enzymatic assistance for disulphide bond formation. Similarly to other giant viruses^47–49^, PkV-RF02 encodes a disulphide isomerase (ORF379) and two sulfhydryl oxidases (ORF378 and ORF492), with ORF379 and ORF378 packaged in the virion (**Supplementary Table 8, Supplementary File 3**) and conserved across nucleocytoviruses^19^. Given that nucleocytovirus membrane and surface proteins can bypass the secretory pathway^50^, ORF379 may promote MCP and mCP folding and assembly at viral factory membranes, while sulfhydryl oxidases regenerate its catalytic activity. The incorporation of these enzymes into virions points to a dedicated system for regulating disulphide bonds.

Three penton variants are present in the PkV-RF02 capsid, in a previously unobserved architectural arrangement. Two were resolved by cryo-EM and occupy most vertices at similar frequencies, although their distribution is not obviously regulated. The third was detected only by XL-MS, likely because its low abundance prevented high-resolution reconstruction. The existence of three vertex types suggests functional specialization. Multiple penton variants could broaden host range, which may be advantageous when hosts occur at low abundance, as is the case for haptophytes^46^. Alternatively, one variant may mediate genome delivery, analogous to the PBCV-1 unique vertex or the APMV stargate^27^, or act as the nucleation site for capsid assembly, as proposed in giant virus assembly models^51^. Determining the functions of these distinct vertex types will require further study. Likewise, our observations on the organization of the PkV-RF02 membrane channel raise new questions about its function. The channel complex anchors the inner membrane to the capsid beneath the trisymmetron, forming the only detectable membrane-capsid connection in the particle, and may therefore contribute to membrane recruitment during assembly. Although it shares limited structural similarity with potassium channels, which are common in giant viruses^31,52^ and possibly involved in membrane depolarization during entry^53,54^, its hexameric architecture and lack of the conserved G(Y/F)G motif^55,56^ argue against a canonical potassium channel function. Whether this complex has been repurposed as an assembly scaffold or retains a role in cation transport during infection remains unclear.

PTMs add another layer of complexity to the virion. Half of all virion proteins, including MCPv1, carry phosphorylation sites. PkV-RF02 encodes and packages several proteins potentially involved in phosphorylation^19^. Coding for serine/threonine kinases is a hallmark of large DNA viruses^57^. The most heavily phosphorylated proteins in PkV-RF02, ORF455 and ORF147, contain intrinsically disordered regions, with several phosphorylation sites in ORF455 mapping to these regions (**Supplementary Fig. 8**). In other varidnaviruses, phosphorylation of intrinsically disordered regions regulates formation of condensates required to form infectious viral progeny^58^. ORF455 localizes to the core (cluster 4) and directly interacts with 11 viral proteins, among them the genome condensing DVNP; whereas ORF147 is membrane-associated (cluster 3) and has 36 detected PPI partners, including proteins in the core and capsid. These observations suggest that ORF455 and ORF147 may act as interaction hubs helping to recruit, concentrate, and compartmentalize virion components during assembly. Cryo-EM revealed extensive O-linked glycosylation of the PkV-RF02 capsid surface, while glycoproteomics detected oxonium ions corresponding to HexNAc, Hex, MeVioNAc, and 2MeVioNAc. Uncommon VioNAc-containing glycans have also been reported in the O-glycosylated surface fibrils of APMV, where they are implicated in host attachment^40,59^. In contrast, chlorovirus MCPs carry N-linked glycans^60–63^. Like other giant viruses, PkV-RF02 encodes components of a glycosylation machinery. Notably, we identified a glycosyltransferase homolog (ORF481) on the inner surface of the pentasymmetron. As with the membrane channel, it remains unclear whether ORF481 contributes to capsid glycosylation or has been co-opted for a structural role. PkV-RF02 also encodes several additional proteins homologous to glycosylation-related enzymes in other giant viruses^64^ (**Supplementary Table 9**).

In conclusion, our results demonstrate that giant virus particles are not merely scaled-up versions of conventional icosahedral capsids, but rather structurally and functionally complex machines. The remarkable degree of molecular specialization, compartmentalization, and biochemical potential revealed here raises fundamental questions about how this complexity contributes to host manipulation, virion stability, and infection success.

## Materials and Methods

### Virus production and purification

The host *Prymnesium kappa RCC3423* was grown in liquid IMR/2 medium^65^ at 15°C with 14:10 hours light:darkness cycles. *P. kappa* cells were diluted to 5×10^4^ algae/ml and incubated for 24 hours. Then, they were infected with 2% PkV-RF02 lysate and collected after 3-4 days when the culture seemed clear. The lysate was centrifuged at 3,500 xg, 20 minutes, at 10°C. The supernatant was concentrated by spinning at 25,000 rpm for 2 hours at 10°C (SW41/SW32 rotor) or alternatively, by ultrafiltration using Amicon® stirred cells with 10 kDa filter. The concentrated lysate was loaded onto a 4-layer OptiPrep™ gradient (45%-38%-32%-25% iodixanol diluted in SM buffer: 50 mM Tris-HCl, 100 mM NaCl, 8 mM MgSO_4_, 0.005% glycerol) and centrifuged at 25,000 rpm at 10°C for at least 19 hours (SW41 rotor). The virus band was collected approximately at 32% iodixanol and stored at 4°C until further use. For experimental assays, the virus was and dialysed against SM buffer for one hour in 0.025 μm pore size MCE Membrane Filters. The quality of purified viral preparations was assessed by analysing (i) structural integrity by negative staining in transmission electron microscopy, (ii) protein composition by denaturing electrophoresis (SDS-PAGE), and (iii) particle concentration by Nanoparticle Tracking Analysis (ZetaView).

### Cryo-EM sample preparation and data collection

Purified PkV-RF02 virions were dialyzed against PM2 buffer (20 mM Tris-HCl, 100 mM NaCl, 5 mM CaCl_2_) for one hour at 4°C. Samples were further concentrated by ultracentrifugation at 28,500 rpm at 10°C for 2 hours (SW55Ti) and deposited in glow-discharged (1-minute, 25 mA), Quantifoil R1.2/1.3 or R0.6/1 Copper/Rhodium (Cu/Rh) grids. Three consecutive incubations on the grid with 3 µl drops of the sample followed by blotting were performed to increase the amount of particles lying in the carbon holes^66^. Excess liquid was blotted and grids were vitrified in a FEI Vitrobot. Before data collection, grids were examined in a 200 kV Talos Arctica (FEI) microscope at the CNB-CSIC cryo-EM facility, to assess particle concentration, ice quality and overall suitability of the sample. For single particle analyses, cryo-EM images were obtained at the European Synchrotron Radiation Facility (ESRF, Grenoble, France) using a 300 kV Titan Krios microscope equipped with a K3 camera at a nominal pixel size of 1.35 Å/px at 64,000 x. Additional parameters are summarized in **Supplementary Table 1**.

### Cryo-EM single particle image processing

All image processing was performed within the Scipion3 framework^67^. Frame alignment was carried out with MotionCor2^68^. The contrast transfer function (CTF) was estimated using CTFFIND4^69^ and Xmipp3, and the best micrographs were selected based on consensus CTF^70^. Particles were picked manually using Xmipp3^70^ and extracted in 1620 x 1620 px box size (binned at 2 Å/px) using RELION^71^. 2D and 3D classifications were conducted in RELION^71^. The best 3D class was further processed by RELION 3D auto-refine followed by CTF refinement^72^. Corrections were applied in the following order: beam tilt refinement, anisotropic magnification, and per particle defocus correction. Finally, Ewald sphere correction was applied^73^, achieving an icosahedrally averaged final map at 4.7 Å resolution (**Extended Fig. 1d**). Additional image processing, reconstruction and refinement parameters are summarized in **Supplementary Table 1.**

For block-based reconstructions of the PkV-RF02 capsid, ten overlapping blocks were defined, to cover the AU. Blocks were centred on capsomers D, e, F, I, O, Q, and Y; as well as at the 2-fold, 3-fold and 5-fold symmetry axes (**Extended Data Fig. 2a, Supplementary Table 2**). The workflow explained in the following lines was accomplished for each of the ten blocks (**Extended Data Fig. 2b**).

Sub-particles were defined by Localrec^74,75^ and extracted in 256 x 256 px boxes from the original images at 1.35 Å/px. Due to uncertainty in the 4.7 Å icosahedral map handedness, the extraction was performed twice by changing the sign of the defocus correction. Both sets of sub-particles were reconstructed in parallel, and the subset yielding higher resolution was selected for further processing^71^. CryoSPARC local refinement was performed^76^ and subsequently, RELION 3D classification without alignment was conducted. The particles assigned to the best classes were selected for RELION 3D auto refine and post-processing, achieving 3-3.5 Å resolution (**Supplementary Fig. 1**). More details of the workflow are summarized in **Supplementary Table 2.**

### Model building and analysis

Initial models for MCPv1 and Pv1 were predicted using AlphaFold 2 or 3^77,78^. ChimeraX was used for rigid fitting of each chain initial model into the sharpened maps^79^. Next, the fitted model of each chain was refined using COOT^80^ and Phenix real-space refine^81^. To identify additional unmodeled densities, we generated remnant maps with ChimeraX by creating a map from the traced model (*molmap* command) and subtracting it from the main cryo-EM map. ModelAngelo was used to generate initial molecular models in the remnant map densities^82^. In the first ModelAngelo search, no sequence was provided, and results were submitted to BLAST (https://blast.ncbi.nlm.nih.gov/Blast.cgi) against the PkV-RF02 proteome (GenBank: PV100844.1). For hits with significant e-value, a second ModelAngelo search was completed with the hit protein sequence. Resulting hits were refined using COOT^80^ and Phenix real-space refine^81^. Most mCPs were modelled using sharpened maps produced by DeepEMhancer^83^. The N-terminal region of Pv2 was traced on the final Relion map, without postprocessing.

Interactions between proteins were analysed using PDBePISA^84^, and RING for π-π contacts, manually defining the threshold at 5 Å^85^. For structural alignment of the models, ChimeraX *matchmaker* command was used^79^. Map and model visualization, including Coulombic electrostatic potential colouring, were carried out using ChimeraX^86^. A statistical framework combining compositional and positional analyses, supported by Monte Carlo simulations, was used to assess whether the distribution and spatial arrangement of penton variants across virions deviate from random expectations (see **Supplementary note 1**). The channel profile was calculated using MOLEonline^87^. Traced models or AlphaFold structure predictions were sent to DALI (PDB25 comparison)^88^ and Foldseek (PDB100) servers^89^ for structural homology searches. Both results were considered to find a consensus.

### Disruption assays

PkV-RF02 virions were subjected to a range of disruptive treatments. These included osmotic shock with 0.5 M KCl for 3 hours, followed by 3 hours of 60 mM KCl^28^; 4 M urea overnight^29^; five cycles of freezing in liquid nitrogen and thawing at 100°C; exposure to pH 2 for up to 6 hours, and pH 2 and 100 °C for 1 hour^30^; and 25 mM Tris(2-carboxyethyl)phosphine (TCEP) with 30 mM chloroacetamide (CAA). The structural integrity of the virions following each treatment was assessed using negative staining.

### Negative-staining electron microscopy

Purified PkV-RF02 samples were incubated for 5 minutes on collodion-carbon coated Gilder copper grids (400 mesh square Cat#G400-C3) previously exposed to glow discharge for 15 seconds at 25 mA, washed in SM buffer or water, negatively stained for 30 seconds with 10 µl of 2% uranyl acetate and blotted. Micrographs were obtained with a 100 kV JEOL JEM 1011 transmission electron microscope at CNB-CSIC Electron Microscopy Facility.

### Preparation of purified cross-linked and non-crosslinked PkV-RF02 particles for MS analyses

PkV-RF02 virions were purified as described above. For bottom-up proteomics, four replicates of purified virus particles were generated from 3 litres of infected *P. kappa* culture sample, which were ultracentrifuged at 25,000 rpm for 2 hours and 10°C (SW28). The pellet was stored at −80°C until further processing.

For XL-MS, particles purified from approximately 15 l of infected *P. kappa* culture were ultracentrifuged at 25,000 rpm for 2 hours and 10°C (SW28). The pellet (around 200 µg) was resuspended in Phosphate Buffered Saline (PBS) and cross-linked by adding disuccinimidyl sulfoxide (DSSO) to 3 mM for 30 minutes at 25°C under shaking. The reaction was quenched by adding Tris-HCl to 50 mM final concentration for 20 minutes at 25°C. The sample was ultracentrifuged at 28,500 rpm for 2 hours at 10°C (SW55Ti) to concentrate cross-linked virion samples in a pellet. The pellet was stored at −80°C until further processing.

For phosphoproteomics and glycoproteomics, virions purified from approximately 15 l of infected *P. kappa* culture were purified and ultracentrifuged at 25,000 rpm for 2 hours and 10°C (SW28). The pellets (approximately 200 µg each) were stored at −80°C until further processing.

### Sample preparation for bottom-up proteomics, XL-MS, phosphoproteomics and glycoproteomics

#### Sample preparation for bottom-up proteomics

The four replicates of purified virus were lysed in 8 M urea in Triethylammonium bicarbonate (TEAB) with sonication using a Bioruptor Pico system (Diagenode) at 10 °C, employing 10 cycles of 30-second sonication pulses followed by 30-second pauses. To degrade genomic DNA, 1 µl of Benzonase® Nuclease (70746, Merck) was added to each sample and incubated at room temperature for 30 minutes. Proteins were subsequently extracted and precipitated using methanol/chloroform precipitation^90^. The resulting protein pellets were resuspended in a buffer containing 1% (w/v) sodium deoxycholate (SDC), 5 mM TCEP, and 40 mM CAA in 50 mM triethylammonium bicarbonate (TEAB). Proteolytic digestion was performed overnight at 37 °C with shaking at 1,000 rpm, using trypsin and endoproteinase Lys-C at enzyme-to-protein ratios of 1:25 and 1:100, respectively. Digestion was halted by acidifying the samples with 1% (v/v) formic acid (FA). Following centrifugation, peptides were desalted using C18 stage-tips^91^, dried under SpeedVac, and stored at –20 °C until LC-MS/MS analysis.

#### Sample preparation for XL-MS

The cross-linked PkV-RF02 sample was resuspended in 100 µL of PBS, sonicated for 15 minutes (30 seconds on, 30 seconds off) in a Bioruptor Pico system at 4°C, and deglycosylated using Protein Deglycosylation Mix II (P6044, NEB) under denaturing conditions following the manufacturer instructions. Samples were lysed by addition of 7 M urea, 5 mM TCEP and 50 mM CAA. Approximately 40 units of Benzonase® Nuclease were added for 1 hour at room temperature. Subsequently, the sample was subjected to sonication in a Sonoplus HD 2070 equipped with a MS73 Sonotrode (Bandelin) on 50% intensity for approximately 1-minute. Proteins were extracted using methanol-chloroform precipitation according to standard protocols^90^, dried and resuspended in digestion buffer (50 mM TEAB pH 8, 1% SDC, 30 mM CAA and 5 mM TCEP) and finally, digested by adding trypsin at an enzyme-to-protein ratio of 1:25 (w/w) and Lys-C at a 1:50 ratio (w/w) and incubated at 37°C overnight in the dark. Digestion was stopped by adding 1% FA. Peptides were desalted using reverse phase columns Sep-Pak C8 (Waters) according to the manufacturer’s instructions, dried in the SpeedVac and stored at −20°C.

#### Sample preparation for phosphoproteomics

Approximately 200 µg of purified PkV-RF02 were resuspended in 20 mM 4-(2-hydroxyethyl)-1-piperazineethanesulfonic acid (HEPES) (pH 7.4) with PhosSTOP™ (Roche, 4906845001) and processed as described for the cross-linked sample. Briefly, the sample was sonicated and deglycosylated under denaturing conditions following the manufacturer instructions. Then, the sample was lysed and proteins were extracted using methanol-chloroform precipitation according to standard protocols^90^. The extracted proteins were dried and digested by trypsin at an enzyme-to-protein ratio of 1:25 (w/w) and Lys-C at a 1:100 ratio (w/w) at 37°C overnight in the dark. Peptides were desalted using reverse phase columns Sep-Pak C18 cartridges (Waters). Peptides were dried by SpeedVac.

#### Sample preparation for glycoproteomics

Approximately 200 µg of purified PkV-RF02 were resuspended in 8 M urea, 5 mM TCEP and 50 mM CAA, and processed as described for the cross-linked sample. Briefly, the sample was lysed and proteins were extracted using methanol-chloroform precipitation according to standard protocols^90^. The extracted proteins were dried and digested as described in *Kang et al.*^92^. Samples were treated with IMPa O-glycoprotease at an enzyme-to-protein ratio of 1:20 overnight at 37°C in 2 M urea, 20 mM Tris-HCl. The sample was further digested overnight at 37°C in the dark with AspN (1:50, w/w). Peptides were desalted using reverse phase columns Sep-Pak C18 cartridges (Waters), and dried by SpeedVac.

### Off-line fractionation of cross-linked peptides

Peptides were quantified by Pierce quantitative colourimetric assay (Thermo Fisher Scientific). Cross-linked peptides were fractionated by strong cation exchange (SCX) chromatography using a PolySULFOETHYL A column (100 x 2.1 mm) in an Agilent 1260 Infinity II system using a 95-minute gradient from buffer A (0.05% FA in 20% ACN) to buffer B (0.05% FA, 0.5 M NaCl in 20% ACN). Gradient was as follows: 0 min 0% buffer B, 0.01 min 2% buffer B, 8.01 min 3% buffer B, 14 min 8% buffer B, 28 min 20% buffer B, 48 min 40% buffer B, 68 min 90% buffer B, 74 min 90% buffer B, 75 min 0% buffer B, 95 min 0% buffer B, flow rate 0.18 ml/min with fraction collection in 100 seconds intervals. Twelve fractions with cross-linked peptides were selected and desalted in C8 stage-tips^91^ following manufacturer instructions. Peptides were dried by SpeedVac and stored at −20°C before LC-MS/MS measurements.

### Phosphopeptide enrichment

Phosphopeptides were enriched by High Select^TM^ reagents for Phosphopeptide Enrichment Kit (Thermo Scientific, A32993) following the manufacturer’s instructions. Briefly, TiO_2_ Spin Tip columns were equilibrated by 20 µl Wash Buffer and Binding/Equilibration Buffer, centrifuging at 3,000 ×g for 2 minutes. Suspended peptides in the Binding/Equilibration Buffer were loaded to the spin tip and centrifuged at 1,000 ×g for 5 minutes, and the sample was reapplied once. The column is consequently washed by applying 20 µl Binding/Equilibration Buffer and Wash buffer, twice and applying 20 µl LC-MS/MS grade water. Elution was carried out by adding 50 μl of Phosphopeptide Elution Buffer and centrifugating 1,000 ×g for 5 minutes, twice. Immediately, peptides were dried by SpeedVac.

### Glycopeptide enrichment

Glycopeptide enrichment was carried out using HyperSep™ Retain AX cartridges according to the manufacturer’s instructions. Briefly, the dried peptide mixture was resuspended in loading buffer (95% ACN, 1% TFA) and applied to pre-equilibrated RAX cartridges. After loading, the cartridges were washed three times, and bound O-glycopeptides were subsequently eluted with 400 μl of elution buffer (50% ACN, 0.1% TFA) into a fresh tube. The eluate was dried in a SpeedVac concentrator and stored at −80°C until LC-MS/MS analysis.

### LC-MS/MS data collection

#### LC-MS/MS for bottom-up proteomics

Four replicate samples were resuspended in 1% ACN and 0.05% TFA and analysed using an Orbitrap Astral mass spectrometer (Thermo Fisher Scientific) operating with instrument control software Xcalibur 4.7 and coupled online to a Vanquish Neo UHPLC system (Thermo Fisher Scientific). Approximately 0.5 µg of peptides were loaded onto a 20 cm in-house packed C18 analytical column (Poroshell 120 EC-C18, 2.7 µm; Agilent Technologies) and separated over a 40-minute gradient from buffer A (0.1% FA in water) to buffer B (0.1% FA in 80% ACN) and a flow rate of 250 nl/min. MS1 spectra were acquired in the Orbitrap at a resolution of 180,000, across a mass range of 350-1400 m/z, with a maximum injection time of 50 ms and an AGC target of 500%. An intensity threshold of 1e^4^ was applied, and precursor ions with charge states of 2-5 were selected for fragmentation via HCD using a normalized collision energy of 30%. Dynamic exclusion was enabled with a duration of 10 seconds. MS2 spectra were acquired using the Astral mass analyser with a scan range of 150-2,000 m/z, a maximum injection time of 2.5 ms, and AGC target set to ‘Standard’.

#### LC-MS/MS for XL-MS

Each of the twelve fractions collected using off-line fractionation of cross-linked peptides was resuspended in 1% ACN and 0.05% TFA and analysed using an Orbitrap Fusion Lumos mass spectrometer (Thermo Fisher Scientific) operating with instrument control software Xcalibur version 4.0 and coupled to an Ultimate 3000 nano HPLC system. A Field Asymmetric Ion Mobility Spectrometry (FAIMS) Pro Duo interface was employed. Peptides were separated with a 180-minute gradient going from buffer A (0.1% FA in water) to buffer B (0.1% FA in ACN) and a flow rate of 250 nl/min. FAIMS compensation voltages were alternated between −50, −60, and −75 V. MS1 parameters were 120,000 resolution, isolation window 1.6 m/z, 375-1,600 m/z scan range, injection time 50 ms and AGC target of 100%. The selected charge states for MS2 were 4-8, and precursors were fragmented by stepped HCD with normalized collision energies of 21%, 27%, and 33%. MS2 spectra were recorded in the Orbitrap with resolution set to 60,000, injection time 118 ms and AGC target 200%.

To generate a targeted database for cross-link search, a small aliquot of digested peptides taken before SCX fractionation was analysed using an Orbitrap Fusion Mass Spectrometer (Thermo Fisher) using the same parameters described for cross-linked peptide analysis, with the exception that no FAIMS was used, precursors of charge 2-4 were fragmented by HCD (normalized collision energy 30 %), and MS2 spectra were recorded in the ion trap with scan rate set to ‘Rapid’ and scan range set to ‘Auto’, 35 ms injection time and AGC target set to ‘Standard’.

#### LC-MS/MS for phosphoproteomics

Phosphopeptides were resuspended in 1% ACN and 0.05% TFA and analysed in an Orbitrap Exploris 480 mass spectrometer (Thermo Fisher Scientific) operated by instrument control software Xcalibur version 4.2 and coupled to a Vanquish Neo UHPLC system using a 120-minute gradient going from buffer A (0.1% FA in water) to buffer B (0.1% FA in 80% ACN). FAIMS was employed with alternating compensation voltages in between −40, −55, and −70 V. MS1 spectra were recorded in the orbitrap with resolution set to 60,000, isolation window of 1.6 m/z, 375-1,500 m/z scan range, injection time 50 ms and AGC target set to ‘Standard’. The selected charge states for fragmentation were 2-5 and precursors were fragmented by HCD with a normalized collision energy of 30%. MS2 spectra were recorded in the orbitrap at a resolution of 30,000, with maximum injection time set to ‘Auto’ and AGC target set to ‘Standard’.

To generate a targeted database for phosphoproteome search, a small aliquot of digested peptides taken before enrichment was analysed using a Orbitrap Exploris 480 mass spectrometer (Thermo Fisher Scientific) using the same parameters described for phosphopeptide analysis, with the exception that FAIMS was at −60 and −70 V, MS1 at 120,000 resolution, maximum injection time at auto, and AGC target at 300% precursors of charge 2-5. MS2 spectra were recorded in the orbitrap at a resolution of 15,000.

#### LC-MS/MS for glycoproteomics

Glycopeptides were fragmented using EThcD^93^ and examined in an Orbitrap Fusion mass spectrometer (Thermo Fisher Scientific) operated by instrument control software Xcalibur version 4.0 and coupled to a Vanquish Neo UHPLC system. Glycopeptides were resuspended in 1% ACN and 0.05% TFA and separated using a 120-minute gradient going from buffer A (0.1% FA in water) to buffer B (0.1% FA in ACN). MS1 spectra were recorded in the Orbitrap with resolution set to 120,000, isolation window 1.6 m/z, 400-1,400 m/z scan range, injection time 50 ms and AGC target set to 40000. The selected charge states for fragmentation were 2-7 and precursors were fragmented by HCD with a normalized collision energy of 35%. EThcD fragmentation was triggered upon detection of oxonium ions (m/z 126.055, 138.0549, 144.0655, 168.0654, 186.076, 204.0865, 274.0921, 292.1027, 366.1395). MS2 spectra following HCD were acquired in the Orbitrap at a resolution of 30,000, with maximum injection time set to 60 ms and AGC target set to 50000. MS2 spectra following EThcD were acquired at 30,000 and 120-4,000 scan range with maximum injection time set to 200 ms and AGC target set to 10000.

### LC-MS/MS data analysis

#### Bottom-up proteomics data-analysis

Raw data files were processed using FragPipe v23 with MSFragger^94^, applying the standard label-free quantification (LFQ) workflow and enabling ‘Match between runs’. Raw files were searched against a concatenated target-decoy database generated from a chimera proteome created merging the PkV-RF02 proteome (GenBank: PV100844.1), two host-related proteomes (*Emiliania huxleyi*, GenBank: GCF_000372725.1, and *Chrysochromulina tobinii*, GenBank: GCA_001275005.1), since *Prymnesium kappa Rcc3423* genome is not sequenced, and reference proteomes from bacterium genera present in the host culture: *Sulfitobacter, Balneola, Hyphomosnas, Marivita, Algimonas, Jannaschia, Hoeflea,* and *Sphingorhabdus*. Protein intensities were extracted from the combined_protein.tsv file.

Intensity values were log-transformed and fitted to known copy numbers on a double-logarithmic scale using R (version 4.5). A set of 12 proteins (ORF313, ORF140, ORF287, ORF356, ORF401, ORF132, ORF357, ORF404, ORF76, ORF481, ORF281, ORF289) with known copy numbers from the cryo-EM molecular model was used as internal standard. The remaining traced proteins were excluded from the analysis due to uncertainties in stoichiometry: penton variants, transmembrane proteins, or proteins presenting faint density at certain positions, which may indicate low occupancy. The slope and intercept from the resulting linear regression were used to estimate the copy numbers of all remaining proteins according to the following equation:

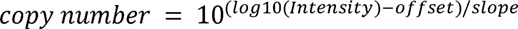

For subsequent analysis, the mean values from four technical replicates were used.

#### XL-MS data analysis

A targeted database for cross-link search was built from the results of linear peptides LC-MS/MS run, which was analysed in FragPipe v22^94^, with MSFragger^95^, applying ‘Default’ parameters. Raw files were searched against the concatenated target-decoy database generated from a chimera proteome as described above for LFQ analysis. Proteins detected in the sample were extracted from the protein.tsv file, and a new FASTA file was generated containing only these proteins, which served as the target database for the cross-linked peptide data search.

Then, XL-MS raw files were analysed in SCOUT1.5.1^96^, with the following settings: 10 ppm MS1 ion mass tolerance, 20 ppm MS2 ion mass tolerance, cysteine carbamidomethylation as fixed modification, methionine oxidation as variable modification, trypsin as enzymatic digestion with 3 missed cleavages as maximum, and DSSO cross-linker, 158.0038 Da (short arm, 54.0106 Da; long arm, 85.9824 Da). The search was completed with 6 amino acid minimum length of peptide and 1% FDR at cross-linked spectrum matches (CSM) level, residue pair level, and PPI level. Residue pair and PPI network results were exported and visualized by Cytoscape v3.7.2^97^ and for residue level representation XlinkCyNET plugin was used^98^.

#### Phosphoproteomics data analysis

As described for XL-MS data analysis, a targeted database was generated for phosphoproteome search as follows: the results of phosphorylated and non-phosphorylated peptide LC-MS/MS were analysed with FragPipe v22, and a new FASTA file was generated containing proteins present in the sample, which served as the target database for the phosphorylated peptide data search.

Then, phosphopeptide data were analysed in FragPipe v22^94^, with MSFragger, applying the LFQ phospho workflow: 20 ppm MS1 ion mass tolerance, 20 ppm MS2 ion mass tolerance, cysteine carbamidomethylation as fixed modification, methionine oxidation, serine, threonine and tyrosine phosphorylation (79.9666 Da) as variable modification, and trypsin as enzymatic digestion with 2 missed cleavages as maximum. Protein identities were extracted from protein.csv and phosphorylation sites were extracted from the site_STY.tsv file.

#### Glycoproteomics data analysis

The glycoproteomics data collected on the glycopeptide enriched fraction was searched using Byonic^99^. Digestion specificity was set to N-terminal cleavage of D allowing for 1 missed cleavage. The search included carbamidomethyl (+57.021464 Da) as fixed modification of cysteine, variable (common 2) oxidation (+15.994915) of methionine and tryptophan, and variable (rare 1) O-linked glycosylation at serine or threonine. We enumerated all possible glycan compositions containing at least 2 HexNAc, 1 Hex, and 1 MeVioNAc residue up to 2,700 Da (total of 265 compositions). The MeVioNAc residue is not part of Byonic’s standard repertoire of glycan residues and was instead encoded as a ‘Pent’ with an additional mass of +69.0578 Da per residue. We conducted a focused search on MCPv1 and used a precursor and fragment mass tolerance of 10 and 20 ppm, respectively. Results were filtered for glycopeptides with score ≥ 400. Please note that all glycan compositions listed represent the combined mass of two putative glycosylation sites per glycopeptide. The glycan oxonium ions were assigned using analysis of the raw MS/MS spectra in Annotator v1.3^100^.

### Clustering of PPI network

To identify functional clusters within the PPI network derived from XL-MS, interaction data were imported and processed in R using the edge betweenness function of the *igraph* package^101^. In brief, PPIs were treated as edges of a graph with weights corresponding to the number of cross-links per PPI, and clusters were identified by unsupervised community clustering following the Girvan-Newman algorithm^32^.

### Mapping of cross-links into models

Cross-link mapping to AlphaFold predicted structures for MCPv2, Pv3, ORF394, rhodopsin (ORF144), disulphide isomerase (ORF379), and DNA-dependent RNA polymerase II, was performed using ChimeraX. For monomeric models of ORF144 and ORF379, only intralink residue pairs were considered, while for multimeric structures, intralink and interlink residue pairs were used. First, amino acids with pLDDT score below 50 were excluded from the analysis. For each residue pair in all five AlphaFold models, Cα-Cα distances were calculated, with distance below or equal to 35 Å considered satisfied, based on the maximum span of the DSSO cross-linker, lysine side chain length and protein flexibility^33^. Models were ranked based on the satisfied cross-link percentage. Cα-Cα distances were plotted in ChimeraX^86^, where distances below or equal to 35 Å were coloured blue and those larger than 35 Å were coloured red.

### Data availability

The PkV-RF02 cryo-EM maps have been deposited at the Electron Microscopy Data Bank (EMDB, http://www.ebi.ac.uk/pdbe/emdb) with accession numbers EMD-55917 (icosahedrally averaged map), 55926 (block D), 55914 (block e), 55913 (block F), 55902 (block I), 55903 (block O), 55911 (block Q), 55915 (block Y), 55901 (block 2F), 55900 (block 3F), 56066 (block 5F1), 55916 (block 5F2). The molecular models are deposited at the Protein Data Bank (PDB, https://www.ebi.ac.uk/pdbe/) with accession numbers 9THG (block D), 9TH1 (block e), 9TH0 (block F), 9TGK (block I), 9TGL (block O), 9TGT (block Q), 9TH2 (block Y), 9TGI (block 2F), 9TGH (block 3F), 9TM8 (block 5F1), 9TH4 (block 5F2). The validation reports are included as Supplementary file S1.

The PkV-RF02 mass-spectrometry data have been deposited at the ProteomeXchange database (https://www.proteomexchange.org/) via PRIDE (https://www.ebi.ac.uk/pride/), with accession number PXD075279 for bottom-up proteomics, XL-MS and phosphoproteomics data; and accession number PXD086706 for glycoproteomics data.

## Acknowledgements

Work supported by grants from the Spanish State Research Agency, with co-funding from the European Regional Development Fund (PID2019-104098GB-I00/AEI/10.13039/501100011033 and PID2022-136456NB-I00/AEI/10.13039/501100011033), and Spanish State Research Agency PID2025-168455NB-I00/AEI/10.13039/501100011033, to C.S.M; and The Research Council of Norway (Norges Forskningsråd) through VirVar-project (294363) to R.-A.S. The CNB-CSIC was further supported by AEI Severo Ochoa Excellence grants SEV-2017-0712/AEI/10.13039/501100011033 and CEX2023-001386-S/AEI/10.13039/501100011033. S.O.U. held a CSIC JAE Severo Ochoa and Maria de Maeztu fellowship (JAE-SOMdM20-20), a predoctoral contract from Spanish Ministry of Science, Innovation and Universities (FPU2020-05148), and an EMBO Scientific Exchange Grant (10981).

We thank Beatriz Martín, Javier Chichón and Noelia Zamarreño from the CNB-CSIC Electron and Cryo-electron Microscopy facilities for the excellent technical support; as well as Gregory Effantin, Eaazhisai Kandiah, and Romain Linares, from European Synchrotron Radiation Facility (ESRF), for Titan Krios data collection under proposals MX-2443 and MX-2369: 10.15151/ESRF-ES-1061180097, 10.15151/ESRF-ES-925344867, 10.15151/ESRF-ES-781088676.

## Author contributions

S.O.-U. and C.S.M. conceived and designed the study, interpreted the data and wrote the manuscript with input from all authors. S.O.-U. and G.N.C. performed cryo-EM sample vitrification and data collection. R.M. performed cryo-EM data processing and penton distribution analyses. S.O.-U., G.N.C., P.H. and C.S.M. contributed to cryo-EM data processing, structural model building and data interpretation. S.O.-U. purified virions for shotgun proteomics and co-purified virions for XL-MS, phosphoproteomics and glycoproteomics analyses, prepared samples for mass spectrometry with L.M., and analysed and interpreted XL-MS, phosphoproteomics and shotgun proteomics data together with P.H., L.M. and F.L. H.W. purified virions for cryo-EM analyses. L.O.L. contributed to large-scale virion production and purification for XL-MS, phosphoproteomics and glycoproteomics analyses. J.S. and R.V. analysed and interpreted glycoproteomics data. H. W. and R.-A.S. provided initial virus stocks, genomic resources and expertise on virus and host biology. C.S.M. supervised the project and acquired funding. All authors discussed the results, critically revised the manuscript and approved the final version.

