## Extended Figures and Tables for "Integrative structural biology of a giant marine virus reveals architectural and functional complexity beyond icosahedral symmetry"

**EXTENDED MATERIAL FOR**

**Short title:** Integrative structural biology of a T=219 virus

**Extended Data Figures**

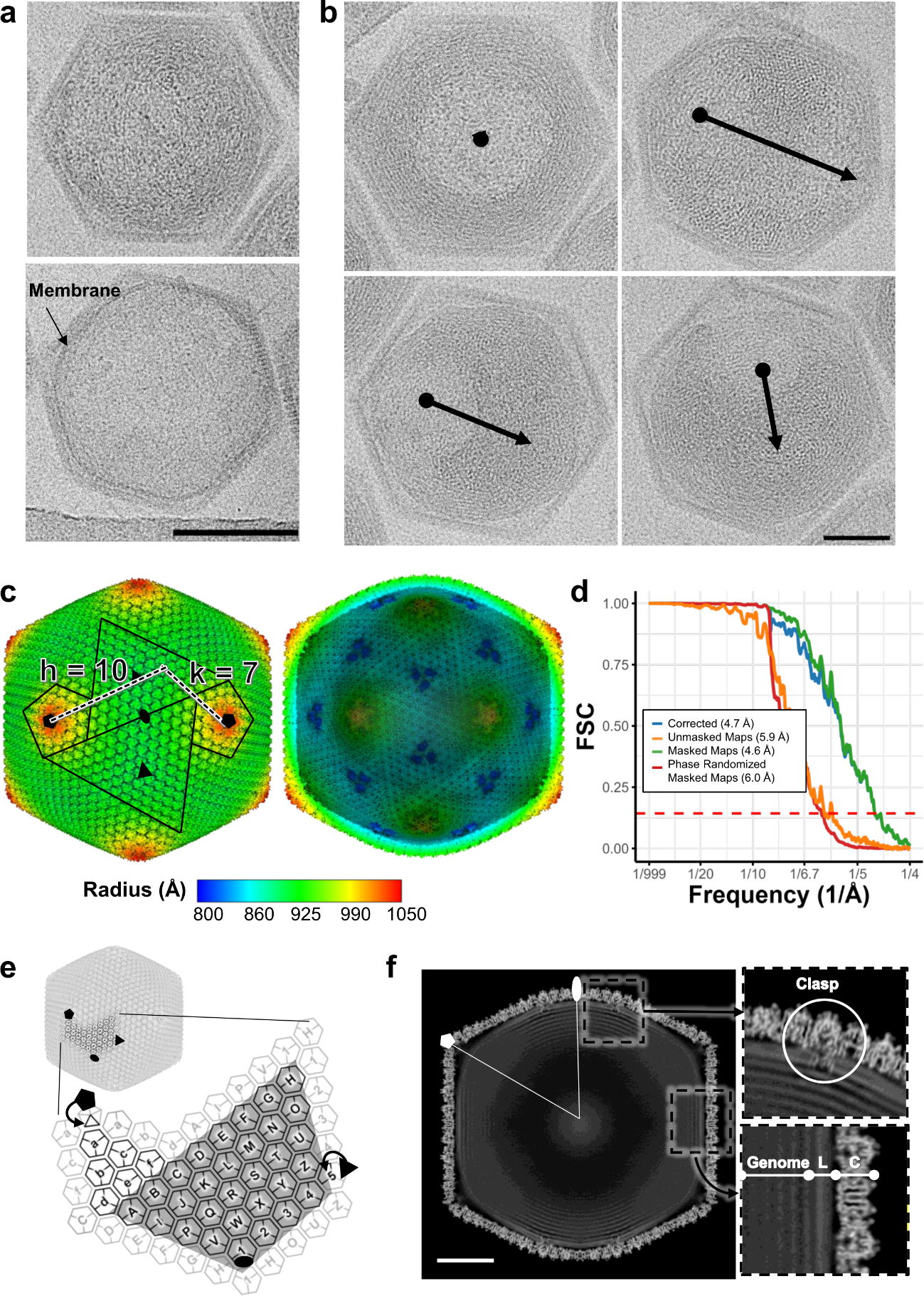

**Extended Data Fig. 1.** **Overall capsid structure of PkV-RF02.** (**a**) Cryo-EM images showing a genome-containing (top) and a genome-less (bottom) PkV-RF02 particle. (**b**) PkV-RF02 particles imaged in different orientations. Arrows indicate the Z-axis of the genome toroidal organization. The scale bar represents 100 nm in (a-b). (**c**) PkV-RF02 cryo-EM map radially coloured. Right: Dashed black lines indicate the h and k lattice coordinates used to calculate the triangulation number (T = h^2^ + hk + k^2^). Black lines indicate pentasymmetron and trisymmetron borders. Black symbols (pentagon, triangle, oval) indicate the 5-fold, 3-fold and 2-fold icosahedral axes. Left: map with the front half computationally removed to show the inner capsid surface. (**d**) Fourier Shell Correlation (FSC) curves indicating a 4.7 Å average resolution in the icosahedrally averaged map, RELION post-process. (**e**) Schematic representation of the icosahedral asymmetric unit. Numbers and letters identify the MCP capsomers. (**f**) Central slice of the PkV-RF02 map. The scale bar represents 50 nm. Dashed squares are zoomed at the right. C, capsid; L, lipid membrane.

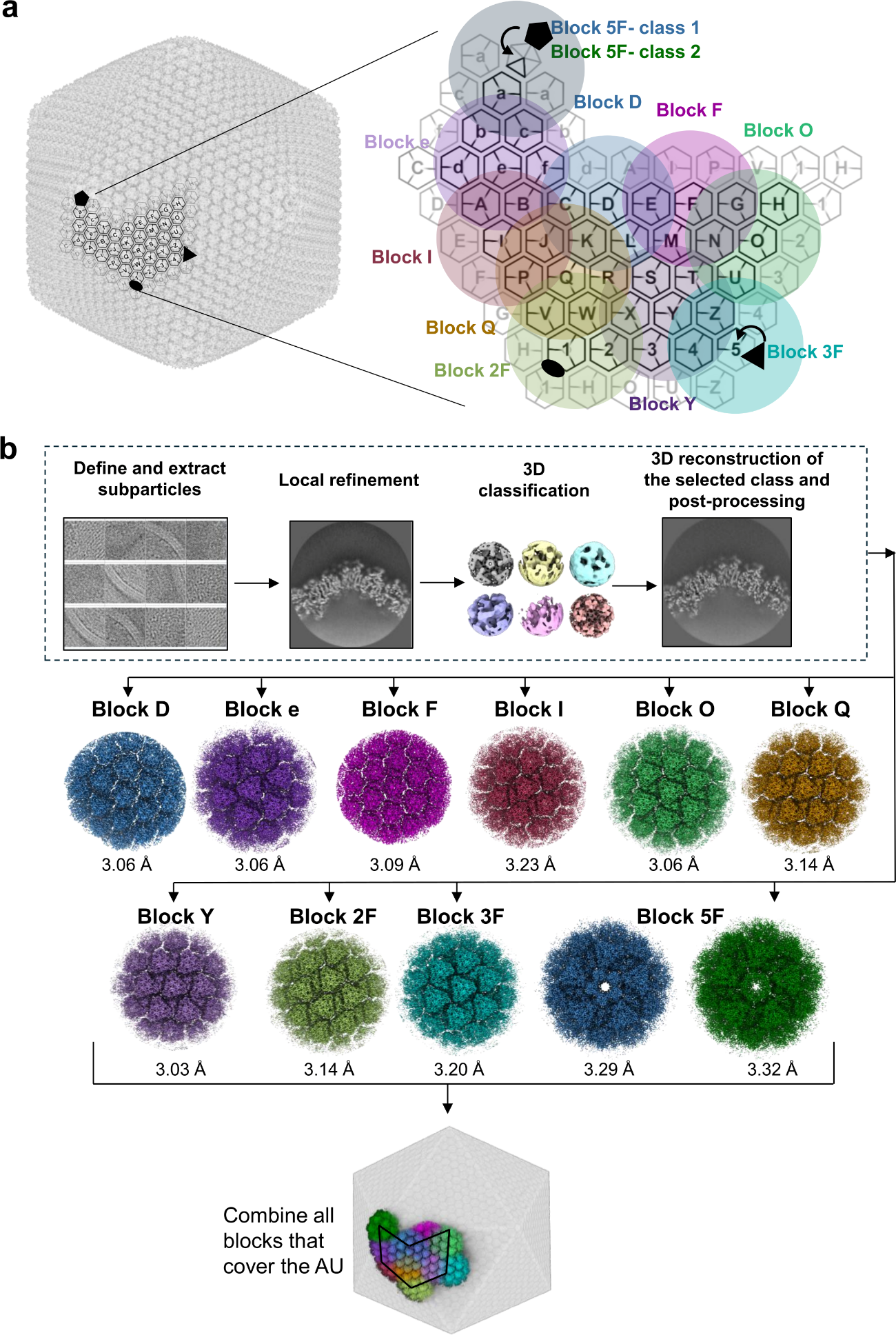

**Extended Data Fig. 2. Block-based reconstruction for PkV-RF02.** (**a**) Schematic representation of ten overlapping blocks that cover the AU. Blocks are denoted by the feature on which they are centred: capsomers D, e, F, I, O, Q, and Y; and the 3-fold (3F), 2-fold (2F) and 5-fold (5F) axes. (**b**) Block-based reconstruction workflow repeated for each of the 10 blocks. Fourier Shell Correlation (FSC) curves for each map are depicted in Supplementary Fig. 1. More details are summarized in Supplementary Table 2.

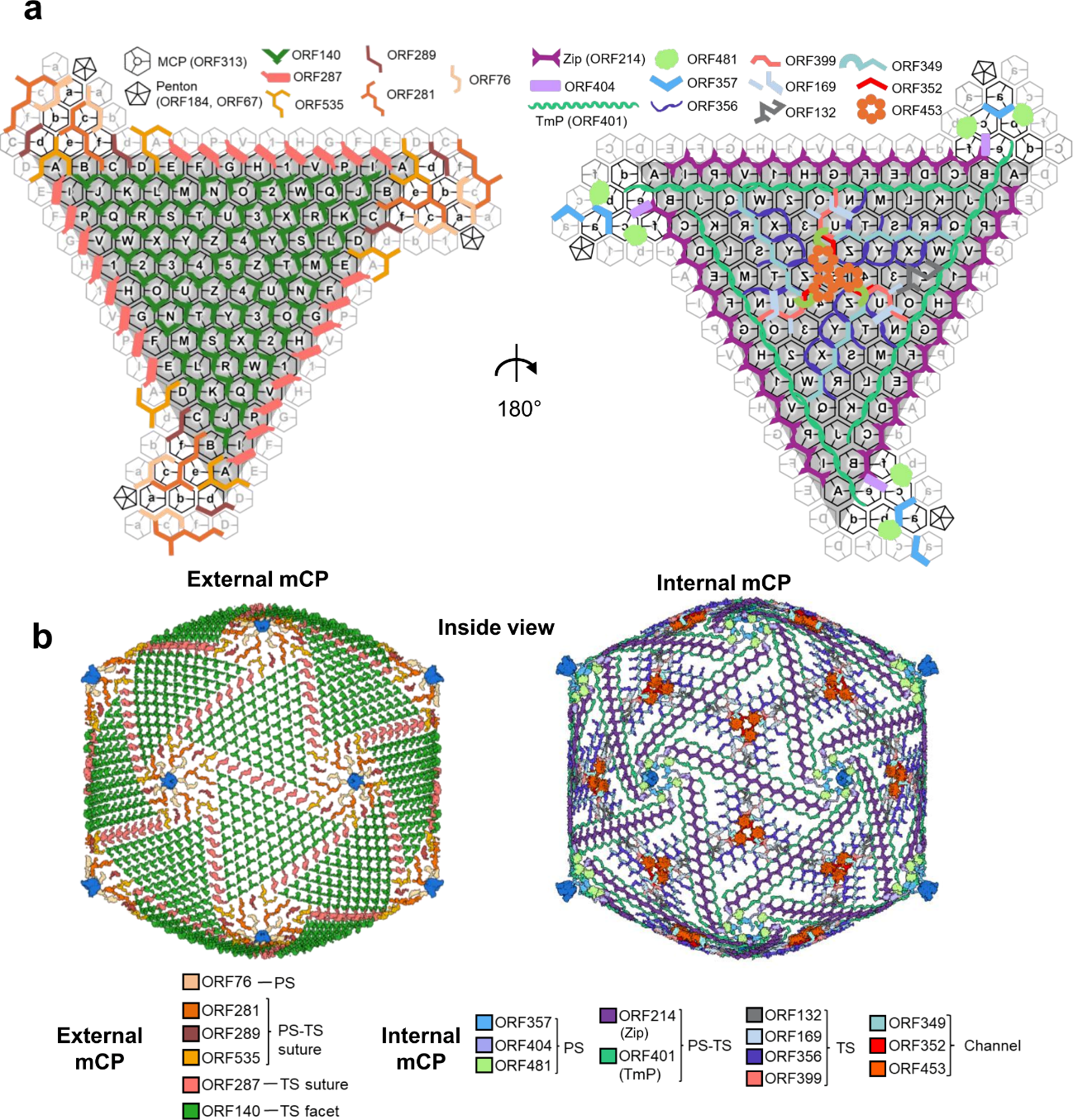

**Extended Data Fig. 3. Minor capsid protein organization of the giant virus PkV-RF02.** (**a**) Schematic organization of one icosahedral facet viewed from outside (left) and inside (right). Capsomers forming the trisymmetron are depicted as grey-filled hexamers, while capsomers in the pentasymmetron are white-filled. Capsomers in surrounding facets are contoured in light grey. Protein colours and shapes are indicated the legend. (**b**) Global view of PkV-RF02 mCP organization. MCP trimers have been computationally removed to show the external and internal mCP network. TS, trisymmetron. PS, pentasymmetron.

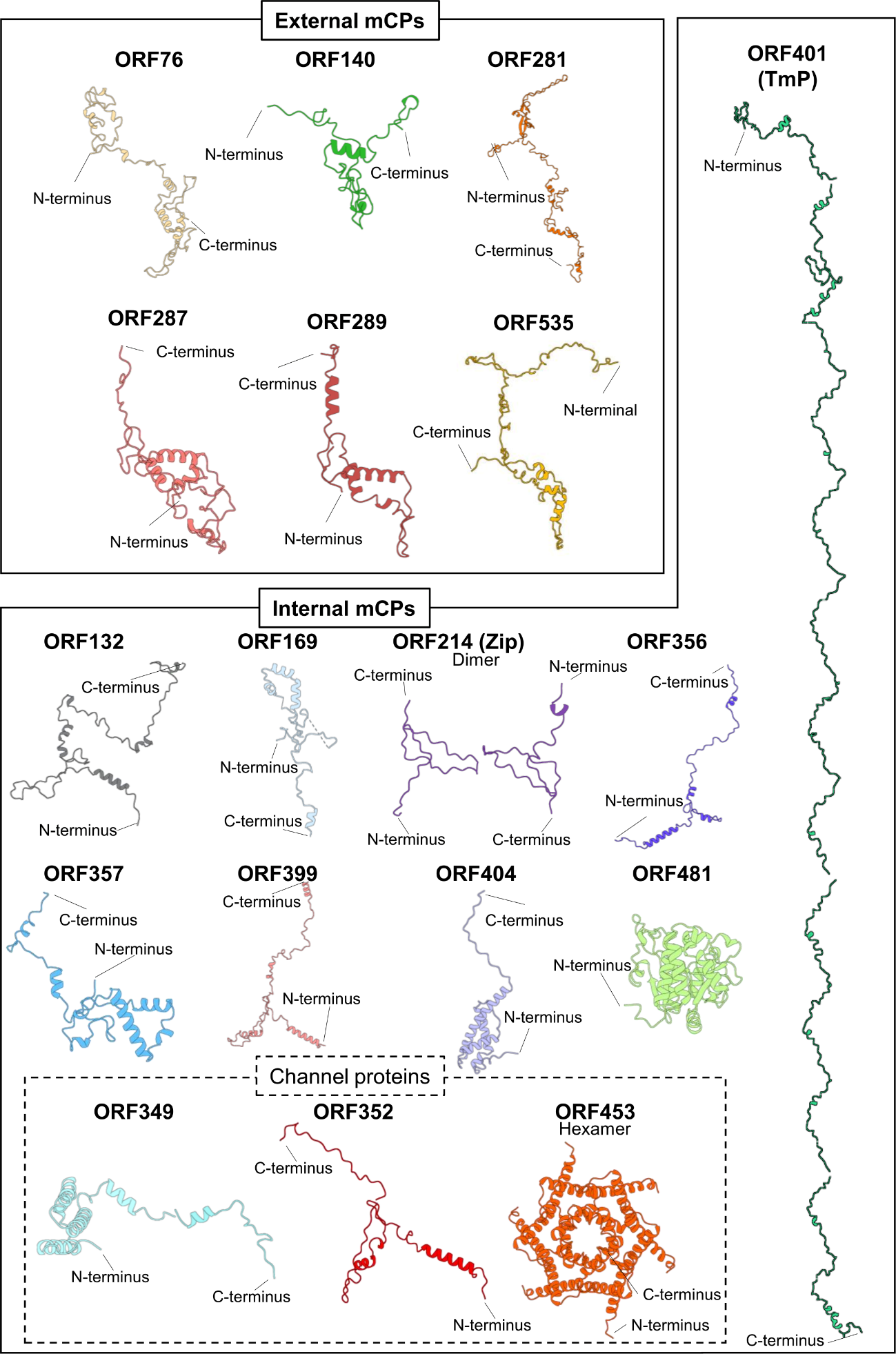

**Extended Data Fig. 4. Structures of PkV-RF02 mCPs.** In cases where the protein forms multimers, the multimeric assembly is displayed. Proteins are coloured as in Extended Figure 3.

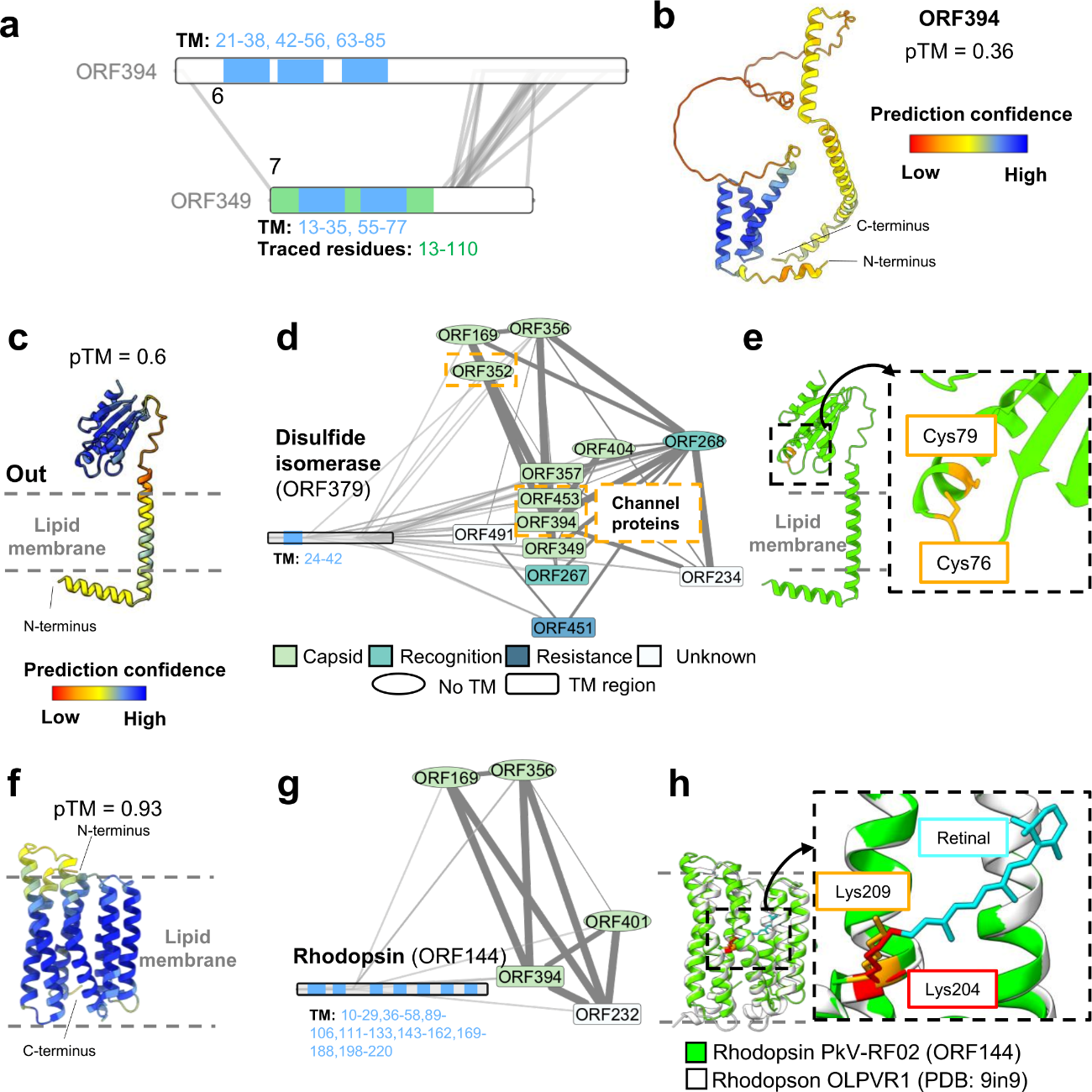

**Extended Data Fig. 5. AlphaFold predictions and XL-MS interaction networks for PkV-RF02 proteins.** (**a**) ORF394-ORF349 interaction network. Transmembrane regions coloured in blue. TM: transmembrane. Traced region in ORF349 is coloured in green. (**b**) AlphaFold predictions for ORF394, coloured by pLDDT. (**c**) AlphaFold prediction for disulphide isomerase (ORF379), coloured by pLDDT. (**d**) Disulphide isomerase (ORF379) interaction network within cluster 3. Channel proteins are indicated in orange. Transmembrane region (TM) coloured in blue. (**e**) Catalytic site of the disulphide isomerase. (**f**) AlphaFold prediction for rhodopsin-like protein (ORF144), coloured by pLDDT. (**g**) Rhodopsin-like protein (ORF144) interaction network within cluster 3. Transmembrane regions (TM) coloured in blue. (**h**) Structural alignment between rhodopsin ORF144 and rhodopsin OLPVR1. Interaction networks in (a), (d) and (g) generated in Cytoscape using XlinkCyNet plugin.

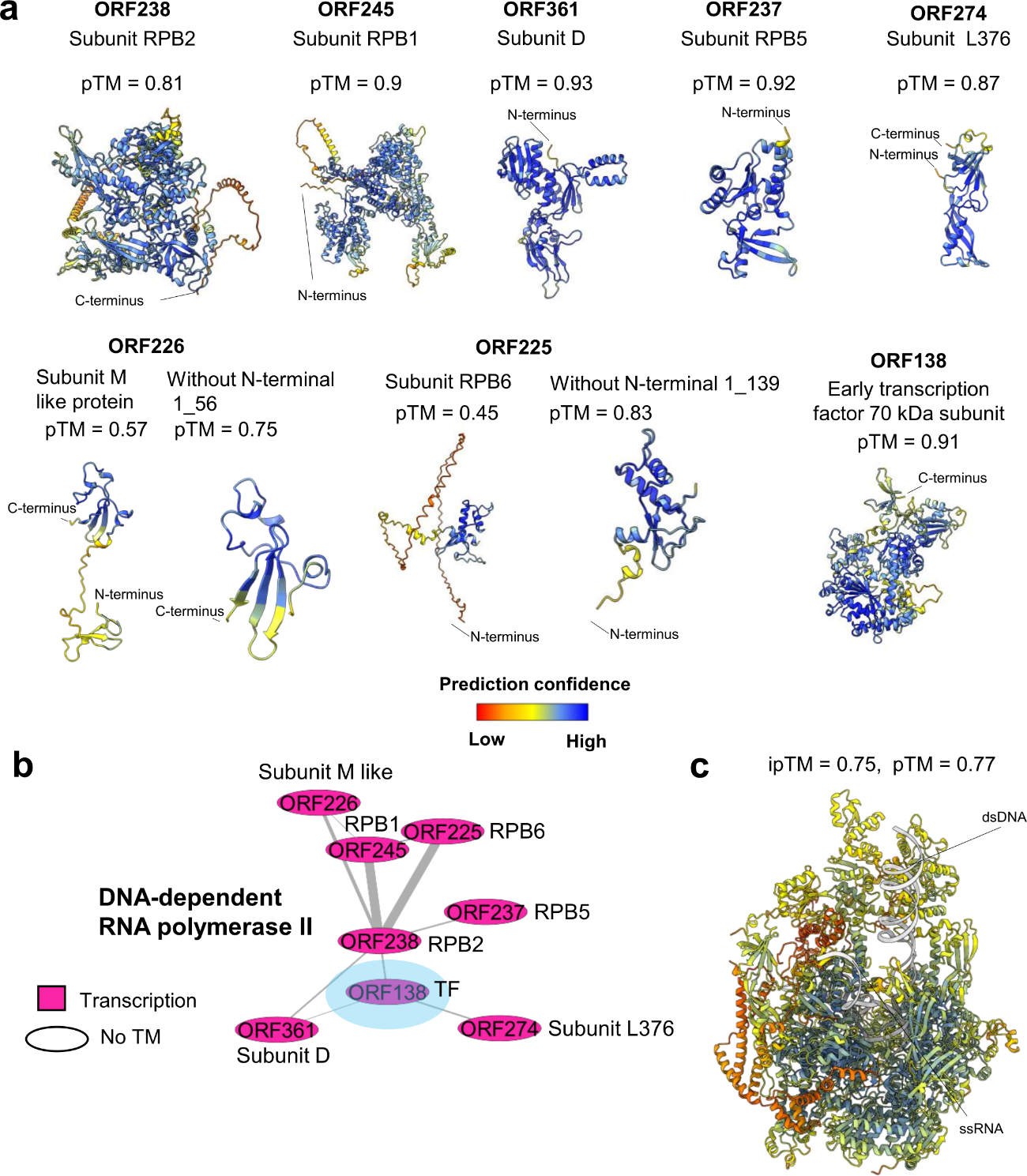

**Extended Data Fig. 6. DNA-dependent RNA polymerase complex components.** (**a**) AlphaFold predictions for the seven subunits of the DNA-dependent RNA polymerase present in the virion and transcription factor (TF) ORF318 coloured by pLDDT. (**b**) Cross-link network of proteins related to DNA-dependent RNA polymerase. Each subunit name is indicated on the side. (**c**) AlphaFold predictions for all the components of DNA-dependent RNA polymerase (b), in complex with dsDNA and ssRNA.

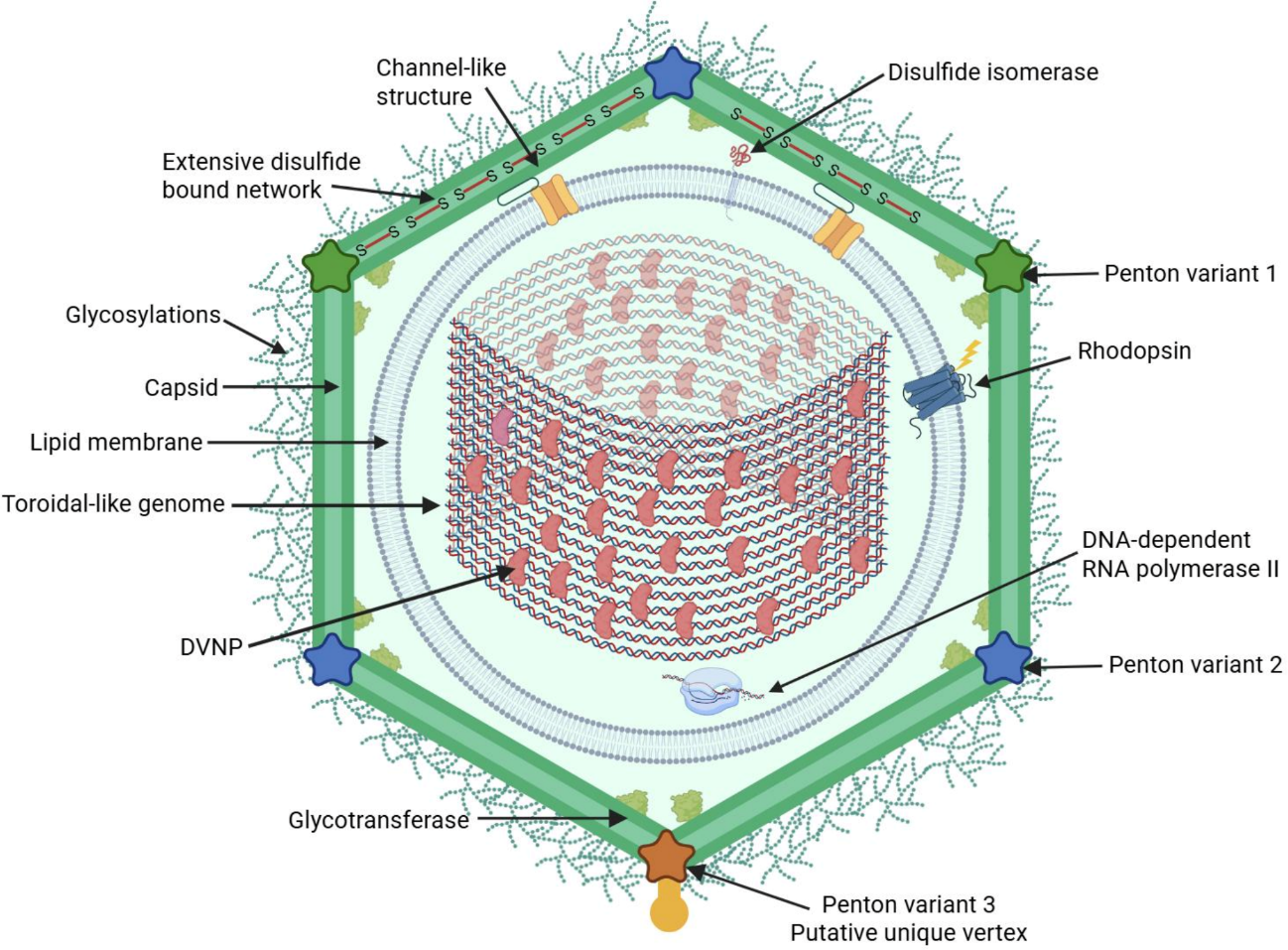

**Extended Data Fig. 7. Schematic organization of PkV-RF02.** Created by Biorender.com.

**Extended data tables**

**Extended Data Table 1. PkV-RF02 capsid proteins identified by the combination of cryo-EM and XL-MS.** Proteins are colour-coded by category.

| **Proteins** | **Description** | **Protein length (aa)** | **Multimerization** | **Chain identifier^*1^** | **Copies per AU^*2^** | **Copies**  **per virion** |
| --- | --- | --- | --- | --- | --- | --- |
| **Major capsid proteins** | | | | | | |
| **ORF313** | MCP variant 1 (MCPv1) | 525 | Trimer | [POS] + [1, 2 ,3] | 109 | 6540 |
| **ORF184** | Penton variant 1 (Pv1) | 316 | Pentamer | l1 | - | - |
| **ORF67** | Penton variant 2 (Pv2) | 447 | Pentamer | k1 | - | - |
| **External minor capsid proteins** | | | | | | |
| **ORF76** |  | 210 | Monomer | 76 | 1 | 30 |
| **ORF140** |  | 108 | Monomer | [o] + [POS] | 22 | 1320 |
| **ORF281** |  | 267 | Monomer | [u] + [POS] | 1 | 60 |
| **ORF287** |  | 167 | Monomer | [x] + [POS] | 4 | 240 |
| **ORF289** |  | 120 | Monomer | [v] + [POS] | 1 | 60 |
| **ORF535** |  | 225 | Monomer | [w] + [POS] | 1 | 60 |
| **Internal minor capsid proteins along the asymmetric unit suture** | | | | | | |
| **ORF214** | Zip protein | 169 | Dimer | [z] + [POS] | 14 | 840 |
| **ORF401** | Tape measure protein (TmP) | 608 | Monomer | [T] + [P, Q] | 1 | 60 |
| **Internal minor capsid proteins** | | | | | | |
| **Pentasymmetron** | | | | | | |
| **ORF357** | *Homologous to P9 of PBCV-1* | 282 | Monomer | 57 | 1 | 60 |
| **ORF404** | *Homologous to P8 of PBCV-1* | 166 | Monomer | [h] + [POS] | 1 | 60 |
| **ORF481** | Putative glycotransferase | 329 | Monomer | [g] + [POS] | 1 | 60 |
| **Trisymmetron** | | | | | | |
| **ORF132** |  | 164 | Monomer | [j] + [POS] | 1 | 60 |
| **ORF169** |  | 138 | Monomer | [n] + [POS] | 1 | 60 |
| **ORF356** |  | 190 | Monomer | [t] + [POS] | 4 | 240 |
| **ORF399** |  | 166 | Monomer | [m] + [POS] | 1 | 60 |
| **Membrane channel** | | | | | | |
| **ORF349** | Provides support to the membrane channel | 147 | Monomer | [p] + [POS] | 4 | 240 |
| **ORF352** | Between capsid and membrane channel | 119 | Monomer | [i] + [1, 2, 3] | 1 | 60 |
| **ORF453** | Membrane channel: creates the pore | 86 | Hexamer | [C] + [A, B, C] | 6 | 360 |
| **Identified by XL-MS^*3^** | | | | | | |
| **ORF394** | Membrane channel: next to ORF349 | 252 | Monomer | - | 1.35 | 80.9 |
| **ORF450** | MCP variant 2 (MCPv2) | 563 | Trimer | - | 0.7 | 44.1 |
| **ORF46** | Penton variant 3 (Pv3) | 402 | Pentamer | - | 0.16 | 10.1 |
| **ORF47** | Putative fibre-like protein | 1312 | Trimer (?) | - | 0.06 | 3.7 |

^*1^Chain identifiers are constructed using a two-character formula [Character 1] + [Character 2] to uniquely identify each chain in Supplementary Table 4. POS stands for capsomer position, see Extended Data Figure 1c.

^*2^AU stands for asymmetric unit. Copy numbers for traced proteins correspond to the number of molecules directly observed in the structural map, whereas copy numbers for proteins identified by XL-MS are estimated by bottom-up proteomics.

^*3^ These proteins are not included in the final PDB but are considered as capsid proteins based on XL-MS data.

**Extended Data Table 2. PTMs of traced proteins: O-linked glycosylations, disulphide bonds, and phosphorylations.** Non-bonded but closely positioned cysteines are highlighted in red.

| **Protein name** | **O-linked glycosylation** | **Intramolecular disulphide bond** | **Inter-molecular disulphide bond** | | **Phosphorylations** |
| --- | --- | --- | --- | --- | --- |
|  |  |  | **Residue 1** | **Residue 2** |  |
| **MCPv1** | Ser357, Ser366, Thr380, Ser382 | Cys174, Cys413 | Cys316 (MCPv1) | Cys165 (MCPv1) | Ser516, Ser524, Ser61, Tyr272, Tyr523, Tyr68 |
|  |  |  | Cys316 (MCPv1) | Cys316 (MCPv1) |  |
|  |  |  | Cys165 (MCPv1) | Cys165 (MCPv1) |  |
| **Pv1** | None | None | Cys145  (Pv1) | Cys316 (MCPv1) | None |
| **Pv2** | None | None | Cys293  (Pv2) | Cys316 (MCPv1) | None |
| **ORF76** | Ser37, Ser76, Ser82 | None | Cys34 (ORF76) | Cys165 (MCPv1) | None |
|  |  |  | Cys62 (ORF76) | Cys165 (MCPv1) |  |
|  |  |  | Cys86 (ORF76) | Cys316 (MCPv1) |  |
|  |  |  | Cys206 (ORF76) | Cys165 (MCPv1) |  |
| **ORF281** | Ser110, Ser174 | Cys188, Cys161 | Cys53 (ORF281) | Cys165 (MCPv1) | Ser55 |
|  |  |  | Cys106 (ORF281) | Cys165 (MCPv1) |  |
|  |  |  | Cys135 (ORF281) | Cys165 (MCPv1) |  |
|  |  |  | Cys178 (ORF281) | Cys165 (MCPv1) |  |
|  |  |  | Cys219 (ORF281) | Cys165 (MCPv1) |  |
|  |  |  | Cys225 (ORF281) | Cys165 (MCPv1) |  |
| **ORF535** | Ser118 | None | Cys75 (ORF535) | Cys316  (MCPv1) | None |
|  |  |  | Cys171 (ORF535) | Cys316 (MCPv1) |  |
|  |  |  | Cys184 (ORF535) | Cys165 (MCPv1) |  |
|  |  |  | Cys213 (ORF535) | Cys165 (MCPv1) |  |
|  |  |  | Cys225 (ORF535) | Cys316 (MCPv1) |  |
