## Supplementary Material for "Integrative structural biology of a giant marine virus reveals architectural and functional complexity beyond icosahedral symmetry"

**Short title:** Integrative structural biology of a T=219 virus

**The PDF file includes:**

**Supplementary notes 1 to 2**

**Supplementary tables 1 to 9**

**Supplementary figures 1 to 10**

**Supplementary references**

**Additional supplementary materials not included in this PDF**:

**Supplementary file 1 (.PDF). Validation reports for PkV-RF02 molecular models of each block.**

**Supplementary file 2 (.xlsx). Structural homology search results of PkV-RF02 proteins.** (a) ORF313 DALI results. (b) ORF313 Foldseek results. (c) ORF450 DALI results. (d) ORF450 Foldseek results. (e) ORF481 DALI results. (f) ORF481 Foldseek results. (g) ORF453 DALI results. (h) ORF453 Foldseek results. (i) ORF46 DALI results. (j) ORF46 Foldseek results. (k) ORF47 C-terminal domain 1 and 2 DALI results. (l) ORF47 C-terminal domain 1 and 2 Foldseek results. (m) ORF47 C-terminal domain 3 and 4 DALI results. (n) ORF47 C-terminal domain 3 and 4 Foldseek results. (o) ORF379 DALI results. (p) ORF379 Foldseek results. (q) ORF144 DALI results. (r) ORF144 Foldseek results.

**Supplementary file 3 (.xlsx). Copy number calculation for PkV-RF02 proteins.**

**Supplementary file 4 (.xlsx). XL-MS results of purified PkV-RF02 analysed by SCOUT.** (a) CSM from PkV-RF02 virions at 1% FDR. (b) Unique lysine-lysine connections of cross-linked proteins from PkV-RF02 virions at 1% FDR. (c) Protein-protein interactions from cross-linked PkV-RF02 virions at 1% FDR. (d) CysNet output from SCOUT to import to Cytoscape. (e) Clustering of proteins detected through XL-MS.

**Supplementary file 5 (.xlsx). Distance measurements of identified cross‑linked sites on PkV‑RF02 proteins.**

**Supplementary file 6 (.xlsx). Phosphoproteome results of PkV-RF02 analysed by FragPipe.** (a) Protein.csv results table. (b) Combined_site_STY_79.9663.csv results table. (c) Phosphorylation site count per protein. (d) Phosphoprotein counts per cluster. (e) Phosphoprotein counts per functional category.

**Supplementary Movie 1.** Structure of PkV-RF02.

### Supplementary Notes

**Supplementary note 1.**

**Statistical analysis of penton organization**

We aimed to determine whether the distribution of penton types across individual icosahedral viral particles is compatible with a model of random assignment. Specifically, we test two null hypotheses:

1. **Compositional Randomness (H₀)**: Penton types are randomly and independently assigned to each virus particle in proportion to their global frequencies.
2. **Positional Randomness (H₀)**: Penton positions in each virus particle are assigned independently, with no spatial clustering of same-variant pentons.

##### Experimental Framework

Using the experimental 3D classification of the cryo-EM subimages, each penton is labeled as Variant 1, Variant 2, or Other. The global frequencies are:

- Variant 1: 0.39
- Variant 2: 0.39
- Other: 0.22

In this study, we assume that all five monomers forming a penton are of the same variant as consistently indicated by the 3D classification results showing homogeneous penton composition. Consequently, each virus particle is represented by a 12-dimensional vector $v_{h}(i)$ where each element corresponds to a penton. The value of the *n*-th element of $v_{h}(i)$ is 1 if penton *n* is of Variant 1, and 0 otherwise. For example, a particle with the first four pentons of Variant 1 and the rest of other variants would be represented as v₁ = (1, 1, 1, 1, 0, 0, 0, 0, 0, 0, 0, 0).

Nonetheless, we acknowledge that the 3D classification process sometimes assigns the five copies of the same subimage-viewed from different orientations-to different classes. This behavior is illustrated in **Supplementary** **Figure 10a** which shows a histogram of subimage assignments per penton to each class, with a clear peak at 5.

##### Analysis of Compositional Randomness

To test if there is a tendency to the assembly of virus particles composed by a single penton variant versus a random distribution, we compare the observed distribution of Variant 1 penton counts per virion to the expected distribution under random assignment using a chi-squared goodness-of-fit test. Specifically, we evaluate the quantity $\left\| v_{h} \right\|$, which represents the number of Variant 1 pentons in each virus particle *h* (that is, $\left\| v_{h} \right\|=\sum_{i=1}^{12} v_{h}(i)$).

The test yielded:

- **Chi-square statistic**: 1894.22
- **p-value**: 0.00000000

This result indicates a statistically significant deviation from randomness, since very small p-values (e.g., < 0.05) mean that the observed distribution is unlikely under the null hypothesis, and larger chi-square values mean a greater discrepancy between observed and expected distributions.

However, a visual inspection of the histograms (see **Supplementary** **Figure 10b**) indicates that the deviation is subtle, primarily involving a slight increase in virus particles with Variant 1 counts near the mode and a decrease at the extremes.

Furthermore, the shape of the experimental histogram strongly disagrees with the hypothesis that a viral particle is composed mainly by pentons of a unique type, or the same number of pentons of Variant 1 than Variant 2, since this distribution would produce peaks in the histogram larger (and smaller) values.

##### Analysis of Positional Randomness

To investigate whether Variant 1 pentons exhibit spatial clustering within a virus particle, we use the 12x12 icosahedral adjacency matrix A, where $A\left( i,j \right)$ is defined as:

$$A(i,j)=\left\{ \begin{aligned} 1 \text{if pentons i and j are adjacent (i.e., connected by an edge of the icosahedron)} \\ 0 otherwise \end{aligned} \right.$$

Using this matrix, we define the clustering score $C_{h}$ for the virus particle h as:

$$C_{h}=v_{h}(i)A(i,j)v_{h}^{T}$$

Where $v_{h}^{T}$is the transpose of the vector $v_{h}$.

This score counts the number of adjacent Variant 1 penton pairs, which is equivalent to the number of edges in the icosahedral structure that connect two Variant 1 pentons.

**Monte Carlo Simulation**

To establish a baseline for clustering, we simulate virus particles $v_{h}$ by randomly assigning penton variants according to their empirical frequencies. Each simulation produces a clustering score, and the collection of these scores allows us to create a reference distribution against which observed clustering can be compared.

**Comparison with Experimental Data**

We compute clustering scores for each virus particle in the experimental dataset and compare the resulting histogram to the simulated baseline using a chi-squared test. The result:

- Chi-square statistic: 2423.86
- p-value: 0.00000000

indicates a statistically significant deviation from the null hypothesis. However, as shown in **Supplementary** **Figure 10c**, the deviation does not support intra-virion cooperativity. In particular the shape of the distribution is inconsistent with the hypothesis that pentons of the same variant tend to aggregate since this situation would produce a sharp peak at the histogram center.

**Supplementary note 2**

**Glycoproteomic analysis of purified PkV-RF02 particles**

We performed glycoproteomics analysis of PkV-RF02 virions to identify site-specific O-glycans. Samples were digested sequentially with ImpA and either trypsin or aspN. Only the aspN digest yielded glycopeptide identifications at the MCPv1 sites, owing to the lack of adjacent trypsin cleavage sites in proximity of the putative O-glycans. Moreover, we found no evidence of ImpA cleavage, indicating the nature of the O-linked glycans is rather distinct from mammalian mucin-type glycans that are required for cleavage by this enzyme. Inspection of the raw MS/MS spectra revealed a striking and recurring pattern of oxonium ions, compatible with the presence of HexNAc (204 Th), HexHexNAc (366 Th), as well as abundant peaks at 202, 188, 184, 170, 166, and 152 Th. The 202 Th oxonium ion is compatible with a glycan residue of C9H15N1O4, such as the rare methyl-N-acetylated viosamine (**Supplementary** **Fig. 8**). The additionally observed oxonium ions can be explained by collision induced loss of water (184, 166 Th), methyl (188 Th), or combinations thereof (170, 152 Th) from the MeVioNAc precursor. Alternatively, the oxonium ions at 188, 170, and 152 Th could also be explained by the presence of unmethylated VioNAc residues, but based on intact glycopeptide masses, this yielded glycan compositions incompatible with the observed pattern of oxonium ions (i.e. glycans without assigned MeVioNAc but nevertheless a prominent 202 Th peak). The most parsimonious explanation for the observed oxonium ions and intact glycopeptide masses is therefore glycans composed of HexNAc, Hex and MeVioNAc, although we do not want to rule out the presence of any other yet unidentified glycan residues. The identified glycopeptides from MCPv1 eluted as 4/5+ charged precursors within 45-50 minutes retention time, with intact glycopeptide masses of up to ca. 5600 Da. Accommodating for the mass of the underlying peptide backbone, the glycans at the combined T354/T356 or T380/S382 sites should be up to 2700 Da in mass. The peptide+HexNAc fragment was often observed in the fragmentation spectra, from which we deduce that the HexNAc residue is most likely to be directly linked to the T/S sites on MCPv1. We therefore completed a final glycopeptide search where we enumerated all possible glycans containing at least 2 HexNAc (one for each site on the glycopeptide), and at least 1 Hex and 1 MeVioNAc residue (to be compatible with the observed oxonium ions) of up to a mass of 2700 Da. This yielded 28 glycopeptide identification with scores above 400. The combined two glycosylation sites on each of the two glycopeptides spanning T354/T356 or T380/S382 consist of 8-13 residues, containing 2-7 HexNAc, 1-6 Hex, and 2-8 MeVioNAc residues. In addition, prominent oxonium ions composed of HexHexNAc, HexNAc2, and MeVioNAc2 with a methyl loss, were also observed. We conclude that the O-linked glycans at MCPv1 sites T354/T356/T380/S382 are each 4-7 residues in length, with HexNAc linked to the peptide, extended with additional HexNAc, Hex, and MeVioNAc residues, containing both HexNAc-HexNAc, Hex-HexNAc, and MeVioNAc-MeVioNAc linkages. We observed no MeVioNAc linkages to either Hex or HexNAc residues, leaving the MeVioNAc position in the glycan structure an open question.

### Supplementary Tables

**Supplementary Table 1. Cryo-EM vitrification conditions, data collection,** **image processing, reconstruction and refinement with icosahedral symmetry.**

| **Vitrification** |  |  |  |
| --- | --- | --- | --- |
| Grids | Quantifoil R1.2/1.3 or R0.6/1 Cu/Rh | | |
| Glow discharge | 25 mA, 1 minute | | |
| Temperature | 20ºC | | |
| Humidity | 95% | | |
| Blot time | 3 seconds | | |
| Force | -3, -4, -5 | | |
| **Data collection** | ***Data collection 1*** | ***Data collection 2*** | ***Data collection 3*** |
| Session DOI | 10.15151/ESRF-ES-1061180097 | 10.15151/ESRF-ES-925344867 | 10.15151/ESRF-ES-781088676 |
| Microscope | Titan Krios | Titan Krios | Titan Krios |
| Camera & acquisition mode | K3, counting mode | K3, counting mode | K3, counting mode |
| Voltage | 300 kV | 300 kV | 300 kV |
| Magnification | 64,000 x | 64,000 x | 64,000 x |
| Nominal pixel size | 1.35 Å/px | 1.35 Å/px | 1.35 Å/px |
| Exposure time | 3 s | 3 s | 2.5 s |
| Total dose rate | 22.9 e^-^/Å^2^ | 22.9 e^-^/Å^2^ | 23.6 e^-^/Å^2^ |
| Number of frames | 24 | 24 | 24 |
| Defocus range, step | -1.4 to -3.0 µm, 0.2 µm | -1.4 to -2.5 µm, 0.2 µm | -0.8 to -2.4 µm, 0.2 µm |
| Micrographs collected | 16,171 | 18,137 | 7,028 |
| Acquisition software | EPU | EPU | EPU |
| **Image processing** |  |  |  |
| Frame alignment software | MotionCor2, Scipion | | |
| CTF estimation software | CTFFIND4, Scipion | | |
| Particle picking software | Xmipp3, Manual picking, Scipion | | |
| Micrographs used | 41,336 | | |
| Particles picked | 71,231 | | |
| **Reconstruction by standard single particle analysis** | | | |
| Software | RELION, Scipion |  |  |
| Particles included | 71,231 |  |  |
| Symmetry imposed | Icosahedral |  |  |
| Rotational accuracy | 0.05 degrees |  |  |
| Translational accuracy | 0.4 pixels |  |  |
| B-factor applied | -182.63 Å^2^ |  |  |
| Final resolution  (gold standard FSC=0.143) | 4.7 Å |  |  |
| Reconstruction pixel size | 2 Å/px |  |  |

**Supplementary Table 2. Block-based reconstruction maps and refinement.**

|  | **D** | **e** | **F** | **I** | **O** | **Q** | **Y** | **2F** | **3F** | | **5F** | | |
| --- | --- | --- | --- | --- | --- | --- | --- | --- | --- | --- | --- | --- | --- |
|  |  |  |  |  |  |  |  |  |  |  | **Class 1** | | **Class 2** |
| **PDB entry** | 9THG | 9TH1 | 9TH0 | 9TGK | 9TGL | 9TGT | 9TH2 | 9TGI | 9TGH | | 9TM8 | | 9TH4 |
| **EMDB entry** | 55926 | 55914 | 55913 | 55902 | 55903 | 55911 | 55915 | 55901 | 55900 | | 56066 | | 55916 |
| **Map centred on capsomer** | D | e | F | I | O | Q | Y | 2-fold symmetry axis | 3-fold symmetry axis | | 5-fold symmetry axis  (2 classes) | | |
| **Sub-particles included** | 2,724,830 | 2,934,220 | 3,232,888 | 1,609,048 | 3,011,071 | 2,737,688 | 3,015,313 | 2,199,219 | | 4,273,860 | 1,670,434 | 1,630,503 | |
| **Sub-particles (%) over total** | 63.75% | 68.65% | 75.64% | 37.64% | 70.45% | 64.05% | 70.55% | 54.45% | | 100% | 39.08% | 38.15% | |
| **Symmetry imposed** | C1 | C1 | C1 | C1 | C1 | C1 | C1 | C1 | | C1 | C1 | C1 | |
| **B-factor applied** | -139.31 Å^2^ | -135.44 Å^2^ | -137.00 Å^2^ | -149.01 Å^2^ | -136.97 Å^2^ | -142.72 Å^2^ | -132.65 Å^2^ | -138.64 Å² | | -157.17 Å^2^ | -134.92 A² | -123.71 Å^2^ | |
| **Final resolution**  **(gold standard FSC = 0.143)** | 3.06 Å | 3.06 Å | 3.09 Å | 3.23 Å | 3.06 Å | 3.14 Å | 3.03 Å | 3.14 Å | | 3.2 Å | 3.29 Å | 3.32 Å | |
| **Pixel size** | 1.35 Å/px | | | | | | | | | | | | |

**Supplementary Table 3. Modelling and validation statistics for PkV-RF02 blocks.**

| **BLOCK** | | **5F class1** | **5F class2** | |  | **e** | | **I** | | **D** | | **F** | | **O** | | **Q** | | **Y** | | **2F** | | **3F** |
| --- | --- | --- | --- | --- | --- | --- | --- | --- | --- | --- | --- | --- | --- | --- | --- | --- | --- | --- | --- | --- | --- | --- |
| Software | |  | | Scipion, ModelAngelo, ChimeraX, Coot, Phenix | | | | | | | | | | | | | | | | | | |
| Chains | | 5 | 5 | |  | 31 | | 12 | | 22 | | 28 | | 20 | | 19 | | 32 | | 10 | | 11 |
| Atoms | | 15305 | 15929 | |  | 96071 | | 38758 | | 54855 | | 70744 | | 44449 | | 53180 | | 82617 | | 26997 | | 20144 |
| Residues | | 1942 | 2025 | |  | 12283 | | 4980 | | 7040 | | 9093 | | 5692 | | 6842 | | 10632 | | 3457 | | 2607 |
| Water | | 0 | 0 | |  | 0 | | 0 | | 0 | | 0 | | 0 | | 0 | | 0 | | 0 | | 0 |
| Ligands | | 0 | 0 | |  | 0 | | 0 | | 0 | | 0 | | 0 | | 0 | | 0 | | 0 | | 0 |
| Bond length (RMSD, Å) | | 0.0039 | 0.0043 | |  | 0.004 | | 0.004 | | 0.004 | | 0.003 | | 0.005 | | 0.005 | | 0.003 | | 0.004 | | 0.004 |
| Bond angles (RMSD, °) | | 0.93 | 0.95 | |  | 0.941 | | 0.931 | | 0.997 | | 0.918 | | 0.959 | | 0.969 | | 0.888 | | 0.929 | | 0.936 |
| MolProbity score | | 1.92 | 1.7 | |  | 1.63 | | 1.65 | | 1.16 | | 1.49 | | 1.61 | | 1.50 | | 1.50 | | 1.48 | | 1.60 |
| Clash score | | 6.78 | 6.97 | |  | 7.57 | | 7.44 | | 7.70 | | 6.67 | | 6.81 | | 6.65 | | 6.22 | | 6.32 | | 5.91 |
|  | **Ramachandran plot (%)** | | | | | | | | | | | | | | | | | | | | | |
| Outliers | | 0.00 | 0.00 | |  | 0.00 | | 0.00 | | 0.00 | | 0.01 | | 0.00 | | 0.00 | | 0.01 | | 0.00 | | 0.04 |
| Allowed | | 3.94 | 4.23 | |  | 3.32 | | 3.63 | | 3.34 | | 2.58 | | 3.22 | | 2.7 | | 2.81 | | 2.68 | | 3.25 |
| Favoured | | 96.06 | 95.77 | |  | 96.68 | | 96.37 | | 96.66 | | 97.41 | | 96.67 | | 97.3 | | 97.18 | | 97.21 | | 96.48 |
|  | **Rama-Z (Ramachandran plot Z-score, RMSD)** | | | | | | | | | | | | | | | | | | | | | |
| whole | | N=1930  1.45  (0.18) | N=2011  1.12  (0.18) | |  | N=12215  0.92  (0.07) | | N=4956  1.38  (0.11) | | N=6996  0.99  (0.10) | | N=9037  1.04  (0.08) | | N=5639  0.99  (0.11) | | N=6804  1.16  (0.10) | | N=10575  0.63  (0.08) | | N=3433  1.24  (0.14) | | N=2579  0.10  (0.17) |
| helix | | N=285  0.36  (0.32) | N=369  0.2  (0.29) | |  | N=2369  0.42  (0.11) | | N=944  1.05  (0.17) | | N=1116  0.23  (0.16) | | N=1664  0.70  (0.13) | | N=998  0.33  (0.17) | | N=1193  0.84  (0.15) | | N=2245  0.15  (0.12) | | N=643  0.92  (0.21) | | N=675  0.88  (0.21) |
| sheet | | N=485  0.72  (0.24) | N=462  0.46  (0.24) | |  | N=3089  0.38  (0.09) | | N=1233  0.54  (0.15) | | N=1716  0.64  (0.12) | | N=2025  0.43  (0.11) | | N=1224  0.57  (0.14) | | N=1644  0.67  (0.13) | | N=2322  0.22  (0.11) | | N=726  0.37  (0.19) | | N=536  0.08  (0.23) |
| loop | | N=1160  1.23  (0.17) | N=1180  1.17  (0.17) | |  | N=6757  0.71  (0.07) | | N=2779  1.00  (0.11) | | N=4164  0.74  (0.09) | | N=5348  0.74  (0.08) | | N=3417  0.74  (0.10) | | N=3967  0.73  (0.09) | | N=6008  0.52  (0.08) | | N=2064  0.93  (0.13) | | N=1368  0.56  (0.17) |
| **Rotamer outliers (%)** | | 2.31 | 1.08 | | 0.78 | | 0.81 | | 1.16 | | 0.91 | | 0.97 | | 1.20 | | 0.73 | | 0.68 | | 1.24 | |
| **Cβ outliers (%)** | | 0 | 0 | | 0 | | 0 | | 0 | | 0 | | 0 | | 0 | | 0 | | 0 | | 0 | |

**Supplementary Table 4.** PkV-RF02 proteins traced in the cryo-EM maps. Colour code as in Extended Data Table 1.

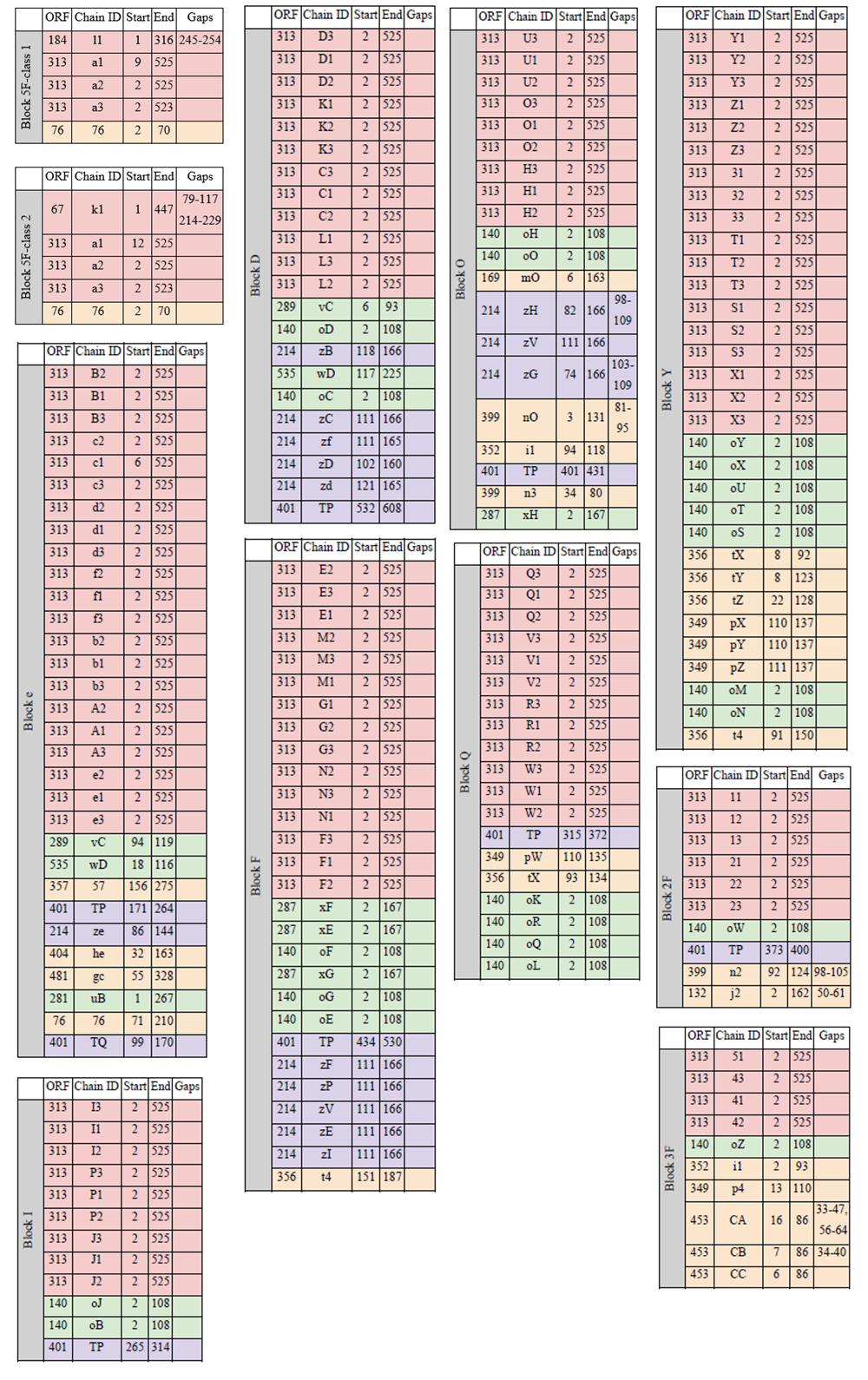

**Supplementary Table 5. RING analysis for π-π interactions in the MCP trimer.** Threshold distance settled manually at 5 Å.

| **Source** | **Target** | **Probability** |
| --- | --- | --- |
| Y1/36/Phe | Y3/26/Trp | 1 |
| Y1/97/Phe | Y1/124/His | 1 |
| Y1/124/His | Y1/333/Phe | 1 |
| Y1/278/Phe | Y1/504/Tyr | 1 |
| Y1/347/His | Y2/383/His | 1 |
| Y1/372/Phe | Y3/386/Phe | 1 |
| Y1/383/His | Y3/347/His | 1 |
| Y1/386/Phe | Y2/372/Phe | 1 |
| Y1/431/Phe | Y1/459/Tyr | 1 |
| Y2/26/Trp | Y3/36/Phe | 1 |
| Y2/97/Phe | Y2/124/His | 1 |
| Y2/124/His | Y2/333/Phe | 1 |
| Y2/278/Phe | Y2/504/Tyr | 1 |
| Y2/347/His | Y3/383/His | 1 |
| Y2/386/Phe | Y3/372/Phe | 1 |
| Y2/431/Phe | Y2/459/Tyr | 1 |
| Y3/97/Phe | Y3/124/His | 1 |
| Y3/124/His | Y3/333/Phe | 1 |
| Y3/278/Phe | Y3/504/Tyr | 1 |
| Y3/431/Phe | Y3/459/Tyr | 1 |

**Supplementary Table 6**. **PDBePISA^2^ analysis of PkV-RF02 MCP intra- and inter-capsomer interactions.** Interface area (Å^2^) is calculated as half the difference between the total accessible surface areas of the isolated and interacting structures. ΔiG represents the solvation-free energy gain upon interface formation, calculated as the difference in total solvation energies between the isolated and interacting structures. A negative ΔiG indicates favourable protein-protein interaction. N_HB_ indicates the number of potential hydrogen bonds formed across the interface. N_SB_ indicates the number of potential salt bridges across the interface. N_DS_ indicates the number of disulfide bounds across the interface. Notice that since PDBePISA considers interactions between atoms, some of the salt bridge atoms could correspond to the same residue. ORF450 corresponds to an AlphaFold model, not to an experimentally traced molecular model.

**Intra-capsomer contacts analysis**

|  | **Chain 1** | **Chain 2** | **Interface area (Å**^2^) | **Δ**^i^**G (kcal/mol)** | **N**_HB_ | **N**_SB_ | **N**_DS_ |
| --- | --- | --- | --- | --- | --- | --- | --- |
| **ORF313**  **capsomer**  **(MCPv1)** | Y1 | Y2 | 4933.2 | -55.9 | 65 | 8 | 1 |
|  | Y2 | Y3 | 4919.6 | -57.7 | 65 | 9 | 1 |
|  | Y1 | Y3 | 4855.4 | -55.6 | 62 | 2 | 1 |

|  | **Chain 1** | **Chain 2** | **Interface area (Å**^2^) | **Δ**^i^**G (kcal/mol)** | **N**_HB_ | **N**_SB_ | **N**_DS_ |
| --- | --- | --- | --- | --- | --- | --- | --- |
| **ORF450**  **capsomer**  **(MCPv2)** | A | B | 4978.8 | -46.3 | 69 | 7 | 0 |
|  | B | C | 4974.9 | -46.4 | 69 | 7 | 0 |
|  | A | C | 4973.6 | -46.1 | 70 | 7 | 0 |

**MCPv1 inter-capsomer contacts analysis**

|  | **Chain 1** | **Chain 2** | **Interface area (Å**^2^) | **Δ**^i^**G (kcal/mol)** | **N**_HB_ | **N**_SB_ | **N**_DS_ |
| --- | --- | --- | --- | --- | --- | --- | --- |
| **Back-to-front** | X3 | Y2 | 517.5 | 2.9 | 9 | 4 | 0 |
|  | X3 | Y1 | 416.3 | -3.5 | 3 | 1 | 1 |
|  | X3 | Y3 | 155.7 | -1.4 | 1 | 0 | 0 |

|  | **Chain 1** | **Chain 2** | **Interface area (Å**^2^) | **Δ**^i^**G (kcal/mol)** | **N**_HB_ | **N**_SB_ | **N**_DS_ |
| --- | --- | --- | --- | --- | --- | --- | --- |
| **Front-to-front** | A2 | e1 | 313.3 | -0.5 | 4 | 2 | 0 |
|  | A3 | e3 | 164.6 | 1.1 | 0 | 0 | 0 |
|  | A2 | e3 | 64.1 | -3.9 | 1 | 0 | 1 |
|  | A1 | e1 | 63.7 | 0.4 | 0 | 0 | 0 |
|  | A3 | e1 | 60.9 | 1.4 | 1 | 1 | 0 |
|  | A1 | e2 | 28.1 | -0.6 | 0 | 0 | 0 |

|  | **Chain 1** | **Chain 2** | **Interface area (Å**^2^) | **Δ**^i^**G (kcal/mol)** | **N**_HB_ | **N**_SB_ | **N**_DS_ |
| --- | --- | --- | --- | --- | --- | --- | --- |
| **Back-to-back** | B2 | E3 | 1276.1 | 2.8 | 17 | 6 | 1 |

**Supplementary Table 7**. **PDBePISA^2^ analysis of PkV-RF02 intra-penton and penton-MCP interactions.** For Pv1, PDBePISA results are presented separately for its two main regions: the main body and C-terminal domain.

|  | **Chain 1** | **Chain 2** | | **Interface area (Å**^2^) | **Δ**^i^**G (kcal/mol)** | **N**_HB_ | **N**_SB_ | **N**_DS_ |
| --- | --- | --- | --- | --- | --- | --- | --- | --- |
| **Pv1**  **ORF184**  (Main body) | J | I | | 2472.6 | -11.0 | 33 | 4 | 0 |
|  | I | H | | 2463.6 | -14.6 | 32 | 2 | 0 |
|  | J | F | | 2453.3 | -13.2 | 36 | 3 | 0 |
|  | G | F | | 2400.0 | -10.8 | 35 | 2 | 0 |
|  | H | G | | 2376.5 | -11.7 | 36 | 2 | 0 |
| Average: | | | | 2433.2 | -12.3 | 34 | 3 | 0 |
| **Pv1**  **ORF184**  (C-terminus) | J | H | | 275.0 | -3.8 | 2 | 0 | 0 |
|  | J | G | | 265.7 | -3.7 | 1 | 0 | 0 |
|  | I | | G | 264.7 | -4.1 | 2 | 0 | 0 |
|  | H | | F | 262.7 | -3.6 | 2 | 0 | 0 |
|  | I | | F | 244.1 | -3.2 | 2 | 0 | 0 |
| Average | | | | 262.4 | -3.7 | 2 | 0 | 0 |

|  | **Chain 1** | **Chain 2** | **Interface area (Å**^2^) | **Δ**^i^**G (kcal/mol)** | **N**_HB_ | **N**_SB_ | **N**_DS_ |
| --- | --- | --- | --- | --- | --- | --- | --- |
| **Pv2**  **ORF67** | D | B | 2034.2 | -17.9 | 25 | 5 | 0 |
|  | D | A | 2029.3 | -15.2 | 30 | 7 | 0 |
|  | E | C | 2016.7 | -13.4 | 27 | 5 | 0 |
|  | E | A | 1995.3 | -17.3 | 27 | 6 | 0 |
|  | C | B | 1980.5 | -16.9 | 30 | 7 | 0 |
| Average: | | | 2011.2 | -16.1 | 28 | 6 | 0 |

|  | **Chain 1** | **Chain 2** | **Interface area (Å**^2^) | **Δ**^i^**G (kcal/mol)** | **N**_HB_ | **N**_SB_ | **N**_DS_ |
| --- | --- | --- | --- | --- | --- | --- | --- |
| **MCPv1-Pv1**  **ORF313-ORF184** | a2 | l1 | 1503.2 | 0.4 | 17 | 9 | 0 |
|  | a3 | l1 | 228.5 | -3.6 | 1 | 0 | 0 |
|  | 76 | l1 | 143.9 | 0.5 | 0 | 0 | 0 |
|  | a1 | l1 | 35.3 | -0.8 | 0 | 0 | 0 |

|  | **Chain 1** | **Chain 2** | **Interface area (Å**^2^) | **Δ**^i^**G (kcal/mol)** | **N**_HB_ | **N**_SB_ | **N**_DS_ |
| --- | --- | --- | --- | --- | --- | --- | --- |
| **MCPv1-Pv2**  **ORF313-ORF67** | a2 | k1 | 1145.2 | -1.0 | 9 | 0 | 1 |
|  | a3 | k1 | 542.6 | -2.1 | 2 | 1 | 0 |
|  | a1 | k1 | 411.5 | 3.4 | 5 | 4 | 0 |

**Supplementary Table 8. Copy number estimation for selected PkV-RF02 proteins in clusters 3 and 4.** The full table is included as Supplementary File 3.

| **ORF number** | **Description** | | **Molecular weight (kDa)** | **Isoelectric point (pI)** | **Copy number per particle** |
| --- | --- | --- | --- | --- | --- |
| **Chromatin remodelling** | |  |  |  |  |
| ORF414 | Dinoflagellate/viral nucleoprotein (DVNP) | | 11.9 | 10.7 | 43.8 |
| **Transcription** |  |  |  |  |  |
| ORF138 | Early transcription factor 70 kDa subunit | | 98.4 | 8.48 | 110.9 |
| ORF238 | DNA-dependent RNA polymerase II subunit RPB2 | | 168.6 | 5.33 | 161.8 |
| ORF245 | DNA-dependent RNA polymerase II subunit RPB1 | | 173.1 | 6.7 | 133.4 |
| ORF361 | DNA-dependent RNA polymerase II subunit D | | 43.2 | 6.46 | 23.1 |
| ORF237 | DNA-dependent RNA polymerase II subunit RPB5 | | 24.9 | 6.6 | 17.6 |
| ORF274 | DNA-dependent RNA polymerase subunit L376 | | 18.4 | 5.24 | 11.7 |
| ORF226 | DNA-dependent RNA polymerase subunit M | | 15 | 7.15 | 10.1 |
| ORF225 | DNA-dependent RNA polymerase II subunit RPB6 | | 29.9 | 4.35 | 24.3 |
| ORF241 | DNA-dependent RNA polymerase, subunit N/RPB10 | | 9.1 | 8.7 | 7.5 |
| **Metabolism** |  |  |  |  |  |
| ORF379 | Disulphide isomerase | | 18.8 | 5.4 | 35.1 |
| ORF144 | Rhodopsinin | | 26.2 | 8.46 | 0.1 |
| ORF378 | ERV/ALR sulfhydryl oxidase | | 23.4 | 9.96 | 3.9 |

**Supplementary Table 9. Proteins encoded by PkV-RF02 related to glycosylation machinery^3^ and close homologs identified using BLASTp.** Description of proteins derived from^4^.

| **ORF number** | **Description** | **Organism** | **Accesion number** | **Query Coverage [%]** | **Identity [%]** | **E value** |
| --- | --- | --- | --- | --- | --- | --- |
| **ORF338** | Glucosamine 6-phosphate N-acetyltransferase | APMV | L316 | 86% | 38.93% | 4e-25 |
| **ORF342** | N-acetylglucosamine-1-phosphate uridyltransferase | APMV | R689 | 82% | 41.43% | 3e-48 |
| **ORF385** | Glutamine-fructose-6-phosphate transaminase | APMV,  PBCV-1 | L619,  A100R | 100%, 100% | 38.39%, 32.11% | 6e-121, 3e-72 |
| **ORF500** | Polysaccharide (O-antigen) biosynthesis protein | *Megavirus chilensis,*  *Moumouvirus australiensis* | mchi_528,  mc_458 | 100%, 99% | 53.40%, 54.35% | 6e-124,  4e-123 |
| **ORF501** | Methyltransferase | *M. chilensis,*  *M. australiensis* | mchi_529, mc_459 | 48%, 37% | 47.04%, 48.28% | 9e-74,  8e-68 |

### Supplementary Figures

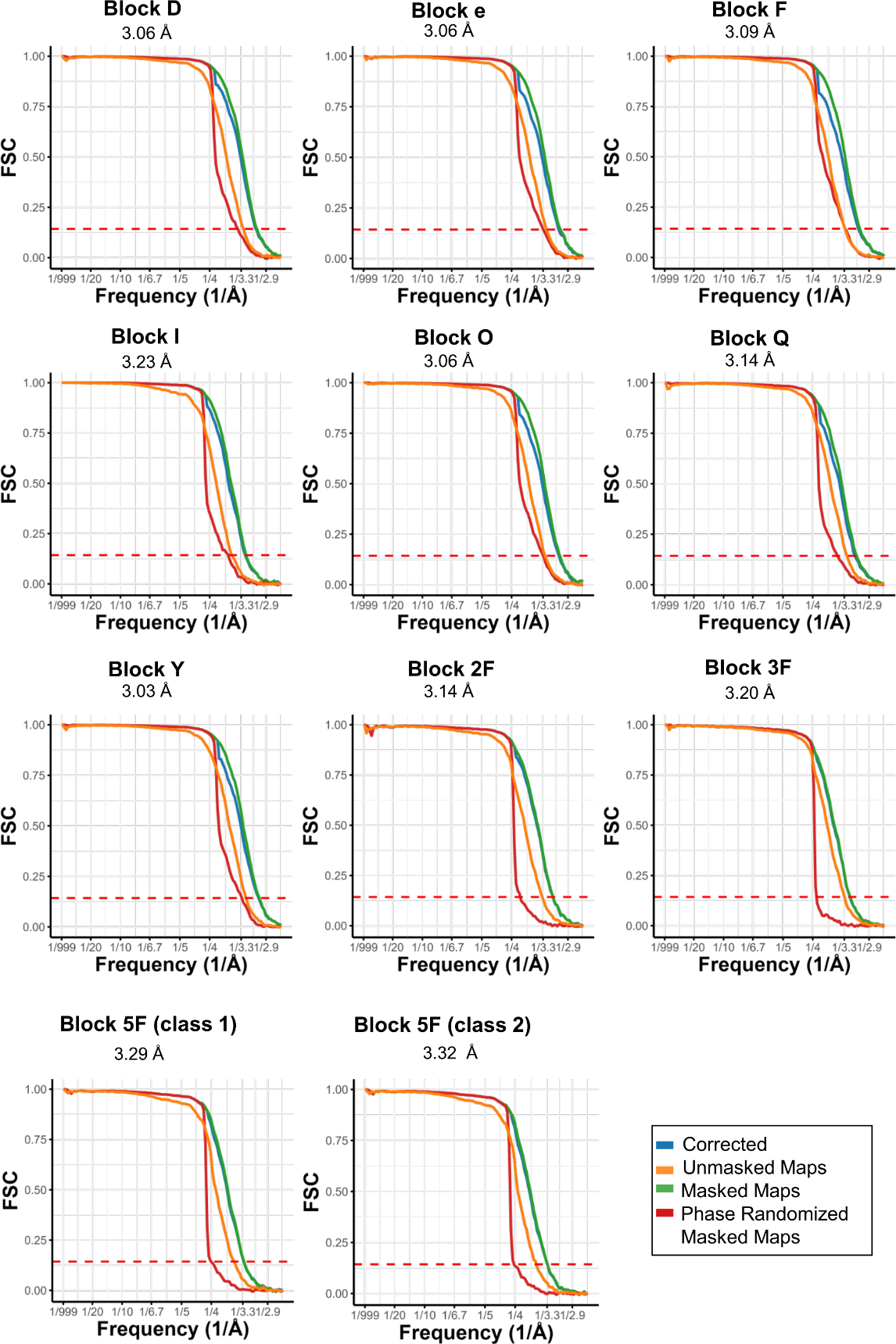

**Supplementary Figure 1. Fourier Shell Correlation (FSC) curves for each block in the block-based image processing workflow.**

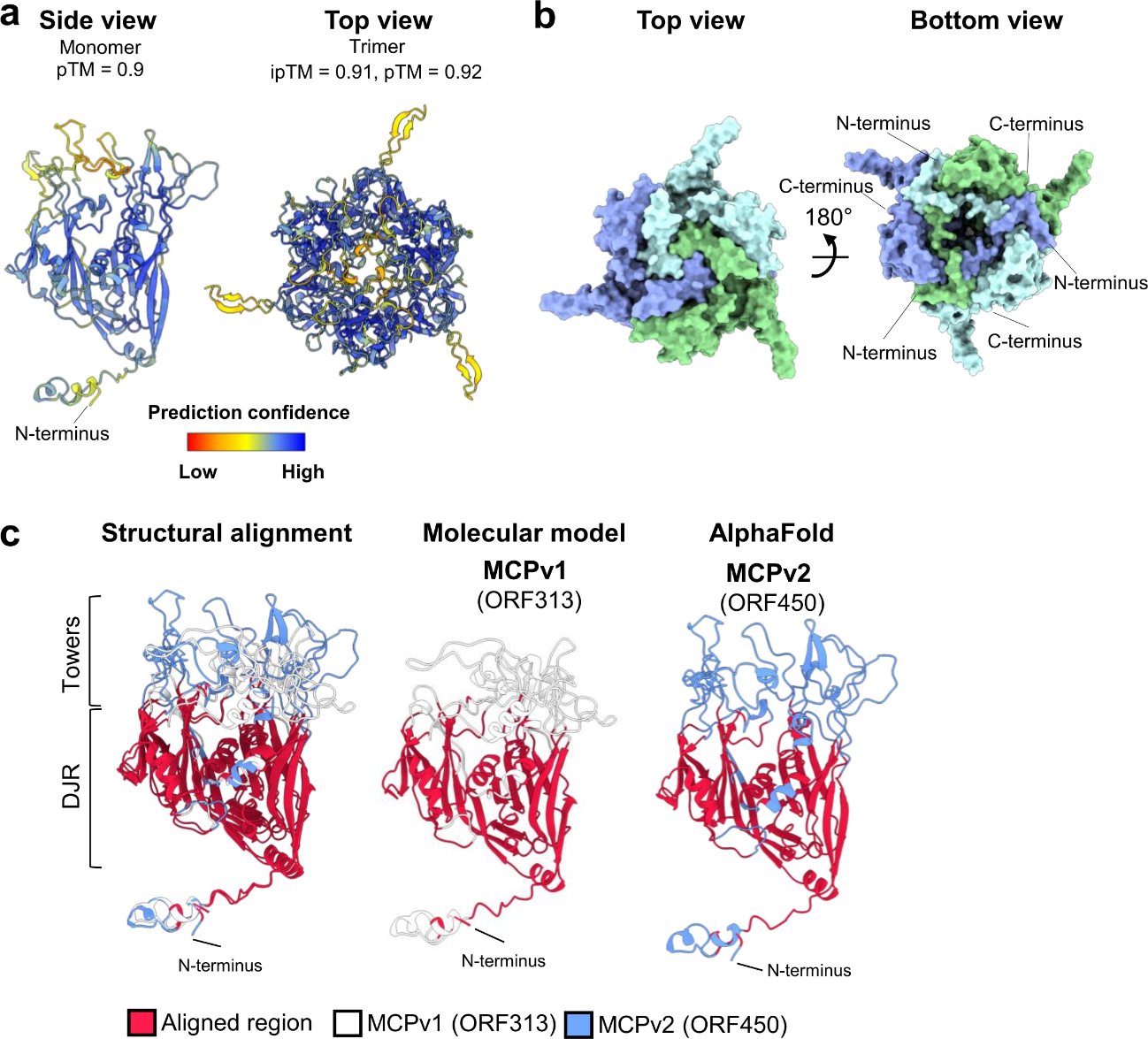

**Supplementary Figure 2. AlphaFold predictions for MCPv2 and structural alignment with MCPv1.** (**a**) MCPv2 (ORF450) AlphaFold prediction for the monomer (left) and trimer (right). Models are coloured by pLDDT. (**b**) MCPv2 trimer prediction coloured by chain. MCPv2 shares structural features with MCPv1, including a closed arrangement at the top of the capsomer and an N-terminal region that embraces adjacent hexamers. (**c**) Structural alignment of MCPv1 traced molecular model and AlphaFold predictions of MCPv2. The most notable difference lies in the tower domains, which appear less well defined in MCPv2. Additionally, MCPv2 does not seem to be capable of forming intra-capsomeric disulphide bonds, as shown by PDBePISA analyses of the trimer prediction (Supplementary Table 6). The AlphaFold model of MCPv2 was submitted to structural homology in DALI and Foldseek. As with MCPv1, the closest matches found were the MCP of EhV-201 (PDB: 8rbt), followed by that of PBCV-1 (PDB: 5tip, glycosylated MCP) (Supplementary File 2c-d).

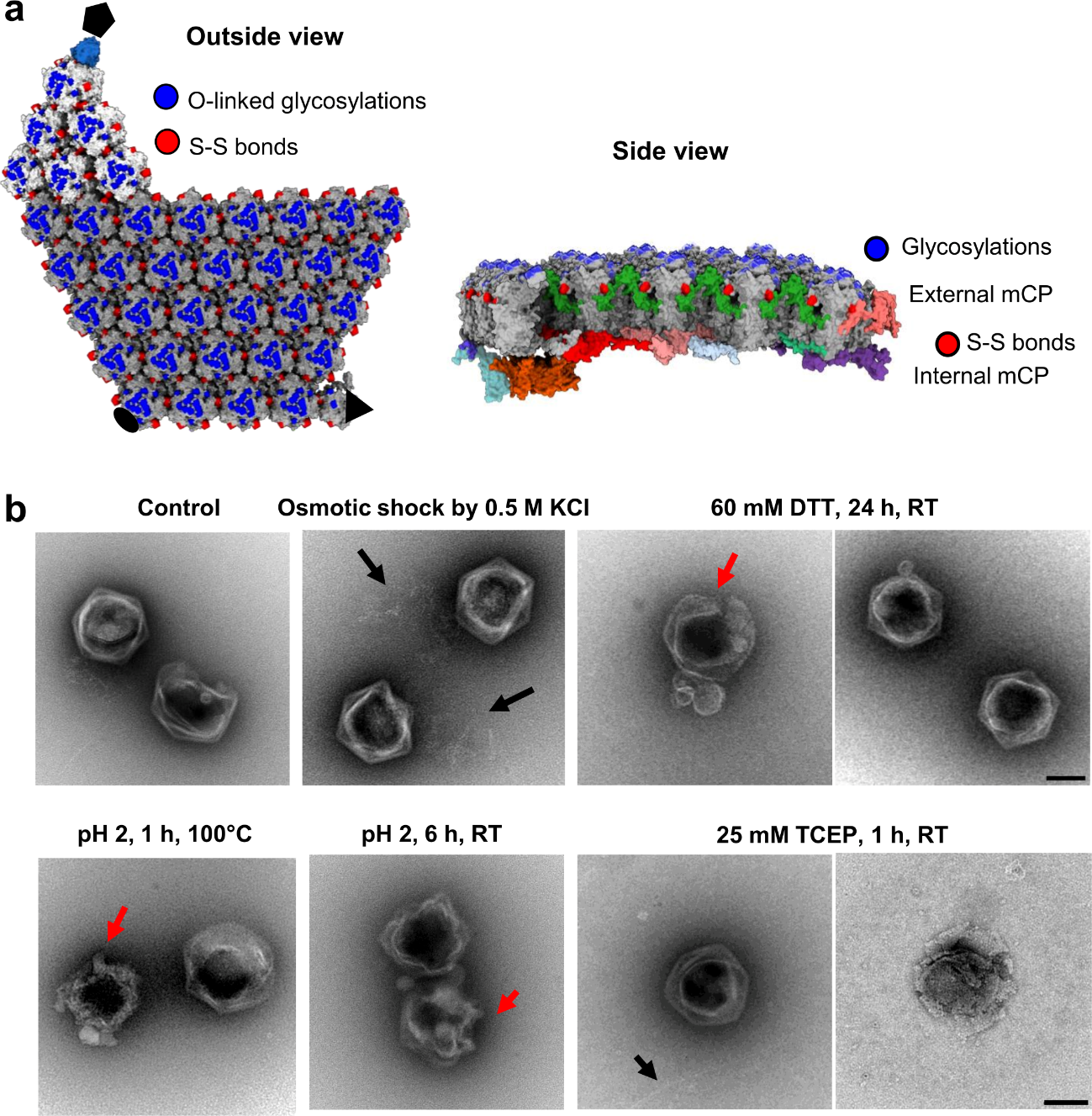
 **Supplementary Figure 3. PkV-RF02 multilayered capsid reinforcement by disulphide bond network and minor coat proteins.** (**a**) Left: top view of the asymmetric unit structure showing localization of inter-capsomer disulphide bonds and O-linked glycosylations. Right: side view of the capsid highlighting the multiple layers of stabilizing elements. Proteins coloured as in Extended Figure 3-4. (**b**) Purified PkV-RF02 particles subjected to chemical treatments visualized by negative staining. Capsids remained largely intact, with only partial opening observed in some particles (red arrows). In certain cases, fibres compatible with released genomes were visible in the background (black arrows). Scale bar represents 100 nm.

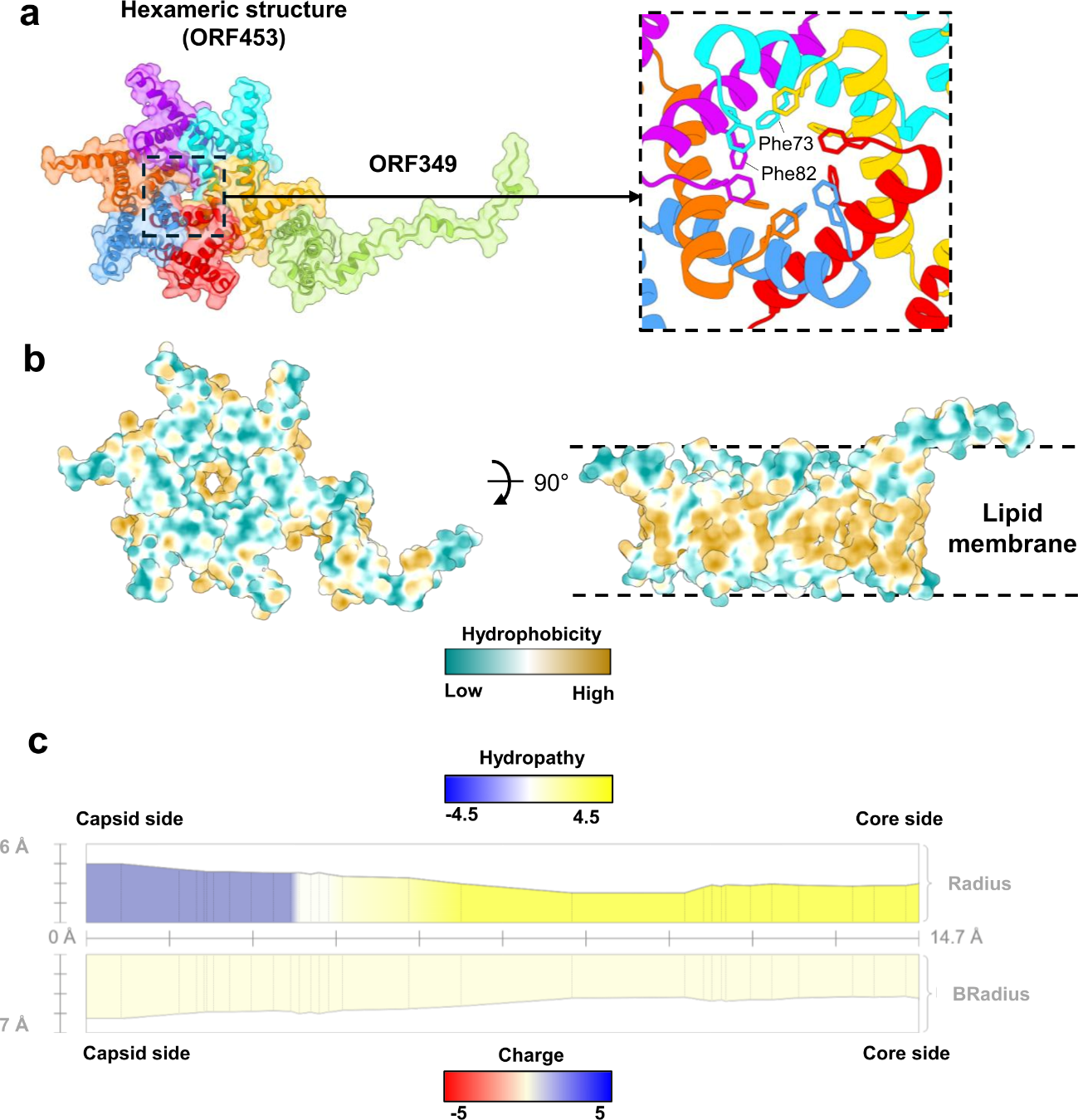

**Supplementary Figure 4. Properties of the PkV-RF02 membrane channel.** (**a**) Channel complex coloured by chain. The region enclosed by the dashed square is zoomed in at the right, showing phenylalanine residues facing the pore. (**b**) Top (left) and side (right) views of the channel structure coloured by surface hydrophobicity. (**c**) Channel profile by MOLE, indicating radius coloured by hydrophobicity and BRadius (calculated from B-factors stored within the structure) coloured by charge.

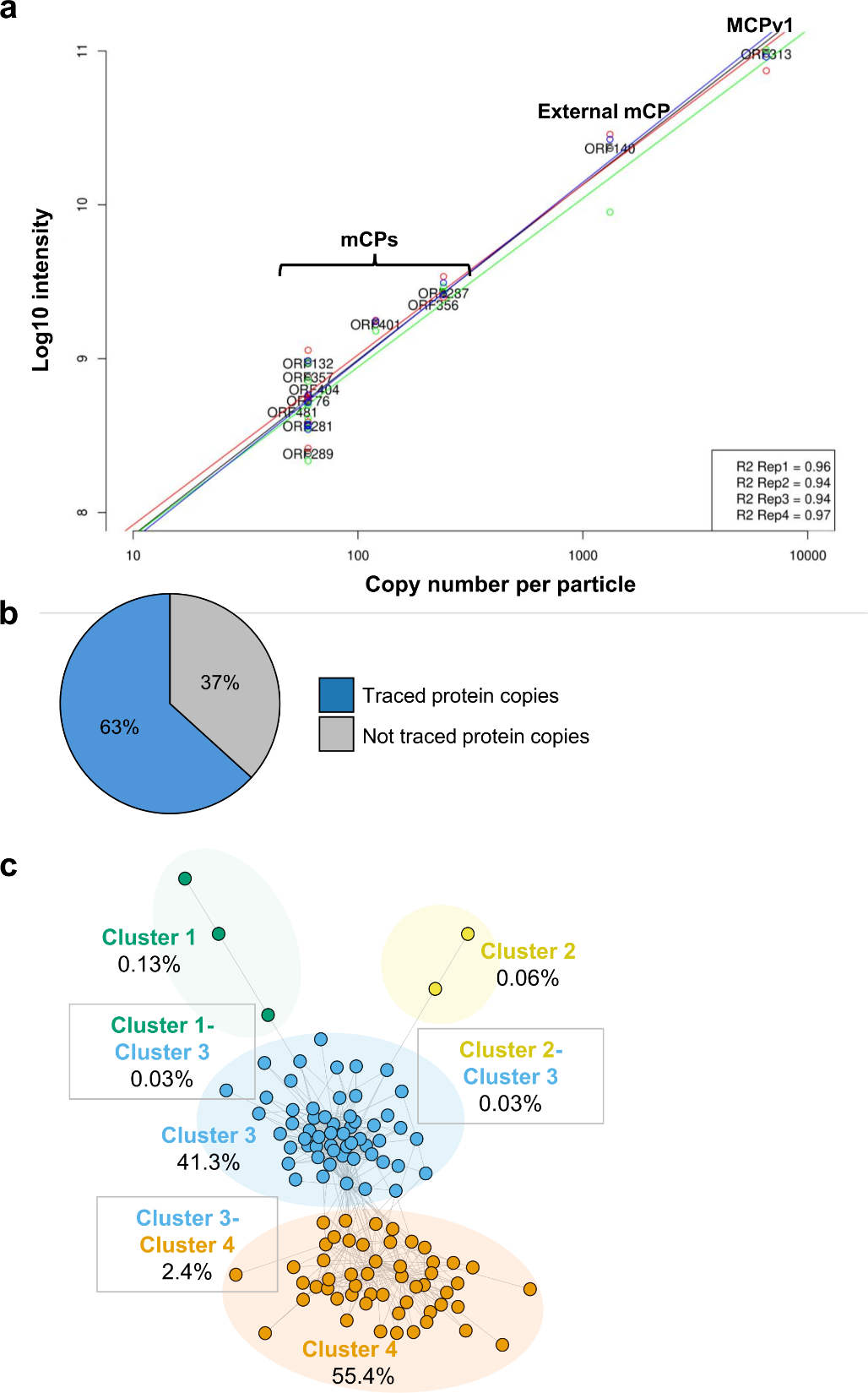

**Supplementary Figure 5. Protein copy number estimation and cross-link distribution within the viral particle.** (**a**) A linear regression model was established using the mass spectrometry signal intensity of shotgun proteomics experiment and capsid protein stoichiometry determined by cryo-EM. This linear relationship enables extrapolation of copy numbers for all other viral proteins based on their intensity values. (**b**) Proportion of traced *vs* non-traced proteins in the particle after copy number estimation. (**c**) Schematic representation of cross-link percentage per cluster pair.

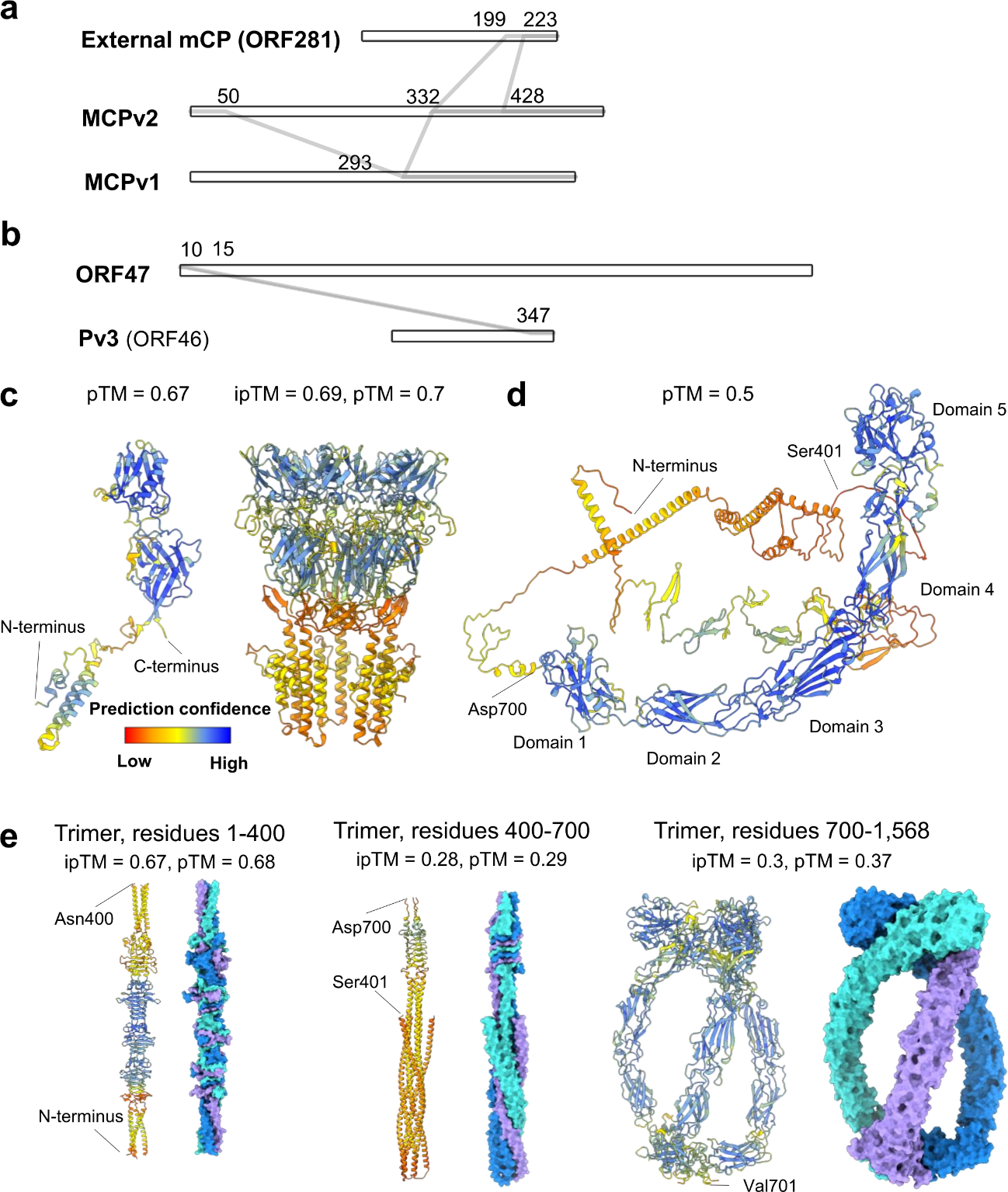

**Supplementary Figure 6. Interaction network for cluster 1 and 2 and cluster 2 components and AlphaFold predictions.** (**a**) Cross-link network of proteins in cluster 1, generated in Cytoscape using XlinkCyNet^5^ plugin. Cross-linked residue numbers are indicated on the top. (**b**) Cross-link interaction network within cluster 2. (**c**) AlphaFold predictions for ORF46 monomer (left) and pentamer (right), coloured by pLDDT. (**d**) AlphaFold predictions for ORF47 protein monomer, coloured by pLDDT. (**e**) AlphaFold predictions for ORF47 trimer for each of the three domains. Models are coloured by pLDDT (left) and by chain (right).

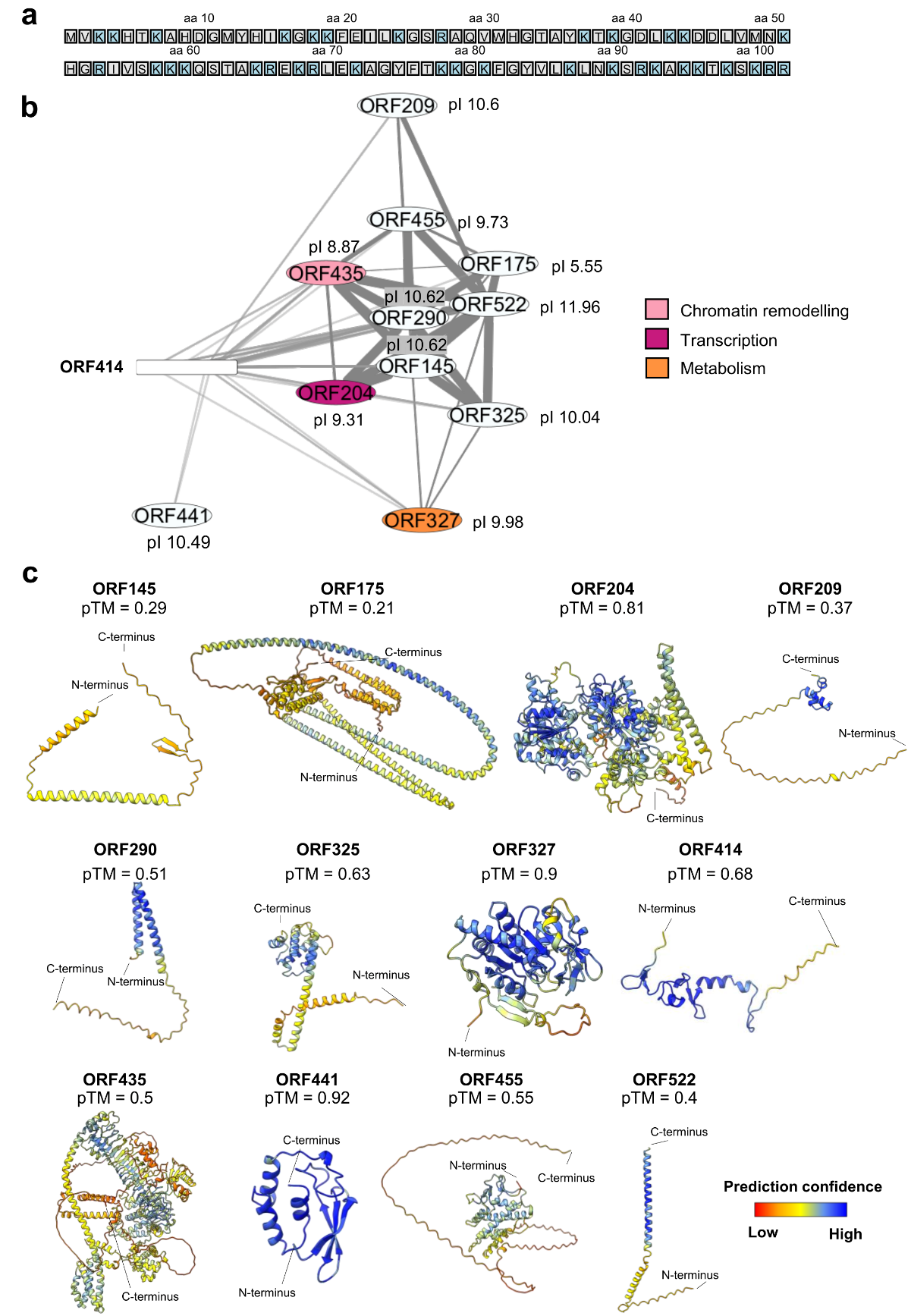

**Supplementary Figure 7. DVNP interactors and their AlphaFold predictions.** (**a**) DVNP (ORF414) sequence with positively charged residues highlighted in light blue. (**b**) Cross-link network of proteins interacting with DVNP (ORF414), generated in Cytoscape using XlinkCyNet^5^ plugin. Each protein isoelectric point (pI) is indicated on the top or side. (**c**) AlphaFold predictions for DVNP interactors, coloured by pLDDT.

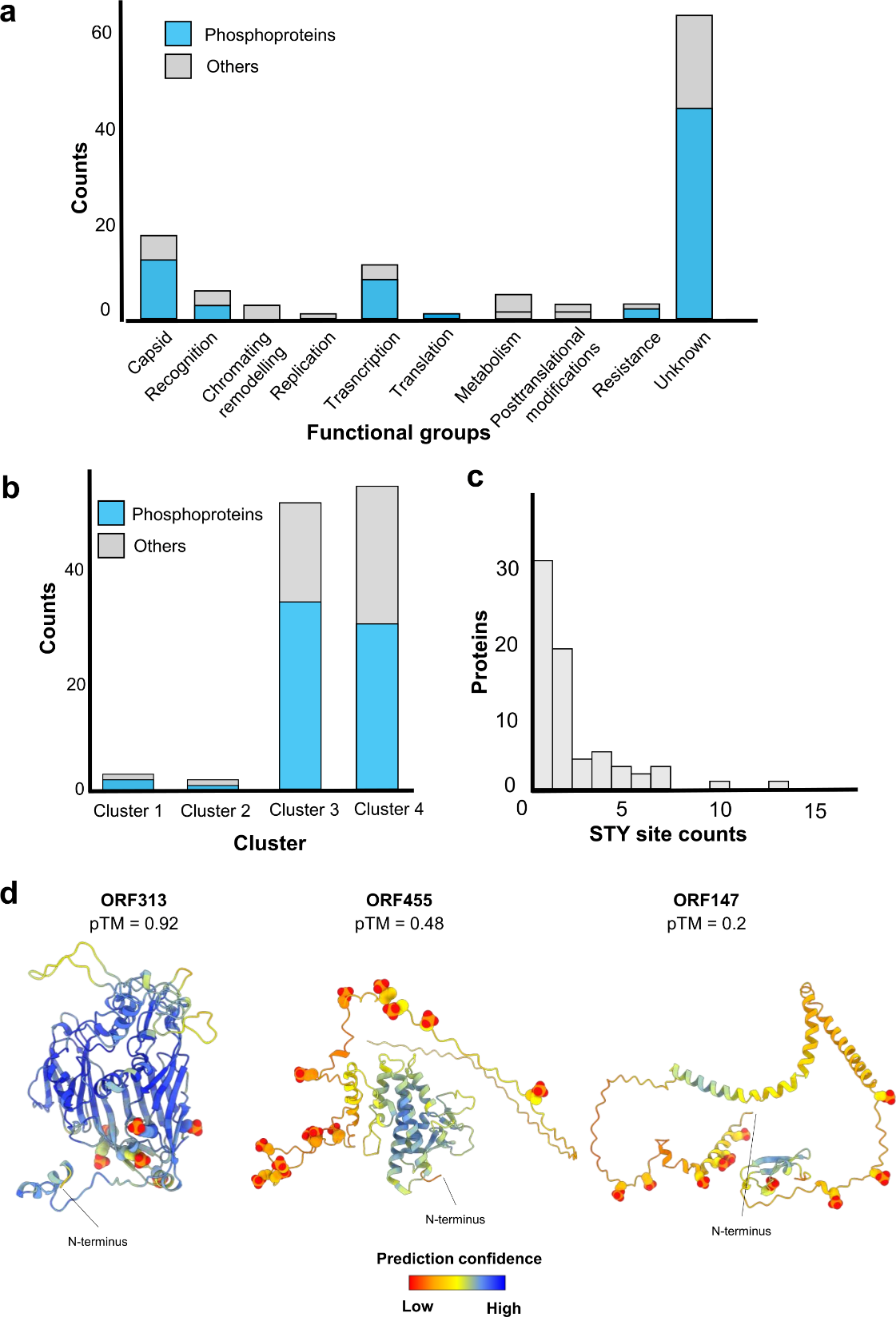

**Supplementary Figure 8. Phosphoproteome analysis of PkV-RF02.** (**a**) Distribution of phosphoproteins by functional categories. (**b**) Phosphoproteins identified within each of the four spatial clusters derived from XL-MS data. (**c**) Number of phosphorylation sites per protein, on serine, threonine and tyrosine (STY) residues. (**d**) AlphaFold predictions with phosphoryl groups at predicted phosphorylation sites. Phosphoryl groups are highlighted in orange-red, and models are coloured by pLDDT.

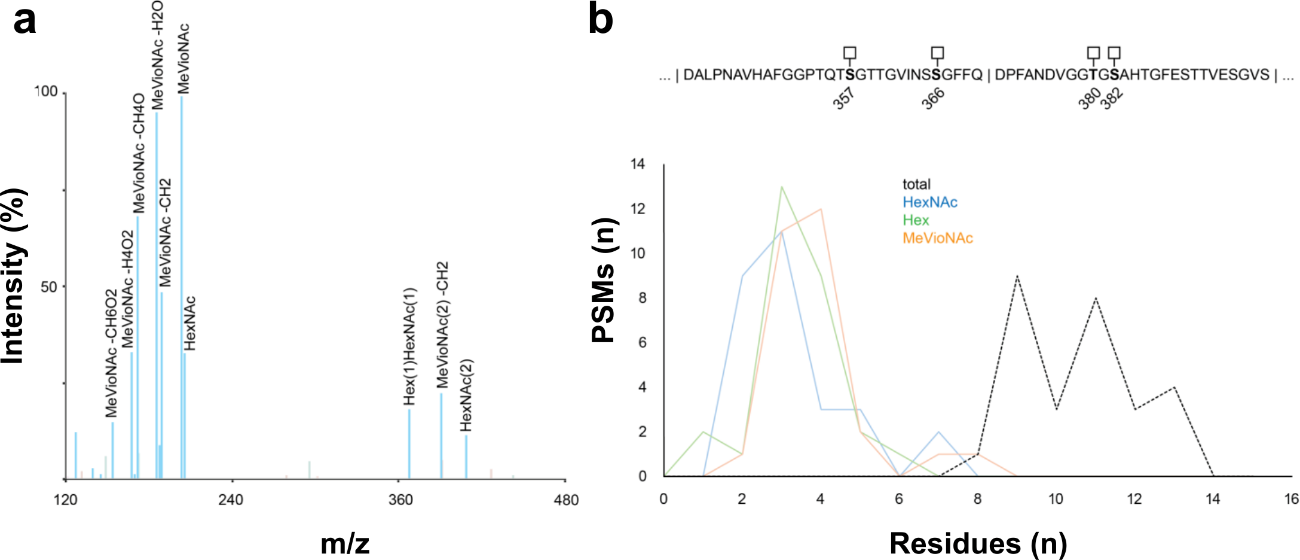

**Supplementary Figure 9.** (**a**) Representative MS/MS spectra showing pattern oxonium ions compatible with the presence of HexNAc (204 Th), HexHexNAc (366 Th), as well as abundant peaks at 202, 188, 184, 170, 166, and 152 Th. The 202 Th oxonium ion is compatible with a glycan residue of C_9_H_15_N_1_O_4_, such as the rare methyl-N-acetylated viosamine. The additionally observed oxonium ions can be explained by collision induced loss of water (184, 166 Th), methyl (188 Th), or combinations thereof (170, 152 Th) from the MeVioNAc precursor. Alternatively, the oxonium ions at 188, 170, and 152 Th could also be explained by the presence of unmethylated VioNAc residues, but based on intact glycopeptide masses, this yielded glycan compositions incompatible with the observed pattern of oxonium ions (i.e. glycans without assigned MeVioNAc but nevertheless a prominent 202 Th peak). (**b**) Glycopeptide identifications with Byonic scores >400, based on a search including all glycans up to 2700 Da containing at least 2 HexNAc, 1 Hex, and 1 MeVioNAc residue. The identified compositions show the total number of glycan residues and the corresponding numbers of HexNAc, Hex, and MeVioNAc units.

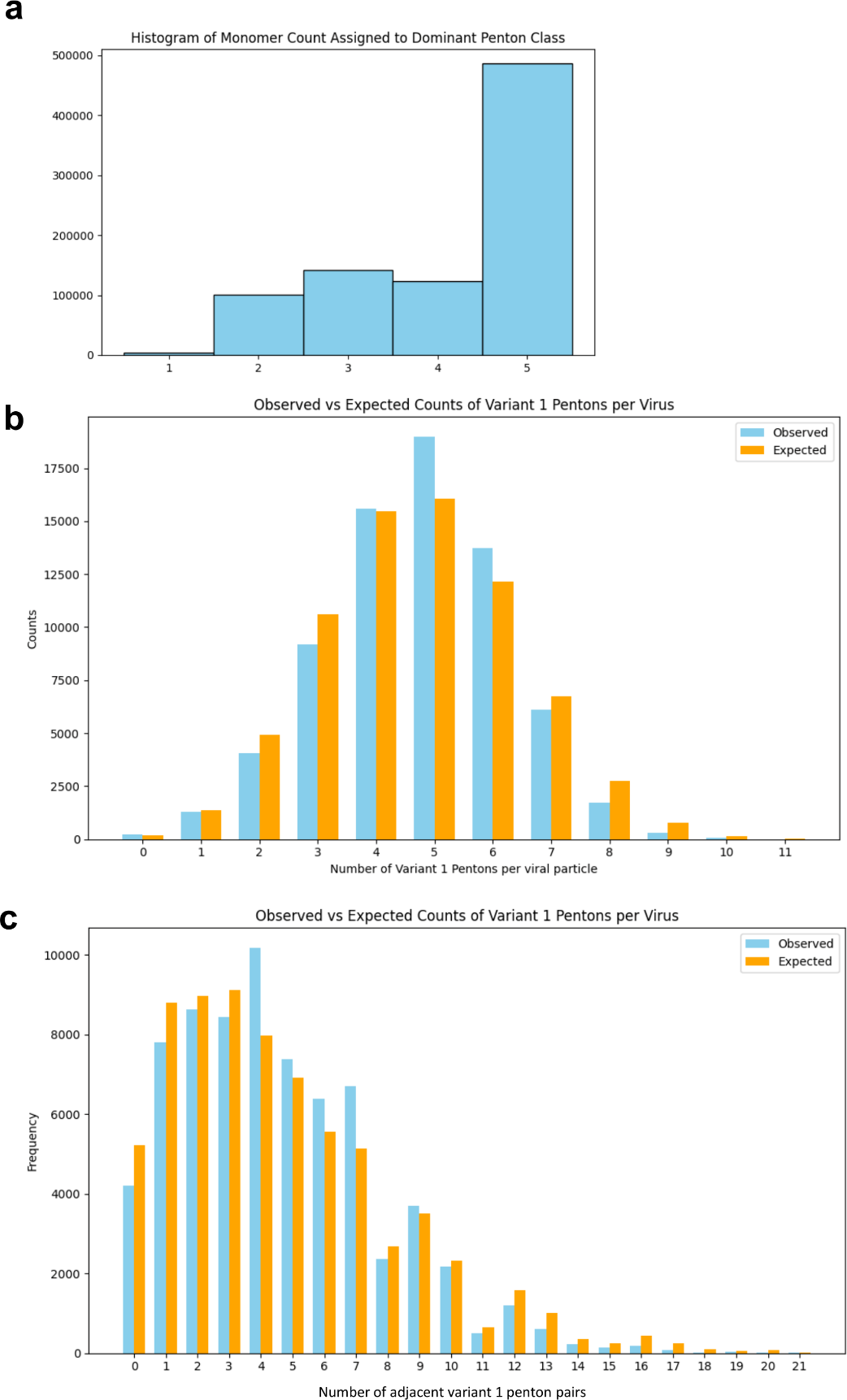

**Supplementary Figure 10. Classification and distribution analysis of penton variants across viral particles.** (**a**) Histogram of monomer count assigned to dominant penton classes. In the cryo-EM block-based reconstruction workflow, the subimages corresponding to the 5F block were classified into six groups. To generate the histogram, we computed the number of subimages assigned to the same class for each vertex. For vertices with subimages distributed across multiple classes, only the majoritarian class was considered. Therefore, the bar at position n in the histogram represents the number of vertices with *n* subimages belonging to the same class. It is important to note that approximately 20% of the subimages were assigned to classes unrelated to Variant 1 or Variant 2 pentons. As a result, we assume that the number of vertices with n = 5 is significantly underestimated due to noise in the data. (**b**) Observed versus expected counts of Pv1 per viral particle. Histograms compare the expected (computer-generated) and observed (experimental) distributions of $\left\| v_{h} \right\|.$ While the shapes of both distributions are visually similar, statistical analysis confirms a significant difference between them. Therefore, we may conclude that the penton variant distribution is close but not exactly what would be expected in a pure random distribution. Bins with less than 5 counts were excluded. (**c**) Comparison of expected (computer-generated) and observed (experimental) distributions of $C_{h}.$ While the shapes of both distributions are visually similar, statistical analysis confirms a significant difference between them. Importantly, the presence of variant *n* penton clusters would be expected to produce peaks at intermediate values, which are not observed. Bins with less than 5 counts were excluded.
