## Supplementary file 1 for "Integrative structural biology of a giant marine virus reveals architectural and functional complexity beyond icosahedral symmetry": Block_e_D_1292152363_val-report-full-annotate_P1.pdf

### Full wwPDB EM Validation Report (i)

Dec 3, 2025 – 10:12 am GMT

| | | |
|---|---|---|
| PDB ID | : | 9TH1 / pdb_00009th1 |
| EMDB ID | : | EMD-55914 |
| Title | : | Structure of PkV-RF02: block e |
| Deposited on | : | 2025-12-02 |
| Resolution | : | 3.06 Å(reported) |

**This wwPDB validation report is for manuscript review**

This is a Full wwPDB EM Validation Report.

This report is produced by the wwPDB biocuration pipeline after annotation of the structure.

We welcome your comments at **
A user guide is available at
https://www.wwpdb.org/validation/2017/EMValidationReportHelp
with specific help available everywhere you see the (i) symbol.

The types of validation reports are described at
http://www.wwpdb.org/validation/2017/FAQs#types.

For Manuscript Review

---

The following versions of software and data (see references (i)) were used in the production of this report:

| | | |
|---|---|---|
| EMDB validation analysis | : | 0.0.1.dev129 |
| MolProbity | : | 4-5-2 with Phenix2.0 |
| Percentile statistics | : | 20231227.v01 (using entries in the PDB archive December 27th 2023) |
| EM percentile statistics | : | 202505.v01 (Using data in the EMDB archive up until May 2025) |
| MapQ | : | 1.9.13 |
| Ideal geometry (proteins) | : | Engh & Huber (2001) |
| Ideal geometry (DNA, RNA) | : | Parkinson et al. (1996) |
| Validation Pipeline (wwPDB-VP) | : | 2.46 |

### 1 Overall quality at a glance (i)

The following experimental techniques were used to determine the structure:
*ELECTRON MICROSCOPY*

The reported resolution of this entry is 3.06 Å.

Percentile scores (ranging between 0-100) for global validation metrics of the entry are shown in the following graphic. The table shows the number of entries on which the scores are based.

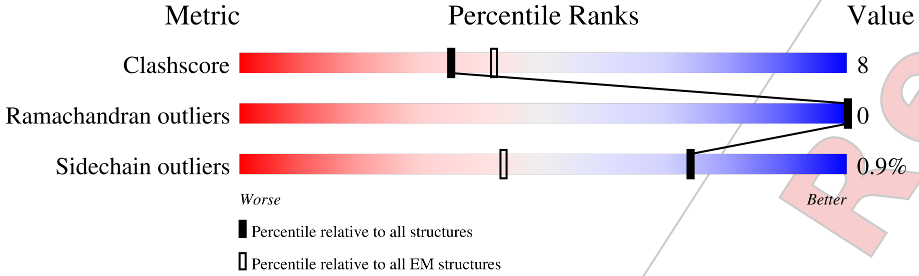


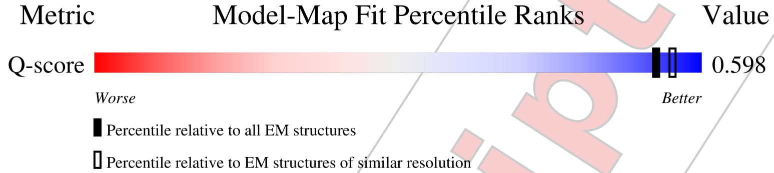


| Metric | Whole archive (#Entries) | EM structures (#Entries) | Similar EM resolution (#Entries, resolution range(Å)) |
|---|---|---|---|
| Clashscore | 210492 | 15764 | - |
| Ramachandran outliers | 207382 | 16835 | - |
| Sidechain outliers | 206894 | 16415 | - |
| Q-score | - | 25397 | 13976 ( 2.56 - 3.56 ) |

The table below summarises the geometric issues observed across the polymeric chains and their fit to the map. The red, orange, yellow and green segments of the bar indicate the fraction of residues that contain outliers for >=3, 2, 1 and 0 types of geometric quality criteria respectively. A grey segment represents the fraction of residues that are not modelled. The numeric value for each fraction is indicated below the corresponding segment, with a dot representing fractions <=5% The upper red bar (where present) indicates the fraction of residues that have poor fit to the EM map (all-atom inclusion < 40%). The numeric value is given above the bar.

| Mol | Chain | Length | Quality of chain |
|---|---|---|---|
| 1 | A1 | 525 | • 80% 19% |
| 1 | A2 | 525 | • 78% 21% |
| 1 | A3 | 525 | • 81% 18% • |
| 1 | B1 | 525 | • 81% 18% |

*Continued on next page...*

For Manuscript Review

*Continued from previous page...*

| Mol | Chain | Length | Quality of chain |
|---|---|---|---|
| 1 | B2 | 525 | 81% 19% |
| 1 | B3 | 525 | 80% 19% |
| 1 | X1 | 525 | 76% 23% |
| 1 | X2 | 525 | 80% 20% |
| 1 | X3 | 525 | 76% 23% |
| 1 | c1 | 525 | 74% 25% |
| 1 | c2 | 525 | 79% 21% |
| 1 | c3 | 525 | 78% 22% |
| 1 | d1 | 525 | 82% 18% |
| 1 | d2 | 525 | 76% 24% |
| 1 | d3 | 525 | 77% 22% |
| 1 | e1 | 525 | 78% 21% |
| 1 | e2 | 525 | 81% 19% |
| 1 | e3 | 525 | 80% 19% |
| 1 | f1 | 525 | 78% 22% |
| 1 | f2 | 525 | 80% 20% |
| 1 | f3 | 525 | 82% 18% |
| 2 | vC | 120 | 20% 78% |
| 3 | wD | 225 | 36% 6% 56% |
| 4 | 57 | 282 | 34% 9% 57% |
| 5 | TP | 608 | 11% 5% 85% |
| 5 | TQ | 608 | 11% 88% |
| 6 | ze | 169 | 31% 65% |
| 7 | he | 166 | 69% 11% 20% |
| 8 | gc | 329 | 22% 67% 16% 17% |

*Continued on next page...*

For Manuscript Review

*Continued from previous page...*

| Mol | Chain | Length | Quality of chain |
|---|---|---|---|
| 9 | uB | 267 | 5% 81% 19% |
| 10 | 76 | 210 | 7% 54% 12% 33% |

For Manuscript Review

### 2 Entry composition (i)

There are 10 unique types of molecules in this entry. The entry contains 96071 atoms, of which 0 are hydrogens and 0 are deuteriums.

In the tables below, the AltConf column contains the number of residues with at least one atom in alternate conformation and the Trace column contains the number of residues modelled with at most 2 atoms.

- Molecule 1 is a protein called Major capsid protein (ORF313).

| Mol | Chain | Residues | Atoms | | | | | AltConf | Trace |
|---|---|---|---|---|---|---|---|---|---|
| | | | Total | C | N | O | S | | |
| 1 | B2 | 524 | 4086 | 2568 | 702 | 795 | 21 | 0 | 0 |
| 1 | B1 | 524 | 4086 | 2568 | 702 | 795 | 21 | 0 | 0 |
| 1 | B3 | 524 | 4086 | 2568 | 702 | 795 | 21 | 0 | 0 |
| 1 | c2 | 524 | 4086 | 2568 | 702 | 795 | 21 | 0 | 0 |
| 1 | c1 | 520 | 4065 | 2555 | 698 | 791 | 21 | 0 | 0 |
| 1 | c3 | 524 | 4086 | 2568 | 702 | 795 | 21 | 0 | 0 |
| 1 | d2 | 524 | 4086 | 2568 | 702 | 795 | 21 | 0 | 0 |
| 1 | d1 | 524 | 4086 | 2568 | 702 | 795 | 21 | 0 | 0 |
| 1 | d3 | 524 | 4086 | 2568 | 702 | 795 | 21 | 0 | 0 |
| 1 | f2 | 524 | 4086 | 2568 | 702 | 795 | 21 | 0 | 0 |
| 1 | f1 | 524 | 4086 | 2568 | 702 | 795 | 21 | 0 | 0 |
| 1 | f3 | 524 | 4086 | 2568 | 702 | 795 | 21 | 0 | 0 |
| 1 | X2 | 524 | 4086 | 2568 | 702 | 795 | 21 | 0 | 0 |
| 1 | X1 | 524 | 4086 | 2568 | 702 | 795 | 21 | 0 | 0 |
| 1 | X3 | 524 | 4086 | 2568 | 702 | 795 | 21 | 0 | 0 |
| 1 | A2 | 524 | 4086 | 2568 | 702 | 795 | 21 | 0 | 0 |
| 1 | A1 | 524 | 4086 | 2568 | 702 | 795 | 21 | 0 | 0 |

*Continued on next page...*

For Manuscript Review

*Continued from previous page...*

| Mol | Chain | Residues | Atoms | | | | | AltConf | Trace |
|---|---|---|---|---|---|---|---|---|---|
| 1 | A3 | 524 | Total 4086 | C 2568 | N 702 | O 795 | S 21 | 0 | 0 |
| 1 | e2 | 524 | Total 4086 | C 2568 | N 702 | O 795 | S 21 | 0 | 0 |
| 1 | e1 | 524 | Total 4086 | C 2568 | N 702 | O 795 | S 21 | 0 | 0 |
| 1 | e3 | 524 | Total 4086 | C 2568 | N 702 | O 795 | S 21 | 0 | 0 |

- Molecule 2 is a protein called ORF289.

| Mol | Chain | Residues | Atoms | | | | AltConf | Trace |
|---|---|---|---|---|---|---|---|---|
| 2 | vC | 26 | Total 188 | C 113 | N 38 | O 37 | 0 | 0 |

- Molecule 3 is a protein called ORF535.

| Mol | Chain | Residues | Atoms | | | | | AltConf | Trace |
|---|---|---|---|---|---|---|---|---|---|
| 3 | wD | 99 | Total 763 | C 470 | N 139 | O 151 | S 3 | 0 | 0 |

- Molecule 4 is a protein called ORF357.

| Mol | Chain | Residues | Atoms | | | | | AltConf | Trace |
|---|---|---|---|---|---|---|---|---|---|
| 4 | 57 | 120 | Total 965 | C 596 | N 170 | O 190 | S 9 | 0 | 0 |

- Molecule 5 is a protein called Tape measure (ORF401).

| Mol | Chain | Residues | Atoms | | | | | AltConf | Trace |
|---|---|---|---|---|---|---|---|---|---|
| 5 | TP | 94 | Total 720 | C 435 | N 135 | O 145 | S 5 | 0 | 0 |
| 5 | TQ | 72 | Total 574 | C 355 | N 104 | O 113 | S 2 | 0 | 0 |

- Molecule 6 is a protein called Zipper protein (ORF214).

| Mol | Chain | Residues | Atoms | | | | AltConf | Trace |
|---|---|---|---|---|---|---|---|---|
| 6 | ze | 59 | Total 448 | C 270 | N 85 | O 93 | 0 | 0 |

- Molecule 7 is a protein called ORF404.

For Manuscript Review

| Mol | Chain | Residues | Atoms | | | | | AltConf | Trace |
|---|---|---|---|---|---|---|---|---|---|
| | | | Total | C | N | O | S | | |
| 7 | he | 132 | 1086 | 683 | 183 | 215 | 5 | 0 | 0 |

- Molecule 8 is a protein called ORF481.

| Mol | Chain | Residues | Atoms | | | | | AltConf | Trace |
|---|---|---|---|---|---|---|---|---|---|
| | | | Total | C | N | O | S | | |
| 8 | gc | 274 | 2267 | 1460 | 371 | 428 | 8 | 0 | 0 |

- Molecule 9 is a protein called ORF281.

| Mol | Chain | Residues | Atoms | | | | | AltConf | Trace |
|---|---|---|---|---|---|---|---|---|---|
| | | | Total | C | N | O | S | | |
| 9 | uB | 267 | 2166 | 1346 | 395 | 414 | 11 | 0 | 0 |

- Molecule 10 is a protein called ORF76.

| Mol | Chain | Residues | Atoms | | | | | AltConf | Trace |
|---|---|---|---|---|---|---|---|---|---|
| | | | Total | C | N | O | S | | |
| 10 | 76 | 140 | 1109 | 675 | 208 | 220 | 6 | 0 | 0 |

For Manuscript Review

### 3 Residue-property plots

These plots are drawn for all protein, RNA, DNA and oligosaccharide chains in the entry. The first graphic for a chain summarises the proportions of the various outlier classes displayed in the second graphic. The second graphic shows the sequence view annotated by issues in geometry and atom inclusion in map density. Residues are color-coded according to the number of geometric quality criteria for which they contain at least one outlier: green = 0, yellow = 1, orange = 2 and red = 3 or more. A red diamond above a residue indicates a poor fit to the EM map for this residue (all-atom inclusion < 40%). Stretches of 2 or more consecutive residues without any outlier are shown as a green connector. Residues present in the sample, but not in the model, are shown in grey.

• Molecule 1: Major capsid protein (ORF313)

Chain B2:

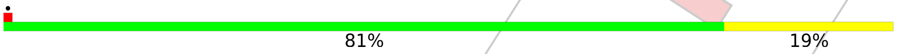


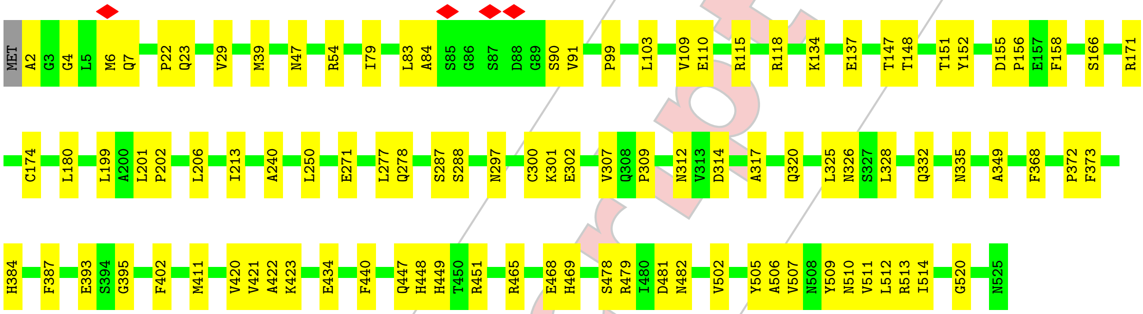


• Molecule 1: Major capsid protein (ORF313)

Chain B1:

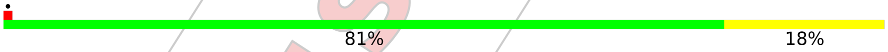


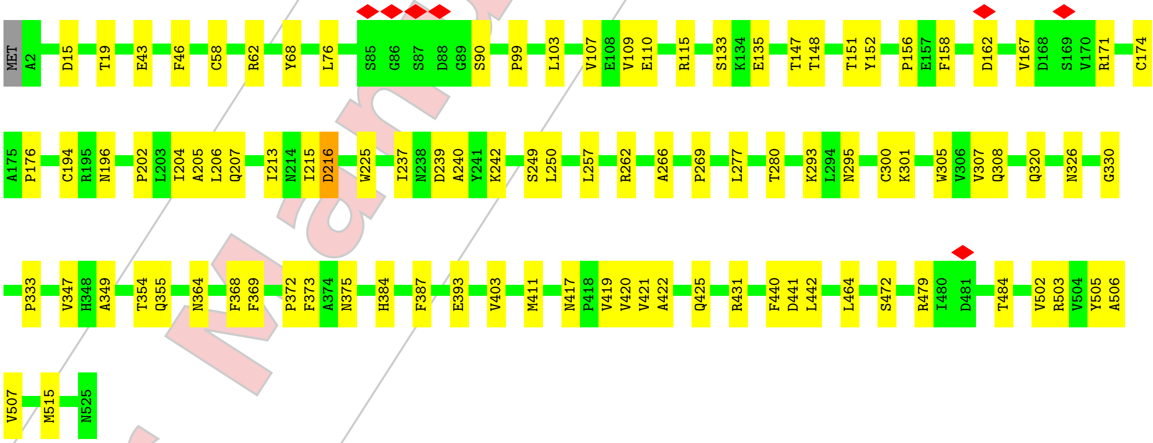


• Molecule 1: Major capsid protein (ORF313)

Chain B3:

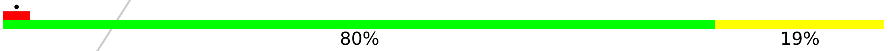


For Manuscript Review

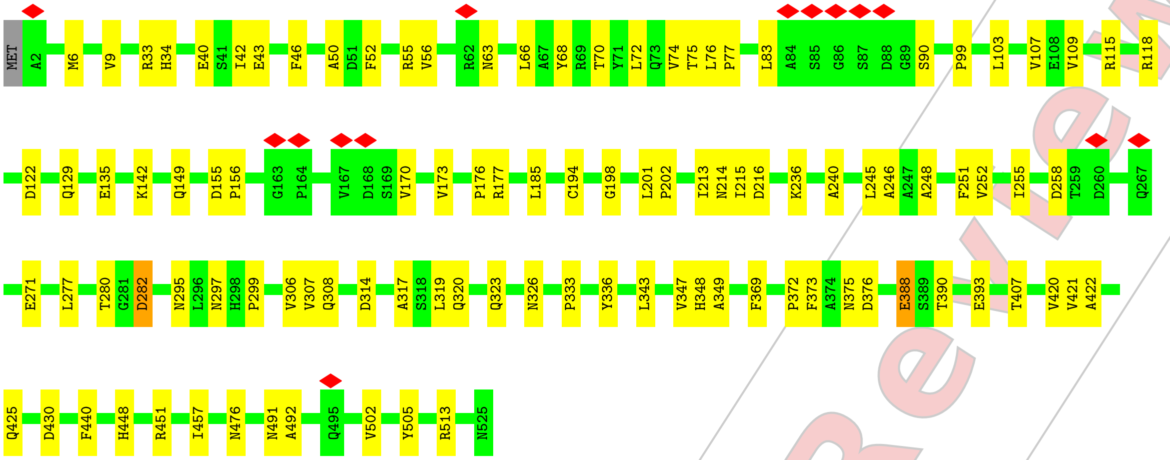


- Molecule 1: Major capsid protein (ORF313)

Chain c2:

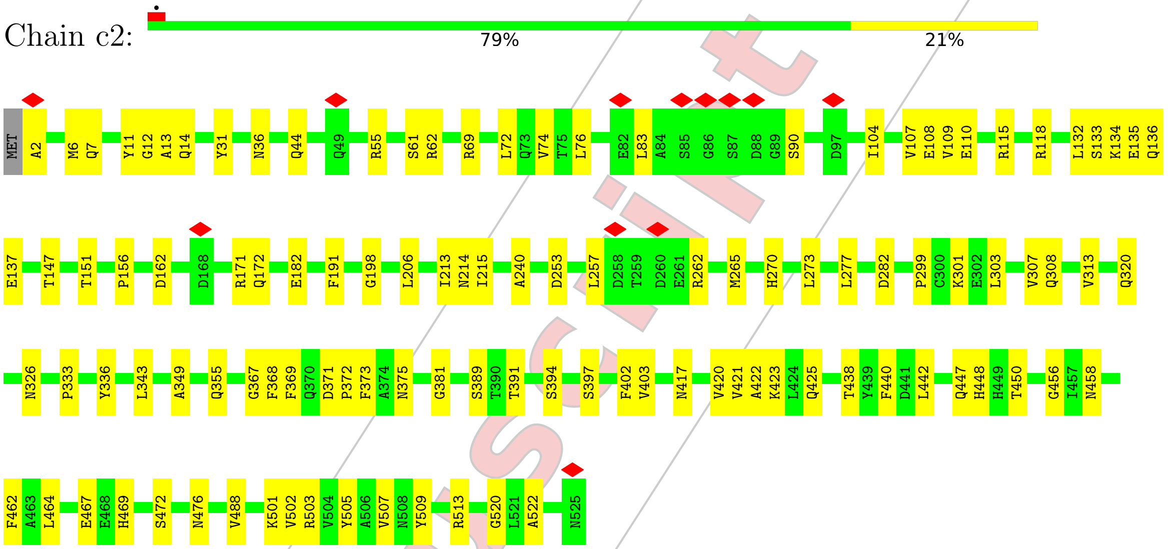


- Molecule 1: Major capsid protein (ORF313)

Chain c1:

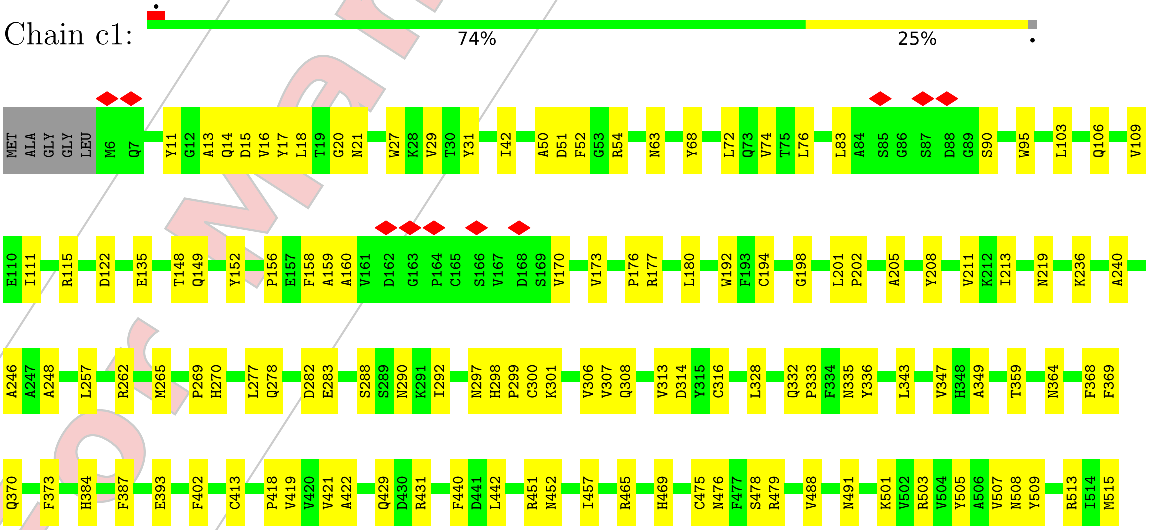


For Manuscript Review

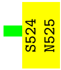


• Molecule 1: Major capsid protein (ORF313)

Chain c3:

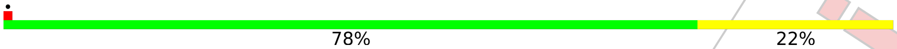


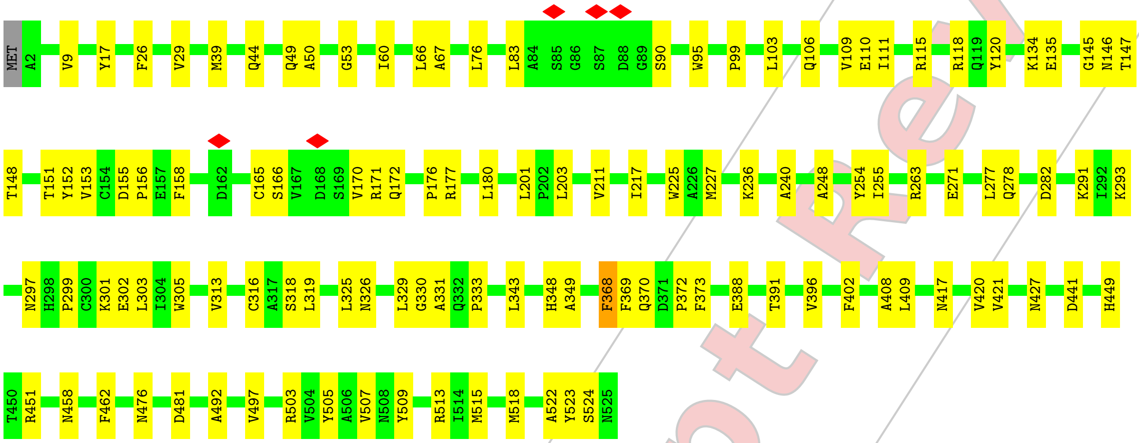


• Molecule 1: Major capsid protein (ORF313)

Chain d2:

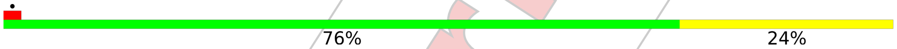


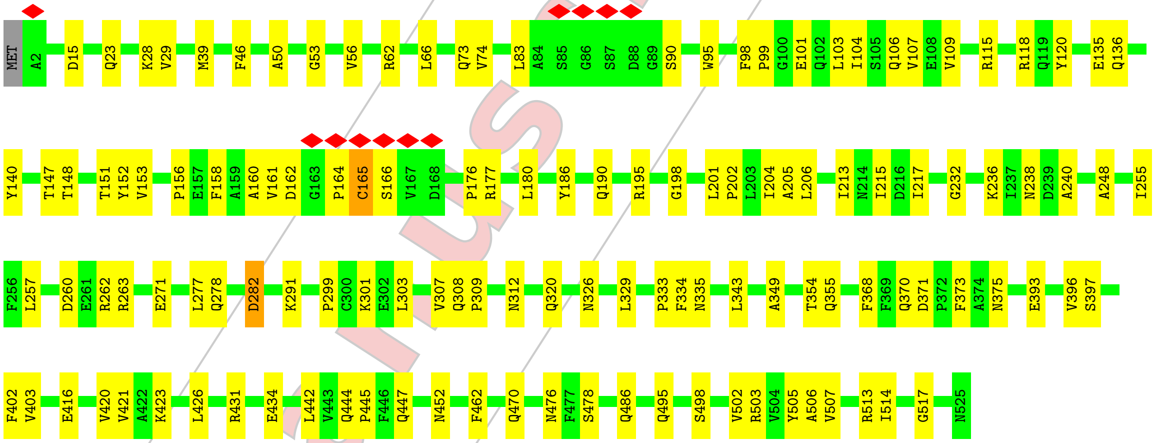


• Molecule 1: Major capsid protein (ORF313)

Chain d1:

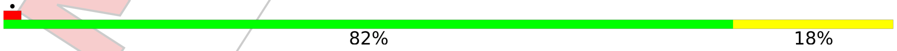


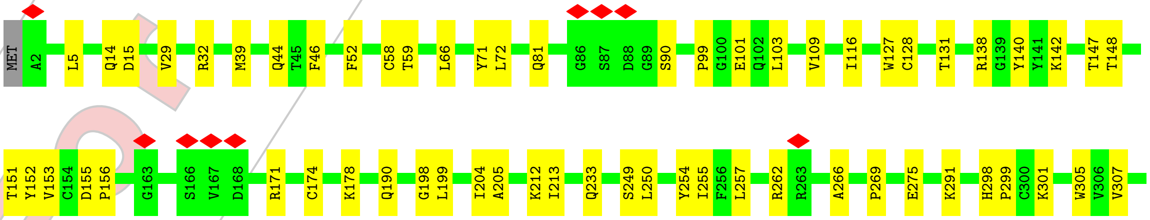

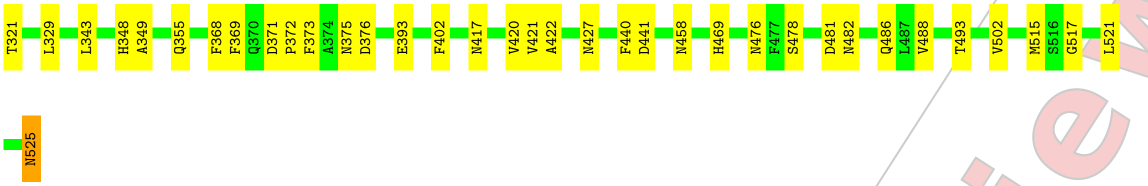


• Molecule 1: Major capsid protein (ORF313)

Chain d3:

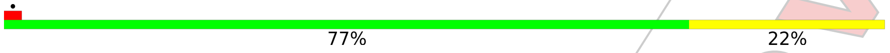


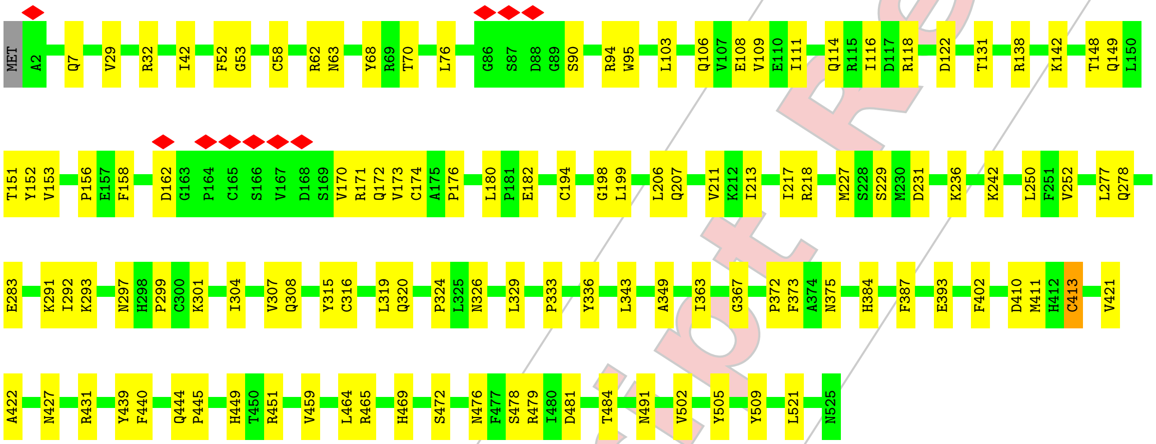


• Molecule 1: Major capsid protein (ORF313)

Chain f2:

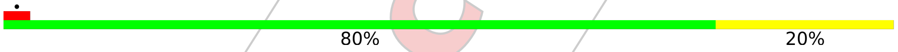


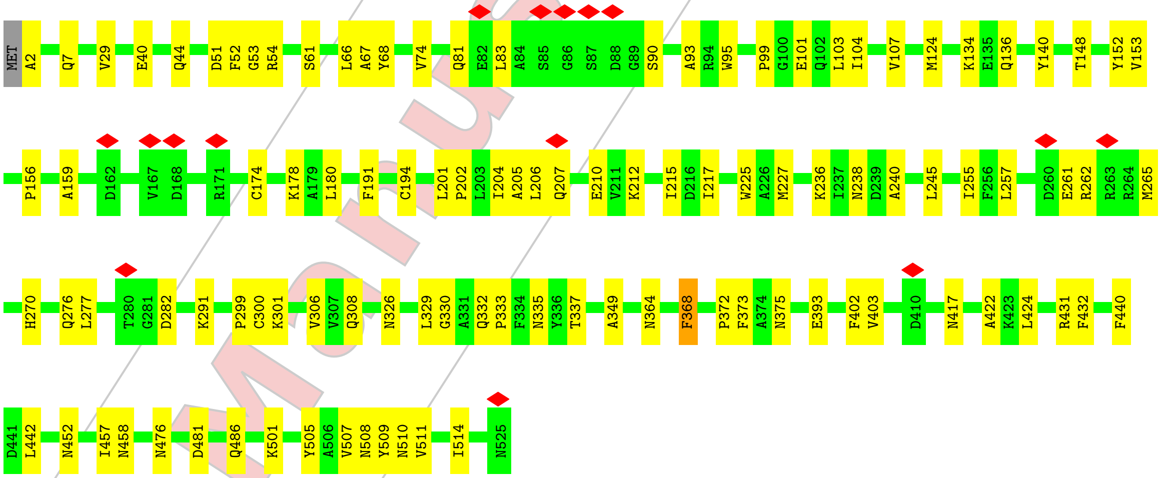


• Molecule 1: Major capsid protein (ORF313)

Chain f1:

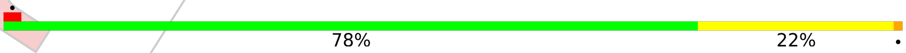


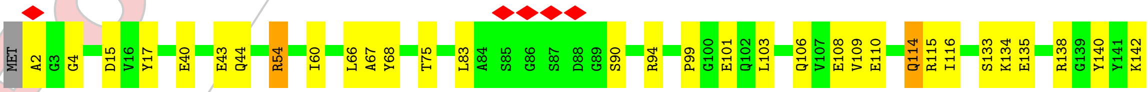


For Manuscript Review

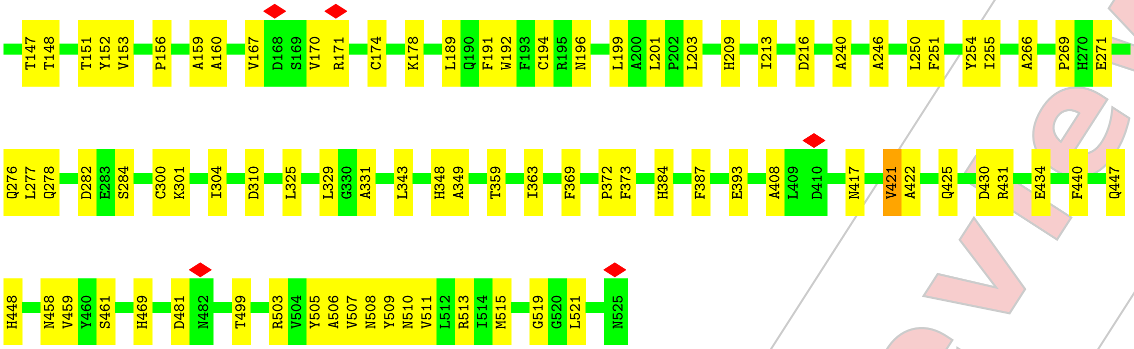


• Molecule 1: Major capsid protein (ORF313)

Chain f3:

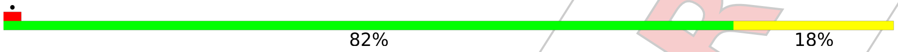


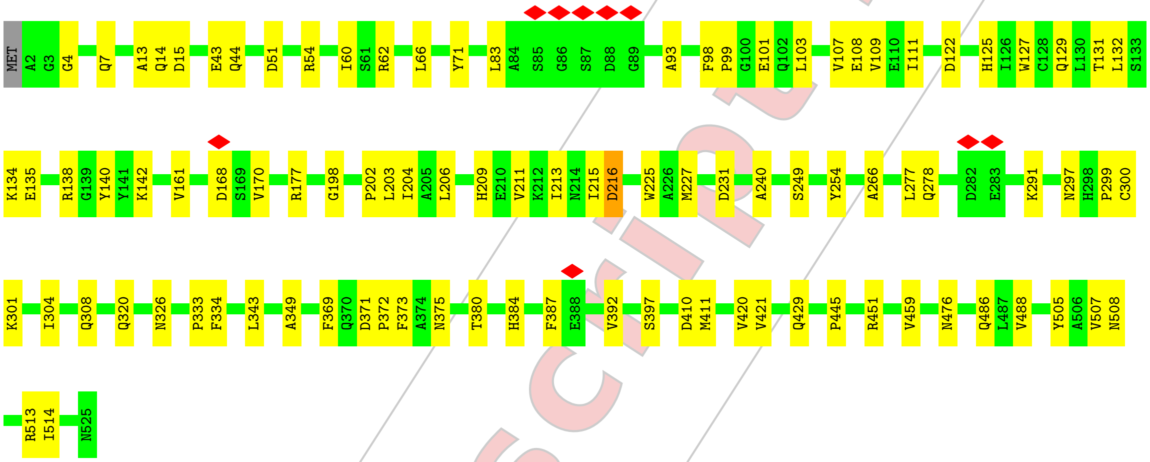


• Molecule 1: Major capsid protein (ORF313)

Chain X2:

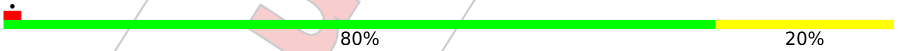


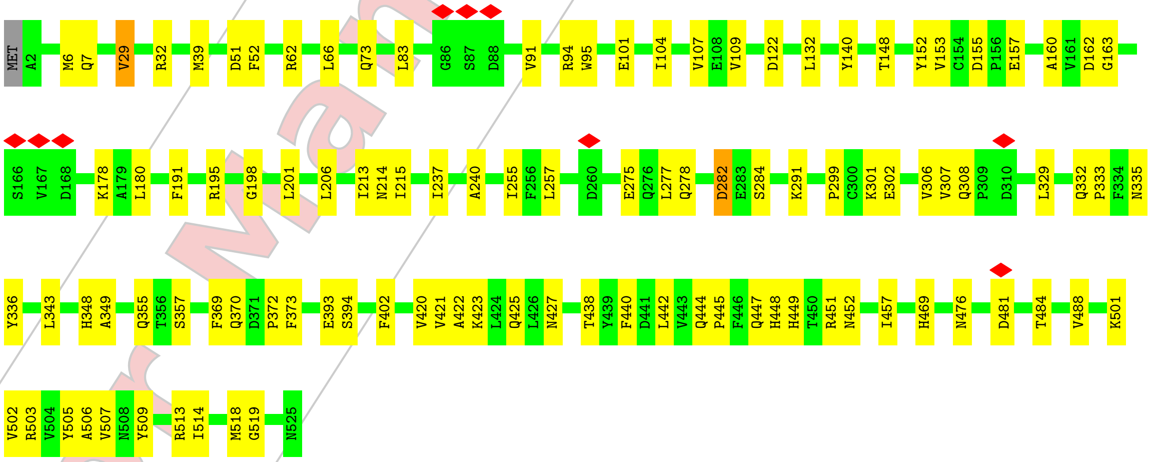


• Molecule 1: Major capsid protein (ORF313)

Chain X1:

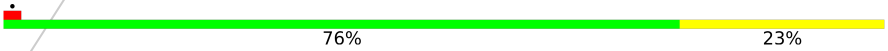


For Manuscript Review

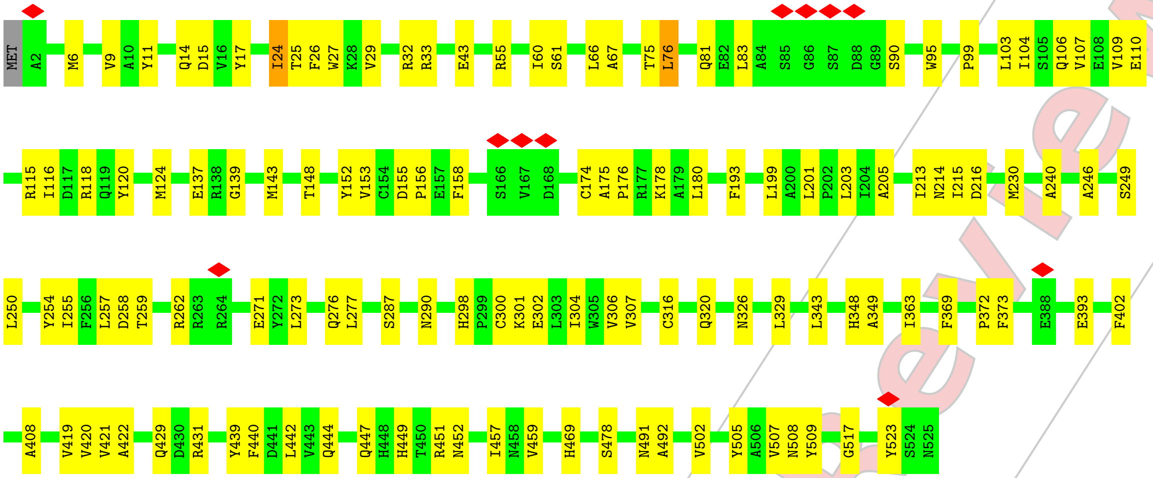


• Molecule 1: Major capsid protein (ORF313)

Chain X3:

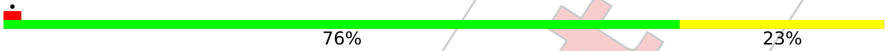


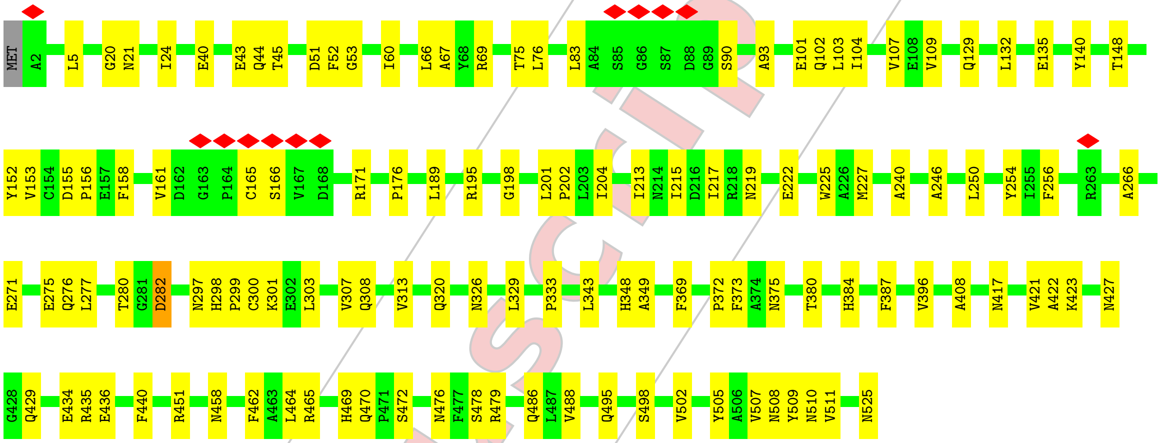


• Molecule 1: Major capsid protein (ORF313)

Chain A2:

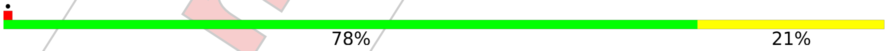


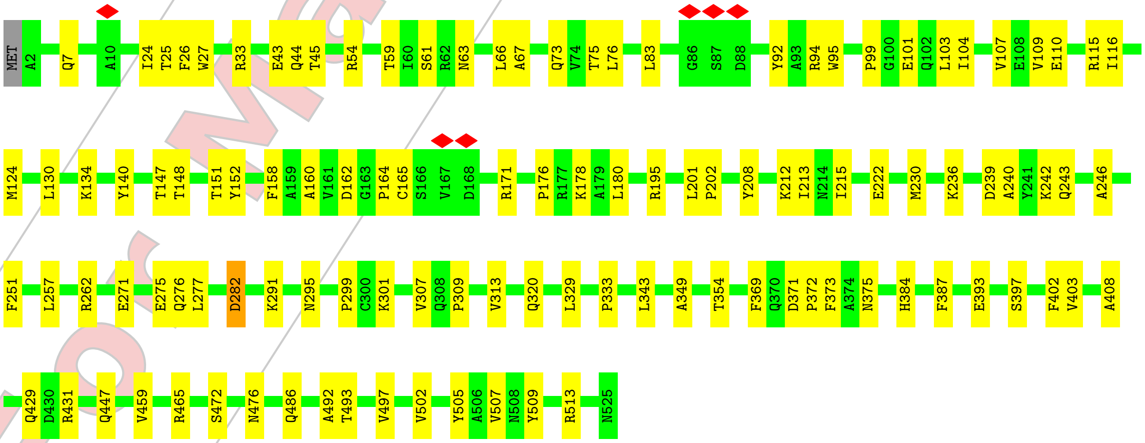


• Molecule 1: Major capsid protein (ORF313)

For Manuscript Review

Chain A1:

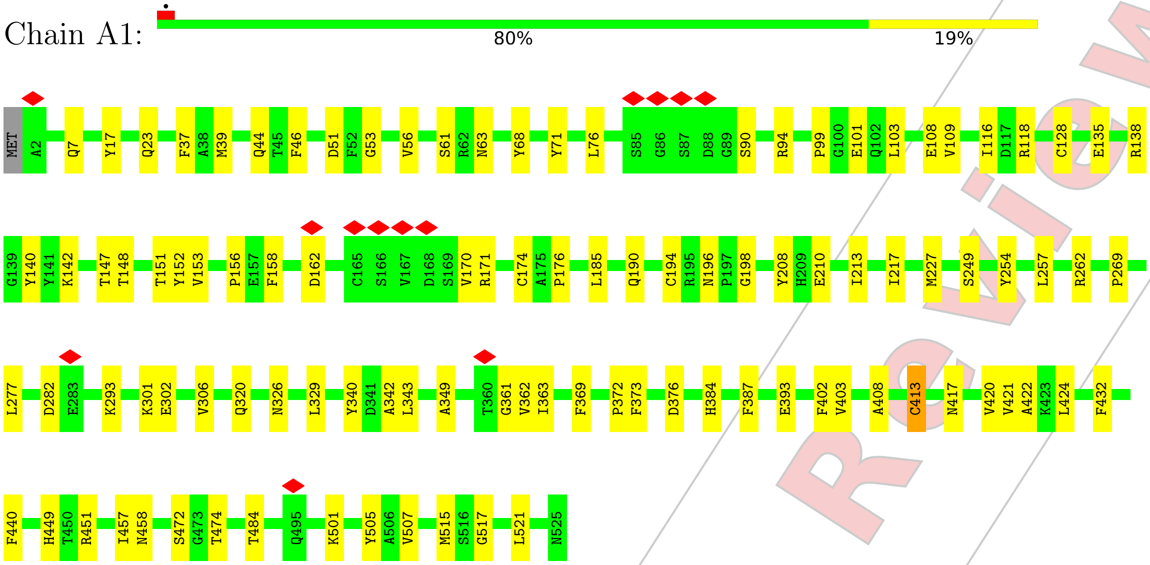


• Molecule 1: Major capsid protein (ORF313)

Chain A3:

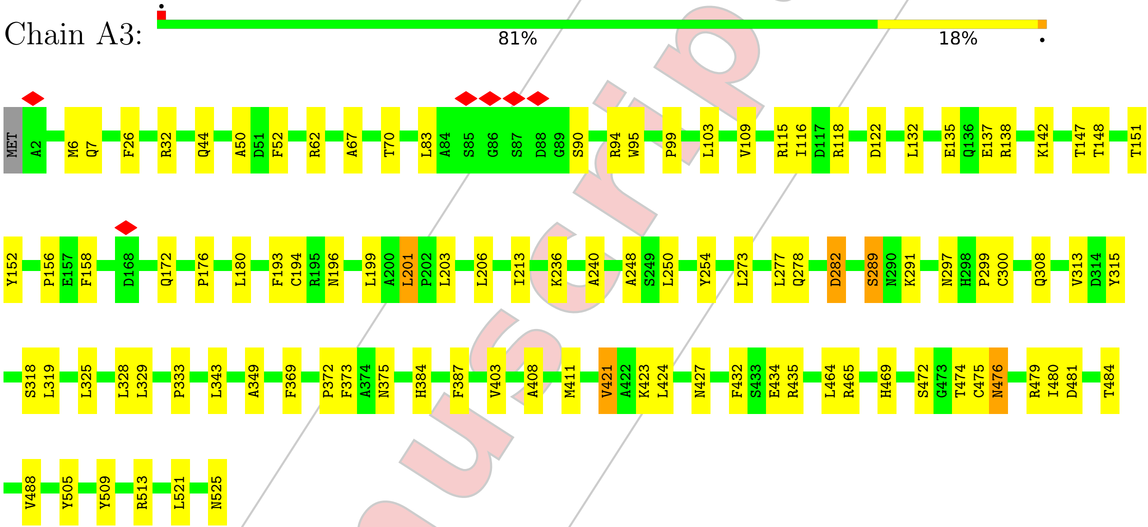


• Molecule 1: Major capsid protein (ORF313)

Chain e2:

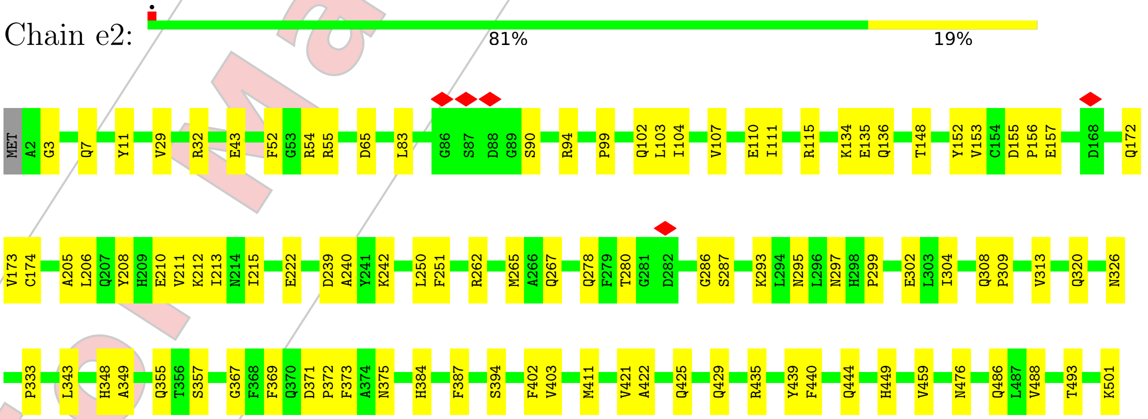


For Manuscript Review

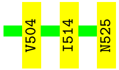


• Molecule 1: Major capsid protein (ORF313)

Chain e1:

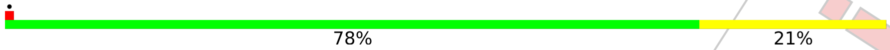


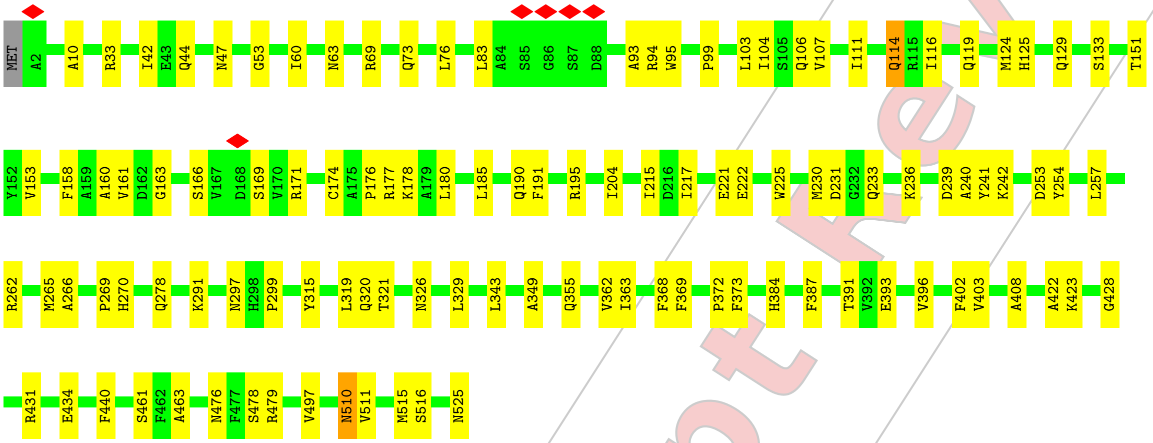


• Molecule 1: Major capsid protein (ORF313)

Chain e3:

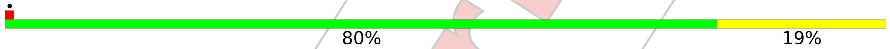


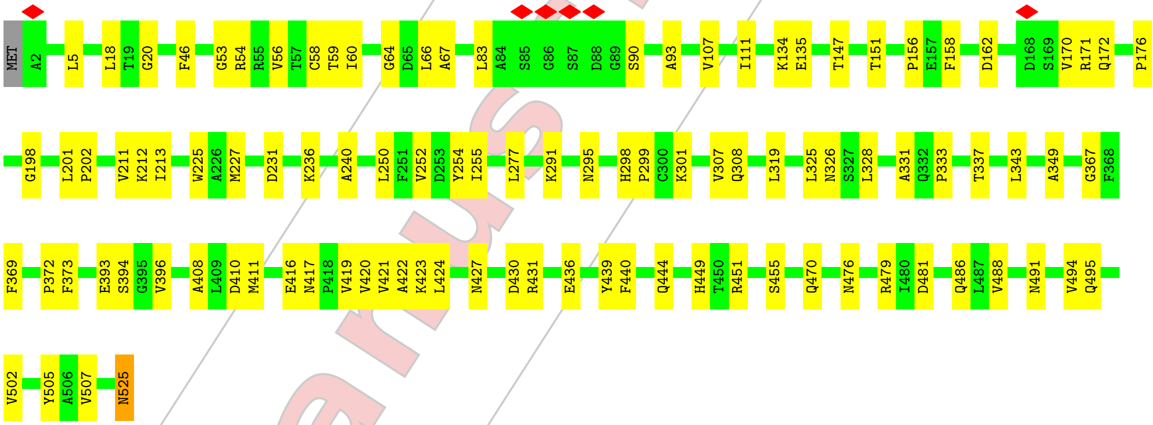


• Molecule 2: ORF289

Chain vC:

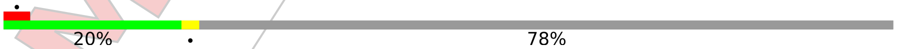


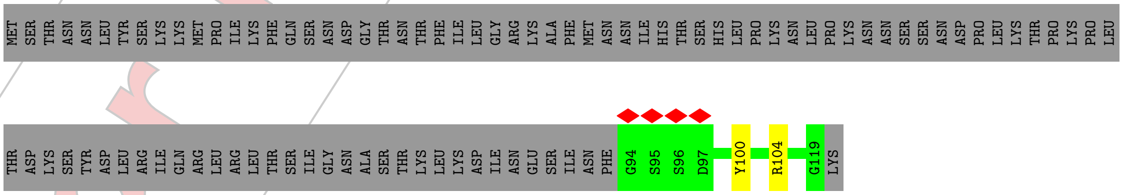


• Molecule 3: ORF535

Chain wD:

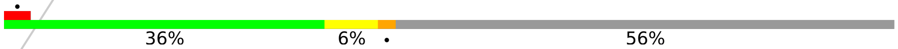


For Manuscript Review

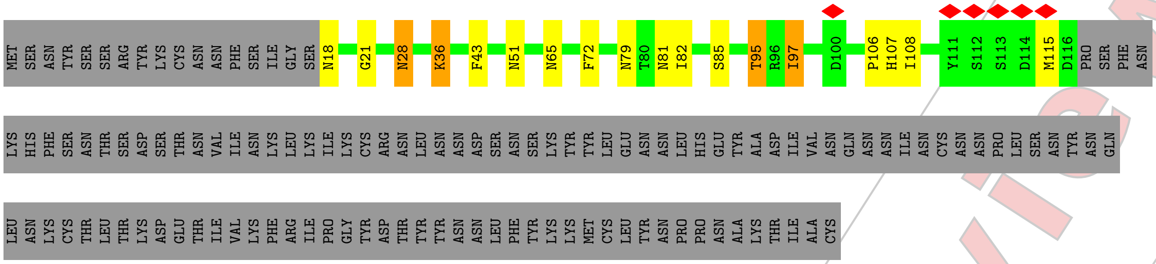


• Molecule 4: ORF357

Chain 57:

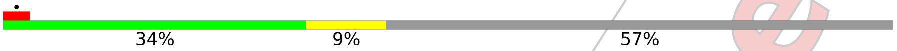


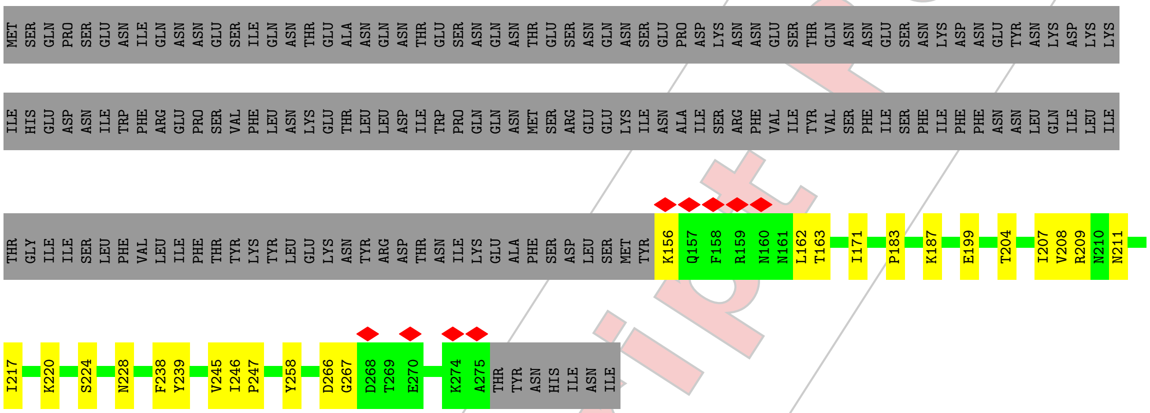


• Molecule 5: Tape measure (ORF401)

Chain TP:

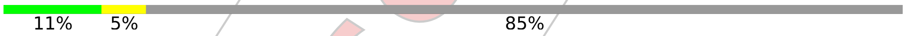


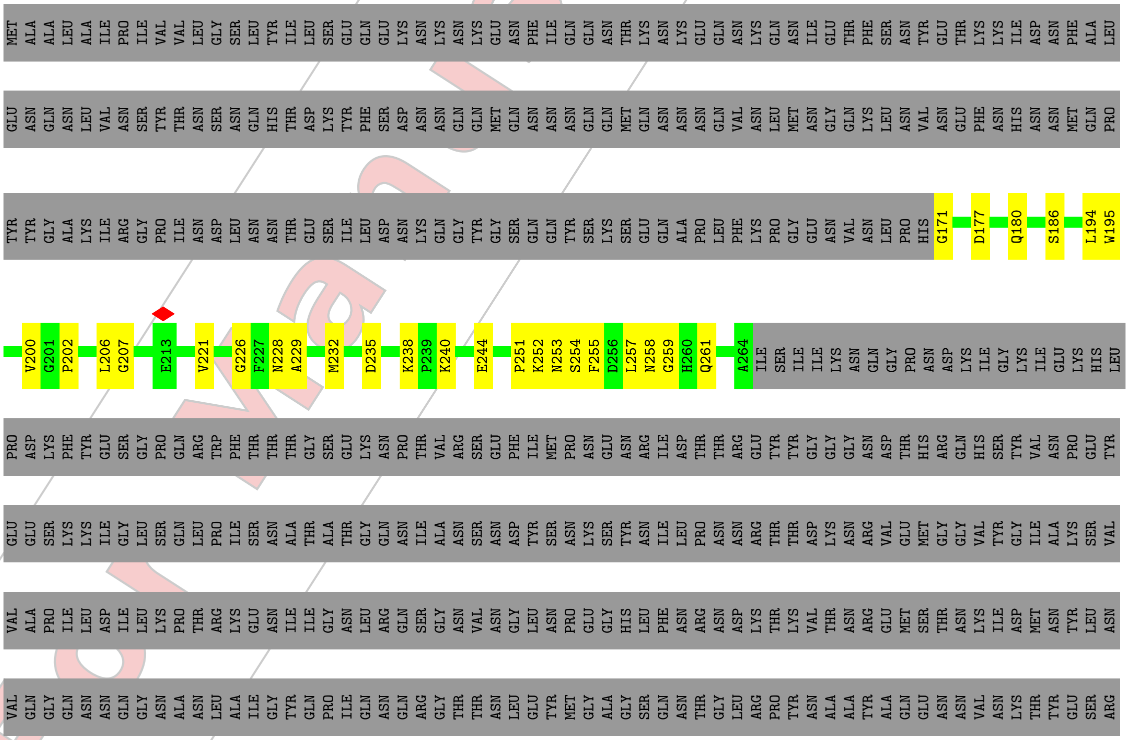


For Manuscript Review

PRO ASN GLN GLY MET ASN LEU PHE ASN ASN HIS ASN ASN VAL GLU ILE TYR LYS ASN GLU ASN ILE LEU LYS ASN ASN ARG GLY ASN ILE SER ASN GLY GLY PRO ASN ILE ILE PRO SER SER GLU PHE MET GLY GLU ILE ASN SER MET GLN SER TYR ASP ASN ASN PHE ASN ALA

SER ARG MET ASP GLU SER LEU LEU ASN ALA PHE LYS ASN ASN PRO TYR THR LYS SER LEU SER SER VAL ALA

• Molecule 5: Tape measure (ORF401)

Chain TQ: 11% 88%

MET ALA ALA LEU ALA ILE PRO ILE VAL VAL LEU GLY SER LEU TYR ILE LEU SER GLU GLN GLU LYS ASN LYS ASN LYS GLU ASN PHE ILE GLN GLN ASN THR LYS ASN LYS GLU GLN ASN LYS GLN ASN ILE GLU THR PHE SER ASN TYR GLU THR LYS LYS ILE ASP ASN PHE ALA LEU

GLU ASN GLN ASN LEU VAL ASN SER TYR THR ASN SER ASN GLN HIS THR ASP LYS TYR PHE SER ASP ASN ASN GLN GLN MET GLN ASN ASN ASN GLN GLN MET GLN ASN ASN ASN Q99 V100 M103 Q106 V110 N111 E112 H115 R127 Q144 E155 P158 L159

H170 GLY THR ALA ASN ASN SER ASP PHE PHE GLN SER ARG MET ASN GLU SER MET LYS MET SER ASN VAL THR LEU TRP GLU PRO GLN ARG VAL GLY PRO GLY LEU ASN LEU GLY TYR GLY SER GLN ASN GLU ASN GLY PHE ASN ARG GLY GLY VAL ASN GLY SER ASP GLY PHE ASN ALA

GLY MET MET SER ARG ASP SER TRP LYS PRO LYS THR VAL ASP GLU LEU ARG VAL ASP ASN ASN PRO LYS ASN SER PHE ASP LEU ASN GLY HIS GLN GLY PRO ALA ILE SER ILE ILE LYS ASN GLN GLY PRO ASN ASP LYS ILE GLY LYS ILE GLU GLN LYS HIS LEU PRO ASP LYS PHE TYR

GLU SER GLY PRO GLN ARG TRP PHE THR THR THR GLY SER GLU LYS ASN PRO THR VAL ARG SER GLU PHE ILE MET PRO ASN GLU ASN ARG ILE ASP THR THR ARG GLU TYR TYR GLY GLY GLY ASN ASP THR HIS ARG GLN HIS SER TYR VAL ASN PRO GLU TYR GLU GLU SER LYS LYS

ILE GLY LEU SER GLN LEU PRO ILE SER ASN ALA THR ALA THR GLY GLN ASN ILE ALA ASN SER ASN ASP TYR SER ASN LYS SER TYR ASN ILE LEU PRO ASN ASN ARG THR THR ASP LYS ASN ARG VAL GLU MET GLY GLY VAL TYR GLY ILE ALA LYS SER VAL VAL ALA PRO ILE LEU

ASP ILE LEU LYS PRO THR ARG LYS GLU ASN ILE ILE GLY ASN LEU ARG GLN SER GLY ASN VAL ASN GLY LEU ASN PRO GLU GLY HIS LEU PHE ASN ARG ASN ASP LYS THR LYS VAL THR ASN ARG GLU MET SER THR ASN LYS ILE ASP MET ASN TYR LEU ASN VAL GLN GLY GLN ASN

ASN GLN GLY ASN ALA ASN LEU ALA ILE GLY TYR GLN PRO ILE GLN ASN GLN ARG GLY THR THR ASN LEU GLU TYR MET GLY ALA GLY SER GLN ASN THR GLY LEU ARG PRO TYR ASN ALA ALA TYR ALA GLN GLU ASN ASN VAL ASN LYS THR TYR GLU SER ARG PRO ASN GLN GLY GLY

MET ASN LEU PHE ASN ASN HIS ASN ASN VAL GLU ILE TYR LYS ASN GLU ASN ILE LEU LYS ASN ASN ARG GLY ASN ILE SER ASN GLY PRO ASN ILE ILE PRO SER SER GLU PHE MET GLY GLU ILE ASN SER MET GLN SER TYR ASP ASN ASN PHE ASN ALA SER ARG MET ASP GLU

SER LEU LEU ASN ALA PHE LYS ASN ASN PRO TYR THR LYS SER LEU SER SER VAL ALA

• Molecule 6: Zipper protein (ORF214)

Chain ze: 31% 65%

MET THR SER PHE LYS LYS MET PHE LYS GLY PHE LYS ASN GLU HIS ILE LEU GLY ILE LEU GLY LEU LEU ILE LEU GLY PHE ALA LEU TYR LYS TYR SER ASP ASN LYS ASN VAL HIS THR LEU GLY MET THR PRO GLY PRO VAL THR LEU LYS SER SER GLU VAL ASN ASN GLN PRO HIS

PRO MET ILE ASN ASN MET MET THR SER MET VAL GLN ASP VAL VAL ASN LYS PRO ALA ALA THR ASN PRO SER ASP L86 E94 Q97 P100 L106 L111 L112 N113 N121 S125 L134 N144 THR ASN CYS PRO TRP ASN ASN ILE SER THR ILE ASP THR ASP LYS

PHE ARG LYS ALA LEU GLU ILE GLY THR GLY ALA

• Molecule 7: ORF404

Chain he: 69% 11% 20%

For Manuscript Review

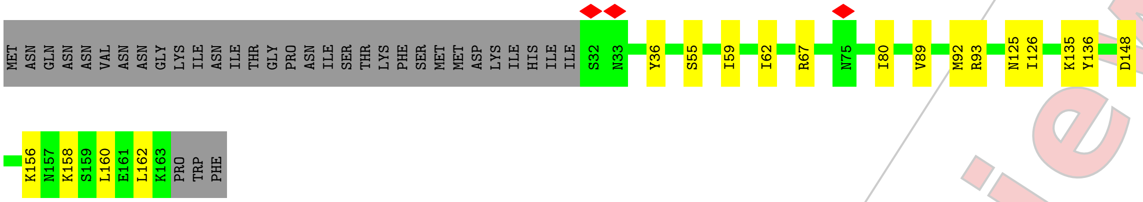


- Molecule 8: ORF481

Chain gc:

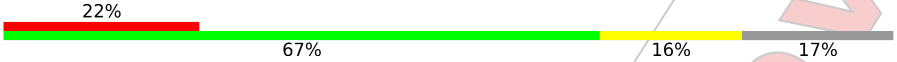


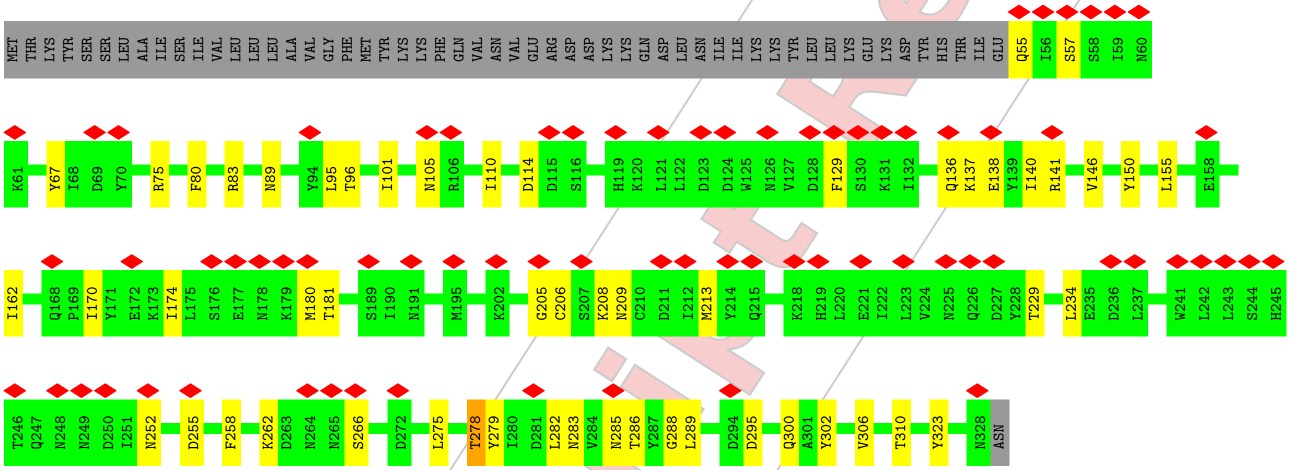


- Molecule 9: ORF281

Chain uB:

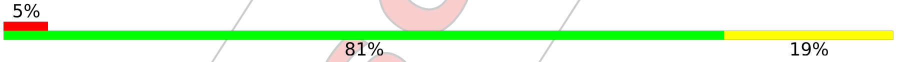


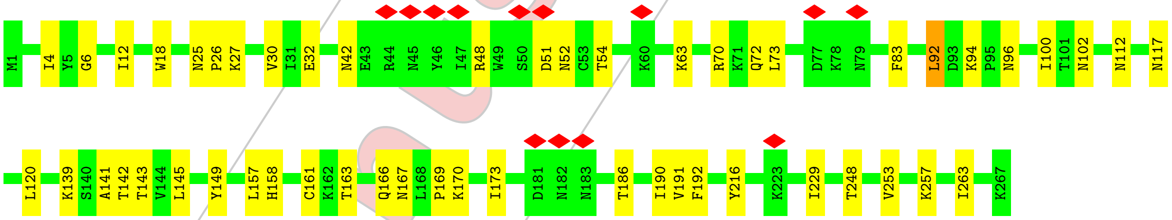


- Molecule 10: ORF76

Chain 76:

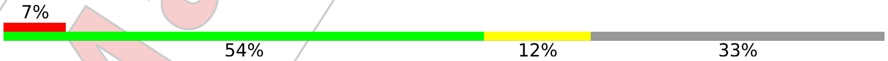


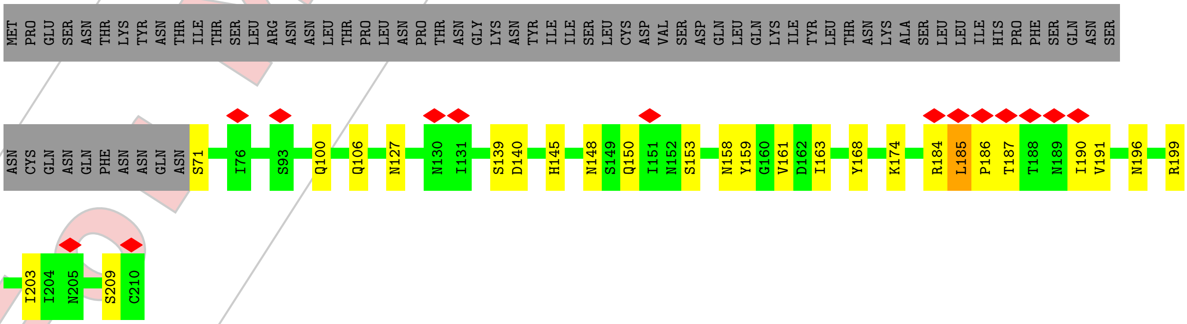


For Manuscript Review

### 4 Experimental information (i)

| Property | Value | Source |
|---|---|---|
| EM reconstruction method | SINGLE PARTICLE | Depositor |
| Imposed symmetry | POINT, Not provided | |
| Number of particles used | 2934220 | Depositor |
| Resolution determination method | FSC 0.143 CUT-OFF | Depositor |
| CTF correction method | PHASE FLIPPING AND AMPLITUDE CORRECTION | Depositor |
| Microscope | TFS KRIOS | Depositor |
| Voltage (kV) | 300 | Depositor |
| Electron dose $(e^-/\text{Å}^2)$ | 22.9 | Depositor |
| Minimum defocus (nm) | 800 | Depositor |
| Maximum defocus (nm) | 2500 | Depositor |
| Magnification | Not provided | |
| Image detector | GATAN K3 (6k x 4k) | Depositor |
| Maximum map value | 0.235 | Depositor |
| Minimum map value | -0.171 | Depositor |
| Average map value | 0.000 | Depositor |
| Map value standard deviation | 0.017 | Depositor |
| Recommended contour level | 0.03 | Depositor |
| Map size (Å) | 345.6, 345.6, 345.6 | wwPDB |
| Map dimensions | 256, 256, 256 | wwPDB |
| Map angles (°) | 90.0, 90.0, 90.0 | wwPDB |
| Pixel spacing (Å) | 1.35, 1.35, 1.35 | Depositor |

For Manuscript Review

### 5 Model quality (i)

#### 5.1 Standard geometry (i)

The Z score for a bond length (or angle) is the number of standard deviations the observed value is removed from the expected value. A bond length (or angle) with $|Z| > 5$ is considered an outlier worth inspection. RMSZ is the root-mean-square of all Z scores of the bond lengths (or angles).

| Mol | Chain | Bond lengths | | Bond angles | |
|---|---|---|---|---|---|
| | | RMSZ | #$\|Z\|$ >5 | RMSZ | #$\|Z\|$ >5 |
| 1 | A1 | 0.13 | 0/4181 | 0.26 | 0/5690 |
| 1 | A2 | 0.14 | 0/4181 | 0.26 | 0/5690 |
| 1 | A3 | 0.14 | 0/4181 | 0.27 | 0/5690 |
| 1 | B1 | 0.13 | 0/4181 | 0.26 | 0/5690 |
| 1 | B2 | 0.15 | 0/4181 | 0.28 | 0/5690 |
| 1 | B3 | 0.13 | 0/4181 | 0.27 | 0/5690 |
| 1 | X1 | 0.13 | 0/4181 | 0.27 | 0/5690 |
| 1 | X2 | 0.13 | 0/4181 | 0.27 | 0/5690 |
| 1 | X3 | 0.14 | 0/4181 | 0.27 | 0/5690 |
| 1 | c1 | 0.14 | 0/4160 | 0.28 | 0/5662 |
| 1 | c2 | 0.12 | 0/4181 | 0.25 | 0/5690 |
| 1 | c3 | 0.14 | 0/4181 | 0.27 | 0/5690 |
| 1 | d1 | 0.13 | 0/4181 | 0.26 | 0/5690 |
| 1 | d2 | 0.14 | 0/4181 | 0.27 | 0/5690 |
| 1 | d3 | 0.14 | 0/4181 | 0.27 | 0/5690 |
| 1 | e1 | 0.15 | 0/4181 | 0.28 | 0/5690 |
| 1 | e2 | 0.15 | 0/4181 | 0.28 | 0/5690 |
| 1 | e3 | 0.15 | 0/4181 | 0.28 | 0/5690 |
| 1 | f1 | 0.14 | 0/4181 | 0.27 | 0/5690 |
| 1 | f2 | 0.12 | 0/4181 | 0.26 | 0/5690 |
| 1 | f3 | 0.13 | 0/4181 | 0.26 | 0/5690 |
| 2 | vC | 0.09 | 0/189 | 0.19 | 0/250 |
| 3 | wD | 0.14 | 0/776 | 0.31 | 0/1047 |
| 4 | 57 | 0.12 | 0/981 | 0.26 | 0/1324 |
| 5 | TP | 0.13 | 0/735 | 0.27 | 0/989 |
| 5 | TQ | 0.12 | 0/586 | 0.34 | 0/792 |
| 6 | ze | 0.11 | 0/455 | 0.22 | 0/624 |
| 7 | he | 0.14 | 0/1103 | 0.26 | 0/1484 |
| 8 | gc | 0.09 | 0/2318 | 0.24 | 0/3138 |
| 9 | uB | 0.13 | 0/2205 | 0.29 | 0/2985 |
| 10 | 76 | 0.11 | 0/1126 | 0.27 | 0/1519 |
| All | All | 0.13 | 0/98254 | 0.27 | 0/133614 |

There are no bond length outliers.

For Manuscript Review

There are no bond angle outliers.

There are no chirality outliers.

There are no planarity outliers.

#### 5.2 Too-close contacts (i)

In the following table, the Non-H and H(model) columns list the number of non-hydrogen atoms and hydrogen atoms in the chain respectively. The H(added) column lists the number of hydrogen atoms added and optimized by MolProbity. The Clashes column lists the number of clashes within the asymmetric unit, whereas Symm-Clashes lists symmetry-related clashes.

| Mol | Chain | Non-H | H(model) | H(added) | Clashes | Symm-Clashes |
|---|---|---|---|---|---|---|
| 1 | A1 | 4086 | 0 | 3903 | 72 | 0 |
| 1 | A2 | 4086 | 0 | 3902 | 81 | 0 |
| 1 | A3 | 4086 | 0 | 3903 | 71 | 0 |
| 1 | B1 | 4086 | 0 | 3903 | 58 | 0 |
| 1 | B2 | 4086 | 0 | 3901 | 65 | 0 |
| 1 | B3 | 4086 | 0 | 3903 | 68 | 0 |
| 1 | X1 | 4086 | 0 | 3903 | 80 | 0 |
| 1 | X2 | 4086 | 0 | 3903 | 71 | 0 |
| 1 | X3 | 4086 | 0 | 3901 | 94 | 0 |
| 1 | c1 | 4065 | 0 | 3880 | 97 | 0 |
| 1 | c2 | 4086 | 0 | 3903 | 83 | 0 |
| 1 | c3 | 4086 | 0 | 3901 | 87 | 0 |
| 1 | d1 | 4086 | 0 | 3903 | 61 | 0 |
| 1 | d2 | 4086 | 0 | 3902 | 89 | 0 |
| 1 | d3 | 4086 | 0 | 3902 | 83 | 0 |
| 1 | e1 | 4086 | 0 | 3902 | 79 | 0 |
| 1 | e2 | 4086 | 0 | 3900 | 71 | 0 |
| 1 | e3 | 4086 | 0 | 3901 | 70 | 0 |
| 1 | f1 | 4086 | 0 | 3902 | 75 | 0 |
| 1 | f2 | 4086 | 0 | 3903 | 69 | 0 |
| 1 | f3 | 4086 | 0 | 3903 | 66 | 0 |
| 2 | vC | 188 | 0 | 193 | 2 | 0 |
| 3 | wD | 763 | 0 | 745 | 19 | 0 |
| 4 | 57 | 965 | 0 | 930 | 14 | 0 |
| 5 | TP | 720 | 0 | 661 | 25 | 0 |
| 5 | TQ | 574 | 0 | 545 | 7 | 0 |
| 6 | ze | 448 | 0 | 435 | 6 | 0 |
| 7 | he | 1086 | 0 | 1082 | 16 | 0 |
| 8 | gc | 2267 | 0 | 2236 | 34 | 0 |
| 9 | uB | 2166 | 0 | 2151 | 44 | 0 |
| 10 | 76 | 1109 | 0 | 1087 | 16 | 0 |

*Continued on next page...*

For Manuscript Review

*Continued from previous page...*

| Mol | Chain | Non-H | H(model) | H(added) | Clashes | Symm-Clashes |
|---|---|---|---|---|---|---|
| All | All | 96071 | 0 | 91989 | 1474 | 0 |

The all-atom clashscore is defined as the number of clashes found per 1000 atoms (including hydrogen atoms). The all-atom clashscore for this structure is 8.

All (1474) close contacts within the same asymmetric unit are listed below, sorted by their clash magnitude.

| Atom-1 | Atom-2 | Interatomic distance (Å) | Clash overlap (Å) |
|---|---|---|---|
| 1:e3:227:MET:HE2 | 1:e3:326:ASN:HD21 | 1.47 | 0.80 |
| 1:B2:47:ASN:O | 1:e3:295:ASN:ND2 | 2.18 | 0.77 |
| 8:gc:181:THR:HG22 | 8:gc:252:ASN:HB2 | 1.67 | 0.75 |
| 10:76:139:SER:HB2 | 10:76:163:ILE:HG22 | 1.69 | 0.75 |
| 1:c3:49:GLN:NE2 | 9:uB:143:THR:O | 2.21 | 0.74 |
| 1:c2:282:ASP:HB2 | 1:c2:501:LYS:HG2 | 1.70 | 0.73 |
| 1:d1:427:ASN:ND2 | 1:d1:481:ASP:OD1 | 2.21 | 0.72 |
| 1:A1:269:PRO:HB3 | 1:A1:515:MET:HG2 | 1.71 | 0.72 |
| 1:X1:298:HIS:H | 1:X1:478:SER:HB2 | 1.55 | 0.71 |
| 1:c2:282:ASP:HB3 | 1:c2:503:ARG:HG2 | 1.72 | 0.71 |
| 1:e1:111:ILE:HG12 | 1:e1:116:ILE:HG13 | 1.73 | 0.71 |
| 1:d2:282:ASP:HB3 | 1:d2:503:ARG:HG2 | 1.72 | 0.71 |
| 3:wD:106:PRO:O | 3:wD:107:HIS:ND1 | 2.23 | 0.71 |
| 1:f1:167:VAL:HG22 | 10:76:150:GLN:HB3 | 1.72 | 0.71 |
| 1:X2:29:VAL:HG23 | 1:X3:198:GLY:HA3 | 1.73 | 0.70 |
| 1:A3:109:VAL:HG22 | 1:A3:213:ILE:HG12 | 1.72 | 0.70 |
| 1:d1:269:PRO:HB3 | 1:d1:515:MET:HG2 | 1.74 | 0.70 |
| 1:B1:62:ARG:NH1 | 1:B1:206:LEU:O | 2.22 | 0.70 |
| 1:c2:13:ALA:H | 7:he:158:LYS:HE3 | 1.57 | 0.70 |
| 1:A2:25:THR:HG22 | 1:A2:27:TRP:H | 1.56 | 0.70 |
| 1:B2:277:LEU:HD11 | 1:B2:505:TYR:HB3 | 1.73 | 0.69 |
| 1:e2:421:VAL:HG12 | 1:e2:488:VAL:HB | 1.74 | 0.69 |
| 1:d2:277:LEU:HD11 | 1:d2:505:TYR:HB3 | 1.75 | 0.69 |
| 1:d1:198:GLY:HA3 | 1:d3:29:VAL:HG13 | 1.73 | 0.69 |
| 1:A3:94:ARG:NH1 | 1:A3:329:LEU:O | 2.25 | 0.69 |
| 1:B3:55:ARG:HG3 | 1:B3:216:ASP:HB3 | 1.74 | 0.68 |
| 1:A3:278:GLN:HE22 | 1:A3:297:ASN:H | 1.41 | 0.68 |
| 1:c3:227:MET:HE2 | 1:c3:326:ASN:HD21 | 1.59 | 0.68 |
| 1:f2:349:ALA:HA | 1:f1:373:PHE:HA | 1.75 | 0.68 |
| 1:A3:480:ILE:O | 5:TP:252:LYS:NZ | 2.27 | 0.68 |
| 1:c2:320:GLN:O | 1:c2:326:ASN:ND2 | 2.28 | 0.67 |
| 1:X3:155:ASP:HB2 | 1:X3:348:HIS:HE2 | 1.60 | 0.67 |
| 1:c2:421:VAL:HG22 | 1:c2:488:VAL:HB | 1.75 | 0.67 |

*Continued on next page...*

For Manuscript Review

*Continued from previous page...*

| Atom-1 | Atom-2 | Interatomic distance (Å) | Clash overlap (Å) |
|---|---|---|---|
| 1:B1:320:GLN:O | 1:B1:326:ASN:ND2 | 2.25 | 0.67 |
| 1:d2:312:ASN:OD1 | 1:d2:335:ASN:ND2 | 2.27 | 0.67 |
| 1:c3:518:MET:HE3 | 8:gc:80:PHE:HB3 | 1.75 | 0.67 |
| 1:c2:6:MET:HE3 | 1:c1:429:GLN:HG3 | 1.77 | 0.67 |
| 1:c3:90:SER:HB3 | 1:c3:156:PRO:HG3 | 1.77 | 0.67 |
| 1:c2:349:ALA:HA | 1:c1:373:PHE:HA | 1.77 | 0.67 |
| 1:c3:9:VAL:HG21 | 4:57:258:TYR:HB2 | 1.77 | 0.66 |
| 1:d2:115:ARG:HD2 | 1:d2:118:ARG:HB2 | 1.78 | 0.66 |
| 1:f3:109:VAL:HG22 | 1:f3:213:ILE:HG12 | 1.77 | 0.66 |
| 1:A1:162:ASP:OD2 | 1:A1:171:ARG:NH1 | 2.28 | 0.66 |
| 1:B2:314:ASP:HB3 | 1:B2:317:ALA:HB3 | 1.77 | 0.66 |
| 1:B2:423:LYS:HD3 | 1:B2:434:GLU:HG3 | 1.76 | 0.66 |
| 1:f1:94:ARG:NH1 | 1:f1:329:LEU:O | 2.28 | 0.66 |
| 1:c2:520:GLY:HA3 | 5:TQ:115:HIS:HB3 | 1.77 | 0.66 |
| 1:A3:308:GLN:NE2 | 1:A3:333:PRO:O | 2.26 | 0.66 |
| 1:c1:525:ASN:ND2 | 1:X3:427:ASN:O | 2.29 | 0.66 |
| 1:c3:44:GLN:NE2 | 1:c3:254:TYR:OH | 2.29 | 0.66 |
| 1:X2:421:VAL:HG22 | 1:X2:488:VAL:HB | 1.76 | 0.66 |
| 1:c2:313:VAL:HG12 | 1:c2:333:PRO:HA | 1.76 | 0.65 |
| 1:d2:442:LEU:HD21 | 1:d1:142:LYS:HE3 | 1.77 | 0.65 |
| 1:e3:277:LEU:HD11 | 1:e3:505:TYR:HB3 | 1.78 | 0.65 |
| 1:X3:213:ILE:HD12 | 1:X3:250:LEU:HD21 | 1.78 | 0.65 |
| 1:e2:90:SER:HB3 | 1:e2:156:PRO:HG3 | 1.78 | 0.65 |
| 1:X3:297:ASN:ND2 | 1:X3:525:ASN:OD1 | 2.30 | 0.65 |
| 1:X3:282:ASP:N | 1:X3:282:ASP:OD1 | 2.29 | 0.65 |
| 1:A1:420:VAL:HG12 | 1:A1:421:VAL:HG23 | 1.79 | 0.65 |
| 1:c1:90:SER:HB3 | 1:c1:156:PRO:HG3 | 1.79 | 0.65 |
| 1:e2:278:GLN:HE22 | 1:e2:297:ASN:H | 1.44 | 0.65 |
| 1:f2:375:ASN:ND2 | 1:f1:393:GLU:OE2 | 2.25 | 0.65 |
| 1:A2:282:ASP:OD1 | 1:A2:282:ASP:N | 2.27 | 0.65 |
| 1:A3:62:ARG:HD3 | 1:A3:203:LEU:HB3 | 1.79 | 0.65 |
| 1:B3:109:VAL:HG22 | 1:B3:213:ILE:HG12 | 1.79 | 0.65 |
| 1:X2:282:ASP:OD1 | 1:X2:282:ASP:N | 2.30 | 0.65 |
| 1:f1:421:VAL:HB | 9:uB:229:ILE:HD11 | 1.78 | 0.64 |
| 1:X2:427:ASN:ND2 | 1:X2:481:ASP:OD1 | 2.30 | 0.64 |
| 1:B2:109:VAL:HG22 | 1:B2:213:ILE:HG12 | 1.79 | 0.64 |
| 1:c3:427:ASN:HD21 | 1:c3:481:ASP:H | 1.46 | 0.64 |
| 1:d2:53:GLY:HA2 | 1:d2:217:ILE:O | 1.96 | 0.64 |
| 1:f1:349:ALA:HA | 1:f3:373:PHE:HA | 1.79 | 0.64 |
| 1:e1:355:GLN:HE21 | 1:e3:393:GLU:HA | 1.62 | 0.64 |
| 1:e3:421:VAL:HG22 | 1:e3:488:VAL:HB | 1.77 | 0.64 |

*Continued on next page...*

For Manuscript Review

Continued from previous page...

| Atom-1 | Atom-2 | Interatomic distance (Å) | Clash overlap (Å) |
|---|---|---|---|
| 1:c3:291:LYS:HE3 | 1:c3:293:LYS:HE2 | 1.79 | 0.64 |
| 1:d1:109:VAL:HG22 | 1:d1:213:ILE:HG12 | 1.78 | 0.64 |
| 1:d2:29:VAL:HB | 1:d3:198:GLY:HA3 | 1.78 | 0.64 |
| 1:d3:108:GLU:OE2 | 3:wD:65:ASN:ND2 | 2.31 | 0.64 |
| 1:f2:236:LYS:NZ | 1:f2:238:ASN:OD1 | 2.30 | 0.64 |
| 1:e3:90:SER:HB3 | 1:e3:156:PRO:HG3 | 1.79 | 0.64 |
| 1:d3:427:ASN:HD21 | 1:d3:481:ASP:H | 1.44 | 0.64 |
| 1:X2:195:ARG:NH2 | 1:X2:275:GLU:OE1 | 2.31 | 0.64 |
| 1:A2:309:PRO:HG3 | 1:A2:493:THR:HG23 | 1.80 | 0.64 |
| 1:c1:282:ASP:HB3 | 1:c1:501:LYS:HG3 | 1.78 | 0.64 |
| 1:e2:384:HIS:HB2 | 1:e2:387:PHE:HD2 | 1.64 | 0.63 |
| 1:A2:429:GLN:HG3 | 1:A3:6:MET:HE3 | 1.79 | 0.63 |
| 1:e3:427:ASN:HD21 | 1:e3:481:ASP:H | 1.45 | 0.63 |
| 9:uB:100:ILE:HD11 | 9:uB:117:ASN:HD22 | 1.63 | 0.63 |
| 1:X2:355:GLN:HE21 | 1:X1:393:GLU:HA | 1.63 | 0.63 |
| 1:e2:135:GLU:HB3 | 1:e3:451:ARG:HA | 1.79 | 0.63 |
| 1:c2:11:TYR:O | 1:c1:479:ARG:NH1 | 2.32 | 0.63 |
| 1:e3:170:VAL:HG12 | 1:e3:172:GLN:H | 1.62 | 0.63 |
| 1:f1:54:ARG:NH2 | 9:uB:149:TYR:O | 2.32 | 0.63 |
| 1:X3:109:VAL:HG12 | 1:X3:213:ILE:HG12 | 1.80 | 0.63 |
| 1:e1:83:LEU:HD13 | 1:e1:240:ALA:HA | 1.78 | 0.63 |
| 1:B1:162:ASP:OD1 | 1:B1:171:ARG:NH1 | 2.32 | 0.63 |
| 10:76:127:ASN:ND2 | 10:76:209:SER:OG | 2.31 | 0.63 |
| 1:B2:411:MET:HE1 | 1:B1:403:VAL:HA | 1.80 | 0.63 |
| 1:f1:425:GLN:NE2 | 1:f1:430:ASP:OD1 | 2.32 | 0.63 |
| 1:X2:277:LEU:HD11 | 1:X2:505:TYR:HB3 | 1.81 | 0.62 |
| 1:B3:282:ASP:N | 1:B3:282:ASP:OD1 | 2.31 | 0.62 |
| 1:f1:266:ALA:O | 7:he:136:TYR:OH | 2.16 | 0.62 |
| 8:gc:146:VAL:HG13 | 8:gc:213:MET:HE2 | 1.82 | 0.62 |
| 1:d2:262:ARG:NH2 | 5:TP:180:GLN:OE1 | 2.33 | 0.62 |
| 1:X2:206:LEU:HD21 | 1:X2:514:ILE:HD12 | 1.82 | 0.62 |
| 1:e3:83:LEU:HD13 | 1:e3:240:ALA:HA | 1.82 | 0.62 |
| 1:c1:421:VAL:HG22 | 1:c1:488:VAL:HB | 1.82 | 0.62 |
| 1:d3:53:GLY:HA2 | 1:d3:217:ILE:O | 1.99 | 0.62 |
| 1:e3:308:GLN:NE2 | 1:e3:333:PRO:O | 2.27 | 0.62 |
| 1:X3:53:GLY:HA2 | 1:X3:217:ILE:O | 1.99 | 0.62 |
| 1:A2:109:VAL:HG22 | 1:A2:213:ILE:HG12 | 1.80 | 0.62 |
| 1:B1:269:PRO:HB3 | 1:B1:515:MET:HG2 | 1.80 | 0.62 |
| 1:c2:420:VAL:HG12 | 1:c2:421:VAL:HG13 | 1.81 | 0.62 |
| 1:d2:198:GLY:HA3 | 1:d1:29:VAL:HG13 | 1.80 | 0.62 |
| 1:f2:417:ASN:ND2 | 1:f2:458:ASN:OD1 | 2.27 | 0.62 |

Continued on next page...

For Manuscript Review

*Continued from previous page...*

| Atom-1 | Atom-2 | Interatomic distance (Å) | Clash overlap (Å) |
|---|---|---|---|
| 1:e1:42:ILE:HG21 | 1:e1:63:ASN:HD22 | 1.65 | 0.62 |
| 1:f1:44:GLN:NE2 | 1:f1:254:TYR:OH | 2.33 | 0.61 |
| 1:A2:497:VAL:HB | 3:wD:115:MET:HE2 | 1.81 | 0.61 |
| 1:B2:384:HIS:HB2 | 1:B2:387:PHE:HD2 | 1.65 | 0.61 |
| 1:f1:301:LYS:N | 1:f1:507:VAL:O | 2.33 | 0.61 |
| 1:A2:271:GLU:OE2 | 1:A2:513:ARG:NH1 | 2.34 | 0.61 |
| 1:A3:421:VAL:HG23 | 1:A3:488:VAL:HB | 1.82 | 0.61 |
| 1:f2:476:ASN:ND2 | 1:f3:15:ASP:OD1 | 2.30 | 0.61 |
| 1:A1:109:VAL:HG22 | 1:A1:213:ILE:HG12 | 1.82 | 0.61 |
| 1:c1:278:GLN:NE2 | 1:c1:297:ASN:O | 2.33 | 0.61 |
| 1:X2:373:PHE:HA | 1:X3:349:ALA:HA | 1.82 | 0.61 |
| 1:c2:14:GLN:HG3 | 7:he:158:LYS:HZ2 | 1.66 | 0.61 |
| 1:c3:106:GLN:OE1 | 1:c3:118:ARG:NH2 | 2.34 | 0.61 |
| 1:e2:320:GLN:O | 1:e2:326:ASN:ND2 | 2.33 | 0.61 |
| 10:76:185:LEU:HD13 | 10:76:199:ARG:HD3 | 1.83 | 0.61 |
| 1:d2:373:PHE:HA | 1:d3:349:ALA:HA | 1.81 | 0.61 |
| 1:B2:320:GLN:O | 1:B2:326:ASN:ND2 | 2.32 | 0.61 |
| 1:B2:349:ALA:HA | 1:B1:373:PHE:HA | 1.83 | 0.61 |
| 1:c3:166:SER:O | 1:c3:171:ARG:NH2 | 2.34 | 0.61 |
| 1:f2:393:GLU:OE2 | 1:f3:375:ASN:ND2 | 2.34 | 0.61 |
| 1:X1:25:THR:HG22 | 1:X1:27:TRP:H | 1.66 | 0.61 |
| 1:X1:60:ILE:O | 1:X1:254:TYR:OH | 2.18 | 0.61 |
| 1:X1:320:GLN:O | 1:X1:326:ASN:ND2 | 2.33 | 0.61 |
| 1:X1:343:LEU:HD21 | 1:X1:408:ALA:HB1 | 1.82 | 0.61 |
| 1:d1:66:LEU:HB2 | 1:d1:255:ILE:HB | 1.84 | 0.60 |
| 1:A3:525:ASN:HD21 | 5:TP:255:PHE:HD2 | 1.48 | 0.60 |
| 1:X3:308:GLN:NE2 | 1:X3:333:PRO:O | 2.33 | 0.60 |
| 1:B2:373:PHE:HA | 1:B3:349:ALA:HA | 1.84 | 0.60 |
| 1:c2:277:LEU:HD11 | 1:c2:505:TYR:HB3 | 1.82 | 0.60 |
| 1:c3:513:ARG:HD3 | 1:c3:515:MET:HE3 | 1.82 | 0.60 |
| 1:f2:53:GLY:HA2 | 1:f2:217:ILE:O | 2.02 | 0.60 |
| 1:c1:115:ARG:NH2 | 1:X3:429:GLN:OE1 | 2.33 | 0.60 |
| 1:d3:42:ILE:HG21 | 1:d3:63:ASN:HD22 | 1.66 | 0.60 |
| 1:f2:301:LYS:N | 1:f2:507:VAL:O | 2.34 | 0.60 |
| 1:f1:384:HIS:HB2 | 1:f1:387:PHE:HD2 | 1.67 | 0.60 |
| 1:e2:355:GLN:NE2 | 1:e2:371:ASP:OD2 | 2.33 | 0.60 |
| 1:e2:375:ASN:ND2 | 1:e1:393:GLU:OE2 | 2.35 | 0.60 |
| 1:c1:503:ARG:NH1 | 9:uB:96:ASN:OD1 | 2.35 | 0.60 |
| 1:d3:94:ARG:NE | 1:d3:329:LEU:O | 2.27 | 0.60 |
| 1:d3:162:ASP:OD1 | 1:d3:171:ARG:NH1 | 2.34 | 0.60 |
| 1:d3:308:GLN:NE2 | 1:d3:333:PRO:O | 2.30 | 0.60 |

*Continued on next page...*

For Manuscript Review

*Continued from previous page...*

| Atom-1 | Atom-2 | Interatomic distance (Å) | Clash overlap (Å) |
|---|---|---|---|
| 1:B2:201:LEU:HD12 | 1:B2:202:PRO:HD2 | 1.84 | 0.60 |
| 1:A2:277:LEU:HD11 | 1:A2:505:TYR:HB3 | 1.84 | 0.60 |
| 1:A2:301:LYS:N | 1:A2:507:VAL:O | 2.32 | 0.60 |
| 1:c1:109:VAL:HG22 | 1:c1:213:ILE:HG12 | 1.83 | 0.59 |
| 1:c1:525:ASN:HD21 | 1:X3:429:GLN:HG2 | 1.67 | 0.59 |
| 1:A1:277:LEU:HD11 | 1:A1:505:TYR:HB3 | 1.84 | 0.59 |
| 1:e3:298:HIS:NE2 | 1:e3:525:ASN:OD1 | 2.34 | 0.59 |
| 1:B2:83:LEU:HD13 | 1:B2:240:ALA:HA | 1.85 | 0.59 |
| 1:B1:262:ARG:NH1 | 5:TP:259:GLY:O | 2.35 | 0.59 |
| 1:A3:513:ARG:NH1 | 5:TP:258:ASN:OD1 | 2.35 | 0.59 |
| 1:X1:83:LEU:HD13 | 1:X1:240:ALA:HA | 1.84 | 0.59 |
| 1:e3:420:VAL:HG12 | 1:e3:421:VAL:HG13 | 1.84 | 0.59 |
| 1:d2:423:LYS:HD3 | 1:d2:434:GLU:HG3 | 1.83 | 0.59 |
| 1:A2:447:GLN:NE2 | 1:A1:71:TYR:OH | 2.34 | 0.59 |
| 1:e1:423:LYS:HD3 | 1:e1:434:GLU:HB3 | 1.83 | 0.59 |
| 9:uB:167:ASN:HA | 9:uB:190:ILE:HD12 | 1.83 | 0.59 |
| 1:c3:76:LEU:HD21 | 1:c3:103:LEU:HD11 | 1.84 | 0.59 |
| 1:d1:420:VAL:HG12 | 1:d1:421:VAL:HG23 | 1.84 | 0.59 |
| 1:f3:51:ASP:HB3 | 1:f3:54:ARG:HE | 1.66 | 0.59 |
| 1:f3:384:HIS:HB2 | 1:f3:387:PHE:HD2 | 1.68 | 0.59 |
| 1:c1:349:ALA:HA | 1:c3:373:PHE:HA | 1.83 | 0.59 |
| 1:e2:402:PHE:HE2 | 1:e3:343:LEU:HD22 | 1.67 | 0.59 |
| 10:76:100:GLN:NE2 | 10:76:196:ASN:O | 2.35 | 0.59 |
| 1:B3:170:VAL:O | 1:B3:177:ARG:NH2 | 2.35 | 0.59 |
| 1:d3:149:GLN:NE2 | 1:d3:173:VAL:O | 2.35 | 0.59 |
| 1:A2:76:LEU:HD21 | 1:A2:103:LEU:HD11 | 1.84 | 0.59 |
| 7:he:59:ILE:HG12 | 7:he:92:MET:HE1 | 1.85 | 0.59 |
| 1:f1:434:GLU:OE1 | 9:uB:216:TYR:OH | 2.14 | 0.59 |
| 1:c3:118:ARG:NH1 | 1:c3:120:TYR:OH | 2.35 | 0.59 |
| 1:X3:301:LYS:N | 1:X3:507:VAL:O | 2.35 | 0.59 |
| 1:B2:297:ASN:O | 1:B2:478:SER:N | 2.36 | 0.58 |
| 1:B2:393:GLU:OE2 | 1:B3:375:ASN:ND2 | 2.31 | 0.58 |
| 1:e2:267:GLN:HB3 | 7:he:135:LYS:HG2 | 1.84 | 0.58 |
| 1:e2:425:GLN:OE1 | 1:e2:486:GLN:NE2 | 2.35 | 0.58 |
| 1:e3:67:ALA:HB3 | 1:e3:201:LEU:HB3 | 1.85 | 0.58 |
| 1:B1:277:LEU:HD11 | 1:B1:505:TYR:HB3 | 1.85 | 0.58 |
| 1:B3:201:LEU:HD12 | 1:B3:202:PRO:HD2 | 1.86 | 0.58 |
| 1:c3:263:ARG:NH2 | 7:he:148:ASP:OD1 | 2.35 | 0.58 |
| 1:c3:318:SER:O | 1:c3:326:ASN:ND2 | 2.31 | 0.58 |
| 1:X3:40:GLU:HB3 | 1:X3:256:PHE:HE2 | 1.69 | 0.58 |
| 1:e2:104:ILE:HG23 | 1:e2:107:VAL:HG23 | 1.83 | 0.58 |

*Continued on next page...*

For Manuscript Review

Continued from previous page...

| Atom-1 | Atom-2 | Interatomic distance (Å) | Clash overlap (Å) |
|---|---|---|---|
| 1:B2:479:ARG:NH2 | 5:TP:251:PRO:O | 2.32 | 0.58 |
| 1:c2:135:GLU:HB3 | 1:c3:451:ARG:HA | 1.85 | 0.58 |
| 1:f1:447:GLN:HE21 | 1:f3:71:TYR:HE2 | 1.50 | 0.58 |
| 1:c1:149:GLN:NE2 | 1:c1:173:VAL:O | 2.33 | 0.58 |
| 1:X3:107:VAL:HG22 | 1:X3:215:ILE:HG12 | 1.86 | 0.58 |
| 1:A3:67:ALA:HB3 | 1:A3:201:LEU:HB3 | 1.85 | 0.58 |
| 1:B2:301:LYS:N | 1:B2:507:VAL:O | 2.36 | 0.58 |
| 1:B1:90:SER:HB3 | 1:B1:156:PRO:HG3 | 1.85 | 0.58 |
| 1:d2:62:ARG:HH12 | 5:TP:171:GLY:HA3 | 1.66 | 0.58 |
| 1:B3:129:GLN:OE1 | 1:B3:451:ARG:NH1 | 2.36 | 0.58 |
| 1:d3:277:LEU:HD11 | 1:d3:505:TYR:HB3 | 1.84 | 0.58 |
| 1:f2:66:LEU:HB2 | 1:f2:255:ILE:HB | 1.86 | 0.58 |
| 1:f1:109:VAL:HG22 | 1:f1:213:ILE:HG12 | 1.84 | 0.58 |
| 1:f1:116:ILE:O | 1:f1:276:GLN:NE2 | 2.36 | 0.58 |
| 1:d2:164:PRO:O | 1:d3:491:ASN:ND2 | 2.36 | 0.58 |
| 1:f3:60:ILE:O | 1:f3:254:TYR:OH | 2.21 | 0.58 |
| 1:e2:280:THR:HG23 | 1:e2:504:VAL:HB | 1.84 | 0.58 |
| 1:B1:349:ALA:HA | 1:B3:373:PHE:HA | 1.86 | 0.58 |
| 1:X3:129:GLN:OE1 | 1:X3:451:ARG:NH1 | 2.36 | 0.58 |
| 1:B1:109:VAL:HG22 | 1:B1:213:ILE:HG12 | 1.85 | 0.58 |
| 1:d3:52:PHE:HE1 | 1:d3:103:LEU:HD22 | 1.69 | 0.58 |
| 1:f1:90:SER:HB3 | 1:f1:156:PRO:HG3 | 1.86 | 0.58 |
| 1:f1:106:GLN:NE2 | 1:f1:108:GLU:OE2 | 2.37 | 0.58 |
| 1:B3:90:SER:HB3 | 1:B3:156:PRO:HG3 | 1.86 | 0.57 |
| 1:A1:349:ALA:HA | 1:A3:373:PHE:HA | 1.84 | 0.57 |
| 1:c3:60:ILE:O | 1:c3:254:TYR:OH | 2.18 | 0.57 |
| 1:d2:375:ASN:ND2 | 1:d1:393:GLU:OE2 | 2.33 | 0.57 |
| 1:f2:51:ASP:OD1 | 1:f2:52:PHE:N | 2.34 | 0.57 |
| 1:f3:291:LYS:HG2 | 1:f3:486:GLN:HG2 | 1.86 | 0.57 |
| 1:X3:280:THR:O | 1:e2:486:GLN:NE2 | 2.36 | 0.57 |
| 1:A2:291:LYS:HD3 | 1:A2:486:GLN:HG2 | 1.85 | 0.57 |
| 1:e2:210:GLU:OE1 | 1:e2:212:LYS:NZ | 2.37 | 0.57 |
| 1:e1:44:GLN:NE2 | 1:e1:254:TYR:OH | 2.36 | 0.57 |
| 1:d3:207:GLN:HE22 | 1:A2:208:TYR:HD1 | 1.50 | 0.57 |
| 1:X2:284:SER:OG | 1:X2:501:LYS:NZ | 2.37 | 0.57 |
| 1:c1:198:GLY:HA3 | 1:c3:29:VAL:HG13 | 1.85 | 0.57 |
| 1:d2:355:GLN:NE2 | 1:d2:396:VAL:O | 2.37 | 0.57 |
| 1:d1:5:LEU:HD22 | 5:TP:180:GLN:HB2 | 1.86 | 0.57 |
| 1:d3:278:GLN:HE22 | 1:d3:297:ASN:H | 1.52 | 0.57 |
| 1:f2:104:ILE:HD12 | 1:f2:124:MET:HE1 | 1.86 | 0.57 |
| 1:X2:349:ALA:HA | 1:X1:373:PHE:HA | 1.86 | 0.57 |

Continued on next page...

For Manuscript Review

*Continued from previous page...*

| Atom-1 | Atom-2 | Interatomic distance (Å) | Clash overlap (Å) |
|---|---|---|---|
| 1:X3:303:LEU:HD13 | 1:X3:462:PHE:HE2 | 1.68 | 0.57 |
| 1:d2:190:GLN:O | 1:d2:195:ARG:NH1 | 2.38 | 0.57 |
| 1:A3:44:GLN:NE2 | 1:A3:254:TYR:OH | 2.36 | 0.57 |
| 1:d2:206:LEU:HD21 | 1:d2:514:ILE:HD12 | 1.85 | 0.57 |
| 1:f2:291:LYS:HG2 | 1:f2:486:GLN:HG2 | 1.85 | 0.57 |
| 1:f2:308:GLN:NE2 | 1:f2:333:PRO:O | 2.37 | 0.57 |
| 1:X1:67:ALA:HB3 | 1:X1:201:LEU:HB2 | 1.85 | 0.57 |
| 1:B1:442:LEU:HD21 | 1:B3:142:LYS:HE3 | 1.87 | 0.57 |
| 1:d2:299:PRO:HA | 1:d2:476:ASN:HA | 1.86 | 0.57 |
| 1:d2:470:GLN:NE2 | 1:d1:32:ARG:O | 2.33 | 0.57 |
| 1:d3:109:VAL:HG22 | 1:d3:213:ILE:HG12 | 1.84 | 0.57 |
| 1:X3:320:GLN:O | 1:X3:326:ASN:ND2 | 2.37 | 0.57 |
| 1:X3:525:ASN:HD21 | 1:e2:429:GLN:HG3 | 1.69 | 0.57 |
| 1:e2:83:LEU:HD13 | 1:e2:240:ALA:HA | 1.86 | 0.57 |
| 1:d1:44:GLN:NE2 | 1:d1:254:TYR:OH | 2.38 | 0.57 |
| 1:f1:277:LEU:HD11 | 1:f1:505:TYR:HB3 | 1.86 | 0.57 |
| 1:c1:429:GLN:OE1 | 1:e2:115:ARG:NH2 | 2.37 | 0.57 |
| 1:A2:349:ALA:HA | 1:A1:373:PHE:HA | 1.85 | 0.57 |
| 1:e2:302:GLU:OE1 | 1:e2:449:HIS:NE2 | 2.37 | 0.57 |
| 1:e3:417:ASN:HD21 | 1:e3:419:VAL:HB | 1.70 | 0.57 |
| 1:B1:307:VAL:HG22 | 1:B1:502:VAL:HG22 | 1.87 | 0.57 |
| 1:f3:44:GLN:NE2 | 1:f3:254:TYR:OH | 2.38 | 0.56 |
| 1:f3:301:LYS:N | 1:f3:507:VAL:O | 2.38 | 0.56 |
| 1:X2:83:LEU:HD13 | 1:X2:240:ALA:HA | 1.86 | 0.56 |
| 1:A2:313:VAL:HG12 | 1:A2:333:PRO:HA | 1.87 | 0.56 |
| 1:A3:52:PHE:HE1 | 1:A3:103:LEU:HD22 | 1.70 | 0.56 |
| 1:f3:161:VAL:HA | 1:f3:177:ARG:HD2 | 1.85 | 0.56 |
| 1:X3:53:GLY:CA | 1:X3:217:ILE:O | 2.52 | 0.56 |
| 1:B1:280:THR:HG21 | 1:B1:295:ASN:HB3 | 1.87 | 0.56 |
| 1:d3:307:VAL:HG22 | 1:d3:502:VAL:HG22 | 1.88 | 0.56 |
| 1:X3:384:HIS:HB2 | 1:X3:387:PHE:HD2 | 1.68 | 0.56 |
| 1:A2:116:ILE:O | 1:A2:276:GLN:NE2 | 2.36 | 0.56 |
| 1:A2:373:PHE:HA | 1:A3:349:ALA:HA | 1.85 | 0.56 |
| 1:c2:115:ARG:HD2 | 1:c2:118:ARG:HB2 | 1.86 | 0.56 |
| 1:c1:160:ALA:O | 1:c1:177:ARG:NH1 | 2.37 | 0.56 |
| 1:d2:232:GLY:O | 1:d2:326:ASN:ND2 | 2.39 | 0.56 |
| 1:d1:369:PHE:HB2 | 1:d1:372:PRO:HB3 | 1.88 | 0.56 |
| 1:B2:481:ASP:OD1 | 1:B2:482:ASN:N | 2.37 | 0.56 |
| 1:X1:304:ILE:HG12 | 1:X1:459:VAL:HG22 | 1.86 | 0.56 |
| 1:c3:313:VAL:HG12 | 1:c3:333:PRO:HA | 1.88 | 0.56 |
| 1:f2:29:VAL:HB | 1:f3:198:GLY:HA3 | 1.87 | 0.56 |

*Continued on next page...*

For Manuscript Review

*Continued from previous page...*

| Atom-1 | Atom-2 | Interatomic distance (Å) | Clash overlap (Å) |
|---|---|---|---|
| 1:c2:513:ARG:NH2 | 1:c3:17:TYR:OH | 2.38 | 0.56 |
| 1:c1:122:ASP:OD1 | 1:c1:336:TYR:OH | 2.20 | 0.56 |
| 1:c1:491:ASN:HB2 | 1:c3:165:CYS:HA | 1.87 | 0.56 |
| 1:f1:384:HIS:HB2 | 1:f1:387:PHE:CD2 | 2.40 | 0.56 |
| 1:X1:81:GLN:NE2 | 1:X1:178:LYS:O | 2.34 | 0.56 |
| 1:A2:257:LEU:HB2 | 1:A2:262:ARG:HG3 | 1.88 | 0.56 |
| 1:d1:59:THR:OG1 | 1:d1:212:LYS:NZ | 2.39 | 0.56 |
| 1:f1:75:THR:HB | 1:f1:246:ALA:HB3 | 1.88 | 0.56 |
| 1:X2:282:ASP:HB3 | 1:X2:503:ARG:HG2 | 1.88 | 0.56 |
| 1:e2:349:ALA:HA | 1:e1:373:PHE:HA | 1.87 | 0.56 |
| 1:c2:307:VAL:HG22 | 1:c2:502:VAL:HG22 | 1.88 | 0.56 |
| 1:c1:51:ASP:OD1 | 1:c1:52:PHE:N | 2.36 | 0.56 |
| 1:d1:90:SER:HB3 | 1:d1:156:PRO:HG3 | 1.87 | 0.56 |
| 1:d3:53:GLY:CA | 1:d3:217:ILE:O | 2.53 | 0.56 |
| 1:f3:111:ILE:HG12 | 1:f3:211:VAL:HG22 | 1.88 | 0.56 |
| 1:c1:17:TYR:OH | 1:c1:21:ASN:OD1 | 2.18 | 0.55 |
| 1:A2:162:ASP:OD2 | 1:A2:171:ARG:NH1 | 2.37 | 0.55 |
| 1:e2:11:TYR:HD2 | 5:TP:207:GLY:HA2 | 1.70 | 0.55 |
| 1:B2:90:SER:HB3 | 1:B2:156:PRO:HG3 | 1.89 | 0.55 |
| 1:c2:301:LYS:N | 1:c2:507:VAL:O | 2.39 | 0.55 |
| 1:c3:417:ASN:ND2 | 1:c3:458:ASN:OD1 | 2.37 | 0.55 |
| 1:d3:384:HIS:HB2 | 1:d3:387:PHE:HD2 | 1.70 | 0.55 |
| 1:X3:277:LEU:HD21 | 1:X3:505:TYR:HB3 | 1.88 | 0.55 |
| 1:A2:492:ALA:HB2 | 1:A1:171:ARG:HE | 1.72 | 0.55 |
| 1:B1:464:LEU:HB2 | 1:B1:472:SER:HB2 | 1.89 | 0.55 |
| 1:f2:206:LEU:HD21 | 1:f2:514:ILE:HD12 | 1.87 | 0.55 |
| 1:X2:122:ASP:OD1 | 1:X2:336:TYR:OH | 2.16 | 0.55 |
| 1:A2:165:CYS:SG | 3:wD:43:PHE:HB3 | 2.45 | 0.55 |
| 1:X2:393:GLU:OE2 | 1:X3:375:ASN:ND2 | 2.39 | 0.55 |
| 1:X1:107:VAL:HG22 | 1:X1:215:ILE:HG12 | 1.89 | 0.55 |
| 1:B2:271:GLU:OE2 | 1:B2:513:ARG:NH1 | 2.39 | 0.55 |
| 1:B1:110:GLU:HG2 | 1:B1:115:ARG:HA | 1.88 | 0.55 |
| 1:X3:83:LEU:HD13 | 1:X3:240:ALA:HA | 1.88 | 0.55 |
| 1:c2:31:TYR:OH | 1:c1:17:TYR:O | 2.18 | 0.55 |
| 1:c3:83:LEU:HD13 | 1:c3:240:ALA:HA | 1.87 | 0.55 |
| 1:f2:53:GLY:CA | 1:f2:217:ILE:O | 2.54 | 0.55 |
| 1:f2:402:PHE:HE2 | 1:f3:343:LEU:HD22 | 1.70 | 0.55 |
| 1:A2:110:GLU:HG2 | 1:A2:115:ARG:HA | 1.87 | 0.55 |
| 1:e2:308:GLN:NE2 | 1:e2:333:PRO:O | 2.31 | 0.55 |
| 1:e2:309:PRO:HG3 | 1:e2:493:THR:HG23 | 1.88 | 0.55 |
| 1:d2:53:GLY:CA | 1:d2:217:ILE:O | 2.54 | 0.55 |

*Continued on next page...*

For Manuscript Review

*Continued from previous page...*

| Atom-1 | Atom-2 | Interatomic distance (Å) | Clash overlap (Å) |
|---|---|---|---|
| 1:d2:257:LEU:O | 1:d2:262:ARG:NH1 | 2.40 | 0.55 |
| 1:d3:227:MET:HE2 | 1:d3:326:ASN:HD21 | 1.71 | 0.55 |
| 1:f2:107:VAL:HG23 | 1:f2:215:ILE:HG12 | 1.89 | 0.55 |
| 1:e3:236:LYS:HD3 | 1:e3:319:LEU:HD23 | 1.88 | 0.55 |
| 1:f3:308:GLN:NE2 | 1:f3:333:PRO:O | 2.33 | 0.55 |
| 8:gc:262:LYS:HB3 | 8:gc:266:SER:HA | 1.89 | 0.55 |
| 1:B1:293:LYS:HD3 | 1:B1:484:THR:HG22 | 1.88 | 0.54 |
| 1:B3:277:LEU:HD11 | 1:B3:505:TYR:HB3 | 1.88 | 0.54 |
| 1:c1:301:LYS:N | 1:c1:507:VAL:O | 2.40 | 0.54 |
| 1:f3:421:VAL:HG22 | 1:f3:488:VAL:HB | 1.89 | 0.54 |
| 1:e2:110:GLU:HG2 | 1:e2:115:ARG:HA | 1.88 | 0.54 |
| 1:B3:6:MET:HE1 | 1:A3:525:ASN:HB3 | 1.88 | 0.54 |
| 1:B3:236:LYS:HD3 | 1:B3:319:LEU:HD13 | 1.89 | 0.54 |
| 1:B3:307:VAL:HG22 | 1:B3:502:VAL:HG22 | 1.88 | 0.54 |
| 1:d2:278:GLN:HG3 | 1:d2:506:ALA:HB3 | 1.89 | 0.54 |
| 1:d1:44:GLN:O | 1:d1:249:SER:OG | 2.24 | 0.54 |
| 1:A1:293:LYS:HD3 | 1:A1:484:THR:HG22 | 1.89 | 0.54 |
| 1:A2:63:ASN:HD21 | 1:e1:428:GLY:HA3 | 1.71 | 0.54 |
| 1:A2:403:VAL:HA | 1:A3:411:MET:HE1 | 1.89 | 0.54 |
| 4:57:183:PRO:HA | 4:57:246:ILE:HG13 | 1.89 | 0.54 |
| 1:X2:32:ARG:O | 1:X3:470:GLN:NE2 | 2.41 | 0.54 |
| 1:X2:104:ILE:HG23 | 1:X2:107:VAL:HG23 | 1.88 | 0.54 |
| 1:A2:375:ASN:ND2 | 1:A1:393:GLU:OE2 | 2.30 | 0.54 |
| 1:B2:402:PHE:HE2 | 1:B3:343:LEU:HD22 | 1.71 | 0.54 |
| 1:c1:469:HIS:O | 1:c1:509:TYR:OH | 2.21 | 0.54 |
| 1:d1:147:THR:O | 1:d1:151:THR:OG1 | 2.26 | 0.54 |
| 1:X2:7:GLN:OE1 | 1:X1:431:ARG:NH2 | 2.41 | 0.54 |
| 1:A3:116:ILE:HG23 | 1:A3:521:LEU:HD21 | 1.88 | 0.54 |
| 1:d2:320:GLN:O | 1:d2:326:ASN:ND2 | 2.40 | 0.54 |
| 1:X1:447:GLN:HB2 | 1:X3:69:ARG:HE | 1.71 | 0.54 |
| 1:e3:147:THR:O | 1:e3:151:THR:OG1 | 2.24 | 0.54 |
| 1:c1:111:ILE:HG12 | 1:c1:211:VAL:HG22 | 1.90 | 0.54 |
| 1:d1:349:ALA:HA | 1:d3:373:PHE:HA | 1.90 | 0.54 |
| 10:76:140:ASP:HB3 | 10:76:174:LYS:HD3 | 1.90 | 0.54 |
| 1:B1:301:LYS:N | 1:B1:507:VAL:O | 2.39 | 0.54 |
| 1:B3:155:ASP:HB2 | 1:B3:348:HIS:HE2 | 1.72 | 0.54 |
| 1:c2:104:ILE:HG23 | 1:c2:107:VAL:HG23 | 1.88 | 0.54 |
| 1:c2:375:ASN:ND2 | 1:c1:393:GLU:OE2 | 2.41 | 0.54 |
| 1:c2:394:SER:HB2 | 1:c3:396:VAL:HB | 1.90 | 0.54 |
| 8:gc:155:LEU:HD12 | 8:gc:205:GLY:HA3 | 1.90 | 0.54 |
| 1:B3:83:LEU:HD13 | 1:B3:240:ALA:HA | 1.88 | 0.54 |

*Continued on next page...*

For Manuscript Review

*Continued from previous page...*

| Atom-1 | Atom-2 | Interatomic distance (Å) | Clash overlap (Å) |
|---|---|---|---|
| 1:c1:308:GLN:NE2 | 1:c1:333:PRO:O | 2.37 | 0.54 |
| 1:X2:420:VAL:HG12 | 1:X2:421:VAL:HG13 | 1.88 | 0.54 |
| 1:X1:6:MET:HE1 | 1:X3:429:GLN:HB2 | 1.90 | 0.54 |
| 1:A2:147:THR:O | 1:A2:151:THR:OG1 | 2.22 | 0.54 |
| 1:d2:447:GLN:NE2 | 1:d1:71:TYR:OH | 2.41 | 0.54 |
| 1:X2:109:VAL:HG22 | 1:X2:213:ILE:HG12 | 1.90 | 0.54 |
| 1:X1:11:TYR:O | 1:X3:479:ARG:NH1 | 2.41 | 0.54 |
| 1:A1:17:TYR:OH | 1:A3:513:ARG:NH2 | 2.40 | 0.54 |
| 1:A1:53:GLY:HA2 | 1:A1:217:ILE:O | 2.07 | 0.54 |
| 1:B2:137:GLU:OE1 | 1:B2:451:ARG:NH1 | 2.31 | 0.53 |
| 1:c2:355:GLN:HE22 | 1:c2:397:SER:HA | 1.72 | 0.53 |
| 1:c2:423:LYS:HE3 | 1:c2:425:GLN:HE21 | 1.73 | 0.53 |
| 1:d2:66:LEU:HB2 | 1:d2:255:ILE:HB | 1.90 | 0.53 |
| 1:d1:343:LEU:HD22 | 1:d3:402:PHE:HE2 | 1.73 | 0.53 |
| 1:B3:388:GLU:HG2 | 1:B3:390:THR:H | 1.72 | 0.53 |
| 1:c3:299:PRO:HA | 1:c3:476:ASN:HA | 1.89 | 0.53 |
| 1:d2:109:VAL:HG12 | 1:d2:213:ILE:HG12 | 1.89 | 0.53 |
| 1:d2:301:LYS:N | 1:d2:507:VAL:O | 2.42 | 0.53 |
| 1:f3:83:LEU:HD13 | 1:f3:240:ALA:HA | 1.89 | 0.53 |
| 1:X1:15:ASP:OD1 | 1:X3:476:ASN:ND2 | 2.42 | 0.53 |
| 1:A3:313:VAL:HG12 | 1:A3:333:PRO:HA | 1.90 | 0.53 |
| 1:X3:343:LEU:HD21 | 1:X3:408:ALA:HB1 | 1.88 | 0.53 |
| 1:A2:236:LYS:HD2 | 3:wD:97:ILE:HD12 | 1.89 | 0.53 |
| 1:B1:308:GLN:NE2 | 1:B1:333:PRO:O | 2.40 | 0.53 |
| 1:B3:320:GLN:O | 1:B3:326:ASN:ND2 | 2.37 | 0.53 |
| 1:c2:299:PRO:HA | 1:c2:476:ASN:HA | 1.90 | 0.53 |
| 1:d1:15:ASP:OD1 | 1:d3:476:ASN:ND2 | 2.41 | 0.53 |
| 1:d1:307:VAL:HG22 | 1:d1:502:VAL:HG22 | 1.90 | 0.53 |
| 1:X1:422:ALA:HB3 | 1:X1:440:PHE:CE2 | 2.43 | 0.53 |
| 1:f3:227:MET:HE2 | 1:f3:326:ASN:HD21 | 1.74 | 0.53 |
| 1:X2:278:GLN:HG3 | 1:X2:506:ALA:HB3 | 1.91 | 0.53 |
| 1:B3:50:ALA:HB3 | 1:B3:248:ALA:HB3 | 1.90 | 0.53 |
| 1:c3:524:SER:OG | 8:gc:83:ARG:NH1 | 2.42 | 0.53 |
| 1:d1:375:ASN:ND2 | 1:d3:393:GLU:OE2 | 2.40 | 0.53 |
| 1:d3:90:SER:HB3 | 1:d3:156:PRO:HG3 | 1.90 | 0.53 |
| 1:d3:116:ILE:HG23 | 1:d3:521:LEU:HD21 | 1.90 | 0.53 |
| 1:f2:134:LYS:HG2 | 1:f1:135:GLU:HG3 | 1.90 | 0.53 |
| 1:B2:307:VAL:HG22 | 1:B2:502:VAL:HG22 | 1.91 | 0.53 |
| 1:f2:332:GLN:OE1 | 1:f2:335:ASN:ND2 | 2.37 | 0.53 |
| 1:e2:52:PHE:HE1 | 1:e2:103:LEU:HD22 | 1.73 | 0.53 |
| 1:e2:206:LEU:HD21 | 1:e2:514:ILE:HD12 | 1.91 | 0.53 |

*Continued on next page...*

For Manuscript Review

Continued from previous page...

| Atom-1 | Atom-2 | Interatomic distance (Å) | Clash overlap (Å) |
|---|---|---|---|
| 1:c1:277:LEU:HD11 | 1:c1:505:TYR:HB3 | 1.91 | 0.53 |
| 1:f2:204:ILE:O | 1:f2:207:GLN:NE2 | 2.42 | 0.53 |
| 1:X1:104:ILE:HD12 | 1:X1:124:MET:HE1 | 1.91 | 0.53 |
| 1:c1:328:LEU:HD21 | 1:c3:368:PHE:HB3 | 1.90 | 0.53 |
| 1:c1:442:LEU:HD23 | 1:c1:452:ASN:HD21 | 1.74 | 0.53 |
| 1:d2:136:GLN:NE2 | 1:d3:445:PRO:O | 2.43 | 0.53 |
| 1:f2:210:GLU:OE1 | 1:f2:212:LYS:NZ | 2.35 | 0.53 |
| 1:f1:114:GLN:HE22 | 1:f1:521:LEU:H | 1.56 | 0.53 |
| 1:X2:95:TRP:CD1 | 1:X2:180:LEU:HD13 | 2.44 | 0.53 |
| 1:X2:308:GLN:NE2 | 1:X2:333:PRO:O | 2.30 | 0.53 |
| 1:X1:109:VAL:HG22 | 1:X1:213:ILE:HG12 | 1.89 | 0.53 |
| 1:A2:7:GLN:NE2 | 1:A1:474:THR:O | 2.42 | 0.53 |
| 1:f3:101:GLU:OE2 | 1:f3:140:TYR:OH | 2.20 | 0.52 |
| 1:X1:273:LEU:O | 1:X1:469:HIS:NE2 | 2.40 | 0.52 |
| 1:e3:423:LYS:HE3 | 1:e3:430:ASP:HB3 | 1.90 | 0.52 |
| 1:d3:236:LYS:HD2 | 1:d3:319:LEU:HD13 | 1.92 | 0.52 |
| 1:X1:213:ILE:HD12 | 1:X1:250:LEU:HD21 | 1.91 | 0.52 |
| 1:X1:349:ALA:HA | 1:X3:373:PHE:HA | 1.91 | 0.52 |
| 6:ze:106:LEU:HD22 | 6:ze:111:LEU:HD11 | 1.92 | 0.52 |
| 2:vC:100:TYR:OH | 2:vC:104:ARG:NH1 | 2.43 | 0.52 |
| 1:B1:355:GLN:HE21 | 1:B3:393:GLU:HA | 1.75 | 0.52 |
| 1:d3:114:GLN:HG2 | 1:e1:525:ASN:HB2 | 1.92 | 0.52 |
| 1:d3:172:GLN:NE2 | 1:d3:367:GLY:O | 2.42 | 0.52 |
| 1:A3:369:PHE:HB2 | 1:A3:372:PRO:HB3 | 1.92 | 0.52 |
| 1:c3:110:GLU:HB3 | 1:c3:115:ARG:HA | 1.91 | 0.52 |
| 1:c3:277:LEU:HD11 | 1:c3:505:TYR:HB3 | 1.91 | 0.52 |
| 1:X3:104:ILE:HG23 | 1:X3:107:VAL:HG23 | 1.92 | 0.52 |
| 1:c2:6:MET:SD | 1:e2:525:ASN:ND2 | 2.83 | 0.52 |
| 1:d2:402:PHE:HE2 | 1:d3:343:LEU:HD22 | 1.74 | 0.52 |
| 1:d3:62:ARG:NH1 | 1:d3:206:LEU:O | 2.41 | 0.52 |
| 1:A2:104:ILE:HD12 | 1:A2:124:MET:HE1 | 1.92 | 0.52 |
| 1:e1:163:GLY:N | 1:e1:166:SER:OG | 2.32 | 0.52 |
| 1:B1:262:ARG:NH2 | 5:TP:261:GLN:O | 2.43 | 0.52 |
| 1:f1:138:ARG:HG2 | 1:f1:142:LYS:HE2 | 1.91 | 0.52 |
| 1:X2:157:GLU:OE2 | 1:X2:357:SER:OG | 2.27 | 0.52 |
| 1:X2:299:PRO:HA | 1:X2:476:ASN:HA | 1.92 | 0.52 |
| 1:A2:134:LYS:HG2 | 1:A1:135:GLU:HG3 | 1.92 | 0.52 |
| 10:76:196:ASN:O | 10:76:196:ASN:ND2 | 2.43 | 0.52 |
| 1:d2:308:GLN:NE2 | 1:d2:333:PRO:O | 2.35 | 0.52 |
| 1:A1:343:LEU:HD21 | 1:A1:408:ALA:HB1 | 1.92 | 0.52 |
| 1:e2:3:GLY:HA2 | 1:e1:431:ARG:HA | 1.92 | 0.52 |

Continued on next page...

For Manuscript Review

*Continued from previous page...*

| Atom-1 | Atom-2 | Interatomic distance (Å) | Clash overlap (Å) |
|---|---|---|---|
| 1:e1:169:SER:O | 1:e1:171:ARG:N | 2.42 | 0.52 |
| 1:c1:83:LEU:HD13 | 1:c1:240:ALA:HA | 1.90 | 0.52 |
| 1:c1:269:PRO:HB3 | 1:c1:515:MET:HG2 | 1.92 | 0.52 |
| 1:d1:128:CYS:SG | 1:d1:190:GLN:NE2 | 2.83 | 0.52 |
| 1:A2:94:ARG:NE | 1:A2:329:LEU:O | 2.28 | 0.52 |
| 1:A2:299:PRO:HA | 1:A2:476:ASN:HA | 1.91 | 0.52 |
| 1:A1:417:ASN:ND2 | 1:A1:458:ASN:OD1 | 2.41 | 0.52 |
| 5:TP:235:ASP:HA | 5:TP:238:LYS:HE2 | 1.92 | 0.52 |
| 1:c1:524:SER:HB3 | 1:X1:6:MET:HE2 | 1.91 | 0.52 |
| 1:f2:277:LEU:HD11 | 1:f2:505:TYR:HB3 | 1.92 | 0.52 |
| 1:X2:32:ARG:H | 1:X3:470:GLN:HE21 | 1.58 | 0.52 |
| 1:A2:59:THR:OG1 | 1:A2:212:LYS:NZ | 2.43 | 0.52 |
| 1:e2:304:ILE:HG12 | 1:e2:459:VAL:HG22 | 1.92 | 0.52 |
| 1:B3:314:ASP:OD1 | 9:uB:248:THR:OG1 | 2.26 | 0.51 |
| 1:c2:109:VAL:HG22 | 1:c2:213:ILE:HG12 | 1.92 | 0.51 |
| 1:d2:73:GLN:HE21 | 1:d2:186:TYR:HE2 | 1.57 | 0.51 |
| 1:f1:510:ASN:OD1 | 1:f1:511:VAL:N | 2.42 | 0.51 |
| 1:X1:55:ARG:HH21 | 1:X1:214:ASN:HD22 | 1.58 | 0.51 |
| 1:A3:83:LEU:HD13 | 1:A3:240:ALA:HA | 1.91 | 0.51 |
| 1:c2:108:GLU:HG3 | 1:c2:214:ASN:HB2 | 1.92 | 0.51 |
| 1:c3:301:LYS:N | 1:c3:507:VAL:O | 2.28 | 0.51 |
| 1:f1:271:GLU:HG2 | 1:f1:513:ARG:HG2 | 1.91 | 0.51 |
| 1:f3:320:GLN:O | 1:f3:326:ASN:ND2 | 2.43 | 0.51 |
| 1:X3:495:GLN:O | 1:X3:498:SER:OG | 2.24 | 0.51 |
| 1:A1:342:ALA:HB3 | 1:A1:413:CYS:HB3 | 1.91 | 0.51 |
| 1:e1:343:LEU:HD21 | 1:e1:408:ALA:HB1 | 1.92 | 0.51 |
| 6:ze:121:ASN:OD1 | 7:he:93:ARG:NH2 | 2.43 | 0.51 |
| 1:f2:101:GLU:OE2 | 1:f2:140:TYR:OH | 2.26 | 0.51 |
| 1:A1:147:THR:O | 1:A1:151:THR:OG1 | 2.26 | 0.51 |
| 9:uB:4:ILE:HD13 | 9:uB:12:ILE:HG21 | 1.93 | 0.51 |
| 1:c2:134:LYS:HG2 | 1:c1:135:GLU:HG3 | 1.93 | 0.51 |
| 1:c2:343:LEU:HD22 | 1:c1:402:PHE:HE2 | 1.75 | 0.51 |
| 1:c1:314:ASP:OD2 | 9:uB:102:ASN:ND2 | 2.43 | 0.51 |
| 1:f2:257:LEU:HD13 | 1:f2:261:GLU:HG3 | 1.92 | 0.51 |
| 1:f2:442:LEU:HD23 | 1:f2:452:ASN:HD21 | 1.75 | 0.51 |
| 1:e3:427:ASN:ND2 | 1:e3:481:ASP:H | 2.08 | 0.51 |
| 1:d2:90:SER:HB3 | 1:d2:156:PRO:HG3 | 1.92 | 0.51 |
| 1:f1:304:ILE:HG12 | 1:f1:459:VAL:HG22 | 1.92 | 0.51 |
| 1:A3:277:LEU:HD11 | 1:A3:505:TYR:HB3 | 1.92 | 0.51 |
| 1:c2:464:LEU:HB2 | 1:c2:472:SER:HB2 | 1.92 | 0.51 |
| 1:c3:155:ASP:HB2 | 1:c3:348:HIS:HE2 | 1.76 | 0.51 |

*Continued on next page...*

For Manuscript Review

*Continued from previous page...*

| Atom-1 | Atom-2 | Interatomic distance (Å) | Clash overlap (Å) |
|---|---|---|---|
| 1:c3:420:VAL:HG12 | 1:c3:421:VAL:HG13 | 1.92 | 0.51 |
| 1:f2:373:PHE:HA | 1:f3:349:ALA:HA | 1.93 | 0.51 |
| 1:X1:277:LEU:HD11 | 1:X1:505:TYR:HB3 | 1.93 | 0.51 |
| 1:A2:393:GLU:OE2 | 1:A3:375:ASN:ND2 | 2.30 | 0.51 |
| 1:A3:147:THR:O | 1:A3:151:THR:OG1 | 2.25 | 0.51 |
| 1:e1:161:VAL:HG12 | 1:e1:177:ARG:HD3 | 1.92 | 0.51 |
| 9:uB:25:ASN:HB3 | 9:uB:26:PRO:HD3 | 1.91 | 0.51 |
| 1:B2:115:ARG:HD2 | 1:B2:118:ARG:HB2 | 1.91 | 0.51 |
| 1:X1:9:VAL:HG12 | 7:he:162:LEU:HG | 1.91 | 0.51 |
| 1:X3:417:ASN:ND2 | 1:X3:458:ASN:OD1 | 2.31 | 0.51 |
| 1:c2:69:ARG:H | 1:c2:253:ASP:HB2 | 1.76 | 0.51 |
| 1:e2:136:GLN:NE2 | 1:e3:449:HIS:O | 2.44 | 0.51 |
| 1:e2:157:GLU:OE2 | 1:e2:357:SER:OG | 2.28 | 0.51 |
| 1:B2:166:SER:H | 1:B3:491:ASN:HD22 | 1.59 | 0.51 |
| 1:c1:42:ILE:HD12 | 1:c1:63:ASN:HB3 | 1.92 | 0.51 |
| 1:d2:393:GLU:OE2 | 1:d3:375:ASN:ND2 | 2.27 | 0.51 |
| 1:f2:90:SER:HB3 | 1:f2:156:PRO:HG3 | 1.92 | 0.51 |
| 1:X3:195:ARG:NH2 | 1:X3:275:GLU:OE1 | 2.33 | 0.51 |
| 1:e3:301:LYS:N | 1:e3:507:VAL:O | 2.36 | 0.51 |
| 5:TP:253:ASN:ND2 | 5:TP:254:SER:O | 2.44 | 0.51 |
| 1:B2:110:GLU:HG2 | 1:B2:115:ARG:HA | 1.92 | 0.51 |
| 1:B3:422:ALA:HB3 | 1:B3:440:PHE:CE2 | 2.45 | 0.51 |
| 1:c2:402:PHE:HE2 | 1:c3:343:LEU:HD22 | 1.76 | 0.51 |
| 1:X2:66:LEU:HB2 | 1:X2:255:ILE:HB | 1.93 | 0.51 |
| 1:A1:176:PRO:HD2 | 1:A1:362:VAL:HG11 | 1.92 | 0.51 |
| 1:A1:227:MET:HE2 | 1:A1:326:ASN:HD21 | 1.76 | 0.51 |
| 4:57:245:VAL:HG12 | 4:57:247:PRO:HD2 | 1.93 | 0.51 |
| 1:c3:99:PRO:HD2 | 1:c3:145:GLY:O | 2.12 | 0.50 |
| 1:c3:299:PRO:HG2 | 1:c3:509:TYR:HB2 | 1.93 | 0.50 |
| 1:d2:147:THR:O | 1:d2:151:THR:OG1 | 2.26 | 0.50 |
| 1:d3:315:TYR:HE1 | 1:d3:333:PRO:HG3 | 1.76 | 0.50 |
| 1:e2:32:ARG:O | 1:e3:470:GLN:NE2 | 2.38 | 0.50 |
| 1:e1:349:ALA:HA | 1:e3:373:PHE:HA | 1.94 | 0.50 |
| 1:B1:107:VAL:HG23 | 1:B1:215:ILE:HG12 | 1.94 | 0.50 |
| 1:d2:442:LEU:HD23 | 1:d2:452:ASN:HD21 | 1.76 | 0.50 |
| 1:f1:213:ILE:HD12 | 1:f1:250:LEU:HD21 | 1.93 | 0.50 |
| 1:X2:343:LEU:HD22 | 1:X1:402:PHE:HE2 | 1.76 | 0.50 |
| 9:uB:253:VAL:HG12 | 9:uB:257:LYS:HE3 | 1.94 | 0.50 |
| 1:B1:411:MET:HE1 | 1:B3:407:THR:HB | 1.92 | 0.50 |
| 1:d2:416:GLU:HG2 | 1:d1:171:ARG:HH12 | 1.77 | 0.50 |
| 1:f1:276:GLN:HE22 | 1:f1:521:LEU:HD11 | 1.76 | 0.50 |

*Continued on next page...*

For Manuscript Review

*Continued from previous page...*

| Atom-1 | Atom-2 | Interatomic distance (Å) | Clash overlap (Å) |
|---|---|---|---|
| 1:f3:107:VAL:HG23 | 1:f3:215:ILE:HG12 | 1.92 | 0.50 |
| 1:X3:158:PHE:CD2 | 1:X3:176:PRO:HB2 | 2.47 | 0.50 |
| 1:e1:99:PRO:O | 1:e1:103:LEU:HG | 2.11 | 0.50 |
| 1:e1:369:PHE:HB2 | 1:e1:372:PRO:HB3 | 1.93 | 0.50 |
| 9:uB:54:THR:HG21 | 10:76:71:SER:HB3 | 1.94 | 0.50 |
| 1:c2:265:MET:HA | 1:c2:270:HIS:CE1 | 2.47 | 0.50 |
| 1:f2:299:PRO:HA | 1:f2:476:ASN:HA | 1.93 | 0.50 |
| 1:f1:17:TYR:OH | 1:f3:513:ARG:NH2 | 2.32 | 0.50 |
| 1:X2:163:GLY:HA2 | 9:uB:83:PHE:H | 1.75 | 0.50 |
| 1:A2:307:VAL:HG22 | 1:A2:502:VAL:HG22 | 1.94 | 0.50 |
| 1:e1:362:VAL:HG12 | 1:e1:363:ILE:HG12 | 1.93 | 0.50 |
| 1:B3:149:GLN:NE2 | 1:B3:173:VAL:O | 2.45 | 0.50 |
| 1:e1:60:ILE:O | 1:e1:254:TYR:OH | 2.25 | 0.50 |
| 1:e1:104:ILE:HG21 | 1:e1:124:MET:HE1 | 1.93 | 0.50 |
| 1:e1:233:GLN:HA | 1:e1:321:THR:HG23 | 1.93 | 0.50 |
| 5:TQ:100:VAL:HG12 | 5:TQ:110:VAL:HG22 | 1.94 | 0.50 |
| 1:c2:62:ARG:NH2 | 1:c2:206:LEU:O | 2.42 | 0.50 |
| 1:X2:301:LYS:N | 1:X2:507:VAL:O | 2.40 | 0.50 |
| 1:e2:102:GLN:NE2 | 1:e2:222:GLU:O | 2.38 | 0.50 |
| 1:B3:216:ASP:HB2 | 9:uB:263:ILE:HG22 | 1.94 | 0.50 |
| 1:d2:136:GLN:NE2 | 1:d3:449:HIS:O | 2.40 | 0.50 |
| 1:d1:233:GLN:HA | 1:d1:321:THR:HG23 | 1.92 | 0.50 |
| 1:f2:83:LEU:HD23 | 1:f2:240:ALA:HA | 1.93 | 0.50 |
| 1:f2:510:ASN:OD1 | 1:f2:511:VAL:N | 2.39 | 0.50 |
| 1:X1:90:SER:HB3 | 1:X1:156:PRO:HG3 | 1.93 | 0.50 |
| 1:X1:439:TYR:OH | 1:X1:444:GLN:NE2 | 2.44 | 0.50 |
| 1:X3:421:VAL:HG22 | 1:X3:488:VAL:HB | 1.94 | 0.50 |
| 1:A2:201:LEU:HD12 | 1:A2:202:PRO:HD2 | 1.93 | 0.50 |
| 1:A1:153:VAL:HG21 | 1:A1:329:LEU:HD13 | 1.93 | 0.50 |
| 1:A1:422:ALA:HB3 | 1:A1:440:PHE:CE2 | 2.47 | 0.50 |
| 1:e2:134:LYS:HB3 | 1:e3:134:LYS:HG3 | 1.93 | 0.50 |
| 1:e2:172:GLN:NE2 | 1:e2:367:GLY:O | 2.45 | 0.50 |
| 1:e1:372:PRO:HG2 | 1:e1:373:PHE:CD2 | 2.47 | 0.50 |
| 1:e1:479:ARG:HH21 | 5:TP:206:LEU:HD12 | 1.77 | 0.50 |
| 5:TP:206:LEU:HD21 | 5:TP:232:MET:HE1 | 1.93 | 0.50 |
| 8:gc:138:GLU:OE2 | 8:gc:141:ARG:NH2 | 2.45 | 0.50 |
| 1:c1:17:TYR:HE1 | 1:c1:20:GLY:HA3 | 1.76 | 0.50 |
| 1:c1:299:PRO:HA | 1:c1:476:ASN:HA | 1.94 | 0.50 |
| 1:d3:138:ARG:HG2 | 1:d3:142:LYS:HE2 | 1.94 | 0.50 |
| 1:X1:302:GLU:OE1 | 1:X1:449:HIS:NE2 | 2.42 | 0.50 |
| 1:e3:343:LEU:HD21 | 1:e3:408:ALA:HB1 | 1.93 | 0.50 |

*Continued on next page...*

For Manuscript Review

*Continued from previous page...*

| Atom-1 | Atom-2 | Interatomic distance (Å) | Clash overlap (Å) |
|---|---|---|---|
| 8:gc:275:LEU:HD12 | 8:gc:302:TYR:HB3 | 1.93 | 0.50 |
| 1:B2:301:LYS:NZ | 1:B2:468:GLU:O | 2.45 | 0.50 |
| 1:c2:55:ARG:NH2 | 1:c2:108:GLU:OE2 | 2.43 | 0.50 |
| 1:d2:23:GLN:OE1 | 1:d3:32:ARG:NH2 | 2.44 | 0.50 |
| 1:d1:101:GLU:OE2 | 1:d1:140:TYR:OH | 2.19 | 0.50 |
| 1:A1:369:PHE:HB2 | 1:A1:372:PRO:HB3 | 1.94 | 0.50 |
| 8:gc:255:ASP:HB3 | 8:gc:258:PHE:HD2 | 1.77 | 0.50 |
| 1:c2:417:ASN:ND2 | 1:c2:458:ASN:OD1 | 2.36 | 0.49 |
| 1:c1:106:GLN:HE21 | 9:uB:94:LYS:HD3 | 1.76 | 0.49 |
| 1:c3:53:GLY:HA2 | 1:c3:217:ILE:O | 2.12 | 0.49 |
| 1:c3:99:PRO:O | 1:c3:103:LEU:HG | 2.11 | 0.49 |
| 1:d2:201:LEU:HD12 | 1:d2:202:PRO:HD2 | 1.92 | 0.49 |
| 1:d2:349:ALA:HA | 1:d1:373:PHE:HA | 1.94 | 0.49 |
| 1:d2:420:VAL:HG12 | 1:d2:421:VAL:HG23 | 1.93 | 0.49 |
| 1:d3:410:ASP:OD1 | 1:d3:410:ASP:N | 2.43 | 0.49 |
| 1:A3:90:SER:HB3 | 1:A3:156:PRO:HG3 | 1.94 | 0.49 |
| 1:e1:278:GLN:NE2 | 1:e1:297:ASN:O | 2.35 | 0.49 |
| 1:d2:343:LEU:HD22 | 1:d1:402:PHE:HE2 | 1.77 | 0.49 |
| 1:d2:452:ASN:O | 1:d1:138:ARG:NH2 | 2.44 | 0.49 |
| 1:X1:258:ASP:OD1 | 1:X1:259:THR:N | 2.43 | 0.49 |
| 1:e3:172:GLN:NE2 | 1:e3:367:GLY:O | 2.46 | 0.49 |
| 1:B2:199:LEU:HD11 | 1:B2:469:HIS:CD2 | 2.47 | 0.49 |
| 1:B1:419:VAL:HG11 | 1:B1:422:ALA:HB2 | 1.93 | 0.49 |
| 1:B3:42:ILE:HG21 | 1:B3:63:ASN:HD22 | 1.77 | 0.49 |
| 1:c3:95:TRP:CD1 | 1:c3:180:LEU:HD13 | 2.47 | 0.49 |
| 1:f2:67:ALA:HB3 | 1:f2:201:LEU:HB3 | 1.93 | 0.49 |
| 1:f2:301:LYS:HD3 | 1:f2:509:TYR:CZ | 2.48 | 0.49 |
| 1:X1:99:PRO:O | 1:X1:103:LEU:HG | 2.12 | 0.49 |
| 1:d2:46:PHE:HB3 | 1:d2:56:VAL:HG22 | 1.95 | 0.49 |
| 1:X2:155:ASP:HB2 | 1:X2:348:HIS:HE2 | 1.78 | 0.49 |
| 1:B2:422:ALA:HB3 | 1:B2:440:PHE:CE2 | 2.48 | 0.49 |
| 1:X2:402:PHE:HE2 | 1:X3:343:LEU:HD22 | 1.78 | 0.49 |
| 1:A2:107:VAL:HG22 | 1:A2:215:ILE:HG23 | 1.93 | 0.49 |
| 1:X3:60:ILE:O | 1:X3:254:TYR:OH | 2.31 | 0.49 |
| 1:A1:90:SER:HB3 | 1:A1:156:PRO:HG3 | 1.94 | 0.49 |
| 1:e3:430:ASP:N | 1:e3:430:ASP:OD1 | 2.44 | 0.49 |
| 1:B1:204:ILE:HG12 | 1:B1:266:ALA:HB2 | 1.95 | 0.49 |
| 1:c3:147:THR:O | 1:c3:151:THR:HG22 | 2.13 | 0.49 |
| 1:f3:278:GLN:NE2 | 1:f3:297:ASN:O | 2.44 | 0.49 |
| 1:A2:73:GLN:OE1 | 1:A3:435:ARG:NH2 | 2.40 | 0.49 |
| 1:A1:53:GLY:CA | 1:A1:217:ILE:O | 2.61 | 0.49 |

*Continued on next page...*

For Manuscript Review

*Continued from previous page...*

| Atom-1 | Atom-2 | Interatomic distance (Å) | Clash overlap (Å) |
|---|---|---|---|
| 1:e1:95:TRP:CD1 | 1:e1:180:LEU:HD13 | 2.48 | 0.49 |
| 1:e3:64:GLY:HA3 | 1:e3:254:TYR:HB3 | 1.93 | 0.49 |
| 1:A2:45:THR:HG22 | 1:e1:291:LYS:HD2 | 1.93 | 0.49 |
| 1:e2:99:PRO:O | 1:e2:103:LEU:HG | 2.12 | 0.49 |
| 1:B2:2:ALA:HB3 | 1:B2:6:MET:HB2 | 1.95 | 0.49 |
| 1:B1:375:ASN:ND2 | 1:B3:393:GLU:OE2 | 2.34 | 0.49 |
| 1:B1:384:HIS:HB2 | 1:B1:387:PHE:HD2 | 1.78 | 0.49 |
| 1:B3:77:PRO:HD3 | 1:B3:245:LEU:HD23 | 1.94 | 0.49 |
| 1:A1:301:LYS:N | 1:A1:507:VAL:O | 2.37 | 0.49 |
| 1:e3:325:LEU:HD13 | 1:e3:331:ALA:HA | 1.95 | 0.49 |
| 1:c3:303:LEU:HD13 | 1:c3:462:PHE:HE2 | 1.78 | 0.48 |
| 1:f1:99:PRO:O | 1:f1:103:LEU:HG | 2.13 | 0.48 |
| 1:f3:122:ASP:OD2 | 1:f3:505:TYR:OH | 2.29 | 0.48 |
| 1:A1:101:GLU:OE2 | 1:A1:140:TYR:OH | 2.24 | 0.48 |
| 1:A1:306:VAL:HG12 | 1:A1:457:ILE:HA | 1.95 | 0.48 |
| 1:e2:372:PRO:HG2 | 1:e2:373:PHE:CD2 | 2.48 | 0.48 |
| 1:e1:94:ARG:HD2 | 1:e1:151:THR:O | 2.12 | 0.48 |
| 4:57:266:ASP:OD1 | 4:57:267:GLY:N | 2.42 | 0.48 |
| 1:B2:134:LYS:HG2 | 1:B1:135:GLU:HG3 | 1.94 | 0.48 |
| 1:d1:153:VAL:HG21 | 1:d1:329:LEU:HD13 | 1.94 | 0.48 |
| 1:X1:307:VAL:HG22 | 1:X1:502:VAL:HG22 | 1.95 | 0.48 |
| 1:A3:199:LEU:HD11 | 1:A3:469:HIS:CD2 | 2.48 | 0.48 |
| 1:B1:364:ASN:HD21 | 1:B1:368:PHE:HB2 | 1.78 | 0.48 |
| 1:B1:369:PHE:HB2 | 1:B1:372:PRO:HB3 | 1.95 | 0.48 |
| 1:B3:107:VAL:HG22 | 1:B3:215:ILE:HG23 | 1.94 | 0.48 |
| 1:c1:95:TRP:CD1 | 1:c1:180:LEU:HD13 | 2.48 | 0.48 |
| 1:d3:304:ILE:HG12 | 1:d3:459:VAL:HG22 | 1.95 | 0.48 |
| 1:f1:60:ILE:O | 1:f1:254:TYR:OH | 2.30 | 0.48 |
| 1:X1:298:HIS:NE2 | 1:X1:523:TYR:O | 2.47 | 0.48 |
| 1:B3:280:THR:HG21 | 1:B3:295:ASN:HB2 | 1.95 | 0.48 |
| 1:c2:469:HIS:O | 1:c2:509:TYR:OH | 2.23 | 0.48 |
| 1:c3:203:LEU:O | 1:e2:208:TYR:OH | 2.29 | 0.48 |
| 1:f1:159:ALA:HA | 1:f1:359:THR:HG22 | 1.96 | 0.48 |
| 1:A1:282:ASP:HB3 | 1:A1:501:LYS:HG3 | 1.94 | 0.48 |
| 1:e2:293:LYS:NZ | 1:e2:295:ASN:OD1 | 2.46 | 0.48 |
| 1:e3:59:THR:OG1 | 1:e3:212:LYS:NZ | 2.47 | 0.48 |
| 5:TP:200:VAL:HG22 | 5:TP:221:VAL:HG12 | 1.93 | 0.48 |
| 9:uB:170:LYS:O | 9:uB:173:ILE:HG12 | 2.13 | 0.48 |
| 1:c1:332:GLN:OE1 | 1:c1:335:ASN:ND2 | 2.45 | 0.48 |
| 1:d2:403:VAL:HA | 1:d3:411:MET:HE1 | 1.95 | 0.48 |
| 1:f1:417:ASN:ND2 | 1:f1:458:ASN:OD1 | 2.31 | 0.48 |

*Continued on next page...*

For Manuscript Review

*Continued from previous page...*

| Atom-1 | Atom-2 | Interatomic distance (Å) | Clash overlap (Å) |
|---|---|---|---|
| 1:e2:65:ASP:OD2 | 1:e2:262:ARG:NH1 | 2.47 | 0.48 |
| 1:c3:282:ASP:HB3 | 1:c3:503:ARG:HG2 | 1.95 | 0.48 |
| 1:d2:165:CYS:SG | 1:d2:166:SER:N | 2.86 | 0.48 |
| 1:d3:422:ALA:HB3 | 1:d3:440:PHE:CE2 | 2.48 | 0.48 |
| 1:f2:422:ALA:HB3 | 1:f2:440:PHE:CE2 | 2.49 | 0.48 |
| 1:X2:481:ASP:OD1 | 1:X2:481:ASP:N | 2.42 | 0.48 |
| 1:X2:518:MET:HE1 | 4:57:239:TYR:H | 1.77 | 0.48 |
| 1:A2:95:TRP:CD1 | 1:A2:180:LEU:HD13 | 2.49 | 0.48 |
| 7:he:80:ILE:HD13 | 7:he:126:ILE:HG21 | 1.95 | 0.48 |
| 1:B2:312:ASN:OD1 | 1:B2:335:ASN:ND2 | 2.46 | 0.48 |
| 1:B1:43:GLU:HG3 | 1:B1:249:SER:OG | 2.13 | 0.48 |
| 1:A2:222:GLU:HG3 | 3:wD:95:THR:HG22 | 1.95 | 0.48 |
| 1:A1:116:ILE:HG23 | 1:A1:521:LEU:HD21 | 1.94 | 0.48 |
| 1:B3:308:GLN:NE2 | 1:B3:333:PRO:O | 2.47 | 0.48 |
| 8:gc:136:GLN:O | 8:gc:140:ILE:HG12 | 2.13 | 0.48 |
| 9:uB:163:THR:HB | 9:uB:166:GLN:HG3 | 1.95 | 0.48 |
| 1:B1:216:ASP:N | 1:B1:216:ASP:OD1 | 2.46 | 0.48 |
| 1:B3:76:LEU:HD11 | 1:B3:185:LEU:HD11 | 1.95 | 0.48 |
| 1:c3:278:GLN:NE2 | 1:c3:297:ASN:O | 2.40 | 0.48 |
| 1:d2:104:ILE:HG23 | 1:d2:107:VAL:HG23 | 1.95 | 0.48 |
| 1:d2:107:VAL:HG22 | 1:d2:215:ILE:HG12 | 1.96 | 0.48 |
| 1:d3:283:GLU:HG2 | 1:d3:292:ILE:HG13 | 1.96 | 0.48 |
| 1:f2:265:MET:O | 1:f2:270:HIS:NE2 | 2.47 | 0.48 |
| 1:f1:83:LEU:HD13 | 1:f1:240:ALA:HA | 1.94 | 0.48 |
| 1:f1:171:ARG:HA | 1:f1:171:ARG:HH11 | 1.79 | 0.48 |
| 1:f1:196:ASN:HD22 | 1:f1:199:LEU:HG | 1.79 | 0.48 |
| 1:A2:158:PHE:CD2 | 1:A2:176:PRO:HB2 | 2.49 | 0.48 |
| 1:A3:122:ASP:OD2 | 1:A3:505:TYR:OH | 2.31 | 0.48 |
| 1:B3:52:PHE:HE1 | 1:B3:103:LEU:HD22 | 1.79 | 0.48 |
| 1:c1:158:PHE:CD2 | 1:c1:176:PRO:HB2 | 2.49 | 0.48 |
| 1:d1:72:LEU:HA | 1:d1:250:LEU:HD23 | 1.96 | 0.48 |
| 1:d1:421:VAL:HB | 1:d1:488:VAL:HB | 1.96 | 0.48 |
| 1:d3:76:LEU:O | 1:d3:182:GLU:HA | 2.14 | 0.48 |
| 1:f1:269:PRO:HB3 | 1:f1:515:MET:HG2 | 1.96 | 0.48 |
| 1:f3:420:VAL:HG12 | 1:f3:421:VAL:HG13 | 1.95 | 0.48 |
| 1:X3:275:GLU:OE2 | 1:X3:301:LYS:NZ | 2.43 | 0.48 |
| 1:A1:138:ARG:HG2 | 1:A1:142:LYS:HE2 | 1.96 | 0.48 |
| 1:e3:158:PHE:CD2 | 1:e3:176:PRO:HB2 | 2.49 | 0.48 |
| 1:e3:369:PHE:HB2 | 1:e3:372:PRO:HB3 | 1.96 | 0.48 |
| 1:c1:122:ASP:OD2 | 1:c1:505:TYR:OH | 2.19 | 0.47 |
| 1:c3:172:GLN:O | 1:c3:177:ARG:NH2 | 2.47 | 0.47 |

*Continued on next page...*

For Manuscript Review

*Continued from previous page...*

| Atom-1 | Atom-2 | Interatomic distance (Å) | Clash overlap (Å) |
|---|---|---|---|
| 1:c3:513:ARG:HB2 | 1:c3:522:ALA:HB2 | 1.95 | 0.47 |
| 1:f3:168:ASP:OD1 | 1:f3:168:ASP:N | 2.47 | 0.47 |
| 1:B2:29:VAL:HB | 1:B3:198:GLY:HA3 | 1.96 | 0.47 |
| 1:c3:236:LYS:HD2 | 1:c3:319:LEU:HD13 | 1.95 | 0.47 |
| 1:d1:355:GLN:HE22 | 1:d1:371:ASP:HB3 | 1.78 | 0.47 |
| 1:d3:68:TYR:O | 1:d3:194:CYS:HA | 2.13 | 0.47 |
| 1:X3:53:GLY:N | 1:X3:217:ILE:O | 2.47 | 0.47 |
| 1:A3:138:ARG:HG2 | 1:A3:142:LYS:HE2 | 1.95 | 0.47 |
| 8:gc:208:LYS:HG2 | 8:gc:209:ASN:HD22 | 1.79 | 0.47 |
| 1:d1:99:PRO:O | 1:d1:103:LEU:HG | 2.14 | 0.47 |
| 1:d3:469:HIS:O | 1:d3:509:TYR:OH | 2.30 | 0.47 |
| 1:f2:107:VAL:HG22 | 1:f2:191:PHE:HZ | 1.79 | 0.47 |
| 1:A2:33:ARG:CZ | 1:A1:23:GLN:HE21 | 2.27 | 0.47 |
| 1:A3:464:LEU:O | 1:A3:465:ARG:HD2 | 2.14 | 0.47 |
| 1:B2:206:LEU:HD21 | 1:B2:514:ILE:HD12 | 1.95 | 0.47 |
| 1:c2:403:VAL:HG21 | 1:c3:408:ALA:HB2 | 1.96 | 0.47 |
| 8:gc:55:GLN:HG3 | 8:gc:57:SER:H | 1.80 | 0.47 |
| 8:gc:283:ASN:ND2 | 8:gc:285:ASN:OD1 | 2.35 | 0.47 |
| 1:c1:219:ASN:ND2 | 9:uB:32:GLU:OE2 | 2.48 | 0.47 |
| 1:c1:298:HIS:CE1 | 1:c1:525:ASN:H | 2.33 | 0.47 |
| 1:f1:134:LYS:HG3 | 1:f3:134:LYS:HB3 | 1.96 | 0.47 |
| 1:X1:316:CYS:O | 1:X1:320:GLN:HG2 | 2.15 | 0.47 |
| 1:e2:373:PHE:HA | 1:e3:349:ALA:HA | 1.97 | 0.47 |
| 6:ze:113:ASN:H | 7:he:125:ASN:ND2 | 2.12 | 0.47 |
| 8:gc:129:PHE:HD2 | 8:gc:137:LYS:HG3 | 1.80 | 0.47 |
| 8:gc:180:MET:HG3 | 8:gc:206:CYS:HB3 | 1.97 | 0.47 |
| 1:B2:309:PRO:HB2 | 1:B2:312:ASN:HD22 | 1.79 | 0.47 |
| 1:B3:9:VAL:HG11 | 5:TP:257:LEU:HD23 | 1.97 | 0.47 |
| 1:c2:7:GLN:OE1 | 1:c1:431:ARG:NH2 | 2.42 | 0.47 |
| 1:c1:16:VAL:HG13 | 1:c1:18:LEU:HG | 1.97 | 0.47 |
| 1:c3:325:LEU:HD12 | 1:c3:331:ALA:HA | 1.97 | 0.47 |
| 1:X1:116:ILE:O | 1:X1:276:GLN:NE2 | 2.32 | 0.47 |
| 1:X3:52:PHE:HE1 | 1:X3:103:LEU:HD21 | 1.80 | 0.47 |
| 1:X3:219:ASN:ND2 | 9:uB:6:GLY:O | 2.48 | 0.47 |
| 1:e2:313:VAL:HG12 | 1:e2:333:PRO:HA | 1.95 | 0.47 |
| 1:e3:213:ILE:HD12 | 1:e3:250:LEU:HD21 | 1.96 | 0.47 |
| 1:e3:436:GLU:CD | 1:e3:436:GLU:H | 2.23 | 0.47 |
| 9:uB:170:LYS:HE3 | 9:uB:173:ILE:HG13 | 1.97 | 0.47 |
| 1:B3:420:VAL:HG12 | 1:B3:421:VAL:HG23 | 1.97 | 0.47 |
| 1:d2:148:THR:O | 1:d2:152:TYR:HB2 | 2.15 | 0.47 |
| 1:d2:204:ILE:CD1 | 1:d2:262:ARG:HG3 | 2.45 | 0.47 |

*Continued on next page...*

For Manuscript Review

*Continued from previous page...*

| Atom-1 | Atom-2 | Interatomic distance (Å) | Clash overlap (Å) |
|---|---|---|---|
| 1:d1:275:GLU:OE2 | 1:d1:301:LYS:NZ | 2.40 | 0.47 |
| 1:d1:422:ALA:HB3 | 1:d1:440:PHE:CE2 | 2.50 | 0.47 |
| 1:d3:479:ARG:O | 5:TP:186:SER:OG | 2.28 | 0.47 |
| 1:f2:40:GLU:OE1 | 1:f1:4:GLY:N | 2.46 | 0.47 |
| 1:f2:136:GLN:NE2 | 1:f3:445:PRO:O | 2.47 | 0.47 |
| 1:f3:277:LEU:HD11 | 1:f3:505:TYR:HB3 | 1.96 | 0.47 |
| 1:X2:160:ALA:HA | 1:X2:178:LYS:HG2 | 1.95 | 0.47 |
| 1:X2:302:GLU:OE1 | 1:X2:449:HIS:NE2 | 2.43 | 0.47 |
| 1:e2:394:SER:HB2 | 1:e3:396:VAL:HB | 1.97 | 0.47 |
| 4:57:171:ILE:HD13 | 4:57:238:PHE:HB3 | 1.97 | 0.47 |
| 8:gc:114:ASP:OD1 | 8:gc:114:ASP:N | 2.47 | 0.47 |
| 1:c2:171:ARG:HD2 | 1:c3:492:ALA:HB2 | 1.97 | 0.47 |
| 1:c3:201:LEU:HD13 | 1:c3:211:VAL:HG21 | 1.97 | 0.47 |
| 1:c3:369:PHE:HB2 | 1:c3:372:PRO:HB3 | 1.96 | 0.47 |
| 1:d2:282:ASP:OD1 | 1:d2:282:ASP:N | 2.44 | 0.47 |
| 1:A2:408:ALA:HB2 | 1:A1:403:VAL:HG21 | 1.97 | 0.47 |
| 1:A1:46:PHE:HB3 | 1:A1:56:VAL:HG21 | 1.97 | 0.47 |
| 1:e2:286:GLY:HA3 | 9:uB:120:LEU:HD22 | 1.96 | 0.47 |
| 3:wD:28:ASN:OD1 | 3:wD:28:ASN:N | 2.47 | 0.47 |
| 1:c2:7:GLN:HG3 | 1:c1:475:CYS:HA | 1.97 | 0.47 |
| 1:c2:381:GLY:O | 1:c2:389:SER:OG | 2.25 | 0.47 |
| 1:c1:51:ASP:HB3 | 1:c1:54:ARG:HG3 | 1.96 | 0.47 |
| 1:d1:298:HIS:NE2 | 1:d1:525:ASN:OD1 | 2.47 | 0.47 |
| 1:A1:171:ARG:HA | 1:A1:171:ARG:HH11 | 1.79 | 0.47 |
| 1:e1:53:GLY:N | 1:e1:217:ILE:O | 2.48 | 0.47 |
| 1:c2:136:GLN:NE2 | 1:c3:449:HIS:O | 2.48 | 0.47 |
| 1:d3:484:THR:HG21 | 1:X3:45:THR:HG21 | 1.97 | 0.47 |
| 1:f1:310:ASP:OD2 | 1:f1:499:THR:OG1 | 2.31 | 0.47 |
| 1:f2:257:LEU:HB2 | 1:f2:262:ARG:HG3 | 1.97 | 0.46 |
| 1:f2:364:ASN:ND2 | 1:f2:368:PHE:O | 2.48 | 0.46 |
| 1:f1:147:THR:O | 1:f1:151:THR:OG1 | 2.29 | 0.46 |
| 1:X1:193:PHE:HA | 1:X1:199:LEU:O | 2.15 | 0.46 |
| 1:X3:369:PHE:HB2 | 1:X3:372:PRO:HB3 | 1.97 | 0.46 |
| 1:X3:422:ALA:HB3 | 1:X3:440:PHE:CE2 | 2.50 | 0.46 |
| 1:A2:44:GLN:NE2 | 1:A2:61:SER:H | 2.13 | 0.46 |
| 1:c2:513:ARG:HH22 | 5:TQ:103:MET:HE1 | 1.80 | 0.46 |
| 1:c1:11:TYR:O | 1:c1:15:ASP:HB2 | 2.15 | 0.46 |
| 1:d3:299:PRO:HA | 1:d3:476:ASN:HA | 1.97 | 0.46 |
| 1:f1:15:ASP:OD1 | 1:f3:476:ASN:ND2 | 2.37 | 0.46 |
| 1:X1:158:PHE:CD2 | 1:X1:176:PRO:HB2 | 2.51 | 0.46 |
| 1:X3:423:LYS:HG2 | 1:X3:486:GLN:HB3 | 1.96 | 0.46 |

*Continued on next page...*

For Manuscript Review

*Continued from previous page...*

| Atom-1 | Atom-2 | Interatomic distance (Å) | Clash overlap (Å) |
|---|---|---|---|
| 10:76:106:GLN:HG3 | 10:76:203:ILE:HG22 | 1.98 | 0.46 |
| 1:B1:99:PRO:O | 1:B1:103:LEU:HG | 2.16 | 0.46 |
| 1:c1:419:VAL:HG11 | 1:c1:422:ALA:HB2 | 1.98 | 0.46 |
| 1:d3:106:GLN:OE1 | 1:d3:118:ARG:NH2 | 2.36 | 0.46 |
| 1:f1:66:LEU:HB2 | 1:f1:255:ILE:HB | 1.96 | 0.46 |
| 1:e1:476:ASN:OD1 | 1:e1:478:SER:OG | 2.28 | 0.46 |
| 1:B2:213:ILE:HD12 | 1:B2:250:LEU:HD21 | 1.96 | 0.46 |
| 1:B1:68:TYR:O | 1:B1:194:CYS:HA | 2.15 | 0.46 |
| 1:B3:299:PRO:HA | 1:B3:476:ASN:HA | 1.96 | 0.46 |
| 1:X2:275:GLU:OE2 | 1:X2:301:LYS:NZ | 2.40 | 0.46 |
| 1:A3:291:LYS:HG3 | 1:A3:484:THR:HG23 | 1.98 | 0.46 |
| 1:B2:301:LYS:HD3 | 1:B2:509:TYR:CE2 | 2.51 | 0.46 |
| 1:B1:372:PRO:HG2 | 1:B1:373:PHE:CD2 | 2.50 | 0.46 |
| 1:c2:162:ASP:OD1 | 1:c2:171:ARG:NH1 | 2.40 | 0.46 |
| 1:c2:308:GLN:NE2 | 1:c2:333:PRO:O | 2.37 | 0.46 |
| 1:c2:438:THR:O | 1:c2:442:LEU:HB2 | 2.15 | 0.46 |
| 1:c1:17:TYR:CE1 | 1:c1:20:GLY:HA3 | 2.50 | 0.46 |
| 1:c1:246:ALA:O | 9:uB:63:LYS:NZ | 2.41 | 0.46 |
| 1:c3:49:GLN:HE22 | 9:uB:145:LEU:H | 1.64 | 0.46 |
| 1:d1:81:GLN:NE2 | 1:d1:178:LYS:O | 2.43 | 0.46 |
| 1:d3:58:CYS:HB3 | 1:d3:250:LEU:HD12 | 1.97 | 0.46 |
| 1:f3:227:MET:HE2 | 1:f3:326:ASN:ND2 | 2.31 | 0.46 |
| 1:A3:158:PHE:CD2 | 1:A3:176:PRO:HB2 | 2.50 | 0.46 |
| 1:e2:355:GLN:HE21 | 1:e2:371:ASP:HB3 | 1.81 | 0.46 |
| 1:e1:222:GLU:HG2 | 1:e1:315:TYR:CD2 | 2.51 | 0.46 |
| 8:gc:282:LEU:HD11 | 8:gc:289:LEU:HD13 | 1.96 | 0.46 |
| 1:B2:4:GLY:N | 1:B3:40:GLU:OE1 | 2.47 | 0.46 |
| 1:B3:115:ARG:HD2 | 1:B3:118:ARG:HB2 | 1.97 | 0.46 |
| 1:c3:158:PHE:CD2 | 1:c3:176:PRO:HB2 | 2.51 | 0.46 |
| 1:d3:170:VAL:HG12 | 1:d3:172:GLN:H | 1.80 | 0.46 |
| 1:f2:95:TRP:CD1 | 1:f2:180:LEU:HD13 | 2.51 | 0.46 |
| 1:X1:199:LEU:HD11 | 1:X1:469:HIS:CD2 | 2.50 | 0.46 |
| 1:X3:423:LYS:HD2 | 9:uB:92:LEU:HD23 | 1.97 | 0.46 |
| 1:A3:115:ARG:HD2 | 1:A3:118:ARG:HB2 | 1.98 | 0.46 |
| 1:e3:298:HIS:CE1 | 1:e3:525:ASN:H | 2.34 | 0.46 |
| 1:B2:166:SER:H | 1:B3:491:ASN:ND2 | 2.14 | 0.46 |
| 1:B1:202:PRO:HB2 | 1:B1:205:ALA:HB3 | 1.98 | 0.46 |
| 1:B3:70:THR:HG23 | 1:B3:252:VAL:HG22 | 1.96 | 0.46 |
| 1:c2:303:LEU:HG | 1:c2:462:PHE:HE2 | 1.81 | 0.46 |
| 1:c2:369:PHE:HB2 | 1:c2:372:PRO:HB3 | 1.98 | 0.46 |
| 1:c1:313:VAL:HG12 | 1:c1:333:PRO:HA | 1.97 | 0.46 |

*Continued on next page...*

For Manuscript Review

*Continued from previous page...*

| Atom-1 | Atom-2 | Interatomic distance (Å) | Clash overlap (Å) |
|---|---|---|---|
| 1:c3:271:GLU:OE2 | 1:c3:513:ARG:NH2 | 2.46 | 0.46 |
| 1:d2:39:MET:O | 1:d3:431:ARG:NH2 | 2.39 | 0.46 |
| 1:d2:262:ARG:NE | 5:TP:177:ASP:OD1 | 2.40 | 0.46 |
| 1:f1:133:SER:HA | 1:f3:135:GLU:HG2 | 1.98 | 0.46 |
| 1:f3:372:PRO:HG2 | 1:f3:373:PHE:CD1 | 2.51 | 0.46 |
| 1:X3:189:LEU:HD11 | 1:X3:250:LEU:HD11 | 1.98 | 0.46 |
| 1:A2:67:ALA:HB3 | 1:A2:201:LEU:HB3 | 1.98 | 0.46 |
| 1:A2:371:ASP:O | 1:A2:397:SER:HB2 | 2.16 | 0.46 |
| 1:A3:469:HIS:O | 1:A3:509:TYR:OH | 2.27 | 0.46 |
| 1:e1:265:MET:O | 1:e1:270:HIS:NE2 | 2.39 | 0.46 |
| 1:d1:305:TRP:CE2 | 1:d1:458:ASN:HB2 | 2.51 | 0.46 |
| 1:d3:293:LYS:HD3 | 1:e1:47:ASN:HB3 | 1.98 | 0.46 |
| 1:d3:320:GLN:O | 1:d3:326:ASN:ND2 | 2.49 | 0.46 |
| 1:f2:51:ASP:HB3 | 1:f2:54:ARG:HD2 | 1.96 | 0.46 |
| 1:A3:479:ARG:NH1 | 5:TP:252:LYS:O | 2.48 | 0.46 |
| 1:B2:384:HIS:HB2 | 1:B2:387:PHE:CD2 | 2.47 | 0.46 |
| 1:B3:425:GLN:NE2 | 1:B3:430:ASP:OD1 | 2.49 | 0.46 |
| 1:c2:447:GLN:HE21 | 1:c2:448:HIS:CE1 | 2.33 | 0.46 |
| 1:d2:307:VAL:HG22 | 1:d2:502:VAL:HG22 | 1.97 | 0.46 |
| 1:d2:309:PRO:HB2 | 1:d2:312:ASN:HD22 | 1.80 | 0.46 |
| 1:f2:2:ALA:HB1 | 6:ze:134:LEU:HD11 | 1.98 | 0.46 |
| 1:X2:423:LYS:HD2 | 1:X2:425:GLN:HE21 | 1.80 | 0.46 |
| 1:X3:225:TRP:CZ2 | 1:X3:227:MET:HG3 | 2.51 | 0.46 |
| 1:A1:451:ARG:HA | 1:A3:135:GLU:HB3 | 1.98 | 0.46 |
| 1:e1:10:ALA:HB1 | 1:e3:479:ARG:HB2 | 1.98 | 0.46 |
| 1:e1:384:HIS:HB2 | 1:e1:387:PHE:HD2 | 1.81 | 0.46 |
| 1:B1:158:PHE:CD2 | 1:B1:176:PRO:HB2 | 2.51 | 0.46 |
| 1:B1:300:CYS:SG | 1:B1:506:ALA:HB1 | 2.56 | 0.46 |
| 1:B3:176:PRO:HB3 | 1:B3:347:VAL:HG21 | 1.98 | 0.46 |
| 1:c1:202:PRO:HB2 | 1:c1:205:ALA:HB3 | 1.98 | 0.46 |
| 1:c1:278:GLN:HE22 | 1:c1:297:ASN:H | 1.64 | 0.46 |
| 1:f1:348:HIS:HB2 | 1:f3:373:PHE:HD2 | 1.80 | 0.46 |
| 1:f3:93:ALA:HA | 1:f3:225:TRP:O | 2.15 | 0.46 |
| 1:f3:170:VAL:O | 1:f3:177:ARG:NH2 | 2.43 | 0.46 |
| 1:X1:104:ILE:HG23 | 1:X1:107:VAL:HG23 | 1.97 | 0.46 |
| 1:A1:424:LEU:HB3 | 1:A1:432:PHE:CE2 | 2.51 | 0.46 |
| 1:A3:50:ALA:HB3 | 1:A3:248:ALA:HB3 | 1.98 | 0.46 |
| 1:e1:106:GLN:HE22 | 3:wD:65:ASN:HA | 1.81 | 0.46 |
| 1:B2:278:GLN:HG3 | 1:B2:506:ALA:HB3 | 1.97 | 0.45 |
| 1:B1:237:ILE:HG22 | 1:B1:240:ALA:H | 1.81 | 0.45 |
| 1:B1:417:ASN:ND2 | 1:B1:441:ASP:OD2 | 2.48 | 0.45 |

*Continued on next page...*

For Manuscript Review

*Continued from previous page...*

| Atom-1 | Atom-2 | Interatomic distance (Å) | Clash overlap (Å) |
|---|---|---|---|
| 1:c1:76:LEU:HD21 | 1:c1:103:LEU:HD11 | 1.98 | 0.45 |
| 1:c1:288:SER:O | 1:c1:290:ASN:ND2 | 2.49 | 0.45 |
| 1:f3:300:CYS:HA | 1:f3:508:ASN:HA | 1.98 | 0.45 |
| 1:X1:153:VAL:HG21 | 1:X1:329:LEU:HD13 | 1.98 | 0.45 |
| 1:e2:343:LEU:HD22 | 1:e1:402:PHE:HE2 | 1.80 | 0.45 |
| 1:e1:53:GLY:CA | 1:e1:217:ILE:O | 2.64 | 0.45 |
| 1:e3:46:PHE:CD1 | 1:e3:58:CYS:HB2 | 2.51 | 0.45 |
| 1:B3:72:LEU:HG | 1:B3:74:VAL:HG23 | 1.98 | 0.45 |
| 1:c1:236:LYS:HB3 | 1:c1:236:LYS:HE2 | 1.78 | 0.45 |
| 1:c1:451:ARG:HA | 1:c3:135:GLU:HB3 | 1.98 | 0.45 |
| 1:f3:43:GLU:HG3 | 1:f3:249:SER:OG | 2.16 | 0.45 |
| 1:X1:17:TYR:OH | 1:X3:271:GLU:OE2 | 2.29 | 0.45 |
| 1:X1:301:LYS:N | 1:X1:507:VAL:O | 2.38 | 0.45 |
| 1:X1:492:ALA:HB2 | 1:X3:171:ARG:HD2 | 1.98 | 0.45 |
| 1:X3:525:ASN:ND2 | 1:e2:429:GLN:HG3 | 2.31 | 0.45 |
| 1:e3:422:ALA:HB3 | 1:e3:440:PHE:CE1 | 2.52 | 0.45 |
| 1:e3:439:TYR:OH | 1:e3:444:GLN:NE2 | 2.44 | 0.45 |
| 1:c2:110:GLU:HG2 | 1:c2:115:ARG:HA | 1.98 | 0.45 |
| 1:f2:7:GLN:OE1 | 1:f1:431:ARG:NH2 | 2.36 | 0.45 |
| 1:f1:369:PHE:HB2 | 1:f1:372:PRO:HB3 | 1.97 | 0.45 |
| 1:X1:137:GLU:OE1 | 1:X1:451:ARG:NH1 | 2.50 | 0.45 |
| 1:A2:243:GLN:HE22 | 3:wD:51:ASN:HA | 1.82 | 0.45 |
| 1:A2:402:PHE:HE2 | 1:A3:343:LEU:HD22 | 1.80 | 0.45 |
| 1:B2:287:SER:HA | 3:wD:21:GLY:HA3 | 1.97 | 0.45 |
| 1:d2:15:ASP:OD2 | 1:d1:476:ASN:ND2 | 2.50 | 0.45 |
| 1:d1:204:ILE:HG13 | 1:d1:266:ALA:HB2 | 1.99 | 0.45 |
| 1:X3:90:SER:HB3 | 1:X3:156:PRO:HG3 | 1.98 | 0.45 |
| 1:A2:301:LYS:HE2 | 1:A2:509:TYR:CE2 | 2.51 | 0.45 |
| 1:A1:44:GLN:NE2 | 1:A1:254:TYR:OH | 2.49 | 0.45 |
| 1:A1:196:ASN:HD22 | 1:A1:198:GLY:H | 1.65 | 0.45 |
| 1:e2:29:VAL:HB | 1:e3:198:GLY:HA3 | 1.98 | 0.45 |
| 1:e2:422:ALA:HB3 | 1:e2:440:PHE:CE2 | 2.51 | 0.45 |
| 7:he:62:ILE:HD12 | 7:he:92:MET:SD | 2.55 | 0.45 |
| 10:76:158:ASN:HB3 | 10:76:161:VAL:HB | 1.98 | 0.45 |
| 1:B2:447:GLN:HE21 | 1:B2:448:HIS:CE1 | 2.35 | 0.45 |
| 1:f3:98:PHE:HB2 | 1:f3:334:PHE:HE2 | 1.81 | 0.45 |
| 1:X3:276:GLN:NE2 | 1:X3:510:ASN:HD22 | 2.14 | 0.45 |
| 1:e2:7:GLN:OE1 | 1:e1:431:ARG:NE | 2.48 | 0.45 |
| 1:e2:111:ILE:HG12 | 1:e2:211:VAL:HG22 | 1.99 | 0.45 |
| 1:e2:213:ILE:HD12 | 1:e2:250:LEU:HD21 | 1.99 | 0.45 |
| 1:e3:93:ALA:HA | 1:e3:225:TRP:O | 2.16 | 0.45 |

*Continued on next page...*

For Manuscript Review

*Continued from previous page...*

| Atom-1 | Atom-2 | Interatomic distance (Å) | Clash overlap (Å) |
|---|---|---|---|
| 5:TP:240:LYS:HB3 | 5:TP:244:GLU:HB2 | 1.98 | 0.45 |
| 1:d1:155:ASP:HB2 | 1:d1:348:HIS:HE2 | 1.81 | 0.45 |
| 1:f1:422:ALA:HB3 | 1:f1:440:PHE:CE2 | 2.52 | 0.45 |
| 1:f3:216:ASP:N | 1:f3:216:ASP:OD1 | 2.48 | 0.45 |
| 1:A2:343:LEU:HD22 | 1:A1:402:PHE:HE2 | 1.81 | 0.45 |
| 1:A1:362:VAL:HG12 | 1:A1:363:ILE:HG12 | 1.99 | 0.45 |
| 1:e1:221:GLU:HG2 | 3:wD:72:PHE:CD2 | 2.52 | 0.45 |
| 1:B1:15:ASP:O | 1:B1:19:THR:N | 2.43 | 0.45 |
| 1:c2:513:ARG:HE | 1:c2:522:ALA:HB2 | 1.81 | 0.45 |
| 1:X2:153:VAL:HG21 | 1:X2:329:LEU:HD13 | 1.98 | 0.45 |
| 1:X1:451:ARG:HA | 1:X3:135:GLU:HB3 | 1.99 | 0.45 |
| 1:A3:289:SER:OG | 3:wD:36:LYS:HB2 | 2.17 | 0.45 |
| 1:e1:93:ALA:HA | 1:e1:225:TRP:O | 2.17 | 0.45 |
| 1:e1:510:ASN:HD22 | 1:e1:511:VAL:N | 2.14 | 0.45 |
| 8:gc:95:LEU:HD21 | 8:gc:310:THR:HG23 | 1.99 | 0.45 |
| 1:B1:147:THR:O | 1:B1:151:THR:OG1 | 2.31 | 0.45 |
| 1:c1:316:CYS:SG | 9:uB:30:VAL:HG11 | 2.57 | 0.45 |
| 1:d2:99:PRO:O | 1:d2:103:LEU:HG | 2.16 | 0.45 |
| 1:d2:291:LYS:HD3 | 1:d2:486:GLN:HG2 | 1.99 | 0.45 |
| 1:d1:148:THR:O | 1:d1:152:TYR:HB2 | 2.16 | 0.45 |
| 1:d1:171:ARG:HD2 | 1:d1:171:ARG:HA | 1.77 | 0.45 |
| 1:d3:53:GLY:N | 1:d3:218:ARG:O | 2.49 | 0.45 |
| 1:d3:70:THR:HG22 | 1:d3:250:LEU:HD22 | 1.99 | 0.45 |
| 1:X1:419:VAL:HG11 | 1:X1:422:ALA:HB2 | 1.98 | 0.45 |
| 1:e3:424:LEU:O | 1:e3:431:ARG:N | 2.42 | 0.45 |
| 1:c1:513:ARG:HH12 | 7:he:160:LEU:HD11 | 1.82 | 0.45 |
| 1:d2:66:LEU:HD11 | 1:d2:257:LEU:HD11 | 1.98 | 0.45 |
| 1:d3:170:VAL:HG13 | 1:d3:363:ILE:HG21 | 1.99 | 0.45 |
| 1:d3:301:LYS:HD3 | 1:d3:509:TYR:CE2 | 2.52 | 0.45 |
| 1:X2:162:ASP:OD1 | 1:X2:162:ASP:N | 2.49 | 0.45 |
| 1:X1:139:GLY:O | 1:X1:143:MET:HG3 | 2.17 | 0.45 |
| 1:X1:287:SER:O | 1:X1:290:ASN:ND2 | 2.41 | 0.45 |
| 1:A3:427:ASN:ND2 | 1:A3:481:ASP:H | 2.14 | 0.45 |
| 4:57:204:THR:O | 4:57:208:VAL:HG23 | 2.17 | 0.45 |
| 4:57:207:ILE:O | 4:57:211:ASN:ND2 | 2.45 | 0.45 |
| 1:B2:171:ARG:HD2 | 1:B3:492:ALA:HB1 | 1.98 | 0.45 |
| 1:B2:325:LEU:HD23 | 1:B2:328:LEU:HD12 | 1.98 | 0.45 |
| 1:c1:16:VAL:HG11 | 1:c3:299:PRO:HG3 | 1.99 | 0.45 |
| 1:c1:219:ASN:H | 1:c1:219:ASN:HD22 | 1.65 | 0.45 |
| 1:c3:153:VAL:HG21 | 1:c3:329:LEU:HD13 | 1.99 | 0.45 |
| 1:d2:271:GLU:OE2 | 1:d2:513:ARG:NH1 | 2.50 | 0.45 |

*Continued on next page...*

For Manuscript Review

*Continued from previous page...*

| Atom-1 | Atom-2 | Interatomic distance (Å) | Clash overlap (Å) |
|---|---|---|---|
| 1:d3:148:THR:O | 1:d3:152:TYR:HB2 | 2.17 | 0.45 |
| 1:f2:300:CYS:HA | 1:f2:508:ASN:HA | 1.99 | 0.45 |
| 1:X1:33:ARG:NE | 1:X3:20:GLY:O | 2.37 | 0.45 |
| 1:X1:95:TRP:CD1 | 1:X1:180:LEU:HD13 | 2.53 | 0.45 |
| 1:X1:175:ALA:HB2 | 1:X1:363:ILE:HD12 | 1.98 | 0.45 |
| 1:X1:442:LEU:HD23 | 1:X1:452:ASN:HD21 | 1.82 | 0.45 |
| 1:A2:384:HIS:HB2 | 1:A2:387:PHE:HD2 | 1.82 | 0.45 |
| 1:A3:372:PRO:HG2 | 1:A3:373:PHE:CD2 | 2.52 | 0.45 |
| 1:e1:479:ARG:HA | 5:TP:202:PRO:HA | 1.98 | 0.45 |
| 1:c2:372:PRO:HG2 | 1:c2:373:PHE:CD2 | 2.52 | 0.44 |
| 1:c3:50:ALA:HB1 | 1:c3:248:ALA:HB3 | 1.99 | 0.44 |
| 1:d2:98:PHE:HB2 | 1:d2:334:PHE:HE2 | 1.82 | 0.44 |
| 1:d1:372:PRO:HG2 | 1:d1:373:PHE:CD2 | 2.52 | 0.44 |
| 1:f2:148:THR:O | 1:f2:152:TYR:HB2 | 2.17 | 0.44 |
| 1:f2:481:ASP:OD2 | 8:gc:300:GLN:NE2 | 2.50 | 0.44 |
| 8:gc:101:ILE:HG12 | 8:gc:110:ILE:HG21 | 1.98 | 0.44 |
| 1:B2:84:ALA:HB2 | 1:B2:91:VAL:HG23 | 2.00 | 0.44 |
| 1:d2:291:LYS:HB2 | 1:d2:291:LYS:HE2 | 1.75 | 0.44 |
| 1:d2:354:THR:OG1 | 1:d1:376:ASP:OD2 | 2.29 | 0.44 |
| 1:d1:199:LEU:HD11 | 1:d1:469:HIS:CD2 | 2.53 | 0.44 |
| 1:d3:227:MET:HE2 | 1:d3:326:ASN:ND2 | 2.32 | 0.44 |
| 1:f2:74:VAL:HG13 | 1:f2:245:LEU:HD22 | 1.98 | 0.44 |
| 1:f2:227:MET:HE2 | 1:f2:326:ASN:OD1 | 2.17 | 0.44 |
| 1:f3:13:ALA:H | 5:TQ:155:GLU:CD | 2.23 | 0.44 |
| 1:f3:62:ARG:HA | 1:f3:203:LEU:HD13 | 2.00 | 0.44 |
| 1:X2:307:VAL:HG22 | 1:X2:502:VAL:HG22 | 2.00 | 0.44 |
| 1:X3:166:SER:HA | 1:X3:171:ARG:NH2 | 2.31 | 0.44 |
| 1:A2:104:ILE:HG23 | 1:A2:107:VAL:HG23 | 1.99 | 0.44 |
| 1:e1:119:GLN:NE2 | 1:e1:190:GLN:O | 2.37 | 0.44 |
| 1:e1:510:ASN:HD22 | 1:e1:511:VAL:H | 1.65 | 0.44 |
| 9:uB:27:LYS:HG3 | 9:uB:100:ILE:HD13 | 1.99 | 0.44 |
| 1:B1:239:ASP:HA | 1:B1:242:LYS:HE2 | 1.99 | 0.44 |
| 1:c2:198:GLY:HA3 | 1:c1:29:VAL:HG13 | 1.99 | 0.44 |
| 1:c1:300:CYS:HA | 1:c1:508:ASN:HA | 1.99 | 0.44 |
| 1:c3:53:GLY:CA | 1:c3:217:ILE:O | 2.65 | 0.44 |
| 1:f1:67:ALA:HB2 | 1:f1:203:LEU:HD11 | 1.99 | 0.44 |
| 1:X2:198:GLY:HA3 | 1:X1:29:VAL:HB | 1.98 | 0.44 |
| 1:A1:68:TYR:O | 1:A1:194:CYS:HA | 2.16 | 0.44 |
| 1:e2:205:ALA:HB2 | 1:e2:265:MET:O | 2.18 | 0.44 |
| 1:B3:107:VAL:HG13 | 1:B3:215:ILE:HG12 | 1.98 | 0.44 |
| 1:B3:122:ASP:OD1 | 1:B3:336:TYR:OH | 2.27 | 0.44 |

*Continued on next page...*

For Manuscript Review

*Continued from previous page...*

| Atom-1 | Atom-2 | Interatomic distance (Å) | Clash overlap (Å) |
|---|---|---|---|
| 1:c2:107:VAL:HG22 | 1:c2:215:ILE:HG12 | 2.00 | 0.44 |
| 1:X3:153:VAL:HG21 | 1:X3:329:LEU:HD13 | 1.99 | 0.44 |
| 1:e2:403:VAL:HA | 1:e3:411:MET:HE1 | 2.00 | 0.44 |
| 1:B3:122:ASP:OD2 | 1:B3:505:TYR:OH | 2.32 | 0.44 |
| 1:c2:2:ALA:HB3 | 1:c2:6:MET:HE2 | 1.99 | 0.44 |
| 1:c2:472:SER:HA | 1:c3:26:PHE:CE2 | 2.52 | 0.44 |
| 1:d1:299:PRO:HA | 1:d1:476:ASN:HA | 1.98 | 0.44 |
| 1:d3:70:THR:HG23 | 1:d3:252:VAL:HG22 | 1.99 | 0.44 |
| 1:f1:2:ALA:HB1 | 1:f3:429:GLN:HB3 | 1.98 | 0.44 |
| 1:f1:209:HIS:HD1 | 1:f1:519:GLY:H | 1.66 | 0.44 |
| 1:f3:304:ILE:HG12 | 1:f3:459:VAL:HG22 | 1.98 | 0.44 |
| 1:X1:205:ALA:O | 1:X1:517:GLY:HA2 | 2.17 | 0.44 |
| 1:X1:491:ASN:HD22 | 1:X3:165:CYS:HA | 1.82 | 0.44 |
| 1:A2:63:ASN:ND2 | 1:e1:428:GLY:HA3 | 2.32 | 0.44 |
| 1:e2:107:VAL:HG22 | 1:e2:215:ILE:HG12 | 1.98 | 0.44 |
| 1:e1:461:SER:OG | 1:e1:463:ALA:O | 2.35 | 0.44 |
| 1:B1:479:ARG:HA | 1:B1:479:ARG:HH11 | 1.83 | 0.44 |
| 1:c2:72:LEU:HG | 1:c2:74:VAL:HG23 | 1.99 | 0.44 |
| 1:c2:422:ALA:HB3 | 1:c2:440:PHE:CE2 | 2.52 | 0.44 |
| 1:c1:170:VAL:O | 1:c1:177:ARG:NH2 | 2.33 | 0.44 |
| 1:c1:316:CYS:SG | 9:uB:112:ASN:ND2 | 2.91 | 0.44 |
| 1:d3:158:PHE:CD2 | 1:d3:176:PRO:HB2 | 2.53 | 0.44 |
| 1:f2:99:PRO:O | 1:f2:103:LEU:HG | 2.17 | 0.44 |
| 1:f2:159:ALA:O | 1:f2:178:LYS:N | 2.45 | 0.44 |
| 1:f1:101:GLU:OE2 | 1:f1:140:TYR:OH | 2.29 | 0.44 |
| 1:f1:469:HIS:O | 1:f1:509:TYR:OH | 2.31 | 0.44 |
| 1:X2:372:PRO:HG2 | 1:X2:373:PHE:CD2 | 2.52 | 0.44 |
| 1:X3:298:HIS:CE1 | 1:X3:525:ASN:H | 2.36 | 0.44 |
| 1:A2:465:ARG:HA | 1:A2:465:ARG:HD3 | 1.83 | 0.44 |
| 1:e2:43:GLU:HB2 | 1:e2:251:PHE:CE2 | 2.53 | 0.44 |
| 1:e2:94:ARG:HD3 | 1:e2:153:VAL:HG23 | 1.99 | 0.44 |
| 6:ze:100:PRO:O | 7:he:67:ARG:NH1 | 2.49 | 0.44 |
| 9:uB:161:CYS:HB2 | 9:uB:186:THR:O | 2.17 | 0.44 |
| 1:B2:147:THR:O | 1:B2:151:THR:OG1 | 2.28 | 0.44 |
| 1:c1:265:MET:O | 1:c1:270:HIS:NE2 | 2.46 | 0.44 |
| 1:c3:225:TRP:CD1 | 1:c3:330:GLY:HA2 | 2.53 | 0.44 |
| 1:d2:426:LEU:HD13 | 1:d2:431:ARG:HB2 | 2.00 | 0.44 |
| 1:d2:431:ARG:NH2 | 1:d3:7:GLN:OE1 | 2.43 | 0.44 |
| 1:f2:306:VAL:HG12 | 1:f2:457:ILE:HA | 2.00 | 0.44 |
| 1:f2:335:ASN:OD1 | 1:f2:337:THR:OG1 | 2.27 | 0.44 |
| 1:X2:369:PHE:HB2 | 1:X2:372:PRO:HB3 | 2.00 | 0.44 |

*Continued on next page...*

For Manuscript Review

*Continued from previous page...*

| Atom-1 | Atom-2 | Interatomic distance (Å) | Clash overlap (Å) |
|---|---|---|---|
| 1:X3:434:GLU:CD | 1:X3:434:GLU:H | 2.25 | 0.44 |
| 1:A2:83:LEU:O | 1:A2:239:ASP:HB2 | 2.18 | 0.44 |
| 1:A1:63:ASN:HB2 | 1:A1:254:TYR:CE2 | 2.53 | 0.44 |
| 1:A3:70:THR:HG22 | 1:A3:194:CYS:SG | 2.58 | 0.44 |
| 1:e1:236:LYS:HE2 | 1:e1:319:LEU:HD13 | 2.00 | 0.44 |
| 8:gc:150:TYR:HB2 | 8:gc:213:MET:HG2 | 2.00 | 0.44 |
| 1:c2:198:GLY:HA2 | 1:c1:27:TRP:C | 2.43 | 0.44 |
| 1:c2:450:THR:OG1 | 1:c2:467:GLU:OE1 | 2.27 | 0.44 |
| 1:c3:95:TRP:HB3 | 1:c3:99:PRO:HG3 | 1.99 | 0.44 |
| 1:c3:282:ASP:OD1 | 1:c3:282:ASP:N | 2.50 | 0.44 |
| 1:X2:107:VAL:HA | 1:X2:214:ASN:O | 2.18 | 0.44 |
| 1:X1:75:THR:HB | 1:X1:246:ALA:HB3 | 1.98 | 0.44 |
| 1:X3:102:GLN:NE2 | 1:X3:222:GLU:O | 2.43 | 0.44 |
| 1:X3:276:GLN:HE22 | 1:X3:510:ASN:HD22 | 1.66 | 0.44 |
| 1:X3:372:PRO:HG2 | 1:X3:373:PHE:CD2 | 2.53 | 0.44 |
| 1:A2:160:ALA:HA | 1:A2:178:LYS:HG2 | 2.00 | 0.44 |
| 1:A1:372:PRO:HG2 | 1:A1:373:PHE:CD2 | 2.53 | 0.44 |
| 1:A3:273:LEU:O | 1:A3:469:HIS:NE2 | 2.45 | 0.44 |
| 1:e2:54:ARG:HD2 | 9:uB:141:ALA:HA | 2.00 | 0.44 |
| 1:e1:125:HIS:CE1 | 1:e1:129:GLN:HE21 | 2.36 | 0.44 |
| 1:e1:221:GLU:HA | 1:e1:241:TYR:CE1 | 2.52 | 0.44 |
| 1:d2:236:LYS:NZ | 1:d2:238:ASN:OD1 | 2.48 | 0.44 |
| 1:d3:464:LEU:HB2 | 1:d3:472:SER:HB2 | 1.99 | 0.44 |
| 1:f3:127:TRP:CE2 | 1:f3:131:THR:HG21 | 2.53 | 0.44 |
| 1:A1:340:TYR:O | 1:A3:172:GLN:NE2 | 2.43 | 0.44 |
| 1:A3:300:CYS:N | 1:A3:475:CYS:O | 2.45 | 0.44 |
| 1:e1:114:GLN:HE21 | 1:e1:114:GLN:HB3 | 1.54 | 0.44 |
| 1:e1:497:VAL:HG11 | 3:wD:82:ILE:HD12 | 1.98 | 0.44 |
| 3:wD:79:ASN:HD21 | 3:wD:81:ASN:HB2 | 1.83 | 0.44 |
| 1:c2:83:LEU:HD23 | 1:c2:240:ALA:HA | 2.00 | 0.43 |
| 1:c1:306:VAL:HG12 | 1:c1:457:ILE:HA | 2.00 | 0.43 |
| 1:c3:388:GLU:OE2 | 1:c3:391:THR:OG1 | 2.30 | 0.43 |
| 1:d2:160:ALA:O | 1:d2:177:ARG:NH1 | 2.49 | 0.43 |
| 1:d3:94:ARG:HD3 | 1:d3:153:VAL:HG23 | 1.99 | 0.43 |
| 1:X3:101:GLU:OE2 | 1:X3:140:TYR:OH | 2.25 | 0.43 |
| 1:e1:269:PRO:HG3 | 1:e1:515:MET:HE3 | 2.00 | 0.43 |
| 2:vC:100:TYR:O | 2:vC:104:ARG:HG2 | 2.18 | 0.43 |
| 4:57:163:THR:N | 4:57:187:LYS:O | 2.49 | 0.43 |
| 9:uB:70:ARG:HH21 | 9:uB:72:GLN:NE2 | 2.16 | 0.43 |
| 1:B2:23:GLN:HE21 | 1:B3:33:ARG:NH2 | 2.17 | 0.43 |
| 1:B2:465:ARG:HD3 | 1:B2:465:ARG:HA | 1.87 | 0.43 |

*Continued on next page...*

For Manuscript Review

Continued from previous page...

| Atom-1 | Atom-2 | Interatomic distance (Å) | Clash overlap (Å) |
|---|---|---|---|
| 1:B1:422:ALA:HB3 | 1:B1:440:PHE:CE2 | 2.53 | 0.43 |
| 1:B3:369:PHE:HB2 | 1:B3:372:PRO:HB3 | 1.99 | 0.43 |
| 1:c1:465:ARG:HA | 1:c1:465:ARG:HD3 | 1.89 | 0.43 |
| 1:d2:158:PHE:CD2 | 1:d2:176:PRO:HB2 | 2.53 | 0.43 |
| 1:d1:257:LEU:HB2 | 1:d1:262:ARG:HG3 | 1.99 | 0.43 |
| 1:d1:417:ASN:HD22 | 1:d1:441:ASP:HB2 | 1.83 | 0.43 |
| 1:d3:343:LEU:HB2 | 1:d3:413:CYS:HB2 | 1.99 | 0.43 |
| 1:X2:101:GLU:OE2 | 1:X2:140:TYR:OH | 2.28 | 0.43 |
| 1:X3:51:ASP:OD1 | 1:X3:52:PHE:N | 2.41 | 0.43 |
| 1:A1:61:SER:O | 1:A1:254:TYR:OH | 2.30 | 0.43 |
| 1:A1:302:GLU:OE1 | 1:A1:449:HIS:NE2 | 2.46 | 0.43 |
| 1:A1:384:HIS:HB2 | 1:A1:387:PHE:HD2 | 1.83 | 0.43 |
| 1:A3:99:PRO:O | 1:A3:103:LEU:HG | 2.16 | 0.43 |
| 8:gc:67:TYR:CD1 | 8:gc:141:ARG:HD2 | 2.53 | 0.43 |
| 9:uB:48:ARG:NH2 | 9:uB:51:ASP:OD1 | 2.51 | 0.43 |
| 1:B2:302:GLU:OE1 | 1:B2:449:HIS:NE2 | 2.45 | 0.43 |
| 1:c2:90:SER:HB3 | 1:c2:156:PRO:HG3 | 2.00 | 0.43 |
| 1:c1:208:TYR:CE1 | 1:X2:62:ARG:HD2 | 2.53 | 0.43 |
| 1:c3:497:VAL:HB | 10:76:145:HIS:O | 2.18 | 0.43 |
| 1:d2:106:GLN:HE21 | 1:d2:120:TYR:HE2 | 1.66 | 0.43 |
| 1:d3:95:TRP:CD1 | 1:d3:180:LEU:HD13 | 2.52 | 0.43 |
| 1:d3:242:LYS:HD3 | 3:wD:79:ASN:HD22 | 1.84 | 0.43 |
| 1:f1:343:LEU:HD21 | 1:f1:408:ALA:HB1 | 2.00 | 0.43 |
| 1:X1:43:GLU:HG3 | 1:X1:249:SER:OG | 2.18 | 0.43 |
| 1:X3:300:CYS:HA | 1:X3:508:ASN:HA | 2.00 | 0.43 |
| 1:X3:384:HIS:HB2 | 1:X3:387:PHE:CD2 | 2.52 | 0.43 |
| 1:e3:53:GLY:O | 1:e3:54:ARG:NH1 | 2.52 | 0.43 |
| 10:76:186:PRO:O | 10:76:187:THR:OG1 | 2.32 | 0.43 |
| 1:B2:512:LEU:HD12 | 1:B2:520:GLY:O | 2.18 | 0.43 |
| 1:B3:271:GLU:HG2 | 1:B3:513:ARG:HG2 | 2.00 | 0.43 |
| 1:d2:476:ASN:C | 1:d2:478:SER:H | 2.25 | 0.43 |
| 1:d1:291:LYS:HG3 | 1:d1:486:GLN:HG2 | 2.00 | 0.43 |
| 1:d3:439:TYR:OH | 1:d3:444:GLN:NE2 | 2.51 | 0.43 |
| 1:f1:425:GLN:HG2 | 1:f1:430:ASP:HA | 2.00 | 0.43 |
| 1:A1:148:THR:O | 1:A1:152:TYR:HB2 | 2.18 | 0.43 |
| 1:e1:163:GLY:H | 1:e1:166:SER:HG | 1.60 | 0.43 |
| 4:57:156:LYS:NZ | 4:57:199:GLU:OE1 | 2.52 | 0.43 |
| 8:gc:286:THR:HG22 | 8:gc:288:GLY:H | 1.82 | 0.43 |
| 1:c3:325:LEU:H | 1:c3:325:LEU:HD23 | 1.84 | 0.43 |
| 1:f2:44:GLN:NE2 | 1:f2:61:SER:OG | 2.49 | 0.43 |
| 1:f2:66:LEU:HD23 | 1:f2:202:PRO:HA | 2.01 | 0.43 |

Continued on next page...

For Manuscript Review

*Continued from previous page...*

| Atom-1 | Atom-2 | Interatomic distance (Å) | Clash overlap (Å) |
|---|---|---|---|
| 1:X1:420:VAL:HG12 | 1:X1:421:VAL:HG23 | 2.00 | 0.43 |
| 1:X3:132:LEU:O | 1:X3:451:ARG:HD2 | 2.19 | 0.43 |
| 1:X3:307:VAL:HG22 | 1:X3:502:VAL:HG22 | 2.00 | 0.43 |
| 1:A2:75:THR:HB | 1:A2:246:ALA:HB3 | 2.00 | 0.43 |
| 1:A3:282:ASP:OD1 | 1:A3:282:ASP:N | 2.52 | 0.43 |
| 1:e2:439:TYR:OH | 1:e2:444:GLN:NE2 | 2.48 | 0.43 |
| 1:e1:104:ILE:HG23 | 1:e1:107:VAL:HG23 | 2.00 | 0.43 |
| 1:e3:291:LYS:HG2 | 1:e3:486:GLN:HG2 | 1.99 | 0.43 |
| 1:B2:332:GLN:OE1 | 1:B2:335:ASN:ND2 | 2.42 | 0.43 |
| 1:c3:146:ASN:O | 1:c3:409:LEU:HD11 | 2.18 | 0.43 |
| 1:f2:372:PRO:HG2 | 1:f2:373:PHE:CD2 | 2.54 | 0.43 |
| 1:A2:369:PHE:HB2 | 1:A2:372:PRO:HB3 | 2.01 | 0.43 |
| 1:A3:325:LEU:HD23 | 1:A3:328:LEU:HD12 | 2.01 | 0.43 |
| 1:e3:5:LEU:HD23 | 1:e3:5:LEU:HA | 1.84 | 0.43 |
| 1:B3:66:LEU:HB2 | 1:B3:255:ILE:HB | 2.01 | 0.43 |
| 1:c2:172:GLN:NE2 | 1:c2:367:GLY:O | 2.51 | 0.43 |
| 1:X3:21:ASN:OD1 | 4:57:220:LYS:NZ | 2.38 | 0.43 |
| 1:A3:196:ASN:HB3 | 1:A3:199:LEU:HB2 | 2.00 | 0.43 |
| 1:A3:434:GLU:CD | 1:A3:434:GLU:H | 2.26 | 0.43 |
| 1:e2:155:ASP:HB2 | 1:e2:348:HIS:HE2 | 1.82 | 0.43 |
| 1:e3:201:LEU:HD12 | 1:e3:202:PRO:HD2 | 1.99 | 0.43 |
| 1:c3:302:GLU:OE1 | 1:c3:449:HIS:NE2 | 2.41 | 0.43 |
| 1:d3:94:ARG:HD2 | 1:d3:151:THR:O | 2.19 | 0.43 |
| 1:f2:424:LEU:HB3 | 1:f2:432:PHE:CE1 | 2.54 | 0.43 |
| 1:f1:160:ALA:HB2 | 1:f1:178:LYS:HE2 | 1.99 | 0.43 |
| 1:f3:380:THR:HG22 | 1:f3:392:VAL:HG22 | 2.01 | 0.43 |
| 1:f3:410:ASP:OD1 | 1:f3:410:ASP:N | 2.50 | 0.43 |
| 1:X3:76:LEU:HD21 | 1:X3:103:LEU:HD11 | 2.00 | 0.43 |
| 1:A3:95:TRP:CD1 | 1:A3:180:LEU:HD13 | 2.54 | 0.43 |
| 1:d2:135:GLU:HB3 | 1:d3:451:ARG:HA | 2.01 | 0.43 |
| 1:d2:153:VAL:HG21 | 1:d2:329:LEU:HD13 | 2.00 | 0.43 |
| 1:d2:260:ASP:OD1 | 1:d2:263:ARG:NH2 | 2.52 | 0.43 |
| 1:f3:206:LEU:HD21 | 1:f3:514:ILE:HG12 | 2.01 | 0.43 |
| 1:X1:32:ARG:HE | 1:X3:24:ILE:HG21 | 1.83 | 0.43 |
| 1:X1:369:PHE:HB2 | 1:X1:372:PRO:HB3 | 2.00 | 0.43 |
| 1:e2:148:THR:O | 1:e2:152:TYR:HB2 | 2.19 | 0.43 |
| 1:e1:153:VAL:HG21 | 1:e1:329:LEU:HD13 | 2.00 | 0.43 |
| 1:e3:66:LEU:HB2 | 1:e3:255:ILE:HB | 2.01 | 0.43 |
| 1:e3:410:ASP:OD1 | 1:e3:410:ASP:N | 2.51 | 0.43 |
| 5:TP:226:GLY:C | 5:TP:228:ASN:H | 2.27 | 0.43 |
| 1:B2:22:PRO:HB3 | 1:B3:34:HIS:C | 2.44 | 0.43 |

*Continued on next page...*

For Manuscript Review

*Continued from previous page...*

| Atom-1 | Atom-2 | Interatomic distance (Å) | Clash overlap (Å) |
|---|---|---|---|
| 1:c1:13:ALA:O | 1:c1:14:GLN:HG2 | 2.18 | 0.43 |
| 1:f1:513:ARG:NH1 | 6:ze:113:ASN:OD1 | 2.52 | 0.43 |
| 1:f3:204:ILE:HG13 | 1:f3:266:ALA:HB2 | 1.99 | 0.43 |
| 1:X2:422:ALA:HB3 | 1:X2:440:PHE:CE2 | 2.53 | 0.43 |
| 1:X3:313:VAL:HG12 | 1:X3:333:PRO:HA | 2.01 | 0.43 |
| 1:X3:469:HIS:O | 1:X3:509:TYR:OH | 2.30 | 0.43 |
| 1:A2:101:GLU:OE2 | 1:A2:140:TYR:OH | 2.24 | 0.43 |
| 1:A1:7:GLN:NE2 | 1:A3:474:THR:O | 2.52 | 0.43 |
| 1:e1:299:PRO:HA | 1:e1:476:ASN:HA | 2.01 | 0.43 |
| 10:76:185:LEU:HD12 | 10:76:185:LEU:HA | 1.74 | 0.43 |
| 1:B3:306:VAL:HG12 | 1:B3:457:ILE:HA | 2.01 | 0.42 |
| 1:c2:133:SER:HA | 1:c1:135:GLU:HG2 | 2.00 | 0.42 |
| 1:d2:28:LYS:HD2 | 1:d2:28:LYS:HA | 1.84 | 0.42 |
| 1:d3:372:PRO:HG2 | 1:d3:373:PHE:CD2 | 2.54 | 0.42 |
| 1:d3:476:ASN:C | 1:d3:478:SER:H | 2.26 | 0.42 |
| 1:f1:189:LEU:HD13 | 1:f1:191:PHE:HE2 | 1.84 | 0.42 |
| 1:f1:199:LEU:HD11 | 1:f1:469:HIS:CD2 | 2.54 | 0.42 |
| 1:f3:278:GLN:HG3 | 1:f3:508:ASN:ND2 | 2.34 | 0.42 |
| 1:X2:91:VAL:HG22 | 1:X2:237:ILE:HD13 | 2.01 | 0.42 |
| 1:X1:55:ARG:HA | 1:X1:216:ASP:HA | 2.00 | 0.42 |
| 1:X3:44:GLN:NE2 | 1:X3:254:TYR:OH | 2.52 | 0.42 |
| 1:A2:26:PHE:HE2 | 1:A1:472:SER:HA | 1.84 | 0.42 |
| 1:e1:396:VAL:HB | 1:e3:394:SER:HB2 | 2.00 | 0.42 |
| 1:e3:111:ILE:HG12 | 1:e3:211:VAL:HG22 | 2.00 | 0.42 |
| 1:e3:162:ASP:N | 1:e3:162:ASP:OD1 | 2.38 | 0.42 |
| 9:uB:169:PRO:HB3 | 9:uB:192:PHE:CG | 2.54 | 0.42 |
| 1:d2:83:LEU:HD23 | 1:d2:240:ALA:HA | 1.99 | 0.42 |
| 1:f1:278:GLN:HB2 | 1:f1:506:ALA:HB3 | 2.01 | 0.42 |
| 1:X2:51:ASP:OD1 | 1:X2:52:PHE:N | 2.49 | 0.42 |
| 1:X2:425:GLN:HB2 | 1:X2:484:THR:HG22 | 2.01 | 0.42 |
| 1:X2:438:THR:O | 1:X2:442:LEU:HB2 | 2.19 | 0.42 |
| 1:X2:442:LEU:HD23 | 1:X2:452:ASN:HD21 | 1.83 | 0.42 |
| 1:X3:204:ILE:HG13 | 1:X3:266:ALA:HB2 | 2.01 | 0.42 |
| 1:X3:298:HIS:H | 1:X3:478:SER:HG | 1.65 | 0.42 |
| 1:e2:369:PHE:HB2 | 1:e2:372:PRO:HB3 | 2.00 | 0.42 |
| 1:e1:422:ALA:HB3 | 1:e1:440:PHE:CE2 | 2.54 | 0.42 |
| 1:e3:60:ILE:HD12 | 1:e3:252:VAL:HG21 | 2.00 | 0.42 |
| 1:e3:162:ASP:HB2 | 1:e3:171:ARG:NH2 | 2.33 | 0.42 |
| 1:e3:494:VAL:O | 9:uB:191:VAL:HG11 | 2.18 | 0.42 |
| 4:57:246:ILE:HB | 4:57:247:PRO:HD3 | 2.01 | 0.42 |
| 1:B2:7:GLN:OE1 | 1:B1:431:ARG:NH2 | 2.52 | 0.42 |

*Continued on next page...*

For Manuscript Review

Continued from previous page...

| Atom-1 | Atom-2 | Interatomic distance (Å) | Clash overlap (Å) |
|---|---|---|---|
| 1:d2:95:TRP:CD1 | 1:d2:180:LEU:HD13 | 2.55 | 0.42 |
| 1:f1:448:HIS:CD2 | 1:f1:461:SER:HB3 | 2.54 | 0.42 |
| 1:f3:138:ARG:HG2 | 1:f3:142:LYS:HE2 | 2.02 | 0.42 |
| 1:f3:371:ASP:O | 1:f3:397:SER:HB2 | 2.19 | 0.42 |
| 1:X2:148:THR:O | 1:X2:152:TYR:HB2 | 2.19 | 0.42 |
| 1:X1:155:ASP:HB2 | 1:X1:348:HIS:CE1 | 2.54 | 0.42 |
| 1:A2:26:PHE:CE2 | 1:A1:472:SER:HA | 2.53 | 0.42 |
| 1:A2:66:LEU:HD11 | 1:A2:257:LEU:HD11 | 2.01 | 0.42 |
| 1:A1:51:ASP:OD1 | 1:A1:51:ASP:N | 2.52 | 0.42 |
| 1:B2:469:HIS:O | 1:B2:509:TYR:OH | 2.30 | 0.42 |
| 1:c1:159:ALA:HA | 1:c1:359:THR:HG23 | 2.00 | 0.42 |
| 1:c3:316:CYS:HB2 | 10:76:153:SER:HA | 2.01 | 0.42 |
| 1:d2:371:ASP:O | 1:d2:397:SER:HB2 | 2.19 | 0.42 |
| 1:d3:122:ASP:OD1 | 1:d3:336:TYR:OH | 2.29 | 0.42 |
| 1:f3:369:PHE:HB2 | 1:f3:372:PRO:HB3 | 2.02 | 0.42 |
| 1:X2:291:LYS:HZ3 | 1:X2:484:THR:HG21 | 1.84 | 0.42 |
| 1:X2:394:SER:HB2 | 1:X3:396:VAL:HB | 2.01 | 0.42 |
| 1:A2:343:LEU:HD21 | 1:A2:408:ALA:HB1 | 2.00 | 0.42 |
| 1:e1:107:VAL:HG22 | 1:e1:215:ILE:HG12 | 2.01 | 0.42 |
| 9:uB:70:ARG:HH21 | 9:uB:72:GLN:HE21 | 1.67 | 0.42 |
| 1:c2:336:TYR:O | 1:c2:456:GLY:HA3 | 2.20 | 0.42 |
| 1:f2:53:GLY:N | 1:f2:217:ILE:O | 2.53 | 0.42 |
| 1:f1:282:ASP:HB3 | 1:f1:503:ARG:HG2 | 2.01 | 0.42 |
| 1:X1:106:GLN:N | 1:X1:216:ASP:OD1 | 2.52 | 0.42 |
| 1:X3:67:ALA:HB3 | 1:X3:201:LEU:HB3 | 2.02 | 0.42 |
| 1:A2:240:ALA:C | 1:A2:242:LYS:H | 2.27 | 0.42 |
| 1:A2:354:THR:OG1 | 1:A1:376:ASP:OD2 | 2.37 | 0.42 |
| 1:A2:369:PHE:O | 1:A2:372:PRO:HD3 | 2.20 | 0.42 |
| 8:gc:96:THR:HG22 | 8:gc:162:ILE:HG12 | 2.00 | 0.42 |
| 1:B2:420:VAL:HG12 | 1:B2:421:VAL:HG23 | 2.00 | 0.42 |
| 1:B1:148:THR:HG23 | 1:B1:152:TYR:CD2 | 2.54 | 0.42 |
| 1:B1:354:THR:OG1 | 1:B3:376:ASP:OD2 | 2.31 | 0.42 |
| 1:B3:46:PHE:HB3 | 1:B3:56:VAL:HB | 2.02 | 0.42 |
| 1:c2:355:GLN:NE2 | 1:c2:371:ASP:OD2 | 2.52 | 0.42 |
| 1:c1:68:TYR:O | 1:c1:194:CYS:HA | 2.19 | 0.42 |
| 1:c1:283:GLU:HG2 | 1:c1:292:ILE:HG13 | 2.02 | 0.42 |
| 1:c3:66:LEU:HB2 | 1:c3:255:ILE:HB | 2.02 | 0.42 |
| 1:f1:192:TRP:HZ2 | 1:f1:201:LEU:HD21 | 1.84 | 0.42 |
| 1:f3:132:LEU:O | 1:f3:451:ARG:HD2 | 2.18 | 0.42 |
| 1:f3:299:PRO:HA | 1:f3:476:ASN:HA | 2.00 | 0.42 |
| 1:X2:6:MET:HE3 | 1:X1:429:GLN:HG3 | 2.02 | 0.42 |

Continued on next page...

For Manuscript Review

*Continued from previous page...*

| Atom-1 | Atom-2 | Interatomic distance (Å) | Clash overlap (Å) |
|---|---|---|---|
| 1:X3:299:PRO:HA | 1:X3:476:ASN:HA | 2.01 | 0.42 |
| 1:A3:299:PRO:HA | 1:A3:476:ASN:HA | 2.01 | 0.42 |
| 1:A3:315:TYR:O | 1:A3:318:SER:OG | 2.31 | 0.42 |
| 1:e1:231:ASP:OD1 | 1:e1:231:ASP:N | 2.52 | 0.42 |
| 4:57:224:SER:O | 4:57:228:ASN:ND2 | 2.46 | 0.42 |
| 8:gc:75:ARG:HD2 | 8:gc:89:ASN:HB2 | 2.02 | 0.42 |
| 10:76:184:ARG:NH2 | 10:76:209:SER:HA | 2.33 | 0.42 |
| 1:B1:425:GLN:HB2 | 1:B1:484:THR:OG1 | 2.19 | 0.42 |
| 1:c2:343:LEU:HA | 1:c1:369:PHE:HE2 | 1.85 | 0.42 |
| 1:f1:284:SER:OG | 9:uB:169:PRO:HG2 | 2.20 | 0.42 |
| 1:f3:99:PRO:O | 1:f3:103:LEU:HG | 2.19 | 0.42 |
| 1:A1:76:LEU:HD11 | 1:A1:185:LEU:HD11 | 2.02 | 0.42 |
| 1:A3:148:THR:O | 1:A3:152:TYR:HB2 | 2.20 | 0.42 |
| 1:A3:236:LYS:HD3 | 1:A3:319:LEU:HD13 | 2.01 | 0.42 |
| 1:B3:43:GLU:HB2 | 1:B3:251:PHE:CE2 | 2.54 | 0.42 |
| 1:B3:317:ALA:HB1 | 1:B3:323:GLN:HB3 | 2.01 | 0.42 |
| 1:c2:12:GLY:HA2 | 1:c1:479:ARG:HH11 | 1.84 | 0.42 |
| 1:c3:67:ALA:HB2 | 1:c3:203:LEU:HD11 | 2.02 | 0.42 |
| 1:c3:343:LEU:HD21 | 1:c3:408:ALA:HB1 | 2.01 | 0.42 |
| 1:c3:515:MET:HG3 | 8:gc:80:PHE:HB2 | 2.02 | 0.42 |
| 1:X1:76:LEU:HD11 | 1:X1:103:LEU:HD11 | 2.02 | 0.42 |
| 1:e2:501:LYS:HB2 | 1:e2:501:LYS:HE3 | 1.83 | 0.42 |
| 1:e1:158:PHE:CD2 | 1:e1:176:PRO:HB2 | 2.55 | 0.42 |
| 1:e1:516:SER:HB2 | 5:TP:194:LEU:HD11 | 2.02 | 0.42 |
| 1:c2:273:LEU:HD22 | 1:c1:31:TYR:CZ | 2.55 | 0.42 |
| 1:c2:373:PHE:HA | 1:c3:349:ALA:HA | 2.02 | 0.42 |
| 1:c3:170:VAL:HB | 1:c3:177:ARG:HH12 | 1.85 | 0.42 |
| 1:c3:523:TYR:O | 1:c3:524:SER:OG | 2.31 | 0.42 |
| 1:f2:93:ALA:HA | 1:f2:225:TRP:O | 2.20 | 0.42 |
| 1:f3:66:LEU:HD23 | 1:f3:202:PRO:HA | 2.02 | 0.42 |
| 1:X2:39:MET:HG2 | 1:X2:255:ILE:HD12 | 2.02 | 0.42 |
| 1:X2:107:VAL:HG22 | 1:X2:215:ILE:HG23 | 2.01 | 0.42 |
| 1:X1:300:CYS:HA | 1:X1:508:ASN:HA | 2.00 | 0.42 |
| 1:A2:148:THR:O | 1:A2:152:TYR:HB2 | 2.20 | 0.42 |
| 7:he:36:TYR:CG | 7:he:89:VAL:HG21 | 2.55 | 0.42 |
| 1:B2:288:SER:HB2 | 3:wD:18:ASN:HB3 | 2.01 | 0.42 |
| 1:B2:510:ASN:OD1 | 1:B2:511:VAL:N | 2.47 | 0.42 |
| 1:c2:134:LYS:HB3 | 1:c3:134:LYS:HG2 | 2.02 | 0.42 |
| 1:c2:257:LEU:HB2 | 1:c2:262:ARG:HG3 | 2.02 | 0.42 |
| 1:c1:343:LEU:HD22 | 1:c3:402:PHE:HE2 | 1.84 | 0.42 |
| 1:A1:320:GLN:O | 1:A1:326:ASN:ND2 | 2.37 | 0.42 |

*Continued on next page...*

For Manuscript Review

Continued from previous page...

| Atom-1 | Atom-2 | Interatomic distance (Å) | Clash overlap (Å) |
|---|---|---|---|
| 1:e1:320:GLN:O | 1:e1:326:ASN:ND2 | 2.53 | 0.42 |
| 4:57:209:ARG:HG2 | 4:57:217:ILE:HG22 | 2.02 | 0.42 |
| 7:he:55:SER:O | 7:he:59:ILE:HG13 | 2.20 | 0.42 |
| 1:d3:229:SER:OG | 1:d3:231:ASP:OD1 | 2.36 | 0.41 |
| 1:f3:108:GLU:O | 1:f3:213:ILE:HA | 2.20 | 0.41 |
| 1:X3:510:ASN:OD1 | 1:X3:511:VAL:N | 2.48 | 0.41 |
| 1:A2:24:ILE:HG21 | 1:A3:32:ARG:HD3 | 2.02 | 0.41 |
| 1:A1:257:LEU:HB2 | 1:A1:262:ARG:HG3 | 2.01 | 0.41 |
| 1:e3:436:GLU:HG3 | 9:uB:157:LEU:HD21 | 2.02 | 0.41 |
| 1:B2:372:PRO:HG2 | 1:B2:373:PHE:CD2 | 2.55 | 0.41 |
| 1:B1:176:PRO:HB3 | 1:B1:347:VAL:HG21 | 2.02 | 0.41 |
| 1:d2:444:GLN:HB2 | 1:d2:445:PRO:HD3 | 2.03 | 0.41 |
| 1:d3:199:LEU:HD11 | 1:d3:469:HIS:CD2 | 2.55 | 0.41 |
| 1:f2:202:PRO:HB2 | 1:f2:205:ALA:HB3 | 2.02 | 0.41 |
| 1:f1:148:THR:HG23 | 1:f1:152:TYR:CD2 | 2.55 | 0.41 |
| 1:X1:26:PHE:CE2 | 1:X3:472:SER:HA | 2.56 | 0.41 |
| 1:X3:464:LEU:O | 1:X3:465:ARG:HD2 | 2.20 | 0.41 |
| 1:A1:162:ASP:OD1 | 1:A1:162:ASP:N | 2.52 | 0.41 |
| 1:A1:208:TYR:HD2 | 1:A1:517:GLY:HA3 | 1.85 | 0.41 |
| 1:e1:239:ASP:HA | 1:e1:242:LYS:HE2 | 2.01 | 0.41 |
| 1:e1:257:LEU:HB2 | 1:e1:262:ARG:HG3 | 2.02 | 0.41 |
| 1:B2:278:GLN:NE2 | 1:B2:300:CYS:SG | 2.93 | 0.41 |
| 1:B1:133:SER:HA | 1:B3:135:GLU:HG2 | 2.02 | 0.41 |
| 1:c2:6:MET:HE2 | 1:c2:6:MET:HB2 | 1.90 | 0.41 |
| 1:c2:107:VAL:HG12 | 1:c2:191:PHE:CZ | 2.55 | 0.41 |
| 1:c2:147:THR:O | 1:c2:151:THR:OG1 | 2.26 | 0.41 |
| 1:c1:148:THR:O | 1:c1:152:TYR:HB2 | 2.21 | 0.41 |
| 1:c1:478:SER:O | 7:he:156:LYS:HD3 | 2.19 | 0.41 |
| 1:c1:525:ASN:HB2 | 1:X3:427:ASN:HB3 | 2.02 | 0.41 |
| 1:f1:110:GLU:HB3 | 1:f1:115:ARG:HA | 2.01 | 0.41 |
| 1:f1:170:VAL:HG13 | 1:f1:363:ILE:HG21 | 2.01 | 0.41 |
| 1:X2:132:LEU:O | 1:X2:451:ARG:HD2 | 2.21 | 0.41 |
| 1:A1:148:THR:HG23 | 1:A1:152:TYR:CD2 | 2.55 | 0.41 |
| 1:A3:62:ARG:NH1 | 1:A3:206:LEU:O | 2.53 | 0.41 |
| 1:e2:287:SER:HA | 9:uB:18:TRP:CG | 2.56 | 0.41 |
| 1:e2:435:ARG:NH2 | 1:e1:73:GLN:OE1 | 2.53 | 0.41 |
| 1:e3:299:PRO:HA | 1:e3:476:ASN:HA | 2.02 | 0.41 |
| 9:uB:73:LEU:HD12 | 9:uB:73:LEU:HA | 1.87 | 0.41 |
| 1:B1:46:PHE:CD1 | 1:B1:58:CYS:HB2 | 2.55 | 0.41 |
| 1:c2:44:GLN:HE22 | 1:c2:61:SER:H | 1.67 | 0.41 |
| 1:c1:384:HIS:HB2 | 1:c1:387:PHE:HD2 | 1.84 | 0.41 |

Continued on next page...

For Manuscript Review

*Continued from previous page...*

| Atom-1 | Atom-2 | Interatomic distance (Å) | Clash overlap (Å) |
|---|---|---|---|
| 1:c3:111:ILE:HG12 | 1:c3:211:VAL:HG22 | 2.02 | 0.41 |
| 1:c3:372:PRO:HG2 | 1:c3:373:PHE:CD2 | 2.56 | 0.41 |
| 1:d2:355:GLN:HE22 | 1:d2:397:SER:HA | 1.85 | 0.41 |
| 1:d3:427:ASN:ND2 | 1:d3:481:ASP:H | 2.15 | 0.41 |
| 1:f2:81:GLN:NE2 | 1:f2:178:LYS:O | 2.41 | 0.41 |
| 1:f1:43:GLU:HB2 | 1:f1:251:PHE:CZ | 2.55 | 0.41 |
| 1:f1:94:ARG:HD2 | 1:f1:153:VAL:HG22 | 2.01 | 0.41 |
| 1:f1:325:LEU:HD22 | 1:f1:331:ALA:HA | 2.02 | 0.41 |
| 1:X2:201:LEU:HD12 | 1:X2:201:LEU:HA | 1.86 | 0.41 |
| 1:X1:120:TYR:O | 1:X1:124:MET:HG3 | 2.21 | 0.41 |
| 1:A2:54:ARG:CZ | 3:wD:85:SER:HB2 | 2.50 | 0.41 |
| 1:A2:99:PRO:O | 1:A2:103:LEU:HG | 2.20 | 0.41 |
| 1:B2:99:PRO:O | 1:B2:103:LEU:HG | 2.20 | 0.41 |
| 1:B2:155:ASP:HB3 | 1:B2:158:PHE:CE2 | 2.55 | 0.41 |
| 1:c1:176:PRO:HB3 | 1:c1:347:VAL:HG21 | 2.01 | 0.41 |
| 1:d2:162:ASP:OD1 | 1:d2:162:ASP:N | 2.50 | 0.41 |
| 1:d1:39:MET:HG2 | 1:d1:255:ILE:HD12 | 2.01 | 0.41 |
| 1:d1:46:PHE:CD1 | 1:d1:58:CYS:HB2 | 2.55 | 0.41 |
| 1:f3:125:HIS:CE1 | 1:f3:129:GLN:HE21 | 2.39 | 0.41 |
| 1:X2:444:GLN:HB2 | 1:X2:445:PRO:HD3 | 2.02 | 0.41 |
| 1:X3:148:THR:O | 1:X3:152:TYR:HB2 | 2.21 | 0.41 |
| 1:A2:92:TYR:CD2 | 1:A2:230:MET:HG3 | 2.56 | 0.41 |
| 1:A2:195:ARG:NH2 | 1:A2:275:GLU:OE1 | 2.54 | 0.41 |
| 1:e2:173:VAL:HG11 | 1:e3:416:GLU:HG3 | 2.03 | 0.41 |
| 1:e1:33:ARG:NE | 1:e3:20:GLY:O | 2.53 | 0.41 |
| 1:e1:133:SER:HA | 1:e3:135:GLU:HG2 | 2.02 | 0.41 |
| 1:e1:204:ILE:HG13 | 1:e1:266:ALA:HB2 | 2.03 | 0.41 |
| 8:gc:278:THR:HG22 | 8:gc:279:TYR:H | 1.85 | 0.41 |
| 1:B2:39:MET:HE1 | 1:B3:448:HIS:HE1 | 1.85 | 0.41 |
| 1:B2:79:ILE:HB | 1:B2:180:LEU:HD12 | 2.02 | 0.41 |
| 1:d1:476:ASN:C | 1:d1:478:SER:H | 2.28 | 0.41 |
| 1:d3:464:LEU:O | 1:d3:465:ARG:HD2 | 2.21 | 0.41 |
| 1:f2:282:ASP:HB3 | 1:f2:501:LYS:HG2 | 2.03 | 0.41 |
| 1:X2:513:ARG:O | 1:X2:519:GLY:HA2 | 2.21 | 0.41 |
| 1:X1:61:SER:O | 1:X1:203:LEU:HD13 | 2.20 | 0.41 |
| 1:X1:106:GLN:OE1 | 1:X1:118:ARG:NE | 2.53 | 0.41 |
| 3:wD:108:ILE:HD12 | 3:wD:108:ILE:HA | 1.86 | 0.41 |
| 9:uB:158:HIS:HA | 9:uB:166:GLN:HE22 | 1.86 | 0.41 |
| 1:B2:148:THR:O | 1:B2:152:TYR:HB2 | 2.19 | 0.41 |
| 1:B1:76:LEU:HD21 | 1:B1:103:LEU:HD11 | 2.02 | 0.41 |
| 1:B3:372:PRO:HG2 | 1:B3:373:PHE:CD2 | 2.55 | 0.41 |

*Continued on next page...*

For Manuscript Review

*Continued from previous page...*

| Atom-1 | Atom-2 | Interatomic distance (Å) | Clash overlap (Å) |
|---|---|---|---|
| 1:c2:132:LEU:HD21 | 1:c2:137:GLU:HA | 2.02 | 0.41 |
| 1:c2:391:THR:OG1 | 1:c3:370:GLN:NE2 | 2.53 | 0.41 |
| 1:c3:39:MET:HG2 | 1:c3:255:ILE:HD12 | 2.03 | 0.41 |
| 1:d2:205:ALA:O | 1:d2:517:GLY:HA2 | 2.21 | 0.41 |
| 1:f2:68:TYR:O | 1:f2:194:CYS:HA | 2.21 | 0.41 |
| 1:f1:300:CYS:HA | 1:f1:508:ASN:HA | 2.02 | 0.41 |
| 1:X2:447:GLN:HE21 | 1:X2:448:HIS:CE1 | 2.39 | 0.41 |
| 1:X1:110:GLU:HB3 | 1:X1:115:ARG:HA | 2.03 | 0.41 |
| 1:X1:153:VAL:HG11 | 1:X1:230:MET:HE2 | 2.02 | 0.41 |
| 1:A1:158:PHE:CD2 | 1:A1:176:PRO:HB2 | 2.55 | 0.41 |
| 1:A1:208:TYR:CD2 | 1:A1:517:GLY:HA3 | 2.55 | 0.41 |
| 1:A3:343:LEU:HD21 | 1:A3:408:ALA:HB1 | 2.03 | 0.41 |
| 1:e3:337:THR:HG22 | 1:e3:455:SER:HB2 | 2.02 | 0.41 |
| 8:gc:289:LEU:HD22 | 8:gc:323:TYR:CD2 | 2.56 | 0.41 |
| 1:c3:148:THR:O | 1:c3:152:TYR:HB2 | 2.20 | 0.41 |
| 1:d1:427:ASN:HA | 1:d1:482:ASN:HB2 | 2.03 | 0.41 |
| 1:f1:40:GLU:OE1 | 1:f3:4:GLY:N | 2.54 | 0.41 |
| 1:X1:301:LYS:HD3 | 1:X1:509:TYR:CE2 | 2.56 | 0.41 |
| 1:A2:431:ARG:NH2 | 1:A3:7:GLN:OE1 | 2.45 | 0.41 |
| 1:A1:39:MET:HB3 | 1:A1:39:MET:HE2 | 1.82 | 0.41 |
| 1:A1:94:ARG:HD2 | 1:A1:151:THR:O | 2.21 | 0.41 |
| 1:A1:361:GLY:O | 1:A1:363:ILE:N | 2.51 | 0.41 |
| 1:A3:193:PHE:HA | 1:A3:199:LEU:O | 2.21 | 0.41 |
| 1:e2:239:ASP:HA | 1:e2:242:LYS:HE2 | 2.02 | 0.41 |
| 1:B1:225:TRP:CD1 | 1:B1:330:GLY:HA2 | 2.56 | 0.41 |
| 1:B1:305:TRP:HA | 1:B1:503:ARG:O | 2.20 | 0.41 |
| 1:B3:55:ARG:NH2 | 1:B3:214:ASN:OD1 | 2.54 | 0.41 |
| 1:c1:50:ALA:HB3 | 1:c1:248:ALA:HB3 | 2.01 | 0.41 |
| 1:c1:72:LEU:HG | 1:c1:74:VAL:HG23 | 2.02 | 0.41 |
| 1:c1:192:TRP:HZ2 | 1:c1:201:LEU:HD21 | 1.85 | 0.41 |
| 1:c1:307:VAL:O | 1:c1:418:PRO:HG2 | 2.21 | 0.41 |
| 1:d2:98:PHE:HB2 | 1:d2:334:PHE:CE2 | 2.56 | 0.41 |
| 1:d2:101:GLU:OE2 | 1:d2:140:TYR:OH | 2.30 | 0.41 |
| 1:d3:111:ILE:HG12 | 1:d3:211:VAL:HG22 | 2.03 | 0.41 |
| 1:f2:276:GLN:HE22 | 1:f2:510:ASN:HD22 | 1.68 | 0.41 |
| 1:f2:501:LYS:HE3 | 1:f2:501:LYS:HB2 | 1.89 | 0.41 |
| 1:X2:73:GLN:OE1 | 1:X3:435:ARG:NH2 | 2.48 | 0.41 |
| 1:X2:306:VAL:HG12 | 1:X2:457:ILE:HA | 2.03 | 0.41 |
| 1:X2:332:GLN:HB3 | 1:X2:335:ASN:HB2 | 2.03 | 0.41 |
| 1:X1:257:LEU:HB2 | 1:X1:262:ARG:HG3 | 2.03 | 0.41 |
| 1:A2:130:LEU:HD21 | 1:A2:459:VAL:HG21 | 2.02 | 0.41 |

*Continued on next page...*

For Manuscript Review

*Continued from previous page...*

| Atom-1 | Atom-2 | Interatomic distance (Å) | Clash overlap (Å) |
|---|---|---|---|
| 1:A2:472:SER:HA | 1:A3:26:PHE:CE2 | 2.56 | 0.41 |
| 1:e2:299:PRO:HA | 1:e2:476:ASN:HA | 2.02 | 0.41 |
| 1:e2:411:MET:HE1 | 1:e1:403:VAL:HA | 2.02 | 0.41 |
| 1:e1:53:GLY:HA2 | 1:e1:217:ILE:O | 2.21 | 0.41 |
| 1:e1:387:PHE:CE1 | 1:e3:231:ASP:HA | 2.56 | 0.41 |
| 1:e3:307:VAL:HG22 | 1:e3:502:VAL:HG22 | 2.03 | 0.41 |
| 1:e3:325:LEU:HD23 | 1:e3:328:LEU:HD12 | 2.02 | 0.41 |
| 1:e3:417:ASN:ND2 | 1:e3:419:VAL:O | 2.54 | 0.41 |
| 8:gc:208:LYS:HG2 | 8:gc:209:ASN:ND2 | 2.36 | 0.41 |
| 8:gc:229:THR:HG22 | 5:TQ:158:PRO:HG2 | 2.02 | 0.41 |
| 9:uB:173:ILE:HD13 | 9:uB:173:ILE:HA | 1.84 | 0.41 |
| 10:76:148:ASN:HA | 10:76:159:TYR:HB2 | 2.03 | 0.41 |
| 1:B3:68:TYR:O | 1:B3:194:CYS:HA | 2.21 | 0.41 |
| 1:B3:75:THR:HB | 1:B3:246:ALA:HB3 | 2.02 | 0.41 |
| 1:c2:36:ASN:HD21 | 5:TQ:144:GLN:NE2 | 2.19 | 0.41 |
| 1:c2:76:LEU:O | 1:c2:182:GLU:HG3 | 2.21 | 0.41 |
| 1:X2:469:HIS:O | 1:X2:509:TYR:OH | 2.26 | 0.41 |
| 1:A2:43:GLU:HB2 | 1:A2:251:PHE:CE2 | 2.56 | 0.41 |
| 1:A2:472:SER:HA | 1:A3:26:PHE:HE2 | 1.86 | 0.41 |
| 1:A1:99:PRO:O | 1:A1:103:LEU:HG | 2.21 | 0.41 |
| 1:A3:132:LEU:HD21 | 1:A3:137:GLU:HA | 2.03 | 0.41 |
| 1:e2:55:ARG:N | 9:uB:139:LYS:O | 2.53 | 0.41 |
| 1:e1:69:ARG:H | 1:e1:253:ASP:HB3 | 1.86 | 0.41 |
| 1:e3:66:LEU:HD23 | 1:e3:202:PRO:HA | 2.02 | 0.41 |
| 1:c2:107:VAL:HG12 | 1:c2:191:PHE:HZ | 1.86 | 0.40 |
| 1:c1:364:ASN:ND2 | 1:c1:369:PHE:O | 2.51 | 0.40 |
| 1:d2:495:GLN:O | 1:d2:498:SER:OG | 2.30 | 0.40 |
| 1:d1:52:PHE:HE1 | 1:d1:103:LEU:HD22 | 1.86 | 0.40 |
| 1:f1:387:PHE:CE1 | 1:f3:231:ASP:HA | 2.56 | 0.40 |
| 1:X1:148:THR:O | 1:X1:152:TYR:HB2 | 2.21 | 0.40 |
| 1:X1:306:VAL:HG12 | 1:X1:457:ILE:HA | 2.03 | 0.40 |
| 1:A1:128:CYS:SG | 1:A1:190:GLN:NE2 | 2.94 | 0.40 |
| 1:A1:408:ALA:HB2 | 1:A3:403:VAL:HG21 | 2.04 | 0.40 |
| 1:A3:464:LEU:HB2 | 1:A3:472:SER:HB2 | 2.02 | 0.40 |
| 1:e2:54:ARG:NH1 | 9:uB:142:THR:HG23 | 2.36 | 0.40 |
| 1:B1:420:VAL:HG12 | 1:B1:421:VAL:HG23 | 2.03 | 0.40 |
| 1:c1:83:LEU:HD23 | 9:uB:42:ASN:ND2 | 2.36 | 0.40 |
| 1:c3:225:TRP:CZ2 | 1:c3:227:MET:HG3 | 2.56 | 0.40 |
| 1:d2:303:LEU:HG | 1:d2:462:PHE:HE2 | 1.86 | 0.40 |
| 1:d1:205:ALA:O | 1:d1:517:GLY:HA2 | 2.22 | 0.40 |
| 1:f2:225:TRP:CD1 | 1:f2:330:GLY:HA2 | 2.57 | 0.40 |

*Continued on next page...*

For Manuscript Review

*Continued from previous page...*

| Atom-1 | Atom-2 | Interatomic distance (Å) | Clash overlap (Å) |
|---|---|---|---|
| 1:f2:403:VAL:HA | 1:f3:411:MET:HE1 | 2.02 | 0.40 |
| 1:f3:98:PHE:HB2 | 1:f3:334:PHE:CE2 | 2.56 | 0.40 |
| 1:X2:32:ARG:CZ | 1:X1:24:ILE:HG21 | 2.50 | 0.40 |
| 1:X2:107:VAL:HG12 | 1:X2:191:PHE:HZ | 1.86 | 0.40 |
| 1:X3:66:LEU:HD23 | 1:X3:202:PRO:HA | 2.03 | 0.40 |
| 1:A2:291:LYS:HE2 | 1:A2:291:LYS:HB2 | 1.86 | 0.40 |
| 1:A1:170:VAL:HG13 | 1:A1:363:ILE:HG21 | 2.03 | 0.40 |
| 1:A3:315:TYR:HE1 | 1:A3:333:PRO:HG3 | 1.86 | 0.40 |
| 1:A3:384:HIS:HB2 | 1:A3:387:PHE:HD2 | 1.86 | 0.40 |
| 1:e1:191:PHE:O | 1:e1:195:ARG:HG3 | 2.21 | 0.40 |
| 5:TP:229:ALA:HA | 5:TP:232:MET:HG3 | 2.03 | 0.40 |
| 8:gc:170:ILE:O | 8:gc:174:ILE:HG13 | 2.22 | 0.40 |
| 1:d2:161:VAL:HA | 1:d2:177:ARG:HD2 | 2.02 | 0.40 |
| 1:d1:116:ILE:HG23 | 1:d1:521:LEU:HD21 | 2.02 | 0.40 |
| 1:d3:250:LEU:HD23 | 1:d3:250:LEU:HA | 1.99 | 0.40 |
| 1:d3:291:LYS:NZ | 1:X3:43:GLU:OE2 | 2.50 | 0.40 |
| 1:f2:153:VAL:HG21 | 1:f2:329:LEU:HD13 | 2.04 | 0.40 |
| 1:f2:431:ARG:NH2 | 1:f3:7:GLN:OE1 | 2.51 | 0.40 |
| 1:X1:66:LEU:HB2 | 1:X1:255:ILE:HB | 2.04 | 0.40 |
| 1:X3:93:ALA:HA | 1:X3:225:TRP:O | 2.22 | 0.40 |
| 1:A2:164:PRO:HB2 | 3:wD:43:PHE:CD2 | 2.57 | 0.40 |
| 1:A1:37:PHE:HE1 | 1:A1:39:MET:HE3 | 1.86 | 0.40 |
| 1:A3:423:LYS:HE2 | 1:A3:423:LYS:HB3 | 1.86 | 0.40 |
| 1:e1:76:LEU:HD11 | 1:e1:185:LEU:HD11 | 2.03 | 0.40 |
| 8:gc:234:LEU:HD11 | 5:TQ:159:LEU:HD22 | 2.03 | 0.40 |
| 1:B2:395:GLY:HA3 | 1:B1:393:GLU:HB3 | 2.03 | 0.40 |
| 1:B3:99:PRO:O | 1:B3:103:LEU:HG | 2.20 | 0.40 |
| 1:c1:219:ASN:ND2 | 1:c1:219:ASN:H | 2.19 | 0.40 |
| 1:c1:422:ALA:HB3 | 1:c1:440:PHE:CE2 | 2.56 | 0.40 |
| 1:d2:50:ALA:HB3 | 1:d2:248:ALA:HB3 | 2.03 | 0.40 |
| 1:d1:127:TRP:CE2 | 1:d1:131:THR:HG21 | 2.56 | 0.40 |
| 1:X2:94:ARG:NH1 | 1:X2:329:LEU:O | 2.54 | 0.40 |
| 1:X3:75:THR:HB | 1:X3:246:ALA:HB3 | 2.04 | 0.40 |
| 1:A3:424:LEU:HB3 | 1:A3:432:PHE:CE2 | 2.57 | 0.40 |
| 1:e1:230:MET:HB2 | 1:e1:230:MET:HE3 | 1.86 | 0.40 |
| 1:e1:515:MET:HG2 | 5:TP:195:TRP:CZ2 | 2.56 | 0.40 |
| 8:gc:101:ILE:O | 8:gc:105:ASN:HB2 | 2.21 | 0.40 |
| 1:B1:257:LEU:HB2 | 1:B1:262:ARG:HG3 | 2.04 | 0.40 |
| 1:c1:257:LEU:HB2 | 1:c1:262:ARG:HG3 | 2.03 | 0.40 |
| 1:c3:305:TRP:HA | 1:c3:503:ARG:O | 2.22 | 0.40 |
| 1:d2:368:PHE:HZ | 1:d3:324:PRO:HA | 1.86 | 0.40 |

*Continued on next page...*

For Manuscript Review

*Continued from previous page...*

| Atom-1 | Atom-2 | Interatomic distance (Å) | Clash overlap (Å) |
|---|---|---|---|
| 1:d1:482:ASN:HB3 | 1:A2:295:ASN:HD21 | 1.87 | 0.40 |
| 1:f1:68:TYR:O | 1:f1:194:CYS:HA | 2.22 | 0.40 |
| 1:A1:108:GLU:HG2 | 1:A1:118:ARG:HG3 | 2.04 | 0.40 |
| 1:e1:160:ALA:HB2 | 1:e1:178:LYS:HE2 | 2.03 | 0.40 |
| 1:e3:491:ASN:OD1 | 1:e3:495:GLN:NE2 | 2.54 | 0.40 |
| 8:gc:295:ASP:OD1 | 8:gc:295:ASP:N | 2.54 | 0.40 |

There are no symmetry-related clashes.

#### 5.3 Torsion angles (i)

##### 5.3.1 Protein backbone (i)

In the following table, the Percentiles column shows the percent Ramachandran outliers of the chain as a percentile score with respect to all PDB entries followed by that with respect to all EM entries.

The Analysed column shows the number of residues for which the backbone conformation was analysed, and the total number of residues.

| Mol | Chain | Analysed | Favoured | Allowed | Outliers | Percentiles | |
|---|---|---|---|---|---|---|---|
| 1 | A1 | 522/525 (99%) | 503 (96%) | 19 (4%) | 0 | 100 | 100 |
| 1 | A2 | 522/525 (99%) | 502 (96%) | 20 (4%) | 0 | 100 | 100 |
| 1 | A3 | 522/525 (99%) | 509 (98%) | 13 (2%) | 0 | 100 | 100 |
| 1 | B1 | 522/525 (99%) | 505 (97%) | 17 (3%) | 0 | 100 | 100 |
| 1 | B2 | 522/525 (99%) | 506 (97%) | 16 (3%) | 0 | 100 | 100 |
| 1 | B3 | 522/525 (99%) | 513 (98%) | 9 (2%) | 0 | 100 | 100 |
| 1 | X1 | 522/525 (99%) | 506 (97%) | 16 (3%) | 0 | 100 | 100 |
| 1 | X2 | 522/525 (99%) | 506 (97%) | 16 (3%) | 0 | 100 | 100 |
| 1 | X3 | 522/525 (99%) | 507 (97%) | 15 (3%) | 0 | 100 | 100 |
| 1 | c1 | 518/525 (99%) | 495 (96%) | 23 (4%) | 0 | 100 | 100 |
| 1 | c2 | 522/525 (99%) | 507 (97%) | 15 (3%) | 0 | 100 | 100 |
| 1 | c3 | 522/525 (99%) | 503 (96%) | 19 (4%) | 0 | 100 | 100 |
| 1 | d1 | 522/525 (99%) | 511 (98%) | 11 (2%) | 0 | 100 | 100 |
| 1 | d2 | 522/525 (99%) | 502 (96%) | 20 (4%) | 0 | 100 | 100 |
| 1 | d3 | 522/525 (99%) | 504 (97%) | 18 (3%) | 0 | 100 | 100 |

*Continued on next page...*

For Manuscript Review

*Continued from previous page...*

| Mol | Chain | Analysed | Favoured | Allowed | Outliers | Percentiles | |
|---|---|---|---|---|---|---|---|
| 1 | e1 | 522/525 (99%) | 503 (96%) | 19 (4%) | 0 | 100 | 100 |
| 1 | e2 | 522/525 (99%) | 504 (97%) | 18 (3%) | 0 | 100 | 100 |
| 1 | e3 | 522/525 (99%) | 508 (97%) | 14 (3%) | 0 | 100 | 100 |
| 1 | f1 | 522/525 (99%) | 506 (97%) | 16 (3%) | 0 | 100 | 100 |
| 1 | f2 | 522/525 (99%) | 509 (98%) | 13 (2%) | 0 | 100 | 100 |
| 1 | f3 | 522/525 (99%) | 507 (97%) | 15 (3%) | 0 | 100 | 100 |
| 2 | vC | 24/120 (20%) | 23 (96%) | 1 (4%) | 0 | 100 | 100 |
| 3 | wD | 97/225 (43%) | 89 (92%) | 8 (8%) | 0 | 100 | 100 |
| 4 | 57 | 118/282 (42%) | 112 (95%) | 6 (5%) | 0 | 100 | 100 |
| 5 | TP | 92/608 (15%) | 87 (95%) | 5 (5%) | 0 | 100 | 100 |
| 5 | TQ | 70/608 (12%) | 65 (93%) | 5 (7%) | 0 | 100 | 100 |
| 6 | ze | 57/169 (34%) | 55 (96%) | 2 (4%) | 0 | 100 | 100 |
| 7 | he | 130/166 (78%) | 127 (98%) | 3 (2%) | 0 | 100 | 100 |
| 8 | gc | 272/329 (83%) | 265 (97%) | 7 (3%) | 0 | 100 | 100 |
| 9 | uB | 265/267 (99%) | 250 (94%) | 15 (6%) | 0 | 100 | 100 |
| 10 | 76 | 138/210 (66%) | 126 (91%) | 12 (9%) | 0 | 100 | 100 |
| All | All | 12221/14009 (87%) | 11815 (97%) | 406 (3%) | 0 | 100 | 100 |

There are no Ramachandran outliers to report.

##### 5.3.2 Protein sidechains ⓘ

In the following table, the Percentiles column shows the percent sidechain outliers of the chain as a percentile score with respect to all PDB entries followed by that with respect to all EM entries.

The Analysed column shows the number of residues for which the sidechain conformation was analysed, and the total number of residues.

| Mol | Chain | Analysed | Rotameric | Outliers | Percentiles | |
|---|---|---|---|---|---|---|
| 1 | A1 | 442/443 (100%) | 438 (99%) | 4 (1%) | 75 | 86 |
| 1 | A2 | 442/443 (100%) | 440 (100%) | 2 (0%) | 86 | 91 |
| 1 | A3 | 442/443 (100%) | 436 (99%) | 6 (1%) | 62 | 79 |
| 1 | B1 | 442/443 (100%) | 436 (99%) | 6 (1%) | 62 | 79 |
| 1 | B2 | 442/443 (100%) | 439 (99%) | 3 (1%) | 81 | 89 |

*Continued on next page...*

For Manuscript Review

*Continued from previous page...*

| Mol | Chain | Analysed | Rotameric | Outliers | Percentiles | |
|---|---|---|---|---|---|---|
| 1 | B3 | 442/443 (100%) | 438 (99%) | 4 (1%) | 75 | 86 |
| 1 | X1 | 442/443 (100%) | 437 (99%) | 5 (1%) | 70 | 83 |
| 1 | X2 | 442/443 (100%) | 438 (99%) | 4 (1%) | 75 | 86 |
| 1 | X3 | 442/443 (100%) | 437 (99%) | 5 (1%) | 70 | 83 |
| 1 | c1 | 441/443 (100%) | 438 (99%) | 3 (1%) | 81 | 89 |
| 1 | c2 | 442/443 (100%) | 441 (100%) | 1 (0%) | 92 | 95 |
| 1 | c3 | 442/443 (100%) | 439 (99%) | 3 (1%) | 81 | 89 |
| 1 | d1 | 442/443 (100%) | 437 (99%) | 5 (1%) | 70 | 83 |
| 1 | d2 | 442/443 (100%) | 438 (99%) | 4 (1%) | 75 | 86 |
| 1 | d3 | 442/443 (100%) | 437 (99%) | 5 (1%) | 70 | 83 |
| 1 | e1 | 442/443 (100%) | 437 (99%) | 5 (1%) | 70 | 83 |
| 1 | e2 | 442/443 (100%) | 441 (100%) | 1 (0%) | 92 | 95 |
| 1 | e3 | 442/443 (100%) | 438 (99%) | 4 (1%) | 75 | 86 |
| 1 | f1 | 442/443 (100%) | 436 (99%) | 6 (1%) | 62 | 79 |
| 1 | f2 | 442/443 (100%) | 440 (100%) | 2 (0%) | 86 | 91 |
| 1 | f3 | 442/443 (100%) | 439 (99%) | 3 (1%) | 81 | 89 |
| 2 | vC | 21/110 (19%) | 21 (100%) | 0 | 100 | 100 |
| 3 | wD | 88/209 (42%) | 84 (96%) | 4 (4%) | 23 | 50 |
| 4 | 57 | 110/268 (41%) | 109 (99%) | 1 (1%) | 75 | 86 |
| 5 | TP | 78/536 (15%) | 78 (100%) | 0 | 100 | 100 |
| 5 | TQ | 64/536 (12%) | 64 (100%) | 0 | 100 | 100 |
| 6 | ze | 53/151 (35%) | 53 (100%) | 0 | 100 | 100 |
| 7 | he | 127/159 (80%) | 127 (100%) | 0 | 100 | 100 |
| 8 | gc | 259/311 (83%) | 257 (99%) | 2 (1%) | 79 | 88 |
| 9 | uB | 258/258 (100%) | 256 (99%) | 2 (1%) | 79 | 88 |
| 10 | 76 | 130/198 (66%) | 126 (97%) | 4 (3%) | 35 | 60 |
| All | All | 10469/12039 (87%) | 10375 (99%) | 94 (1%) | 74 | 86 |

All (94) residues with a non-rotameric sidechain are listed below:

| Mol | Chain | Res | Type |
|---|---|---|---|
| 1 | B2 | 54 | ARG |
| 1 | B2 | 174 | CYS |

*Continued on next page...*

For Manuscript Review

Continued from previous page...

| Mol | Chain | Res | Type |
|---|---|---|---|
| 1 | B2 | 368 | PHE |
| 1 | B1 | 167 | VAL |
| 1 | B1 | 174 | CYS |
| 1 | B1 | 196 | ASN |
| 1 | B1 | 207 | GLN |
| 1 | B1 | 216 | ASP |
| 1 | B1 | 250 | LEU |
| 1 | B3 | 258 | ASP |
| 1 | B3 | 282 | ASP |
| 1 | B3 | 297 | ASN |
| 1 | B3 | 388 | GLU |
| 1 | c2 | 368 | PHE |
| 1 | c1 | 368 | PHE |
| 1 | c1 | 370 | GLN |
| 1 | c1 | 413 | CYS |
| 1 | c3 | 109 | VAL |
| 1 | c3 | 368 | PHE |
| 1 | c3 | 441 | ASP |
| 1 | d2 | 74 | VAL |
| 1 | d2 | 165 | CYS |
| 1 | d2 | 282 | ASP |
| 1 | d2 | 370 | GLN |
| 1 | d1 | 14 | GLN |
| 1 | d1 | 174 | CYS |
| 1 | d1 | 368 | PHE |
| 1 | d1 | 493 | THR |
| 1 | d1 | 525 | ASN |
| 1 | d3 | 131 | THR |
| 1 | d3 | 174 | CYS |
| 1 | d3 | 316 | CYS |
| 1 | d3 | 413 | CYS |
| 1 | d3 | 421 | VAL |
| 1 | f2 | 174 | CYS |
| 1 | f2 | 368 | PHE |
| 1 | f1 | 54 | ARG |
| 1 | f1 | 114 | GLN |
| 1 | f1 | 174 | CYS |
| 1 | f1 | 216 | ASP |
| 1 | f1 | 421 | VAL |
| 1 | f1 | 481 | ASP |
| 1 | f3 | 14 | GLN |
| 1 | f3 | 209 | HIS |

Continued on next page...

For Manuscript Review

*Continued from previous page...*

| Mol | Chain | Res | Type |
|---|---|---|---|
| 1 | f3 | 216 | ASP |
| 1 | X2 | 29 | VAL |
| 1 | X2 | 257 | LEU |
| 1 | X2 | 282 | ASP |
| 1 | X2 | 370 | GLN |
| 1 | X1 | 14 | GLN |
| 1 | X1 | 24 | ILE |
| 1 | X1 | 76 | LEU |
| 1 | X1 | 174 | CYS |
| 1 | X1 | 271 | GLU |
| 1 | X3 | 5 | LEU |
| 1 | X3 | 161 | VAL |
| 1 | X3 | 282 | ASP |
| 1 | X3 | 380 | THR |
| 1 | X3 | 436 | GLU |
| 1 | A2 | 282 | ASP |
| 1 | A2 | 320 | GLN |
| 1 | A1 | 174 | CYS |
| 1 | A1 | 210 | GLU |
| 1 | A1 | 249 | SER |
| 1 | A1 | 413 | CYS |
| 1 | A3 | 201 | LEU |
| 1 | A3 | 250 | LEU |
| 1 | A3 | 282 | ASP |
| 1 | A3 | 289 | SER |
| 1 | A3 | 421 | VAL |
| 1 | A3 | 476 | ASN |
| 1 | e2 | 174 | CYS |
| 1 | e1 | 114 | GLN |
| 1 | e1 | 174 | CYS |
| 1 | e1 | 368 | PHE |
| 1 | e1 | 391 | THR |
| 1 | e1 | 510 | ASN |
| 1 | e3 | 18 | LEU |
| 1 | e3 | 56 | VAL |
| 1 | e3 | 107 | VAL |
| 1 | e3 | 525 | ASN |
| 3 | wD | 28 | ASN |
| 3 | wD | 36 | LYS |
| 3 | wD | 95 | THR |
| 3 | wD | 97 | ILE |
| 4 | 57 | 162 | LEU |

*Continued on next page...*

For Manuscript Review

*Continued from previous page...*

| Mol | Chain | Res | Type |
|---|---|---|---|
| 8 | gc | 278 | THR |
| 8 | gc | 306 | VAL |
| 9 | uB | 52 | ASN |
| 9 | uB | 92 | LEU |
| 10 | 76 | 168 | TYR |
| 10 | 76 | 185 | LEU |
| 10 | 76 | 190 | ILE |
| 10 | 76 | 191 | VAL |

Sometimes sidechains can be flipped to improve hydrogen bonding and reduce clashes. All (333) such sidechains are listed below:

| Mol | Chain | Res | Type |
|---|---|---|---|
| 1 | B2 | 14 | GLN |
| 1 | B2 | 23 | GLN |
| 1 | B2 | 47 | ASN |
| 1 | B2 | 207 | GLN |
| 1 | B2 | 268 | ASN |
| 1 | B2 | 276 | GLN |
| 1 | B2 | 295 | ASN |
| 1 | B2 | 320 | GLN |
| 1 | B2 | 326 | ASN |
| 1 | B2 | 355 | GLN |
| 1 | B2 | 370 | GLN |
| 1 | B2 | 375 | ASN |
| 1 | B2 | 425 | GLN |
| 1 | B2 | 447 | GLN |
| 1 | B2 | 486 | GLN |
| 1 | B2 | 508 | ASN |
| 1 | B1 | 34 | HIS |
| 1 | B1 | 44 | GLN |
| 1 | B1 | 47 | ASN |
| 1 | B1 | 63 | ASN |
| 1 | B1 | 106 | GLN |
| 1 | B1 | 114 | GLN |
| 1 | B1 | 129 | GLN |
| 1 | B1 | 136 | GLN |
| 1 | B1 | 149 | GLN |
| 1 | B1 | 196 | ASN |
| 1 | B1 | 219 | ASN |
| 1 | B1 | 267 | GLN |
| 1 | B1 | 290 | ASN |
| 1 | B1 | 355 | GLN |

*Continued on next page...*

For Manuscript Review

*Continued from previous page...*

| Mol | Chain | Res | Type |
|---|---|---|---|
| 1 | B1 | 412 | HIS |
| 1 | B1 | 425 | GLN |
| 1 | B1 | 486 | GLN |
| 1 | B3 | 14 | GLN |
| 1 | B3 | 34 | HIS |
| 1 | B3 | 44 | GLN |
| 1 | B3 | 63 | ASN |
| 1 | B3 | 106 | GLN |
| 1 | B3 | 149 | GLN |
| 1 | B3 | 207 | GLN |
| 1 | B3 | 219 | ASN |
| 1 | B3 | 276 | GLN |
| 1 | B3 | 320 | GLN |
| 1 | B3 | 444 | GLN |
| 1 | B3 | 447 | GLN |
| 1 | B3 | 452 | ASN |
| 1 | B3 | 491 | ASN |
| 1 | B3 | 508 | ASN |
| 1 | c2 | 14 | GLN |
| 1 | c2 | 23 | GLN |
| 1 | c2 | 44 | GLN |
| 1 | c2 | 47 | ASN |
| 1 | c2 | 63 | ASN |
| 1 | c2 | 73 | GLN |
| 1 | c2 | 106 | GLN |
| 1 | c2 | 129 | GLN |
| 1 | c2 | 268 | ASN |
| 1 | c2 | 320 | GLN |
| 1 | c2 | 323 | GLN |
| 1 | c2 | 326 | ASN |
| 1 | c2 | 355 | GLN |
| 1 | c2 | 425 | GLN |
| 1 | c2 | 447 | GLN |
| 1 | c2 | 508 | ASN |
| 1 | c1 | 44 | GLN |
| 1 | c1 | 106 | GLN |
| 1 | c1 | 129 | GLN |
| 1 | c1 | 136 | GLN |
| 1 | c1 | 219 | ASN |
| 1 | c1 | 267 | GLN |
| 1 | c1 | 268 | ASN |
| 1 | c1 | 290 | ASN |

*Continued on next page...*

For Manuscript Review

*Continued from previous page...*

| Mol | Chain | Res | Type |
|---|---|---|---|
| 1 | c1 | 370 | GLN |
| 1 | c1 | 412 | HIS |
| 1 | c1 | 452 | ASN |
| 1 | c1 | 491 | ASN |
| 1 | c1 | 525 | ASN |
| 1 | c3 | 44 | GLN |
| 1 | c3 | 49 | GLN |
| 1 | c3 | 207 | GLN |
| 1 | c3 | 268 | ASN |
| 1 | c3 | 298 | HIS |
| 1 | c3 | 326 | ASN |
| 1 | c3 | 355 | GLN |
| 1 | c3 | 370 | GLN |
| 1 | c3 | 427 | ASN |
| 1 | c3 | 482 | ASN |
| 1 | c3 | 508 | ASN |
| 1 | c3 | 525 | ASN |
| 1 | d2 | 44 | GLN |
| 1 | d2 | 73 | GLN |
| 1 | d2 | 106 | GLN |
| 1 | d2 | 114 | GLN |
| 1 | d2 | 268 | ASN |
| 1 | d2 | 276 | GLN |
| 1 | d2 | 312 | ASN |
| 1 | d2 | 335 | ASN |
| 1 | d2 | 447 | GLN |
| 1 | d2 | 452 | ASN |
| 1 | d2 | 491 | ASN |
| 1 | d2 | 495 | GLN |
| 1 | d2 | 508 | ASN |
| 1 | d1 | 14 | GLN |
| 1 | d1 | 44 | GLN |
| 1 | d1 | 47 | ASN |
| 1 | d1 | 136 | GLN |
| 1 | d1 | 149 | GLN |
| 1 | d1 | 209 | HIS |
| 1 | d1 | 219 | ASN |
| 1 | d1 | 243 | GLN |
| 1 | d1 | 267 | GLN |
| 1 | d1 | 276 | GLN |
| 1 | d1 | 308 | GLN |
| 1 | d1 | 355 | GLN |

*Continued on next page...*

For Manuscript Review

*Continued from previous page...*

| Mol | Chain | Res | Type |
|---|---|---|---|
| 1 | d1 | 412 | HIS |
| 1 | d1 | 427 | ASN |
| 1 | d1 | 444 | GLN |
| 1 | d1 | 448 | HIS |
| 1 | d1 | 452 | ASN |
| 1 | d1 | 482 | ASN |
| 1 | d1 | 486 | GLN |
| 1 | d3 | 44 | GLN |
| 1 | d3 | 114 | GLN |
| 1 | d3 | 207 | GLN |
| 1 | d3 | 276 | GLN |
| 1 | d3 | 278 | GLN |
| 1 | d3 | 298 | HIS |
| 1 | d3 | 320 | GLN |
| 1 | d3 | 326 | ASN |
| 1 | d3 | 425 | GLN |
| 1 | d3 | 427 | ASN |
| 1 | d3 | 444 | GLN |
| 1 | d3 | 476 | ASN |
| 1 | d3 | 495 | GLN |
| 1 | d3 | 508 | ASN |
| 1 | f2 | 23 | GLN |
| 1 | f2 | 44 | GLN |
| 1 | f2 | 49 | GLN |
| 1 | f2 | 106 | GLN |
| 1 | f2 | 114 | GLN |
| 1 | f2 | 125 | HIS |
| 1 | f2 | 276 | GLN |
| 1 | f2 | 412 | HIS |
| 1 | f2 | 429 | GLN |
| 1 | f2 | 447 | GLN |
| 1 | f2 | 452 | ASN |
| 1 | f2 | 482 | ASN |
| 1 | f2 | 508 | ASN |
| 1 | f1 | 34 | HIS |
| 1 | f1 | 44 | GLN |
| 1 | f1 | 47 | ASN |
| 1 | f1 | 129 | GLN |
| 1 | f1 | 136 | GLN |
| 1 | f1 | 267 | GLN |
| 1 | f1 | 355 | GLN |
| 1 | f1 | 447 | GLN |

*Continued on next page...*

For Manuscript Review

*Continued from previous page...*

| Mol | Chain | Res | Type |
|---|---|---|---|
| 1 | f1 | 525 | ASN |
| 1 | f3 | 14 | GLN |
| 1 | f3 | 34 | HIS |
| 1 | f3 | 44 | GLN |
| 1 | f3 | 125 | HIS |
| 1 | f3 | 190 | GLN |
| 1 | f3 | 320 | GLN |
| 1 | f3 | 444 | GLN |
| 1 | f3 | 447 | GLN |
| 1 | f3 | 508 | ASN |
| 1 | X2 | 44 | GLN |
| 1 | X2 | 63 | ASN |
| 1 | X2 | 129 | GLN |
| 1 | X2 | 146 | ASN |
| 1 | X2 | 149 | GLN |
| 1 | X2 | 268 | ASN |
| 1 | X2 | 276 | GLN |
| 1 | X2 | 298 | HIS |
| 1 | X2 | 370 | GLN |
| 1 | X2 | 412 | HIS |
| 1 | X2 | 427 | ASN |
| 1 | X2 | 447 | GLN |
| 1 | X2 | 452 | ASN |
| 1 | X2 | 495 | GLN |
| 1 | X1 | 73 | GLN |
| 1 | X1 | 114 | GLN |
| 1 | X1 | 207 | GLN |
| 1 | X1 | 267 | GLN |
| 1 | X1 | 326 | ASN |
| 1 | X1 | 355 | GLN |
| 1 | X1 | 412 | HIS |
| 1 | X1 | 491 | ASN |
| 1 | X3 | 34 | HIS |
| 1 | X3 | 44 | GLN |
| 1 | X3 | 106 | GLN |
| 1 | X3 | 125 | HIS |
| 1 | X3 | 136 | GLN |
| 1 | X3 | 146 | ASN |
| 1 | X3 | 190 | GLN |
| 1 | X3 | 207 | GLN |
| 1 | X3 | 214 | ASN |
| 1 | X3 | 219 | ASN |

*Continued on next page...*

For Manuscript Review

*Continued from previous page...*

| Mol | Chain | Res | Type |
|---|---|---|---|
| 1 | X3 | 276 | GLN |
| 1 | X3 | 297 | ASN |
| 1 | X3 | 298 | HIS |
| 1 | X3 | 320 | GLN |
| 1 | X3 | 444 | GLN |
| 1 | X3 | 447 | GLN |
| 1 | X3 | 470 | GLN |
| 1 | X3 | 482 | ASN |
| 1 | X3 | 486 | GLN |
| 1 | X3 | 508 | ASN |
| 1 | X3 | 525 | ASN |
| 1 | A2 | 23 | GLN |
| 1 | A2 | 34 | HIS |
| 1 | A2 | 44 | GLN |
| 1 | A2 | 47 | ASN |
| 1 | A2 | 63 | ASN |
| 1 | A2 | 106 | GLN |
| 1 | A2 | 114 | GLN |
| 1 | A2 | 119 | GLN |
| 1 | A2 | 243 | GLN |
| 1 | A2 | 268 | ASN |
| 1 | A2 | 295 | ASN |
| 1 | A2 | 323 | GLN |
| 1 | A2 | 370 | GLN |
| 1 | A2 | 412 | HIS |
| 1 | A2 | 447 | GLN |
| 1 | A2 | 482 | ASN |
| 1 | A2 | 495 | GLN |
| 1 | A2 | 508 | ASN |
| 1 | A1 | 23 | GLN |
| 1 | A1 | 49 | GLN |
| 1 | A1 | 114 | GLN |
| 1 | A1 | 119 | GLN |
| 1 | A1 | 129 | GLN |
| 1 | A1 | 149 | GLN |
| 1 | A1 | 196 | ASN |
| 1 | A1 | 267 | GLN |
| 1 | A1 | 276 | GLN |
| 1 | A1 | 320 | GLN |
| 1 | A3 | 196 | ASN |
| 1 | A3 | 278 | GLN |
| 1 | A3 | 320 | GLN |

*Continued on next page...*

For Manuscript Review

*Continued from previous page...*

| Mol | Chain | Res | Type |
|---|---|---|---|
| 1 | A3 | 427 | ASN |
| 1 | A3 | 447 | GLN |
| 1 | A3 | 476 | ASN |
| 1 | A3 | 495 | GLN |
| 1 | A3 | 508 | ASN |
| 1 | A3 | 525 | ASN |
| 1 | e2 | 23 | GLN |
| 1 | e2 | 34 | HIS |
| 1 | e2 | 129 | GLN |
| 1 | e2 | 233 | GLN |
| 1 | e2 | 278 | GLN |
| 1 | e2 | 297 | ASN |
| 1 | e2 | 298 | HIS |
| 1 | e2 | 312 | ASN |
| 1 | e2 | 355 | GLN |
| 1 | e2 | 412 | HIS |
| 1 | e2 | 429 | GLN |
| 1 | e2 | 447 | GLN |
| 1 | e2 | 508 | ASN |
| 1 | e1 | 14 | GLN |
| 1 | e1 | 44 | GLN |
| 1 | e1 | 47 | ASN |
| 1 | e1 | 63 | ASN |
| 1 | e1 | 106 | GLN |
| 1 | e1 | 129 | GLN |
| 1 | e1 | 136 | GLN |
| 1 | e1 | 219 | ASN |
| 1 | e1 | 312 | ASN |
| 1 | e1 | 320 | GLN |
| 1 | e1 | 355 | GLN |
| 1 | e1 | 412 | HIS |
| 1 | e1 | 482 | ASN |
| 1 | e1 | 510 | ASN |
| 1 | e3 | 106 | GLN |
| 1 | e3 | 149 | GLN |
| 1 | e3 | 196 | ASN |
| 1 | e3 | 209 | HIS |
| 1 | e3 | 267 | GLN |
| 1 | e3 | 268 | ASN |
| 1 | e3 | 295 | ASN |
| 1 | e3 | 297 | ASN |
| 1 | e3 | 312 | ASN |

*Continued on next page...*

For Manuscript Review

Continued from previous page...

| Mol | Chain | Res | Type |
|---|---|---|---|
| 1 | e3 | 323 | GLN |
| 1 | e3 | 326 | ASN |
| 1 | e3 | 355 | GLN |
| 1 | e3 | 425 | GLN |
| 1 | e3 | 427 | ASN |
| 1 | e3 | 429 | GLN |
| 1 | e3 | 447 | GLN |
| 1 | e3 | 476 | ASN |
| 1 | e3 | 491 | ASN |
| 1 | e3 | 495 | GLN |
| 3 | wD | 18 | ASN |
| 3 | wD | 51 | ASN |
| 3 | wD | 60 | ASN |
| 3 | wD | 67 | ASN |
| 3 | wD | 79 | ASN |
| 4 | 57 | 160 | ASN |
| 4 | 57 | 161 | ASN |
| 4 | 57 | 237 | GLN |
| 4 | 57 | 250 | GLN |
| 5 | TP | 175 | ASN |
| 5 | TP | 253 | ASN |
| 5 | TP | 260 | HIS |
| 6 | ze | 92 | ASN |
| 6 | ze | 140 | ASN |
| 7 | he | 106 | ASN |
| 7 | he | 125 | ASN |
| 7 | he | 144 | HIS |
| 8 | gc | 109 | HIS |
| 8 | gc | 209 | ASN |
| 8 | gc | 219 | HIS |
| 8 | gc | 230 | ASN |
| 9 | uB | 10 | ASN |
| 9 | uB | 24 | ASN |
| 9 | uB | 37 | ASN |
| 9 | uB | 72 | GLN |
| 9 | uB | 91 | ASN |
| 9 | uB | 112 | ASN |
| 9 | uB | 117 | ASN |
| 9 | uB | 130 | ASN |
| 9 | uB | 166 | GLN |
| 9 | uB | 187 | ASN |
| 10 | 76 | 96 | ASN |

Continued on next page...

For Manuscript Review

*Continued from previous page...*

| Mol | Chain | Res | Type |
|---|---|---|---|
| 10 | 76 | 102 | GLN |
| 10 | 76 | 122 | ASN |
| 10 | 76 | 127 | ASN |
| 10 | 76 | 178 | GLN |
| 5 | TQ | 111 | ASN |
| 5 | TQ | 115 | HIS |
| 5 | TQ | 142 | ASN |
| 5 | TQ | 144 | GLN |
| 5 | TQ | 156 | GLN |

##### 5.3.3 RNA (i)

There are no RNA molecules in this entry.

#### 5.4 Non-standard residues in protein, DNA, RNA chains (i)

There are no non-standard protein/DNA/RNA residues in this entry.

#### 5.5 Carbohydrates (i)

There are no oligosaccharides in this entry.

#### 5.6 Ligand geometry (i)

There are no ligands in this entry.

#### 5.7 Other polymers (i)

There are no such residues in this entry.

#### 5.8 Polymer linkage issues (i)

There are no chain breaks in this entry.

For Manuscript Review

### 6 Map visualisation (i)

This section contains visualisations of the EMDB entry EMD-55914. These allow visual inspection of the internal detail of the map and identification of artifacts.

Images derived from a raw map, generated by summing the deposited half-maps, are presented below the corresponding image components of the primary map to allow further visual inspection and comparison with those of the primary map.

#### 6.1 Orthogonal projections (i)

##### 6.1.1 Primary map

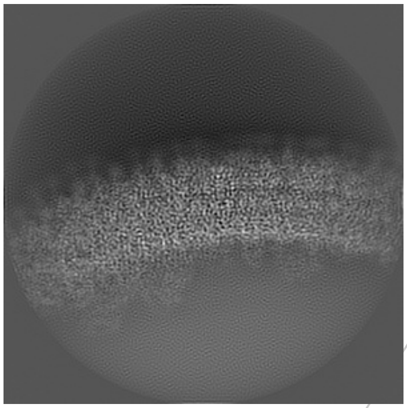
X

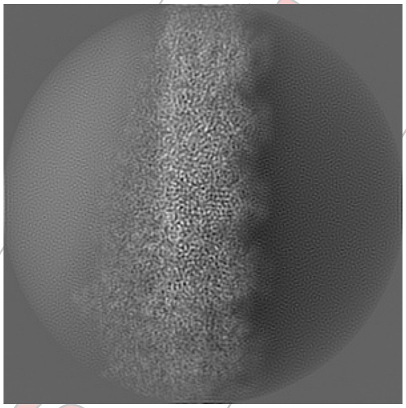
Y

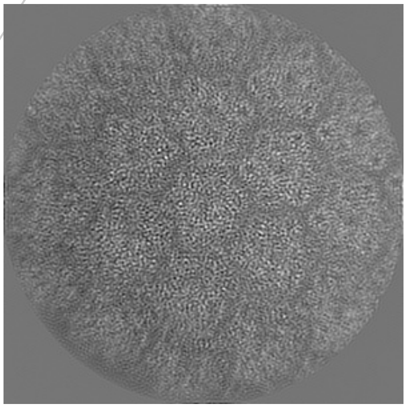
Z

##### 6.1.2 Raw map

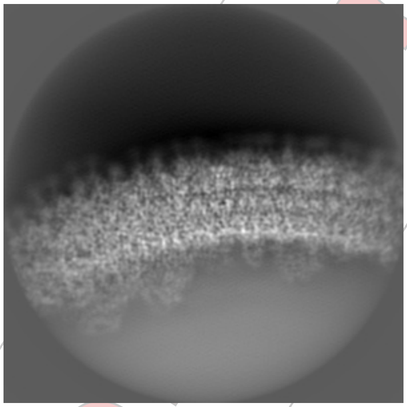
X

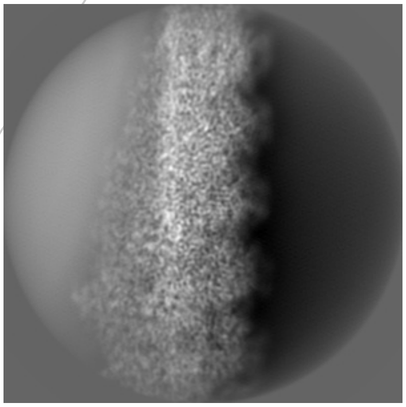
Y

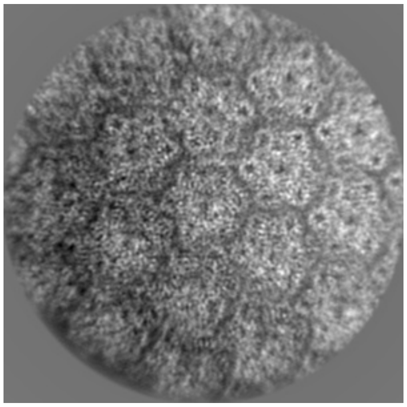
Z

The images above show the map projected in three orthogonal directions.

For Manuscript Review

#### 6.2 Central slices (i)

##### 6.2.1 Primary map

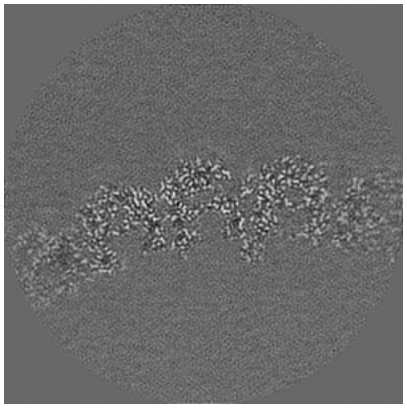
X Index: 128

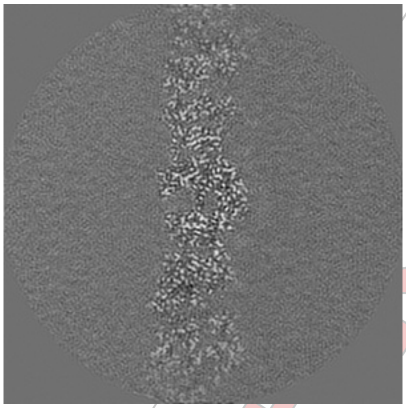
Y Index: 128

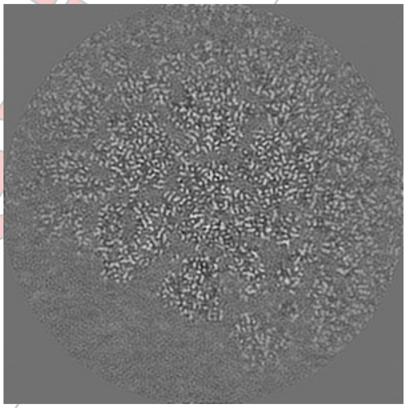
Z Index: 128

##### 6.2.2 Raw map

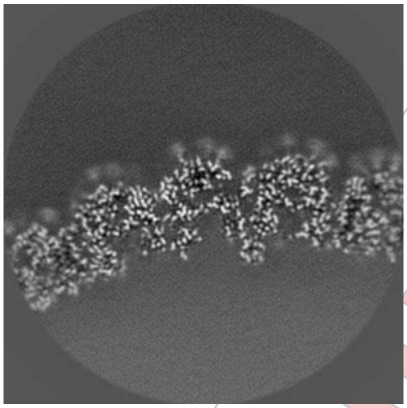
X Index: 128

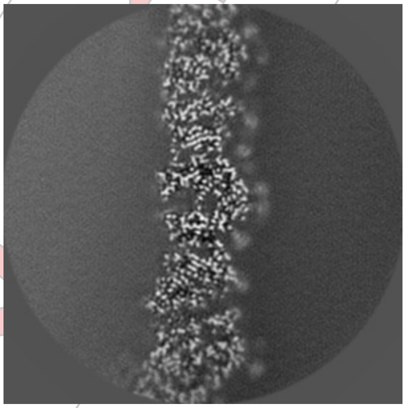
Y Index: 128

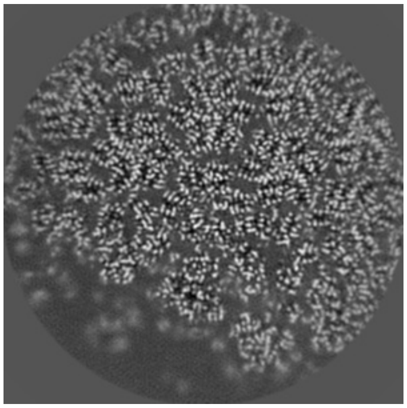
Z Index: 128

The images above show central slices of the map in three orthogonal directions.

For Manuscript Review

#### 6.3 Largest variance slices (i)

##### 6.3.1 Primary map

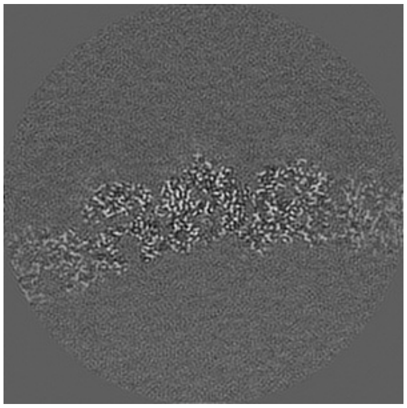
X Index: 122

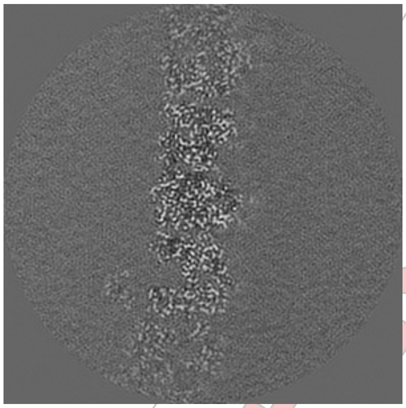
Y Index: 112

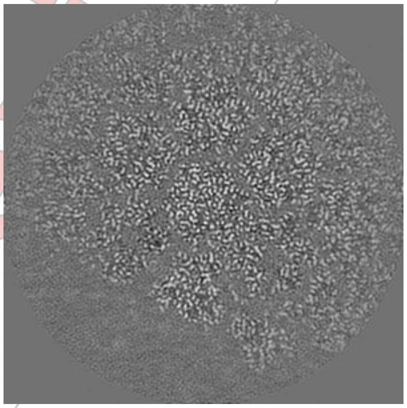
Z Index: 126

##### 6.3.2 Raw map

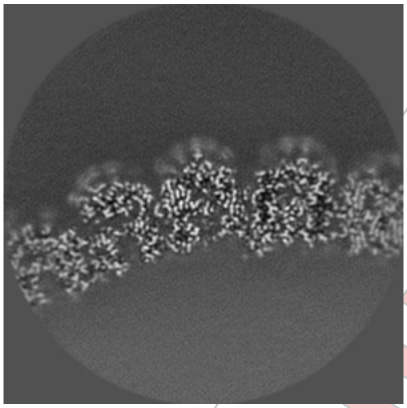
X Index: 122

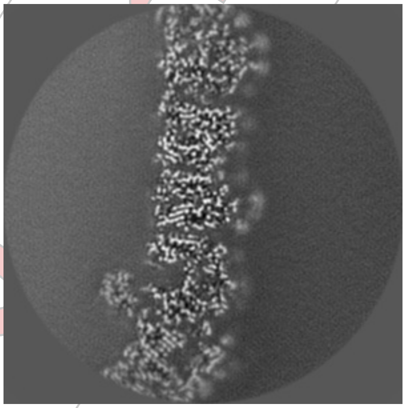
Y Index: 110

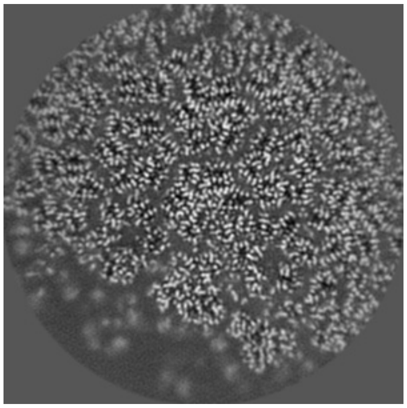
Z Index: 126

The images above show the largest variance slices of the map in three orthogonal directions.

For Manuscript Review

#### 6.4 Orthogonal standard-deviation projections (False-color) (i)

##### 6.4.1 Primary map

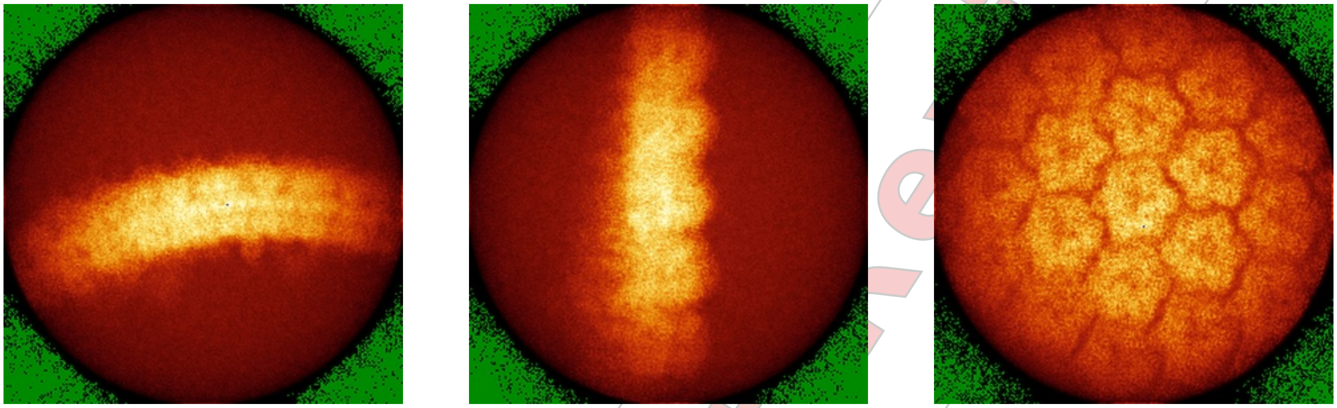

X Y Z

##### 6.4.2 Raw map

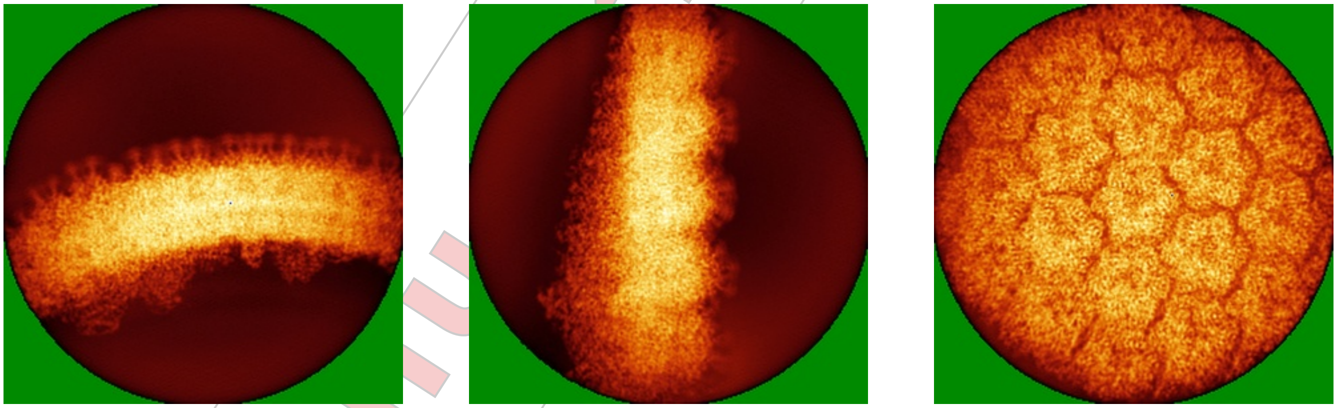

X Y Z

The images above show the map standard deviation projections with false color in three orthogonal directions. Minimum values are shown in green, max in blue, and dark to light orange shades represent small to large values respectively.

For Manuscript Review

#### 6.5 Orthogonal surface views (i)

##### 6.5.1 Primary map

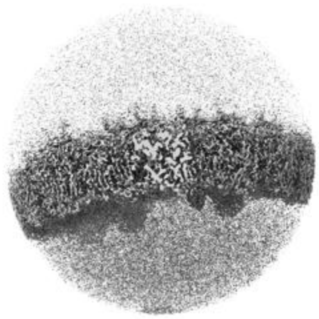

X

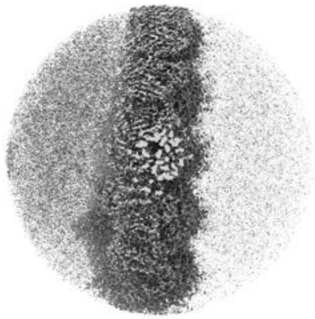

Y

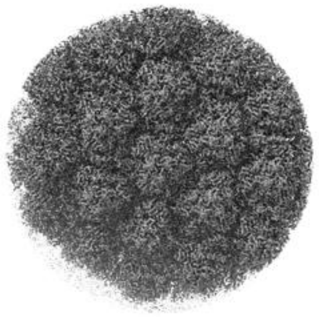

Z

The images above show the 3D surface view of the map at the recommended contour level 0.03. These images, in conjunction with the slice images, may facilitate assessment of whether an appropriate contour level has been provided.

##### 6.5.2 Raw map

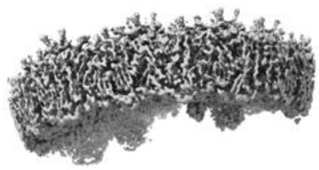

X

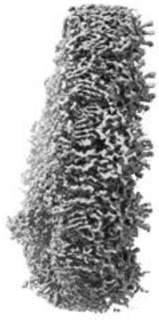

Y

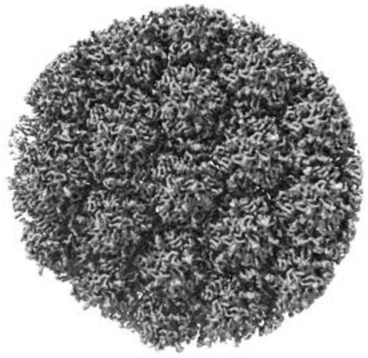

Z

These images show the 3D surface of the raw map. The raw map's contour level was selected so that its surface encloses the same volume as the primary map does at its recommended contour level.

#### 6.6 Mask visualisation (i)

This section was not generated. No masks/segmentation were deposited.

For Manuscript Review

### 7 Map analysis (i)

This section contains the results of statistical analysis of the map.

#### 7.1 Map-value distribution (i)

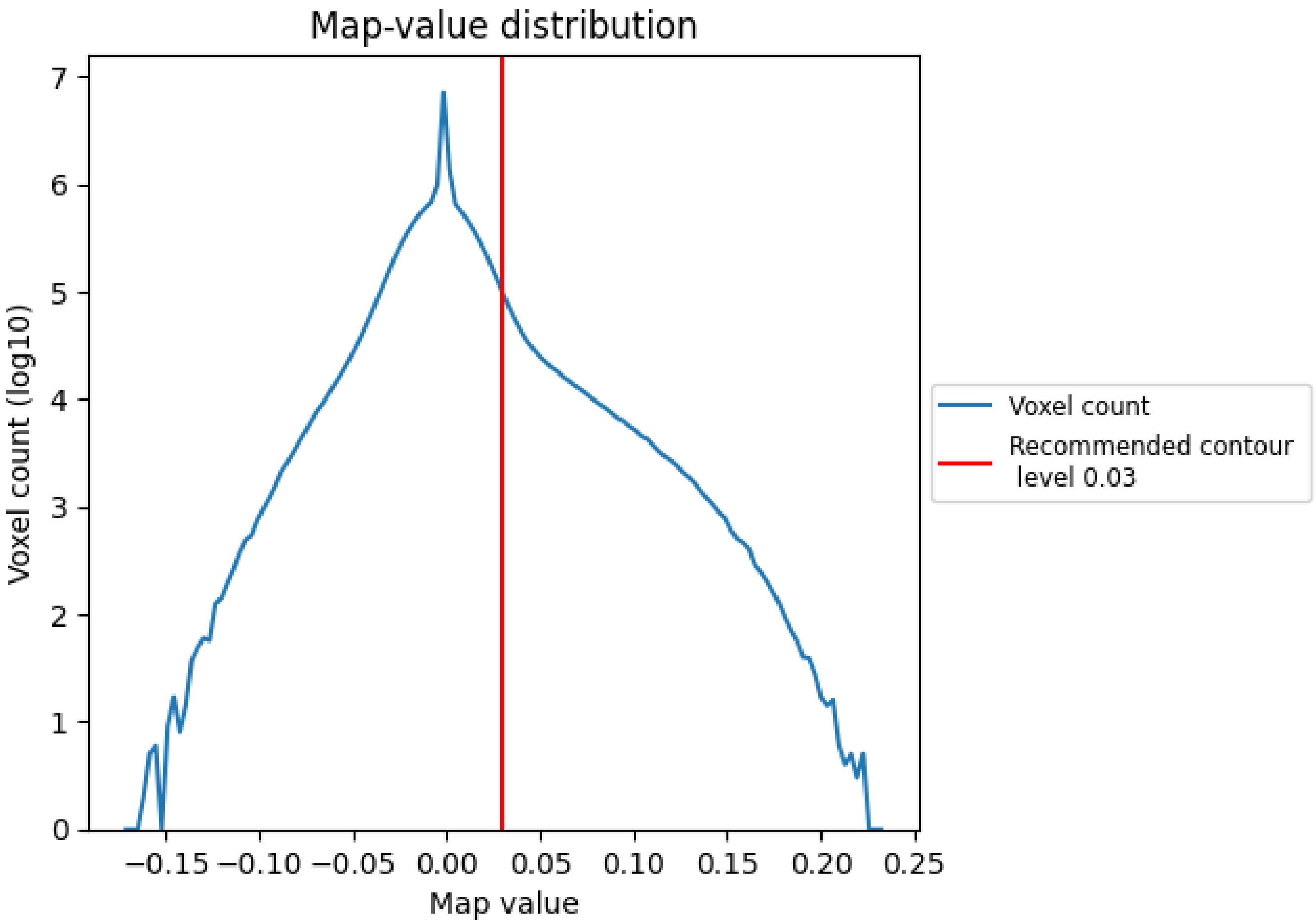


The map-value distribution is plotted in 128 intervals along the x-axis. The y-axis is logarithmic. A spike in this graph at zero usually indicates that the volume has been masked.

For Manuscript Review

##### 7.2 Volume estimate (i)

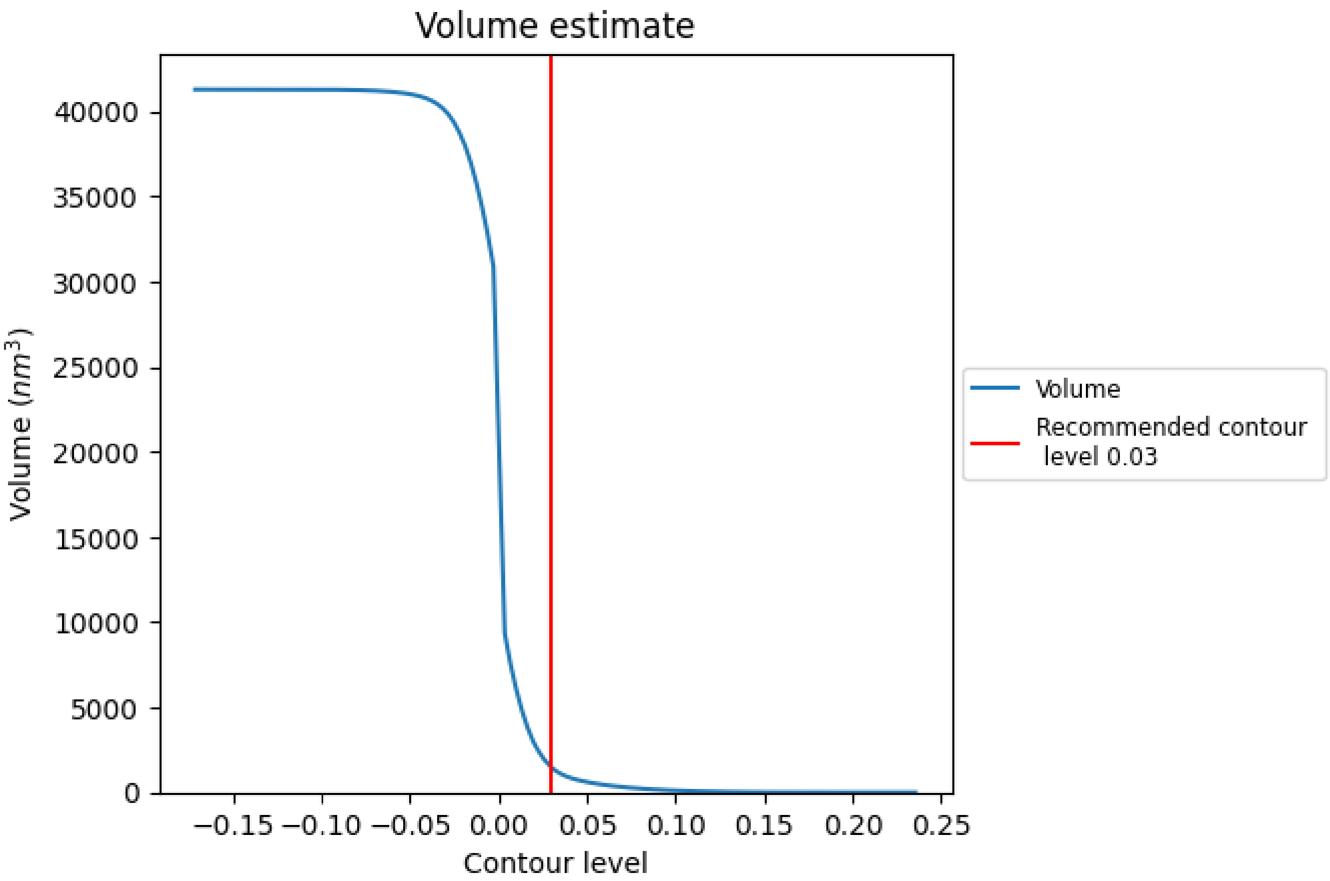


The volume at the recommended contour level is 1471 $nm^3$; this corresponds to an approximate mass of 1329 kDa.

The volume estimate graph shows how the enclosed volume varies with the contour level. The recommended contour level is shown as a vertical line and the intersection between the line and the curve gives the volume of the enclosed surface at the given level.

For Manuscript Review

#### 7.3 Rotationally averaged power spectrum (i)

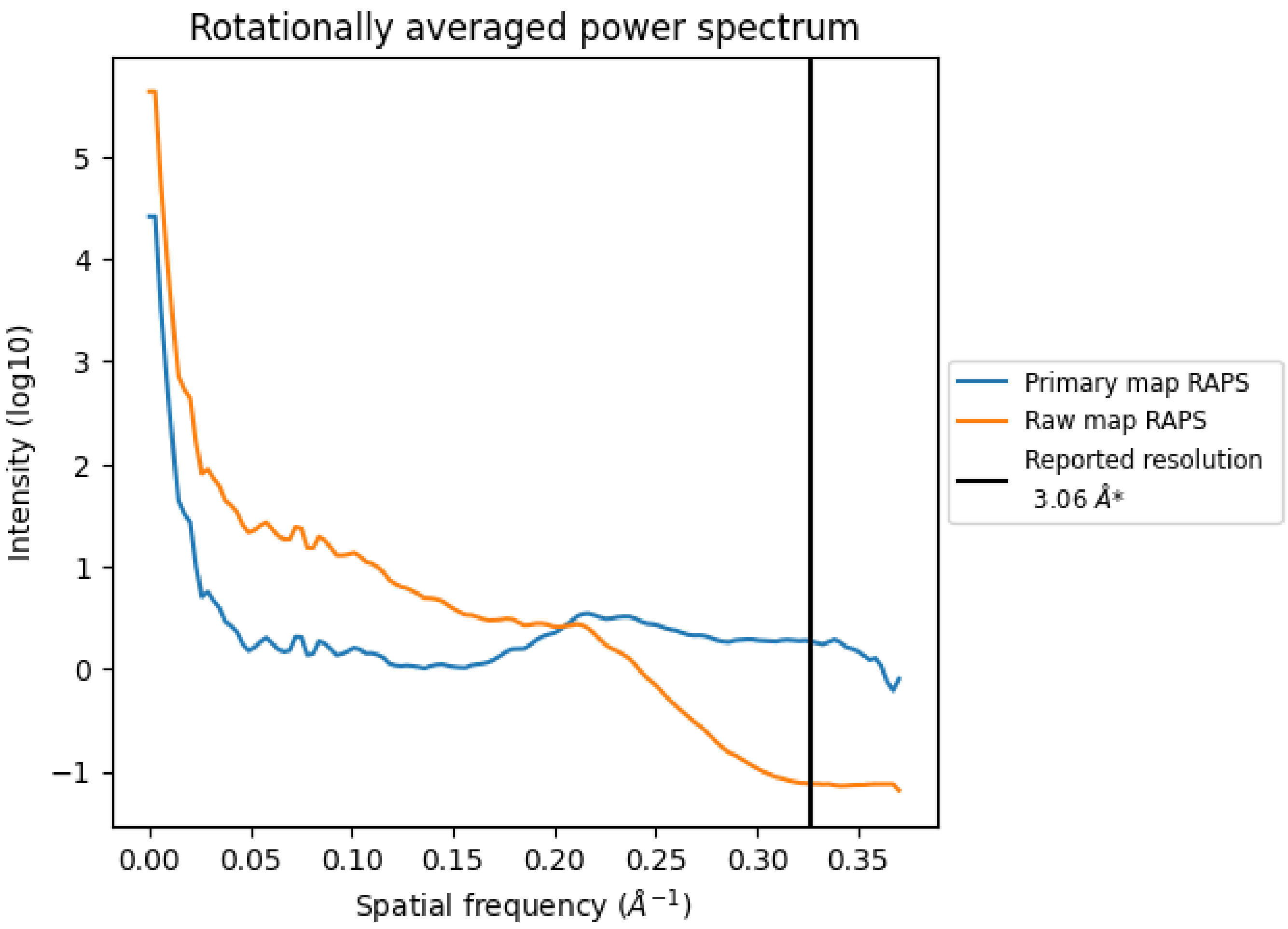


*Reported resolution corresponds to spatial frequency of 0.327 Å$^{-1}$

For Manuscript Review

### 8 Fourier-Shell correlation (i)

Fourier-Shell Correlation (FSC) is the most commonly used method to estimate the resolution of single-particle and subtomogram-averaged maps. The shape of the curve depends on the imposed symmetry, mask and whether or not the two 3D reconstructions used were processed from a common reference. The reported resolution is shown as a black line. A curve is displayed for the half-bit criterion in addition to lines showing the 0.143 gold standard cut-off and 0.5 cut-off.

#### 8.1 FSC (i)

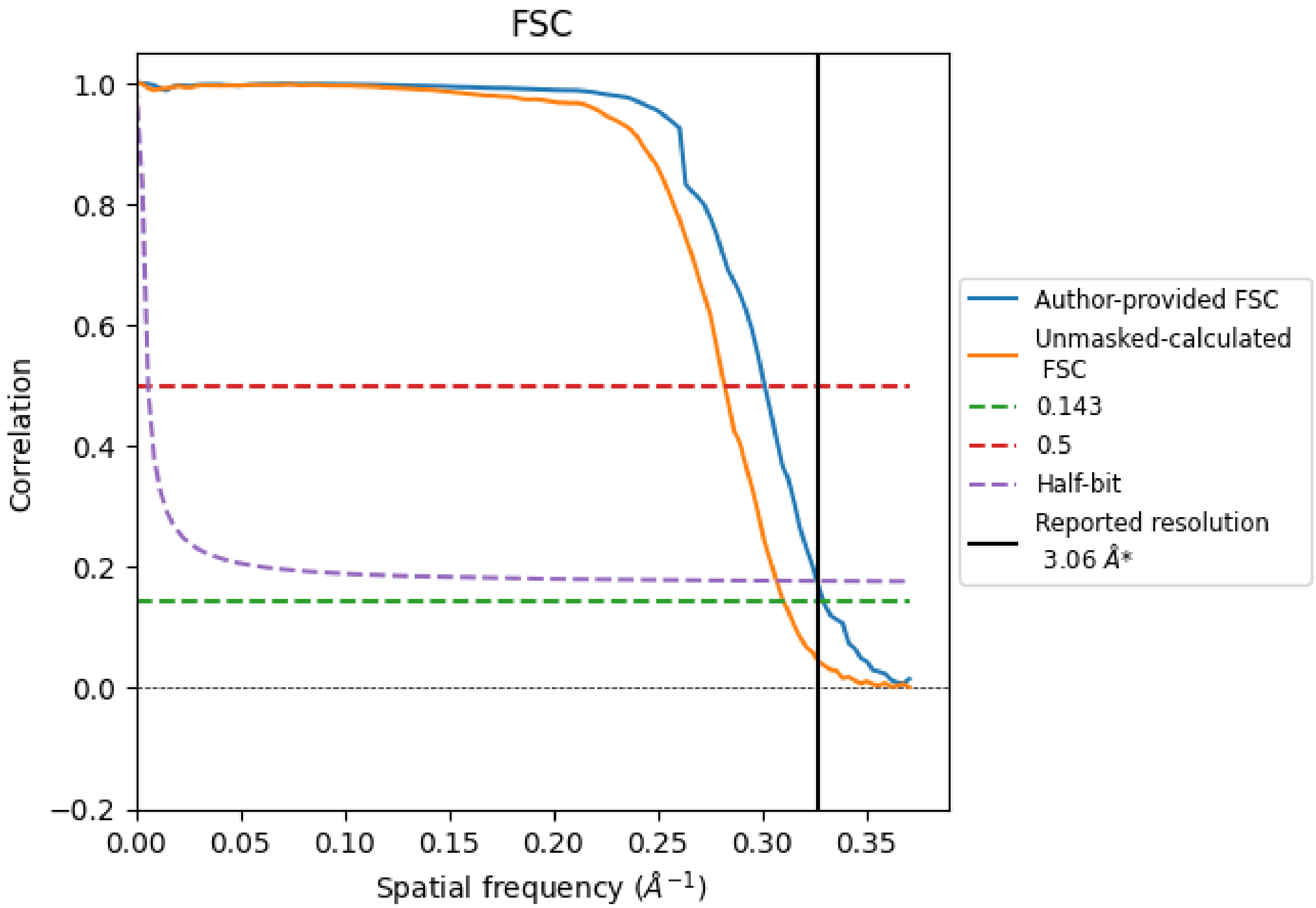


*Reported resolution corresponds to spatial frequency of 0.327 Å$^{-1}$

For Manuscript Review

#### 8.2 Resolution estimates (i)

| Resolution estimate (Å) | Estimation criterion (FSC cut-off) | | |
|---|---|---|---|
| | 0.143 | 0.5 | Half-bit |
| Reported by author | 3.06 | - | - |
| Author-provided FSC curve | 3.04 | 3.32 | 3.07 |
| Unmasked-calculated* | 3.22 | 3.55 | 3.26 |

*Resolution estimate based on FSC curve calculated by comparison of deposited half-maps.

For Manuscript Review

### 9 Map-model fit (i)

This section contains information regarding the fit between EMDB map EMD-55914 and PDB model 9TH1. Per-residue inclusion information can be found in section 3 on page 8.

#### 9.1 Map-model overlay (i)

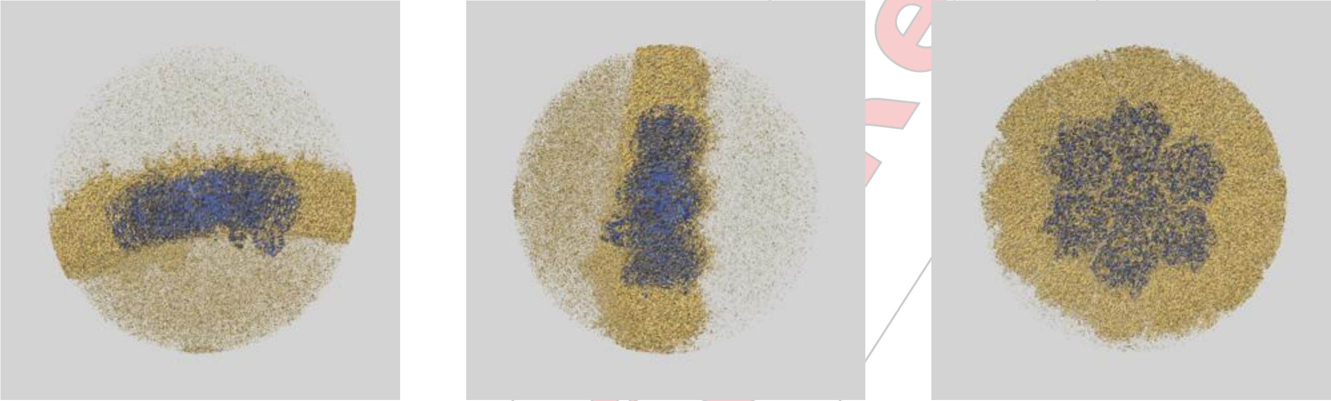

X Y Z

The images above show the 3D surface view of the map at the recommended contour level 0.03 at 50% transparency in yellow overlaid with a ribbon representation of the model coloured in blue. These images allow for the visual assessment of the quality of fit between the atomic model and the map.

For Manuscript Review

#### 9.2 Q-score mapped to coordinate model (i)

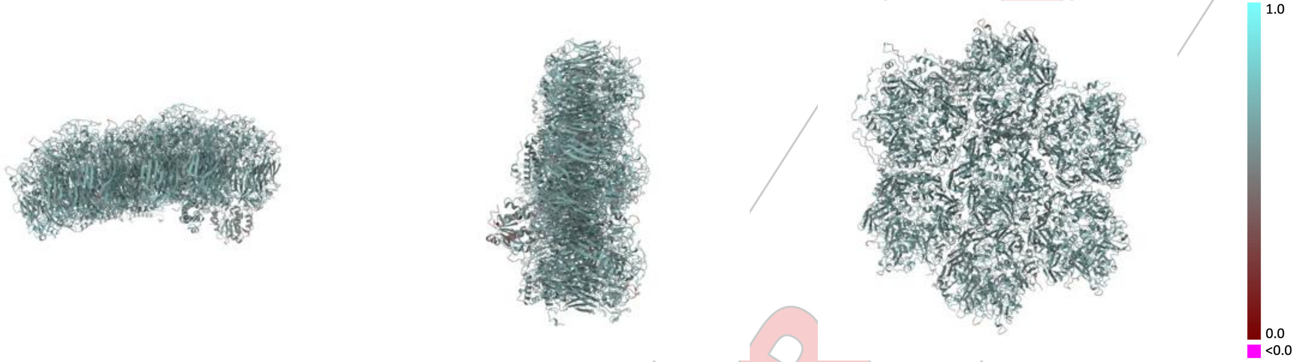


X Y Z

The images above show the model with each residue coloured according its Q-score. This shows their resolvability in the map with higher Q-score values reflecting better resolvability. Please note: Q-score is calculating the resolvability of atoms, and thus high values are only expected at resolutions at which atoms can be resolved. Low Q-score values may therefore be expected for many entries.

#### 9.3 Atom inclusion mapped to coordinate model (i)

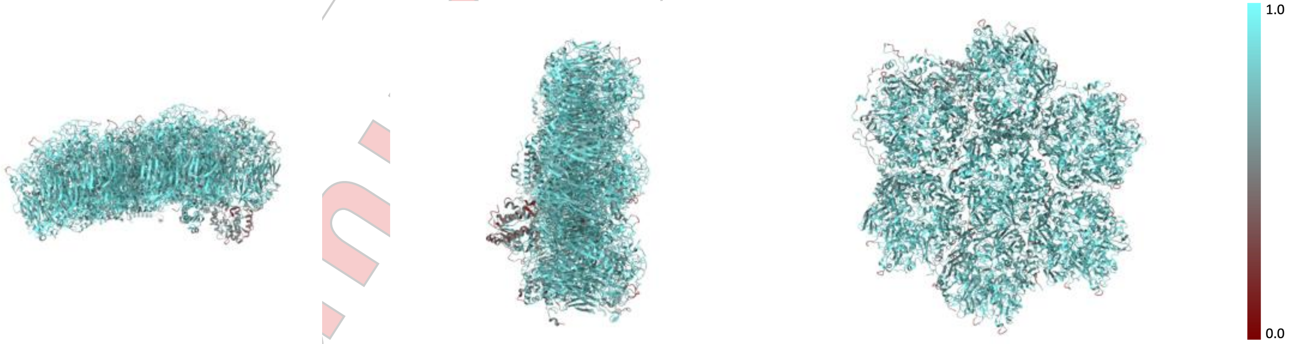


X Y Z

The images above show the model with each residue coloured according to its atom inclusion. This shows to what extent they are inside the map at the recommended contour level (0.03).

For Manuscript Review

#### 9.4 Atom inclusion (i)

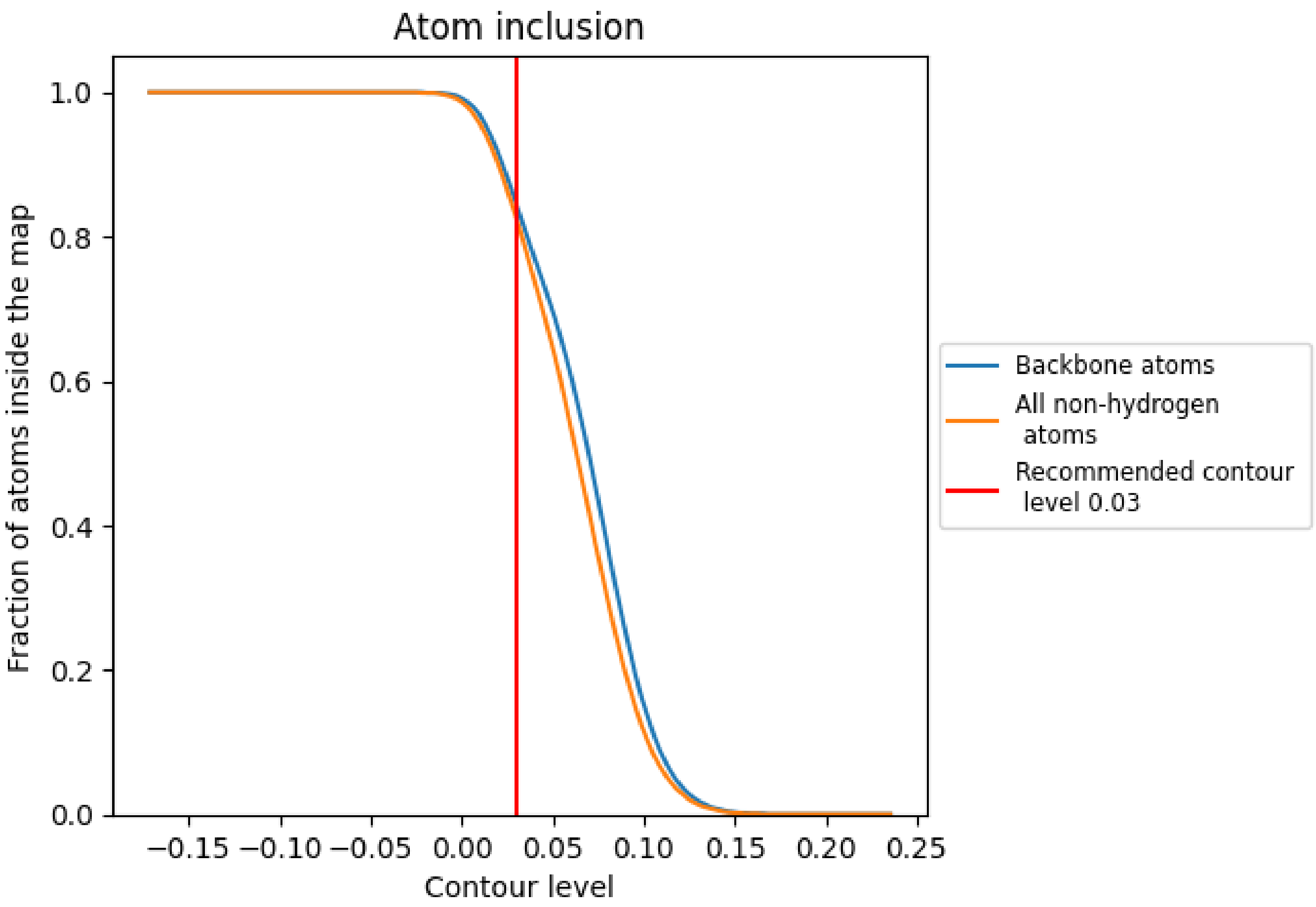


At the recommended contour level, 84% of all backbone atoms, 83% of all non-hydrogen atoms, are inside the map.

For Manuscript Review

#### 9.5 Map-model fit summary (i)

The table lists the average atom inclusion at the recommended contour level (0.03) and Q-score for the entire model and for each chain.

| Chain | Atom inclusion | Q-score |
|---|---|---|
| All | 0.8260 | 0.5980 |
| 57 | 0.7110 | 0.5620 |
| 76 | 0.6910 | 0.5570 |
| A1 | 0.8250 | 0.5940 |
| A2 | 0.8550 | 0.6050 |
| A3 | 0.8530 | 0.6080 |
| B1 | 0.8370 | 0.5980 |
| B2 | 0.8630 | 0.6080 |
| B3 | 0.8250 | 0.6000 |
| TP | 0.8350 | 0.6040 |
| TQ | 0.6570 | 0.5500 |
| X1 | 0.8240 | 0.5950 |
| X2 | 0.8270 | 0.5990 |
| X3 | 0.8550 | 0.6060 |
| c1 | 0.8310 | 0.5980 |
| c2 | 0.7920 | 0.5900 |
| c3 | 0.8390 | 0.6000 |
| d1 | 0.8340 | 0.5990 |
| d2 | 0.8210 | 0.5980 |
| d3 | 0.8570 | 0.6080 |
| e1 | 0.8830 | 0.6130 |
| e2 | 0.8830 | 0.6150 |
| e3 | 0.8810 | 0.6150 |
| f1 | 0.8490 | 0.6050 |
| f2 | 0.8010 | 0.5890 |
| f3 | 0.8120 | 0.5950 |
| gc | 0.5550 | 0.5310 |
| he | 0.7870 | 0.5800 |
| uB | 0.7760 | 0.5860 |
| vC | 0.6610 | 0.5480 |
| wD | 0.7700 | 0.5630 |
| ze | 0.7510 | 0.5770 |

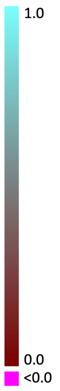


For Manuscript Review
